# Imminent invasion of the chytrid fungus threatens the last naïve amphibian biodiversity hotspots

**DOI:** 10.64898/2026.02.02.703360

**Authors:** Amadeus Plewnia, Frank Pasmans, Tobias Hildwein, Kevin P. Mulder, Jonas Henn, Andrew J. Crawford, Luis Alberto Rueda Solano, Nico Fuhrmann, Arved Lühmann, Jesse Erens, Sandra Victoria Flechas, Alessandro Catenazzi, Laura Victoria Rivera Jaimes, Victor L. N. Araújo, Claudia Lansac, Christopher Heine, Romario Salas, Aldair A. Barros-Granados, Alejandra Barrios Yepes, Laura V. Rengifo-Saavedra, Anna E. Savage, Valentina Vásquez, Jaime Culebras, Jose Daniel Barros Castañeda, José Luis Pérez-González, Sintana Rojas-Montaño, Jefferson Villalba, Juan M. Guayasamin, Sandra P. Galeano, Ben C. Scheele, Simon Clulow, An Martel, Stefan Lötters

## Abstract

While the amphibian chytrid fungus *Batrachochytrium dendrobatidis* (*Bd*) is driving catastrophic biodiversity loss worldwide, some amphibian communities persist seemingly unaffected despite occurring in climates conducive to pathogen establishment. These amphibian communities may remain epidemiologically naïve. As mitigation of *Bd* is rarely successful after establishment, identifying remaining *Bd*-free refuges is imperative. Presently, the only known large-scale *Bd*-free refuge is the island of New Guinea (NG), safeguarding Australasia’s amphibian phylogenetic diversity otherwise devastated by *Bd*. Following extensive multi-year disease surveillance, we here uncover a second large-scale *Bd-*free refuge in the Sierra Nevada de Santa Marta (SNSM), a Neotropical biodiversity hotspot in northern Colombia. We detected no evidence of *Bd* in SNSM-wide screening, while we uncovered the presence of hypervirulent *Bd*-GPL in adjacent areas of the tropical Andes. Population genomic analyses in an SNSM-endemic anuran found no evidence for demographic bottlenecks indicative of cryptic epizootic decline. Niche modelling highlights the high risk for *Bd* establishment and *Bd*-induced declines in both the SNSM and NG, and the important role of lowland environmental barriers in restricting *Bd* invasion. Infection trials using three SNSM-endemic amphibians reveal varying disease susceptibility. Together, these data identify the SNSM as an epidemiologically naïve refuge likely facing imminent *Bd* invasion, which could result in the loss of at least 25 endemic amphibian species. We highlight the urgent need for proactive conservation action and strict implementation of biosecurity to safeguard the unique and vast amphibian diversity of the world’s last major *Bd*-free refuges.

**Significance Statement:** Amphibian chytridiomycosis caused by *Batrachochytrium dendrobatidis* (*Bd*) has driven unprecedented global biodiversity loss. The Neotropics and Australasia comprise epicenters of declines. Identifying remaining *Bd*-free refuges is crucial to curb further amphibian extinctions, but so far contemporary absence of *Bd* has only been demonstrated for New Guinea. Here we identify the last known major *Bd*-free biodiversity hotspot in the Neotropics: the Sierra Nevada de Santa Marta (SNSM) in Colombia. Our results show that amphibian communities in this hotspot are immunologically naïve despite occurrence of hypervirulent *Bd* lineages nearby and climatic conditions within the SNSM conducive to *Bd*-induced declines. This creates an imminent risk for *Bd*-driven declines and highlights the urgent need for preventive actions to avert another wave of biodiversity loss.

## Introduction

The Anthropocene is characterized by the global emergence of infectious diseases, posing major challenges to human and wildlife health (1-3). Amphibian chytridiomycosis, caused by the fungal pathogen *Batrachochytrium dendrobatidis* (*Bd*), is considered the most devastating wildlife disease ever recorded (2, 4). Since the 1980s, it has resulted in catastrophic amphibian declines and extinctions worldwide (5, 6), further accelerating the amphibian extinction crisis (7). While the pathogen has been detected in virtually all regions surveyed, *Bd*-induced declines are distributed unequally across the globe (6, 8). Humid montane environments have emerged as prime landscapes for *Bd* epizootics (8, 9). Neotropical and Australian highlands stand out as epicenters in terms of the number of species affected and the spatial extent of community-wide declines (10). In both regions, *Bd* advanced across the landscape in a wavelike pattern from one or multiple introductions (Fig. 1A, B), with spatiotemporally structured near-extinction of once diverse and abundant amphibian communities (11-13).

**Figure 1.**
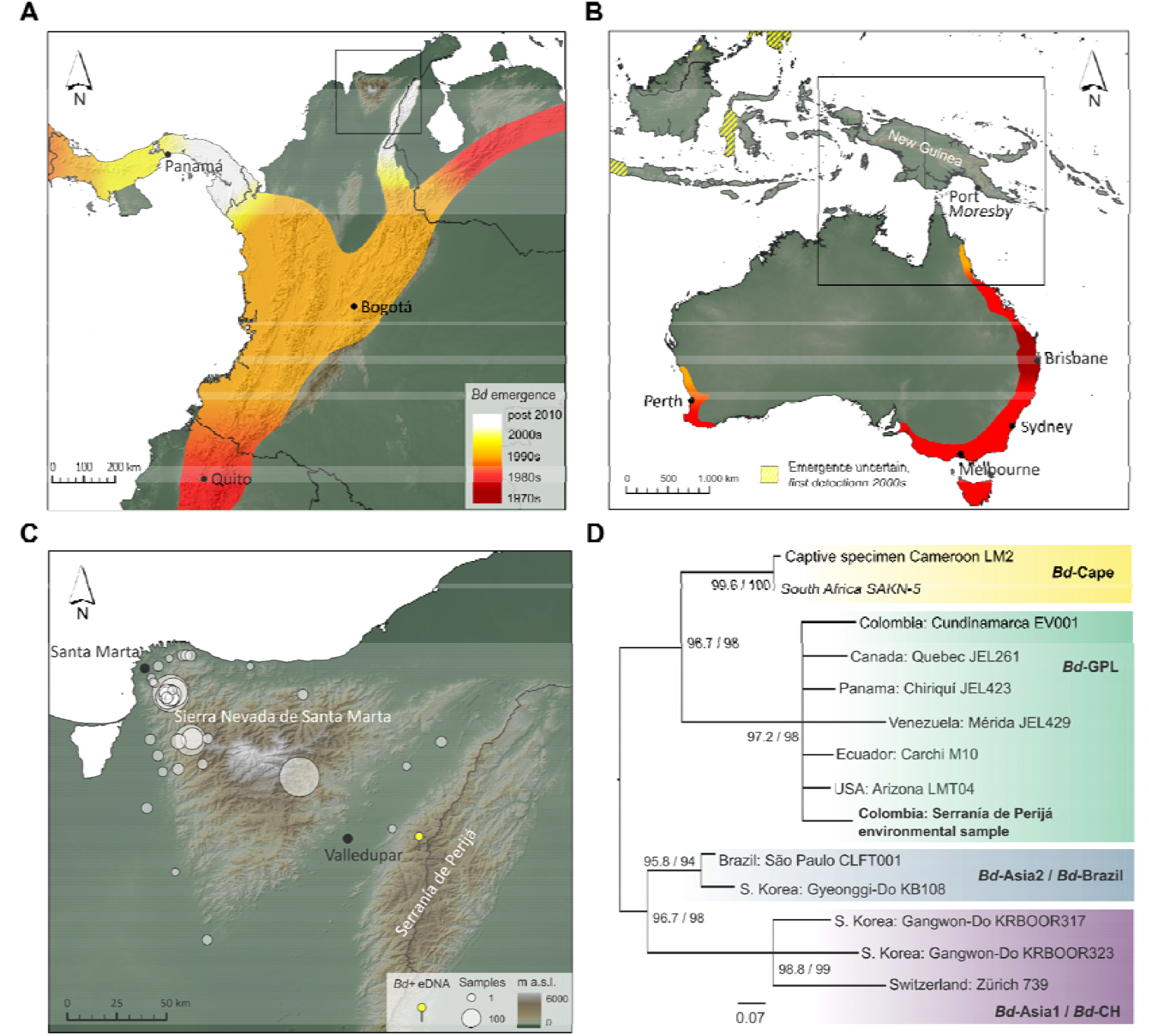
Invasion of *Batrachochytrium dendrobatidis* (*Bd*) in the global epicenters of amphibian declines. A) Hypothetical spatiotemporal spread of *Bd* in the northern Andes and adjacent Central America, approximated by *Bd* presence records and years when likely *Bd*-induced host declines were first observed. Information derived from (12, 31), updated with ‘year last seen’ of highly susceptible harlequin toads from (26) and additional records from (25) and this study. B) Hypothetical spatiotemporal spread in Australasia and adjacent SE Asia, derived from *Bd* presence data and temporal onset of host declines reported in (5, 13, 79). Note that *Bd* might be native in adjacent SE Asia, but surveys were only carried out post 2000. C) *Bd* sampling in the Sierra Nevada de Santa Marta and adjacent Colombian versant of the Serranía de Perijá (SP) along the Venezuelan border; gray circles are *Bd*-negative, the yellow dot is *Bd*-positive (black dots are major cities). Black inset in A) shows location of C) and Fig. 3A and 3B. Black inset in B) shows the location of Fig. 3C and 3D. D) Maximum likelihood phylogeny of *Bd* based on a concatenated alignment of 26 genome-wide loci totaling 6,035 bp generated by sequence capture from the positive sample collected from the SP (see Methods). Values indicated on the nodes are derived from ultrafast bootstrap and SH-aLRT test, respectively. Nodes <90% bootstrap support were collapsed. The SP sample (highlighted in bold) is placed phylogenetically in the hypervirulent GPL clade, and this site is proximal to the *Bd*-free SNSM, and may be the most likely source of colonization of the SNSM by *Bd*.

Amidst the epicenters of global amphibian declines, some communities persist seemingly unaffected (14-17). In regions providing environmental conditions that favor *Bd*-driven amphibian declines, the factors allowing host community survival remain enigmatic. Elevated tolerance or resistance might drive population survival (18) but appear unlikely to explain persistence of intact multi-host communities. Instead, local absence of hypervirulent *Bd* lineages within *Bd*-infested regions could explain amphibian community survival (15, 17). Geographic or environmental barriers might sustain localized pathogen absence long term (17). Such *Bd*-free refuges play a crucial role in averting future disease-driven biodiversity loss and potentially saving extant yet potentially highly susceptible species. Mitigating the impact of emerging infectious disease is most successful in a pre-invasion stage (19-21), thus the timely identification of disease-free refuges is critical to safeguard biodiversity.

The only major *Bd*-free refuge that has so far been uncovered is the island of New Guinea (NG) (15, 16, 22). NG is an amphibian hotspot, containing over 6% of the world’s amphibian species on less than 1% of its landmass (15). Many species are phylogenetically related to the Australian anuran lineages that have suffered extreme *Bd*-induced declines and extinctions (16). NG thus safeguards a substantial portion of Australasia’s evolutionary amphibian diversity (23). In the Neotropical realm, major disease-free refuges have not yet been identified. However, severe declines of amphibian populations throughout the region sharply contrasts with the seemingly unaffected, abundant amphibian communities of the Sierra Nevada de Santa Marta (SNSM) in northern Colombia (14, 24-26). The SNSM is an isolated coastal massif reaching 5,710 meters above sea level and a hotspot of amphibian richness and endemism, harboring at least 25 endemic species, including several endemic genera (27-30, *SI Appendix*, Tables S1, S2). Climatic conditions and community composition closely mirror other Neotropical montane systems that have experienced severe *Bd*-driven declines, and phylogenetically close Andean relatives of SNSM endemics have collapsed elsewhere (25, 26, 31).

Here, we evaluate whether the SNSM represents a *Bd*-free refuge. We predict (i) pre-epizootic amphibian host densities across species, (ii) presence of hypervirulent *Bd* lineages in adjacent mountain ranges, (iii) genome-wide SNP data of a focal species to indicate stable, diverse populations, (iv) environmental conditions conducive to establishment of *Bd* and *Bd*-driven host declines, and (v) host susceptibility to chytridiomycosis. To test these predictions and assess the vulnerability of amphibians of the SNSM, we combine mountain-range wide *Bd* surveillance, *Bd* lineage genotyping, host population genomics, pathogen niche modelling, and experimental infections. Further, we extend our niche modelling of the climatic envelope associated with *Bd*⍰ driven declines to New Guinea, allowing us to evaluate the risk of community⍰ wide declines should *Bd* be introduced. Finally, we outline a series of actions to avert yet another wave of disease-driven amphibian declines and extinctions.

## Results

### Landscape-scale screening uncovers the last major pathogen-free Neotropical biodiversity hotspot

Using a combination of quantitative PCR and CRISPR diagnostics on 2,212 amphibian skin swabs from at least 33 species and 27 environmental DNA (eDNA) samples, we were unable to detect *Bd* in any of the 34 sampled localities across the SNSM and adjacent lowlands (Fig. 1C; see *SI Appendix*, section 1, for a discussion of previous screening data). Bayesian hierarchical modelling yielded a posterior mean probability of 0.0297 per site that *Bd* was present but overlooked (95% highest posterior density 2.6e^−^□–0.0872, when assuming a test sensitivity of 0.75). Locality-wise posterior probabilities of infection were uniformly low across all sites (*SI Appendix*, Fig. S1). In the closest Andean mountain range to the SNSM, the Serranía de Perijá (SP) 20 km E of the SNSM, we detected *Bd* in an eDNA sample (Fig. 1C). Sequence capture confirmed *Bd* presence in the environmental sample from this site. Phylogenomic reconstruction placed the strain from the SP in the hypervirulent *Bd*-GPL clade (Fig. 1D). For details and GenBank accession numbers see *SI Appendix*, Table S3.

The SNSM is home to at least 48 amphibian species, including at least three endemic genera and 25 endemic species that occur exclusively in montane regions (*SI Appendix*, Tables S1, S2). Stream surveys from January 2024 in the SNSM revealed amphibian encounter rates of 26.5 ± 7.0 (mean ± SD) individuals per person per hour (range 17–39, n = 10 localities). Transect-based surveys of *Cryptobatrachus boulengeri* conducted in June 2023 showed a mean encounter rate of 16.3 ± 13.0 individuals per person per hour (range 1.6–49.7, n = 25 localities) and yearly surveys (2019–2024) of *Atelopus laetissimus* yielded 0.049 individuals m^−2^ (range 0.020–0.080) averaged across years (*SI Appendix*, Table S4).

### *Population genomics fail to find evidence for potentially* Bd*-induced bottlenecks in the recent past*

To test whether amphibian populations in the SNSM bear genomic signatures of recent bottlenecks, which could have been caused by past cryptic waves of *Bd* followed by population recovery, we generated whole-genome sequencing data for 19 individuals of the endemic harlequin toad, *Atelopus nahumae*, from one site in the northwestern SNSM (*SI Appendix*, Fig. S2). Observed heterozygosity exceeded heterozygosity expected under Hardy-Weinberg equilibrium in every individual genome (Fig. 2A), and all three inbreeding estimators (Fhat1–Fhat3) yielded negative values, indicating heterozygote excess and the absence of inbreeding or allele loss (*SI Appendix*, Table S5). Runs of homozygosity spanned very short distances (0–100 kb) with only a small number of stretches exceeding 300 kb, inconsistent with a recent bottleneck and increased levels of inbreeding (Fig. 2B) (32). Tajima’s D was strongly skewed towards negative values across genomic windows, consistent with a large, genetically diverse contemporary population (Fig. 2C).

**Figure 2.**
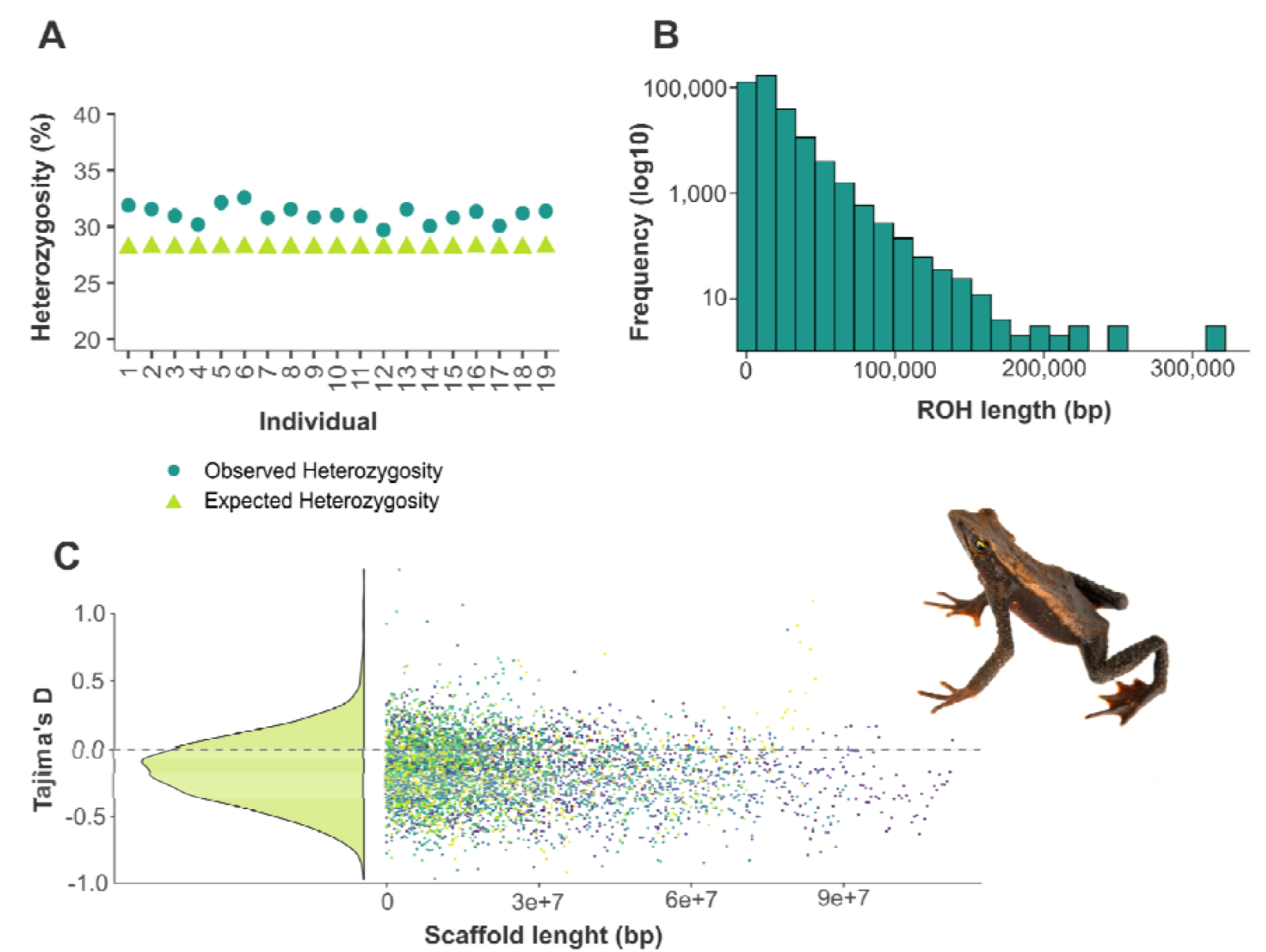
Host population genomics find no genomic signatures indicative of past *Bd*-induced declines in the Sierra Nevada de Santa Marta. A) Observed heterozygosity (turquoise dots) exceeds heterozygosity expected under Hardy-Weinberg equilibrium (green triangles) averaged across SNPs per individual in the representative species, *Atelopus nahumae*. B) Frequency of consecutive homozygous sections in the genome over all individuals. C) Tajima’s D for the *A. nahumae* population calculated over 500,000 bp windows for scaffolds ≥ 10 mbp of each individual. Photo: Jaime Culebras, Photo Wildlife Tours.

Population structure analyses further supported demographic stability. Principal component analysis revealed no clustering of individuals (*SI Appendix*, Fig. S3), and ADMIXTURE cross-validation identified K = 1 as the best-supported model, indicating a single cohesive population with no evidence of fragmentation, admixture, or hidden substructure (*SI Appendix*, Fig. S4). Based on the effective population size (*N*_e_) estimations across the last 150 generations, there are no signs of a rapid population contraction followed by recovery (*SI Appendix*, Fig. S5). See *SI Appendix*, section 2 for details and *SI Appendix*, Table S5 for sample accession numbers (BioProject PRJNA1390455).

### *Niche models reveal environmental conditions conducive to* Bd*-induced amphibian declines and narrow lowland barriers to* Bd *invasion*

We modelled *Bd* environmental suitability based on ∼6,000 global presence records using six climate variables selected in stepwise optimized experimental runs. Probability of occurrence and binary predictors revealed high environmental suitability for *Bd* across the entire SNSM below the snowline (Fig. 3A, *SI Appendix*, Fig. S6, S7). Environmental suitability was equally high in the adjacent SP and Andean mountain ranges in line with the pathogen’s known distribution in these areas. In contrast, lowland environments surrounding the SNSM, including the ∼20 km wide dry valley of the Río César (Fig. 1C) between SNSM and SP, showed low suitability for *Bd* (Fig. 3A). Binary predictions classified this region as mostly unsuitable. However, some binary predictors recovered larger areas within the Río César valley as suitable (*SI Appendix*, Fig. S6). For the second *Bd*-free refuge (15), New Guinea (NG), species distribution modelling revealed high environmental suitability across most of the island with highest suitability in the central highlands and lowest in northern lowlands (Fig. 3C). Binary predictions identified unsuitable regions throughout the lowlands north of the central mountain range, and along the eastern coast (*SI Appendix*, Fig. S6).

**Figure 3.**
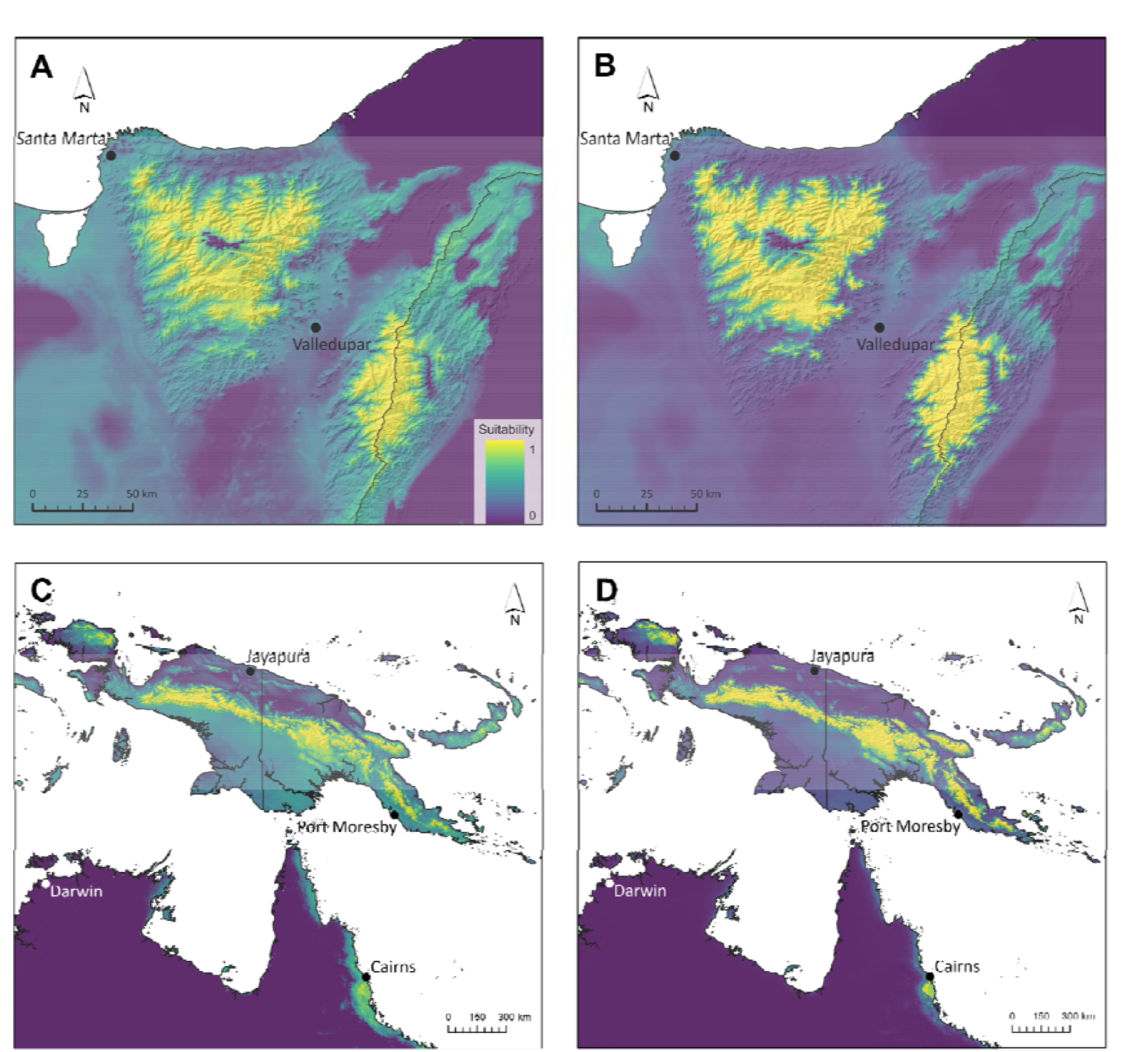
Niche models recover environmental barriers to pathogen introduction and suggest high risk for community declines following *Bd* invasion in the last major *Bd*-free refuges. A) Environmental suitability for *Bd* presence in the Sierra Nevada de Santa Marta (SNSM). Lowland climatic barriers of low environmental suitability towards the adjacent Serranía de Perijá and Andes are the likely cause of contemporary pathogen absence in the SNSM. B) Climatic envelope of *Bd*-driven declines suggests high risk for SNSM endemic amphibian communities. C) Environmental suitability for *Bd* presence in New Guinea (NG). D) Climatic envelope of *Bd*-driven declines suggests high risk for montane amphibian communities in NG. Warmer colors suggest higher suitability.

To gain further insight on whether *Bd* introduction may result in amphibian declines, we analyzed the climatic envelope associated with *Bd*-driven declines from other regions of the world using the same environmental predictors and georeferenced mass mortality events, declines, and extinctions likely induced by *Bd* (*SI Appendix*, Fig. S7). The resulting model showed a narrower range of environmental suitability for *Bd*-induced declines compared to *Bd* presence, with contraction towards tropical highlands, including high suitability for *Bd*-driven declines across all montane regions of SNSM and SP (Fig. 3B, *SI Appendix*, Fig. S7). In NG, the central montane regions showed high suitability for *Bd*-driven declines with low suitability in both northern and southern lowland areas (Fig. 3D, *SI Appendix*, Fig. S6, S7).

### Experimental infections highlight community susceptibility and high risk through reservoir dynamics

To assess direct susceptibility to *Bd*, we experimentally infected three SNSM-endemic amphibian species. Selected species included the stream-associated anurans, *Atelopus laetissimus* and *Cryptobatrachus boulengeri*, and the bromeliad-associated salamander, *Bolitoglossa savagei*. Infected *A. laetissimus* showed no mortality with two of nine inoculated individuals clearing infection. In the remaining seven individuals, infection loads remained lower than in the other species but increased towards the end of the experiment in five individuals (Fig. 4A; *SI Appendix*, Fig. S8, Table S7). Experimental infection of three additional specimens at a later stage and with higher zoospore dose recovered the same non-lethal infection pattern (*SI Appendix*, Table S7). In *B. savagei*, all experimentally inoculated individuals presented an increase in infection load with a single specimen reaching the clinical endpoint for euthanasia at week 10 (Fig. 4B; *SI Appendix*, Fig. S8), and three individuals presenting skin sloughing towards the end of the experiment. Infection loads did not saturate but increased during the experimental period in the surviving specimens of *B. savagei* (Fig. 4B; *SI Appendix*, Fig. S8, Table S7). In contrast, all infected *C. boulengeri* were euthanized following *Bd*-induced loss of self-righting ability within 7 weeks post inoculation after a short period of increased skin sloughing and abnormal hind limb posture (Fig. 4C). We did not observe significant differences in body mass change between infected and control individuals during the experiment (linear model on proportional mass change; *A. laetissimus*: p = 0.607, *C. boulengeri*: p = 0.535, *B. savagei*: p = 0.385, Fig. 4). Histopathology confirmed epidermal colonization by *Bd* in all three species.

**Figure 4.**
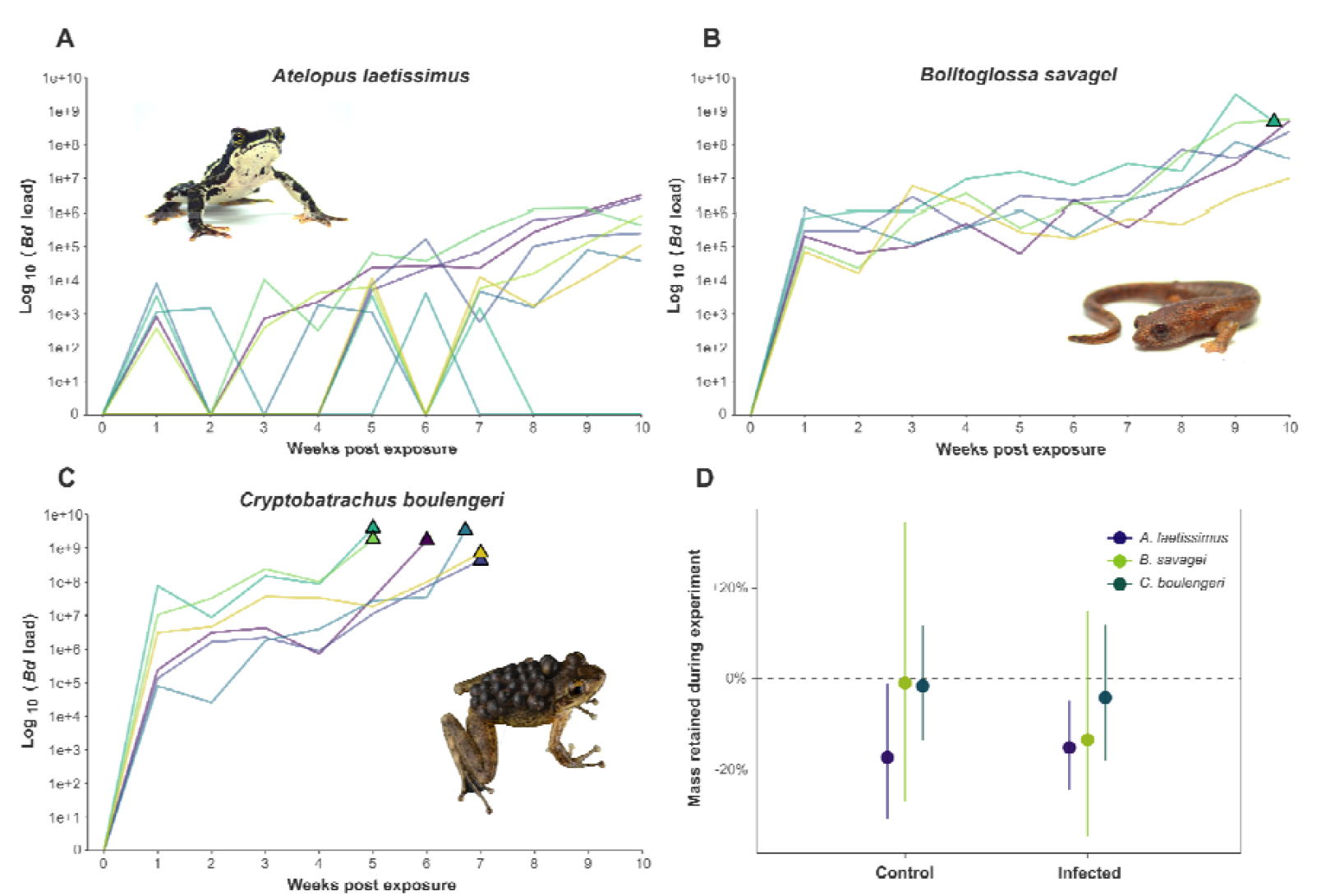
Experimental infection of three amphibian species from the Sierra Nevada de Santa Marta, Colombia. A) *Atelopus laetissimus*, tolerant to *Bd* infection, B) *Bolitoglossa savagei*, low susceptibility to *Bd* infection with delayed mortality underscoring potential reservoir function, C) *Cryptobatrachus boulengeri*, high susceptibility to *Bd* infection. Triangles depict mortality (euthanasia following chytridiomycosis-induced loss of self-righting ability). D) Percentage of body mass retained over the infection trial suggesting no *Bd*-induced effects.Photos: Amadeus Plewnia and Jaime Culebras, Photo Wildlife Tours.

## Discussion

Chytridiomycosis has significantly eroded amphibian diversity at the level of genes, species, and communities. Our results identify a rare and globally significant exception to this pervasive pattern. Five decades after *Bd* invaded Neotropical highlands and Australia, we provide multiple lines of evidence that the seemingly unaffected amphibian communities in the SNSM, bordered by a *Bd*-infested Andean landscape, remain epidemiologically naïve rather than having recovered in a post-epizootic state. However, proving pathogen absence is intrinsically difficult (33, 34). By combining extensive surveillance with negative results, high densities of experimentally susceptible host species, and a flagship species lacking a genomic signature of population decline, we provide independent lines of evidence for the absence of *Bd* in SNSM amphibian communities.

Identifying *Bd*-free refuges embedded within the world’s epicenters of amphibian decline is critical to safeguarding the remaining phylogenetic diversity of amphibians threatened by disease, especially when these occur in diversity hotspots (15, 17). NG and SNSM currently represent a narrow window of opportunity to prevent *Bd* introduction. *Bd*-free refuges represent unique pre-epizootic reference ecosystems. Intact montane amphibian assemblages, in the absence of *Bd* but in a climate conducive to its establishment and impact, offer the opportunity to study pristine community dynamics, and to design and apply conservation strategies before declines occur. Amphibian host abundances in the SNSM were equal to pre-epizootic abundances recorded for closely related Neotropical communities that have since suffered dramatic declines (11, 31). Host genomic diversity as shown for the SNSM-endemic harlequin toad *A. nahumae* indicates long-term demographic stability and substantial adaptive potential, yet such variation is likely to be rapidly eroded following pathogen invasion, as observed elsewhere (35, 36). Today, however, the high standing genetic variation provides a particularly strong basis for informed selection and management of *ex-situ* founder populations (37).

Niche models indicate high suitability for *Bd* presence and conditions conducive to *Bd*-induced declines throughout both refuges. While NG amphibian communities might benefit from oceanic isolation and lowland environments poorly suitable for *Bd* along potential coastal entry points, the SNSM is separated only by a narrow region of low environmental suitability for *Bd* in the surrounding lowlands and the dry valley to the east towards the Andes. These lowland environments likely act as a climatic barrier that has so far prevented host-mediated *Bd* dispersal from infected regions, such as the nearby Serranía de Perijá (SP) (17). In parallel, relatively low human impact in the montane zones of both SNSM and SP (*SI Appendix*, Fig. S9) combined with thermal inactivation of the pathogen’s heat-sensitive zoospores in lowland surroundings may have prevented anthropogenic translocation of *Bd* (38). However, *Bd* exhibits substantial ecological and evolutionary diversity (39-41), and other hypervirulent lineages with distinct thermal growth patterns could pass the lowland barrier into the SNSM. Further, binary predictors recover patches of environmentally suitable habitat throughout the lowland barrier. Coupled with the presence of *Bd*-GPL less than 20 km from the SNSM, *Bd* introduction is highly likely. In the case of NG, potential *Bd* emergence might be more complex, with different major *Bd* clades occurring in surrounding regions, such as the hypervirulent *Bd* GPL clade in Australia and the poorly studied yet possibly less virulent Asia3 clade on Sulawesi (41). Overall, increasing human activity such as the ongoing expansion of infrastructure, tourism, and trade in SNSM and NG will further exacerbate the risk for unintended human-mediated pathogen introduction through contaminated materials or stow-away amphibians (16, 42, 43).

Our experimental infections highlight the likely severity of future *Bd* invasion for SNSM communities but also raise hope for the persistence of some species. While *Cryptobatrachus boulengeri* showed rapid, lethal chytridiomycosis, *Atelopus laetissimus* and *Bolitoglossa savagei* developed increasing infection loads with the latter showing delayed mortality and no loss of infection, potentially contributing to reservoir dynamics with prolonged transmission and host-mediated spread (44, 45). The elevated tolerance of *A. laetissimus* is exceptional given the unparalleled chytridiomycosis-driven declines in the genus (26, 31) and high individual susceptibility to experimental infection in other *Atelopus* species (44, 46, 47). Understanding the mechanisms that enable *A. laetissimus* to survive with *Bd* infection might provide rare insights for safeguarding the >100 highly threatened emblematic harlequin toad species and underscores the importance of epidemiologically naïve refuges to conserve the global amphibian phylogenetic diversity. This tolerance is likely not the result of past pathogen-host coevolution, as we found no evidence of potentially *Bd*-induced declines in the closely related and syntopic *A. nahumae*. Together with *A. nahumae, A. laetissimus* is part of the phylogenetically most basal clade in the genus (48) and given the apparent lack of epizootic declines in the sister genus *Oreophrynella* (38), a phylogenetic signal in susceptibility might explain the elevated tolerance. However, our experiments evaluated only adults, while other life stages may exhibit higher disease susceptibility and disproportionately influence population trajectories during an epizootic (49, 50).

The SNSM harbors at least 25 endemic amphibian species, including evolutionarily distinct and endemic genera and several harlequin toad (*Atelopus*) species that are deeply embedded in regional indigenous culture, where the toad’s seasonal activity patterns inform agricultural cycles (*SI Appendix*, Table S1). In NG, the number of endemic taxa is even higher, with dozens of genera and hundreds of species (15, 23), yet experimental infection trials are lacking. Relative to ∼90 potentially extinct species and a further >400 declined species associated with *Bd* (6), pathogen introduction to these two large *Bd*-free refuges could strongly exacerbate global phylogenetic and functional amphibian diversity loss.

These insights argue for an explicit and unified emergency framework to safeguard remaining *Bd*-free refuges threatened by likely *Bd* introduction. Mitigation and prevention strategies for amphibian chytrid fungi have been developed and tested extensively over the past decades (19-21, 51-53). Core elements include (i) strict biosecurity protocols for human activities, (ii) continued active surveillance of *Bd* coupled with host population monitoring in refuges and adjacent areas, (iii) targeted susceptibility and mitigation studies spanning the evolutionary diversity of amphibian species in the refugia, and (iv) rapid development of *ex situ* capacity for the most vulnerable and evolutionarily distinct species in close collaboration with local and indigenous stakeholders. These efforts must be accompanied by an active, systematic search for additional refuges that may still persist undetected, particularly in yet poorly surveyed regions that harbor a climate conducive to *Bd*-driven declines.

Unlike during the initial waves of ‘enigmatic’ amphibian declines four decades ago, conservation science now has tailored much effort into researching pathogen dynamics, improving surveillance tools, and applying experimental mitigation approaches to detect, anticipate, and prevent pathogen emergence. Whether these tools can be translated into effective prevention will be determined by outcomes in the world’s last major *Bd*-free refuges. The SNSM and NG thus stand as parallel, irreplaceable ecosystems for averting further disease-driven biodiversity loss, raising hope for safeguarding global phylogenetic and functional amphibian diversity.

## Materials and Methods

### Pathogen screening

Permission to collect samples was granted by Autoridad Nacional de Licencias Ambientales (ANLA), Colombia, under research and collection permits RCM0014-00-2024 and RCI003-00-2020, with mobilization permits P02182S9571_N0055, P02182S9691_N0057 and P11487S10331_N0002. Some samples were exported to Germany under ANLA permit Nos. 003790, 003880 and 004120 validated by Secretaría Distrital de Ambiente, Distrito Capital de Bogotá. We conducted several field expeditions during both dry and rainy seasons from 2021 to 2025 targeting premontane and montane amphibian communities in 34 localities in the SNSM (Fig. 1C). Additional field expeditions during June to September of 2024 targeted adjacent lowland ecosystems as well as the neighboring mountain range Serranía de Perijá (SP; Fig. 1C). Amphibians were sought during nocturnal encounter surveys in an opportunistic manner resulting in the collection of 2,212 swab samples (Fig. 1C; *SI Appendix*, Table S6). The spatial structure of sampling was limited by the geopolitical situation, which precluded a larger sampling in the eastern, northeastern and southern slopes of the SNSM and adjacent lowlands (Fig. 1C). Swabbing followed 54 with swabs stored in 180 µl DNA/RNA Shield (Zymo Research, Freiburg) for DNA preservation or dry for on-site processing. In addition, we collected 27 water samples (*SI Appendix*, Table S6) for environmental detection of *Bd* as described in 55.

We deployed two diagnostic approaches (sample information in *SI Appendix*, Table S6): all eDNA and most swab samples were processed with quantitative PCR (qPCR) in duplicate following 56. For qPCR, samples were extracted using the DNeasy Blood & Tissue Kit (Qiagen, Hilden) with deviations from the manufacturer’s protocol as described in 57 for swabs and in 55 for environmental samples. qPCR reactions were performed in 20µl volumes on a StepOnePlus platform using TaqMan Fast Advanced reagents (Thermo Fisher, Waltham) and triplicate synthetic standards (10^1^-10^6^ ITS copies). We considered samples with sigmoidal amplification curves and Cq ≤ 40 positive, with a standard curve efficiency of 90-110% and R^2^ _≥_ 0.98 required to accept the results of a plate. To guide decisions on sampling design, some swab samples were processed on-site deploying CRISPR diagnostics (CRISPR-Dx) (*SI Appendix*, Table S6), as follows. We adapted the Cas12-based rapid assay, FINDeM (58), by using the improved fast DNA extraction protocol of 55, followed by isothermal Recombinase Polymerase Amplification and Cas12-activated cleavage of a fluorescence reporter. Performance of both methods has been compared previously (58, 59). FINDeM reaction conditions followed the supplementary protocol in 59 and reactions were run in triplicate alongside controls and visualized in the field with a handheld UV flashlight and UV glasses.

We assessed the posterior probability of infection per site for diagnostic zero detections as described in 33 using WinBUGS14 (60) with skin swab data only. We assumed three test sensitivity values for pathogen diagnostics (0.5, 0.75 and 1.0) and perfect specificity given zero detections in our dataset. For posterior probability estimates, we further combined samples by nearest-neighbor distance of ≤1 km.

As our eDNA screening in areas surrounding the SNSM led to the detection of *Bd* in environmental samples from the SP to the southeast, we employed a sequence-capture approach on the positive sample as an independent verification of *Bd* presence and to obtain data for phylogenomic placement of the *Bd* lineage. Sequence capture followed 61, with library preparation using the Watchmaker Genomics DNA Library Prep Kit. The resulting 26 loci were concatenated into a 6,035 bp alignment with no missing data. Besides the SP sample, the alignment contained a representative panel of whole-genome and sequence-capture *Bd* samples to determine the phylogenetic position of the SP-derived lineage (*SI Appendix*, Table S3). We inferred a maximum likelihood phylogeny using IQ-TREE2 version 2.3.6 (62) visualized assuming midpoint rooting as implemented in FigTree version 1.4.4 (63).

### Host population genomics

We selected *Atelopus nahumae* (Anura: Bufonidae) as a model species for population genomic profiling due to the high susceptibility of harlequin toads (*Atelopus*) to *Bd* (12, 26) and its stream associated life history in montane forests, where *Bd*-driven declines have been common in other harlequin toad species (see above). We evaluated i) genome-wide heterozygosity as an indicator for genetic variation for assessing the potential for genetic adaptation to any future outbreak of *Bd* and ii) the potential presence of demographic bottlenecks or inbreeding in the recent past that could indicate epizootic decline and subsequent recovery. We collected toeclips of 19 specimens from the northwestern SNSM (*SI Appendix*, Fig. S2) and extracted DNA using the DNeasy Blood & Tissue Kit. We sequenced whole genomes using DNBSeq technology (T7 platform; BGI, Hong Kong) from PCR-free, 150 bp paired-end libraries, targeting 10× assuming a genome size of 3.5 Gb (64) with coverage reported in *SI Appendix*, Table S5. Reads were adapter-trimmed and base-quality-filtered using the software FASTP version 0.23.4 (65) (Phred ≥ 20) before mapping against the *A. laetissimus* reference genome (GenBank accession PRJNA1142550; 64) using the software BWA-MEM2 version 2.2.1 (66). After stringent filtering for mapping quality using SAMtools version 1.18 (67) (MAPQ ≥ 30), we used ANGSD version 0.940 (68) to identify SNPs. ANGSD utilizes genotype likelihoods, which handle genotyping and mapping errors better than other SNP calling software at low coverage (68). Subsequently, tfam, tped were generated as input for PLINK version 1.9 (69). Using PLINK we generated VCF files from the tped and tfam as input for iSMC version 1.0.0 (70). We estimated inbreeding coefficients (Fhat1–Fhat3, following 71) (*SI Appendix*, Table S5) and population structure with PCA in PLINK. In brief, Fhat1 estimates the variance-standardised relatedness of an individual with the average individual of the population according to Hardy-Weinberg expectations. Fhat2 corresponds to the basic calculation of excess homozygosity. Fhat3 is based on Wright’s F (72) and calculates the correlation between uniting gametes. We calculated Tajima’s D with VCFtools (73) for scaffolds ≥10 mbp. Genome-wide heterozygosity and Tajima’s D provide information on overall genetic diversity and allele frequency distributions and can indicate populations inconsistent with recent severe bottlenecks. However, these metrics alone cannot distinguish long-term demographic stability from population recovery following a recent decline, as rapid post-bottleneck expansion may restore heterozygosity and generate negative Tajima’s D values. We therefore additionally quantified Runs of Homozygosity using BCFtools (74) for scaffolds ≥10 mbp, which provide insights into recent severe bottlenecks and inbreeding over few generations (32, 75), when potential cryptic *Bd*-induced declines would have been most likely. Population structure was further examined using ADMIXTURE (version 1.3.0; 76). In addition, we explored recent changes in effective population size (*N*_*e*_). We used iSMC to calculate the per-individual recombination rate (*SI Appendix*, Table S5) to obtain the cM/Mbp values necessary for implementation of GONE2 (77). Because iSMC is based on the sequentially Markovian coalescent (SMC) method, it is designed for large timeframes of hundreds of thousands of years. Accurate estimations of the past 100 years are not possible with SMC methods. The software package GONE2 uses linkage disequilibrium to estimate *N*_e_ in recent past generations. We therefore implemented GONE2 to accurately estimate the recent demographic history of *A. nahumae*. A mutation rate of 3e^-9^ per site per generation was assumed based on an estimation for *Atelopus manauensis* (78). Details on population genomic methods are provided in *SI Appendix*, section 2.

### *Niche Modelling for* Bd *presence and* Bd*-induced declines*

To assess environmental suitability for *Bd* throughout SNSM and NG, we compiled global occurrence records for *Bd* (spanning the period 1981–2024) from the Amphibian Disease Portal (79; downloaded 21 May 2025). We refrained from using regional subsets of *Bd* occurrences, as applied in previous work (80), because lineage-specific training data are not available at sufficient resolution and the complete global niche is most relevant for risk assessment given that the source of any future introduction into the *Bd*-free refuges cannot be predicted. We removed duplicate records by coordinates, resulting in ∼6,000 unique geo-referenced occurrences, 75% of which were used as training points in machine-learning correlative presence-only distribution modelling following the principle of maximum entropy. Climatic predictors were clipped to global land surfaces and included 66 numeric CHELSA version 2.1 variables (*SI Appendix*, Table S8, 30 arc seconds resolution, 1981–2010; 81-84), which provide increased spatial resolution compared to previous environmental suitability models for *Bd* (9, 79). We modeled *Bd* environmental suitability in MaxEnt v3.4.4 following 9. In brief, we used linear (L), product (P), and quadratic (Q) features to reduce overfitting (85). We ran ten replicate models and assessed model performance using area under the Receiver Operating Characteristic curve and omission rate (86). Predictor variables were reduced from 66 to six based on contribution metrics, jackknife tests, and response curve interpretability in two stepwise exploratory runs (*SI Appendix*, Fig. S10, S11). Selected variables include Annual Precipitation (bio12), Growing Season Length (gsl), Number of Growing Degree Days above 0 °C (ngd0), Net Primary Production on Land as Carbon Mass Flux (npp), Monthly Potential Evapotranspiration (pet_penman_min) and Surface Downwelling Shortwave Flux in Air (rsds_range) (*SI Appendix*, Fig. S12). Continuous clog-log outputs were further converted to binary predictions using three common thresholds (*SI Appendix*, Table S9): 10^th^ percentile training presence, maximum training sensitivity plus specificity, and maximum test sensitivity plus specificity (86-88). To further assess the putatively more confined climatic envelope in which *Bd* induces amphibian declines (89, 90), in a second run we compiled and georeferenced 969 global locality records at which amphibian mass mortalities, declines or extinctions most likely induced by *Bd* have been reported (*SI Appendix*, Fig. S12). We used the six climatic variables selected with presence data for MaxEnt modelling as described above.

### Infection trials

We used three representative species from the premontane and high montane SNSM amphibian communities to assess susceptibility to *Bd* infection. Selected species were phylogenetically unrelated and ecologically distinct (30), including the bromeliad-associated salamander, *Bolitoglossa savagei* (Plethodontidae) and the stream-associated anurans, *Cryptobatrachus boulengeri* (Hemiphractidae) and *Atelopus laetissimus* (Bufonidae). All animal experiments were approved by the institutional animal care and use committee (*Comité Institucional para el Cuidado y Uso de Animales, CICUA*) of Universidad de los Andes, Bogotá under code C.FUA_24-020. Collection of live individuals was authorized by Autoridad Nacional de Licencias Ambientales (ANLA) under Resolución 002166 with mobilization permit P11487S10853_N0005. Animals were collected during nocturnal surveys and transported to a controlled climate chamber (∼20°C, ∼80% relative humidity, 12 h light cycle) at the Universidad de los Andes, Bogotá. All frogs tested negative upon arrival using CRISPR-Dx, as described above. Live animals were weighed before inoculation and after euthanasia on a digital scale to the nearest 0.01 g and measured with calipers to the nearest 0.1 mm prior to inoculation. Animals were housed individually in terraria with wet paper towels as substrate, plastic hides and ad-libitum provision of feeder insects. Individuals were randomly assigned to treatment groups. After a one-week acclimatization period, we experimentally infected six individuals per species alongside six negative controls. The experiment was terminated 10 weeks after inoculation. Experimental setup followed 91: In brief, we used the hypervirulent *Bd*-GPL isolate JEL423 for infection trials, cultured on 0.75× strength TGhL plate media at 20°C. For the inoculum, we rinsed plates with 3 ml sterile distilled water and counted spores using a Neubauer improved hemocytometer. Spores were diluted to a final concentration of 5 × 10^6^ spores ml^-1^ and amphibians were inoculated in random order with 1 ml for 24 h in individual plastic tubes. Negative controls were sham-inoculated. In the case of *A. laetissimus*, due to slow infection progression during the first weeks of the experiment, incongruent with the high susceptibility of other *Atelopus* species (47), we inoculated three additional specimens that were not inoculated in the first trial. Inoculation of the additional specimens was conducted as described above but using a higher dose (8 × 10^6^ spores ml^-1^) four weeks post initial inoculation, to confirm tolerance of *A. laetissimus* to experimental infection with an increased sample size and an independent *Bd* inoculum (*SI Appendix*, Table S7). Individuals were inspected daily for clinical signs and skin swabs were collected every seven days and at the end of the experiment. Swabs were preserved in DNA/RNA Shield and analyzed with qPCR as described above. We set the loss of self-righting ability as a humane endpoint for euthanasia with an overdose of lidocaine. Upon euthanasia, we preserved skin sections in 5 % buffered formalin for histopathologic confirmation of epidermal *Bd* colonization (91). Sublethal effects were assessed as proportional change in body mass using the ln ratio of terminal to initial body mass, ln(terminal weight/initial weight). For each species, we fit linear models with infection status (inoculated vs. control) as the focal effect, using the natural logarithm of initial mass as covariate. Inference used HC3 heteroscedasticity-robust standard errors to reduce small sampling bias (92).

### Hygiene and biosafety protocols

During all field surveys, each individual was handled with a fresh pair of nitrile gloves (93). Prior to each field visit, boots and other equipment were thoroughly cleaned and subsequently disinfected with 70% ethanol or Virkon S and no sampling consumables previously brought to other field sites were employed. Infection experiments were carried out in a climate chamber at Universidad de los Andes with all materials in the climate chamber being disinfected with 70% ethanol and waste being autoclaved at the end of the experiment.

## Acknowledgments

We are grateful to Amanda B. Quezada Riera, Andrés Rocha Usuga, María José Navarrete Méndez and Viktoria Ferner for their help in the field. We thank Angie Sánchez Galán of the Museo de Historia Natural C. J. Marinkelle at the Universidad de los Andes for her help with obtaining permits. We are further grateful to Deborah Bower and Philipp Böning for valuable discussions and to Elvia Mora, Juana Diaz, Karin Fischer and Sabine Naber for their help in the lab. We want to thank the Arhuaco and Kogi communities of the Sierra Nevada de Santa Marta as well as the families from San Pedro de la Sierra and Palmor for their hospitality and support throughout our field expeditions. We are grateful to Fundación Atelopus for supporting logistics on field expeditions. We further thank the Colombian environmental authorities for issuing permits. This work was partially funded by the Forschungsfonds of Trier University, the Wilhelm-Peters-Fonds of the German Society of Herpetology DGHT, and the National Geographic Society (Grant number NGS-66509C-20). Computational resources on the HPC Elwetritsch at RPTU were provided by Rechenzentrumsallianz Rheinland-Pfalz (RARP). Amadeus Plewnia is funded by the Research Foundation Flanders (FWO) under PhD fellowship fundamental research 1104226N.

