## Supplementary material for "Imminent invasion of the chytrid fungus threatens the last naïve amphibian biodiversity hotspots": SI Appendix

#### Supporting Information Text

##### Section 1 – Discussion of previous *Batrachochytrium dendrobatids* (*Bd*) records from the Sierra Nevada de Santa Marta

(25) present positive *Bd* records in their Colombia-wide revision of the pathogen's distribution for the Sierra Nevada de Santa Marta (SNSM). Records are based solely on end-point PCR testing with the Bd1a/Bd2a primer pair (94). Positives were reported from two localities, "Reserva Buenavista" (*Atelopus laetissimus* and *Cryptobatrachus boulengeri*), a locality on Río Ancho in La Guajira on the northern slope of the Sierra Nevada de Santa Marta (SNSM) and from "SNSM" (*A. laetissimus* and *Serranobatrachus* sp. [as *Pristimantis* sp. 25]), a locality referring to the Cuchilla San Lorenzo in Magdalena on the northwestern flank of SNSM. The respective samples have been collected in the early 2010s by members of our team (L. A. Rueda Solano and colleagues) and further processed by members of our team (V. Flechas and colleagues). The samples have not been preserved and hence direct examination is impossible. While we acknowledge that these records could comprise true positives, we consider them as likely false positives for the following reasons: First, we have commonly observed issues of false positives with end-point PCR and electrophoresis verification due to the Bd1a/Bd2a primer pair commonly amplifying non-target fragments and forming secondary structures that are visible in gel electrophoresis. Second, we have included both localities in our recent sampling (29 samples from the first and 1,024 from the latter locality), all testing negative. Third, no declines or mortality events have been reported from both sites during the screening in the 2010s despite end-point positives also being found in the experimentally highly susceptible *C. boulengeri.* Fourth, both in the early 2010s (14) and today, densities remain similar to other pre-epizootic sites, including those for the highly susceptible *C. boulengeri* which occurs in both regions. Fifth, GONE2 analysis of *Atelopus nahumae* from the second site, Cuchilla San Lorenzo, rejects the presence of past genomic bottlenecks as expected in case of a post-epizootic recovery. In this site, *A. nahumae* occurs in syntopy with the species tested positive by (25). Importantly, (25) compiled a large country-wide dataset which included multiple distinct sampling subsets collected by different teams in distinct ecoregions. As such, it could be possible that the SNSM subset contributed from one exploration might have been contaminated while the remaining data presented by (25) comprises true positives. Multiple of the Andean and Chocoan sites where *Bd* was detected by (25) have been confirmed recently using quantitative PCR (95).

### Section 2 – Population genomic methods

For this study, we used the genome assembly of *Atelopus laetissimus* as a reference (64; Genbank Accession number PRJNA1142550). *Atelopus laetissimus* belongs to the Sierra Nevada clade of the genus *Atelopus,* as does *A. nahumae* studied in this paper (48). The reference genome was indexed for downstream analysis using *bwa-mem2* (66) and *samtools* (67) with the *index* and *faidx* commands, respectively. We trimmed adapters and filtered raw reads for Phred score < 20 using *fastp* (65). The sequences were then mapped against the reference genome using *bwa-mem2*. The resulting sam-formatted alignment files were then converted into bam-files, quality filtered with a threshold of 30, sorted according to their position in the genome, and indexed using *samtools*.

***angsd.*** We used the program suite *angsd* (68) to calculate genotype likelihoods from the bam-files using the *-GL 1* function. Specific input files (tfam and tped) required for the software PLINK were compiled in *angsd* with the function *-doPLINK2*. SNPs were filtered according to significance and genotype quality (-SNP_pval 1e-6, -postCutoff 0.99, -geno_minDepth 4).

***PLINK.*** For the following analyses, we used the software *PLINK* version 1.9 (69). The required input files (tfam and tped) contain genotype data as well as SNP and sample metadata. Different filtering steps were applied for different types of analysis, following (96). For genetic diversity functions (*-ibc* and *-het*), filtering included relatedness (*--rel-cutoff 0.4*), SNP and individual missing values (*--geno 0.5*; *--mind 0.5*), and the frequency and number of rare alleles (*--maf 0.05*; *--mac 2*). Furthermore *PLINK* only allows a maximum of 200 chromosomes or scaffolds. Given the high number of scaffolds of the only available reference genome, the *A. nahumae* dataset likewise contains >200 scaffolds. We therefore only considered scaffolds ≥10 mbp. The function *-ibc* was used to calculate three inbreeding coefficients (Fhat1–Fhat3) based on differing statistical approaches and definitions of inbreeding. Observed and expected homozygosity were calculated using the function *-het*. For Tajima's D calculations, we filtered out variants which have Hardy-Weinberg equilibrium exact test p-value below $1e^{-6}$ (*--hwe 1e-6*), variants with missing call rates exceeding 0.25 and samples with missing call rates exceeding 0.5 (*--geno 0.25*; *--mind 0.5*) and further filtered for minor allele count (*--mac 1*). Calculations were then performed with VCFtools (73) function *--TajimaD* for 500 kb windows of scaffolds ≥10 mbp. A VCF file with scaffolds >=10 Mbp, filtered for relatedness (--rel-cutoff 0.4), missing data (--geno 0.5; --mind 0.5) and rare variants (--maf 0.05; --mac 2) was used for counting runs of homozygosity in BCFtools (97) using the roh command. Prior to a principal component analysis (PCA) and *ADMIXTURE* analysis, the data were filtered for Hardy-Weinberg equilibrium (*--hwe 1e-6*), missing values (*--geno 0.5*; *--mind 0.5*), rare variants (*--maf 0.05*; *--mac 2*), linkage disequilibrium (*--indep-pairwise 50 10 0.5*) and relatedness (*--rel-cutoff 0.4*). None of the 19 individuals were removed by the latest filtering step. Again, only scaffolds ≥10 mbp were used for PCA and *ADMIXTURE*. Based on this subset, a PCA with 18 principal components was performed using the *-pca* function.

***ADMIXTURE.*** Based on the filtered data set described above an *ADMIXTURE* (version 1.3.0; 76) analysis was conducted. The optimal number of clusters (*K*) was determined by comparing cross-validation (CV) errors across multiple *K*-values, with lower CV errors indicating a better model fit.

***iSMC.*** To create single individual VCF files, which is one possible input for iSMC, a VCF was created in *PLINK* from the generated tped and tfam files using the command *-recode vcf* and extracted only the scaffold "ptg000004l" as it was among the longest with over 100 Mbp. Since *iSMC* requires a masking file, for example in BED format, we generated a one-lined BED file using "bcftools query -f'%CHROM\t%POS0\t%END\t%ID\n'" on the VCF file and extracted the very last SNP. This makes *iSMC* to utilize all other SNPs in the VCF for the calculation. We ran iSMC version 1.0.0. and calculated for the numerically first four individuals. For the iSMC parameter file (opt.bpp) we deviated from default parameters with the following settings: function_tolerance = 1e-1; number_intervals = 30;

number_rho_categories = 5; number_ne_categories = 1; number_theta_categories = 1; rho_var_model = Gamma; fragment_size = 2,000,000. Resulting rho and theta values (Tab. S5) from iSMC were transformed using the following equation to obtain a recombination rate in the unit cM/Mbp, which is required as GONE2 input: r = ((((2 x rho x µ) / theta) [M/bp]) x 100 [cM/bp]) x 1,000,000 [cM/Mbp] (98). This changed equation results from the implementation of rho in the current Version of *iSMC* as rho= 2 x Ne x r (Julien Y. Dutheil *in litt.* 11.12.2025). Using the four individuals (Accesion Numbers SAMN54183605-SAMN54183612), iSMC calculated rho = 0.0289979668739593; theta = 0.00934062603536013. Resulting in an r = 1.862699573 cM/Mbp.

***GONE2.*** Because SMC methods like *iSMC* utilize recombination rates (few per generation) and mutation rates (even more rare), recent timeframes are unreliable to estimate (77). GONE2 includes a linkage-disequilibrium based approach leading to a higher reliability in estimating Ne over the last 150 generations. For *Atelopus nahumae*, this corresponds to approximately 300 years, assuming a generation time of 2 years, and thus covers the period that is decisive for the question addressed in this study. As for iSMC, we only used scaffold "ptg000004l" for which we subsetted to 2 million SNPs in the VCF files. We used default parameters for unphased diploid genomes in GONE2, supplying a recombination rate of 1.863, rounded from the iSMC calculation. The resulting slowly increasing shape across more than 50 Generations (i.e. > 100 years) and sharp decrease of $N_e$ in the past ~5 generations (~10 years) is reminiscent of signals in metapopulations (77) or alternatively artifacts from expansive genetic changes maybe caused by secondary contact of differentiated populations. As PCA and Admixture do not hint towards subpopulations in the dataset, it appears as this is an artifact from an extensive genetic diversity as result of population expansion or secondary contacts (99). This could potentially lead to a lack of continuity between physical and genetic maps causing the recent 5 generations to drastically decrease in $N_e$. Low linkage disequilibrium (LD) among closely linked polymorphisms combined with high LD among more distantly linked polymorphisms could result from selective sweeps associated with secondary contact. This is supported by observations of high contemporary numbers of individuals in the field and all other indices that contradict an ongoing population contraction.

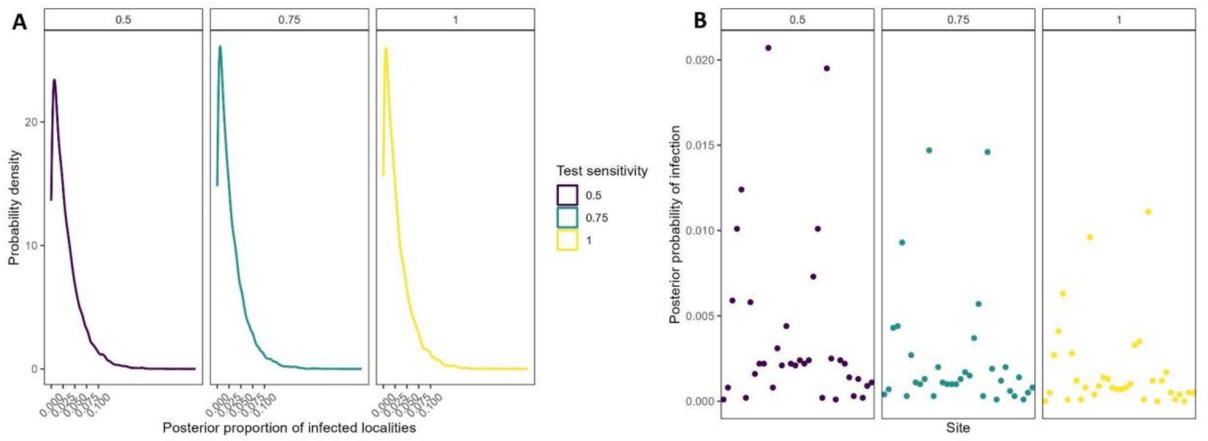

**Figure S1.** A) Distribution of the posterior probability that a randomly chosen site was *Bd*-infected under three different test-sensitivity assumptions (0.5, 0.75, and 1.0). B) Posterior probability of infection for individual sites under the same test-sensitivity assumptions.

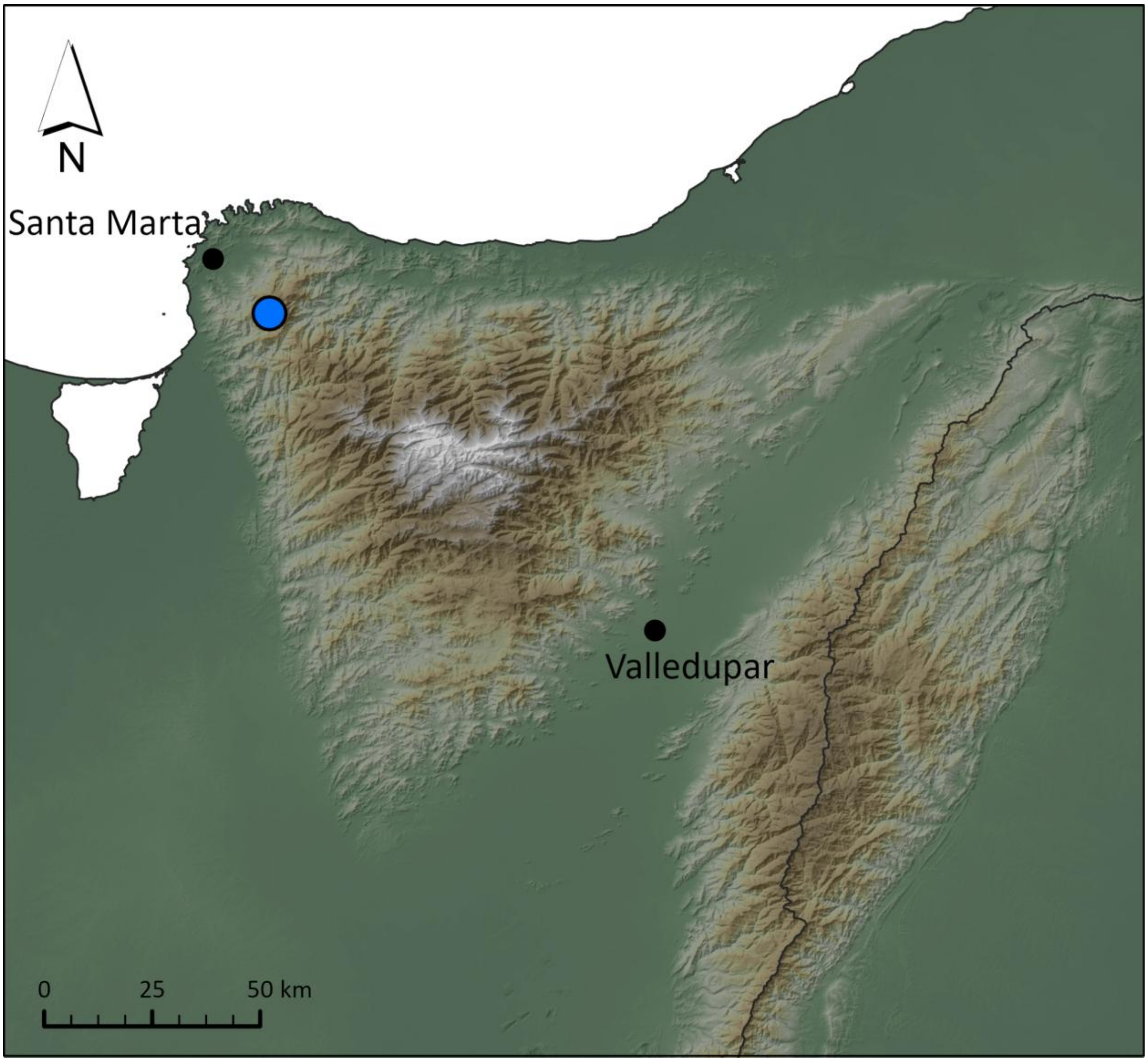

**Figure S2.** Sampling locality (blue dot) for population genomic analyses of *Atelopus nahumae*.

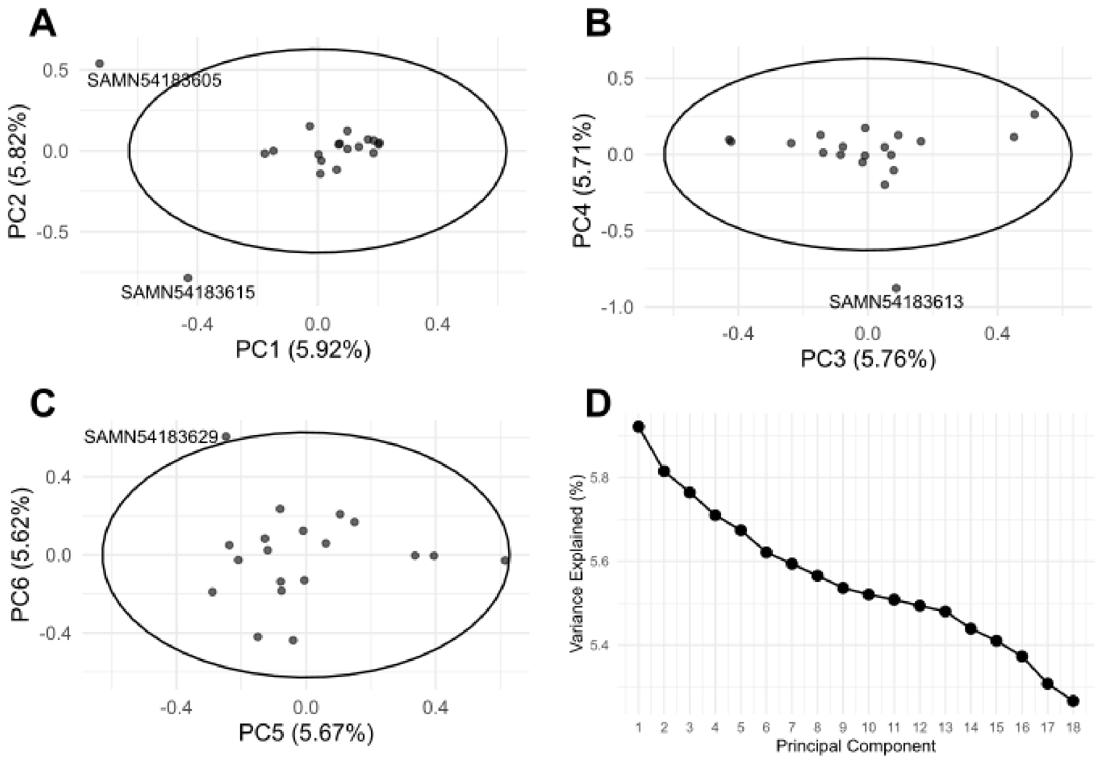

**Figure S3.** Principal component analysis (PCA) of genetic variation among individuals, based on SNP data from scaffolds >10mbp. (A) PC1 vs. PC2; (B) PC3 vs. PC4; (C) PC3 vs PC4. Each point represents one individual. Ellipses indicate the 95% confidence interval based on a multivariate normal distribution. Labelled individuals fall outside this interval and may be considered outliers. (D) Scree plot showing the proportion of variance explained by the 18 principal components (PCs). Each point represents the percentage of total variance explained by the corresponding PC.

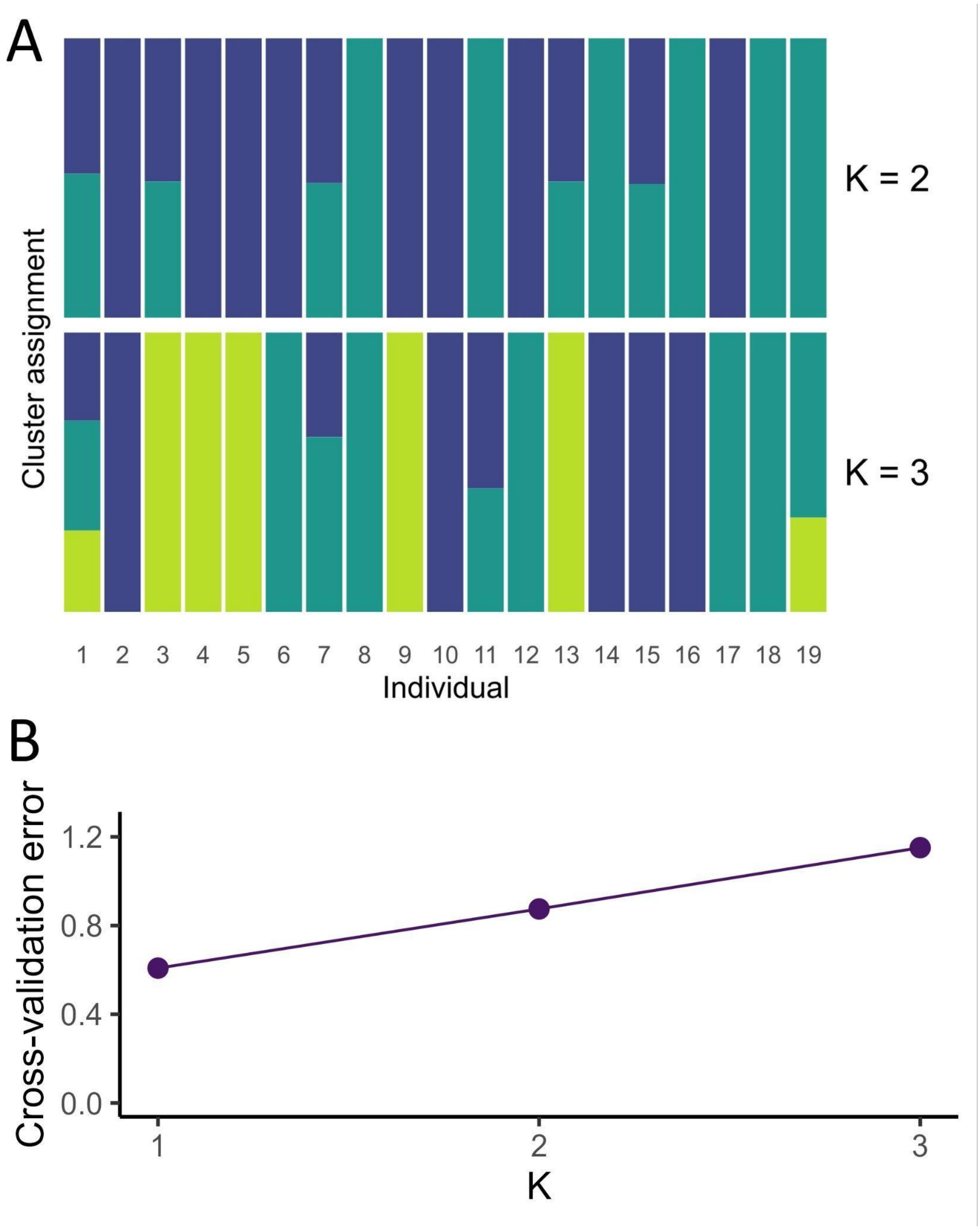

**Figure S4.** (A) ADMIXTURE barplots showing estimated ancestry proportions for each individual based on SNP data, assuming two different scenarios of genetic clustering (K = 2 and K = 3 genetic clusters). Each vertical bar represents one individual. Colouration indicates the proportion of ancestry assigned to each inferred genetic cluster. (B) Cross-validation error estimates from ADMIXTURE analysis for K = 1 to 3, supporting a single cluster for the studied population. Each point corresponds to the cross-validation error for a given K.

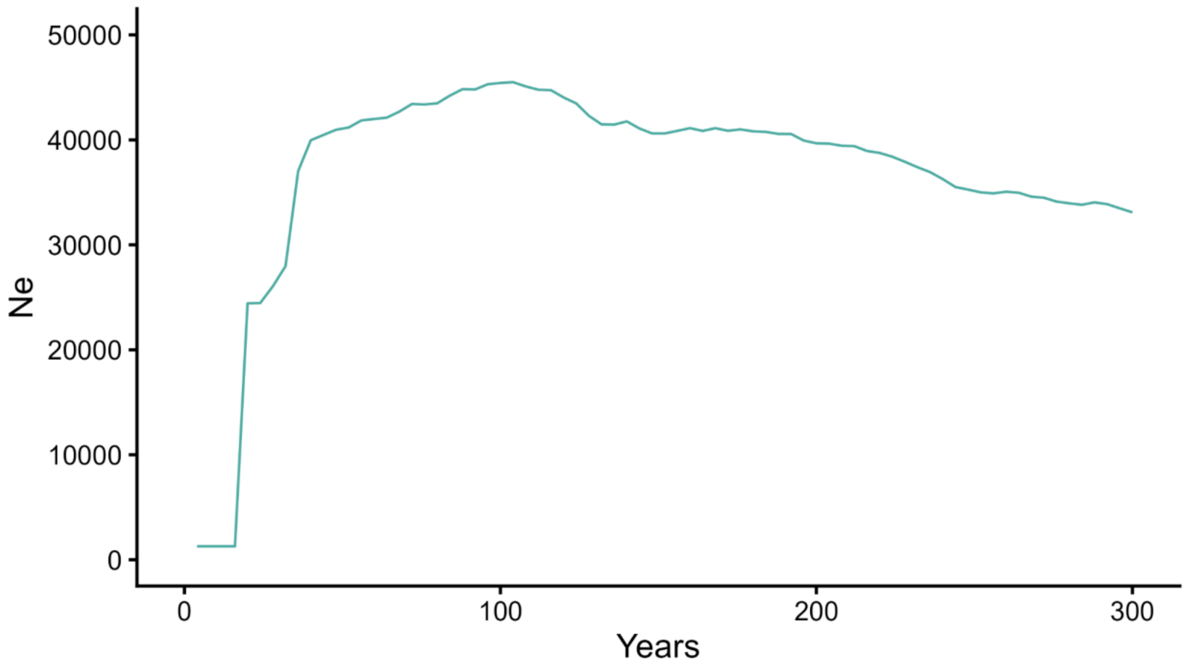

**Figure S5.** Effective population size of *A. nahumae* over past 150 generations (about 300 years) inferred with the linkage-disequilibrium-based software GONE2. No past genomic bottleneck followed by recovery detected. The sharp decrease of Ne towards the present is a known artifact in expanding diverse populations as described above.

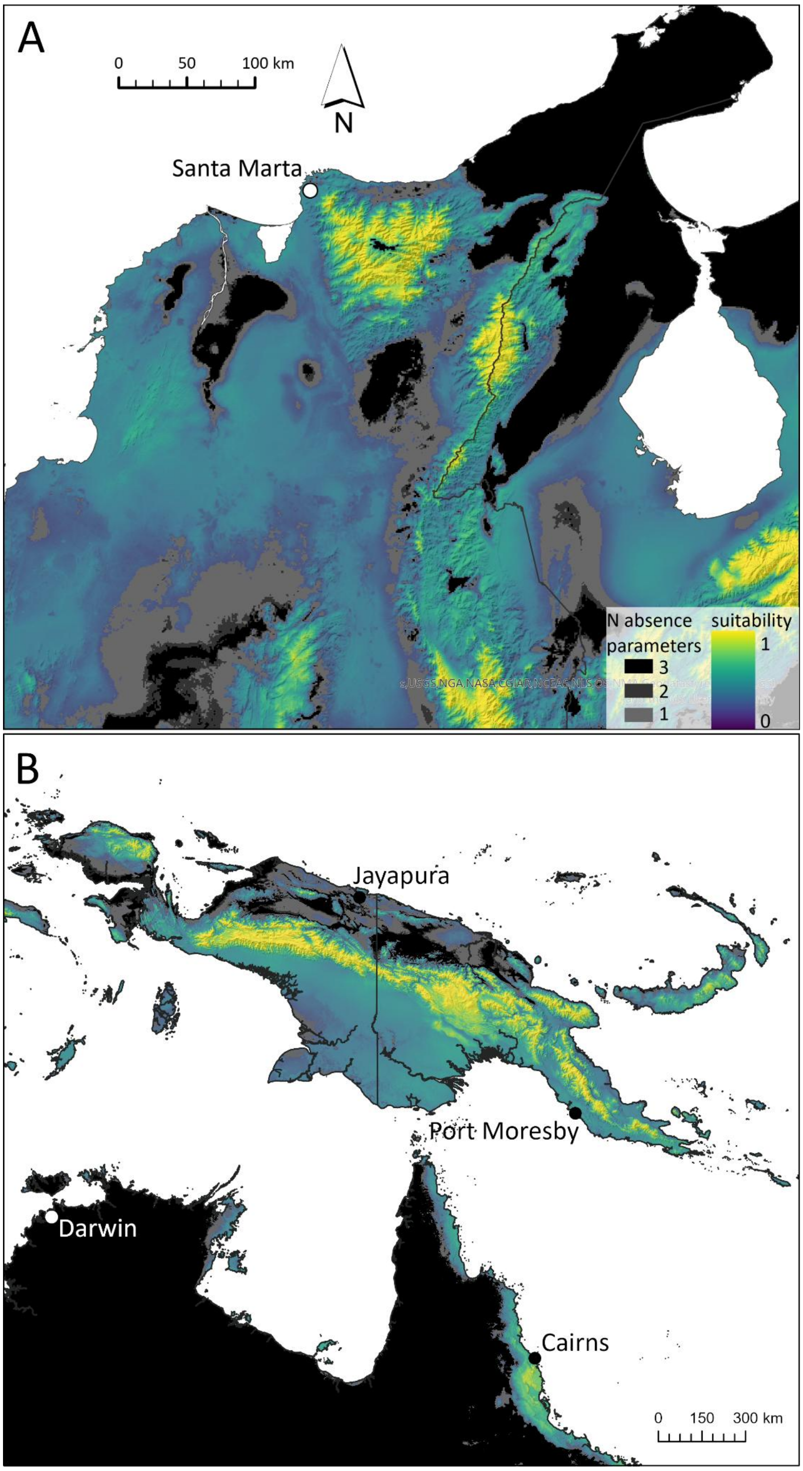

**Figure S6.** Mapped species distribution model for environmental suitability for *Bd* presence (with higher suitability indicated by warmer colors) in the (A) SNSM and (B) NG, overlaid by presence/absence classification using the parameters 10 percentile training presence clog-log threshold, maximum training sensitivity plus specificity clog-log threshold and maximum test sensitivity plus specificity clog-log threshold (with darker colors in grey-scale indicating several parameters suggestion absence). Note lowland barrier between SNSM and SP as well as lowland barriers extending along the foothills of the Andean central and eastern Cordillera.

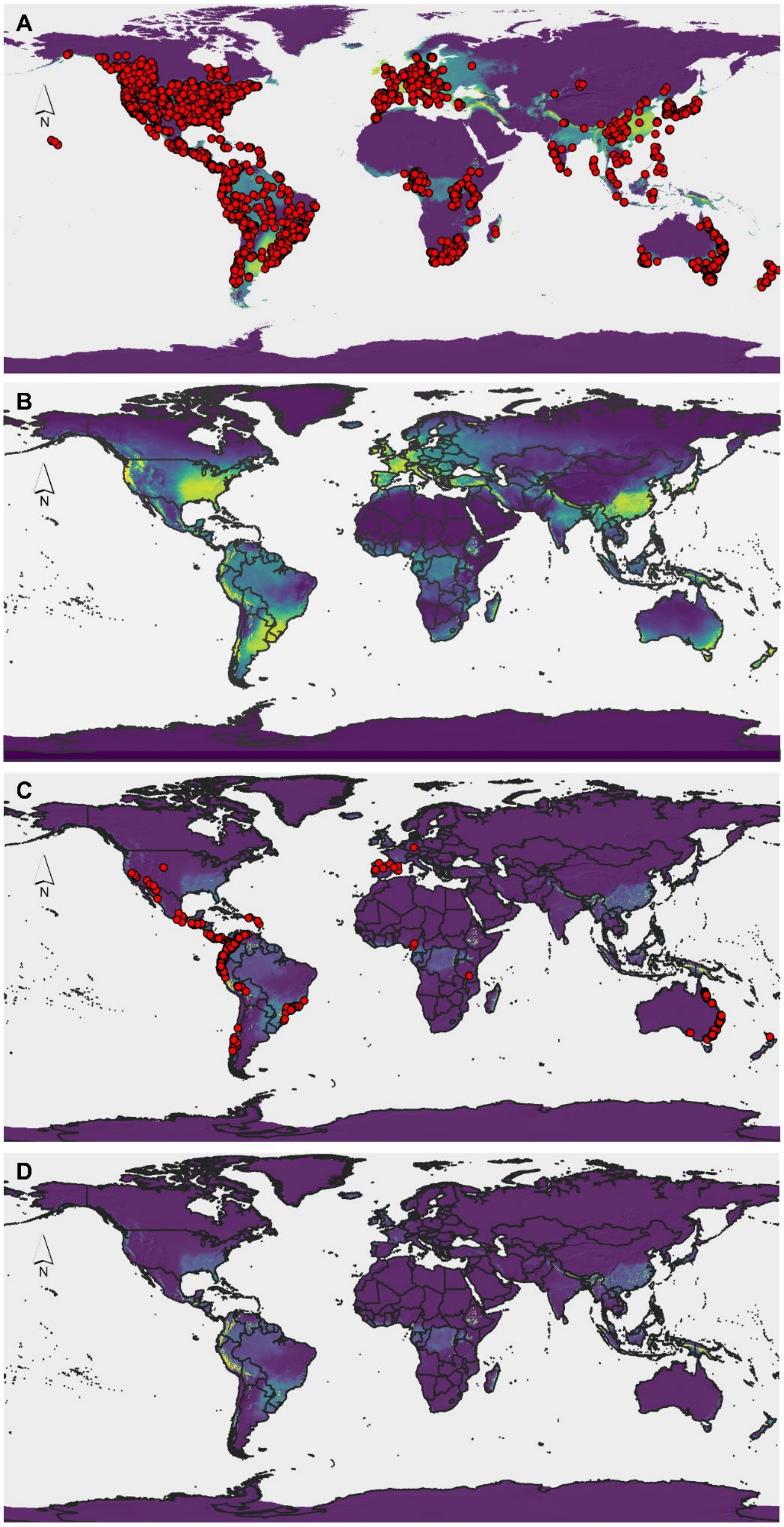

**Figure S7.** Global extent of environmental suitability for *Bd* presence and *Bd*-induced declines. (A) Global occurrence records used for the presence model. (B) Global model for environmental suitability of *Bd* presence. (C) Records of *Bd*-induced population declines and extinctions. (D) Global model of the climatic envelope for *Bd*-driven host declines. Higher environmental suitability is indicated by warmer colors; color scale as in Fig. 3. Note the strongly reduced spatial extent which predicts high environmental suitability for *Bd*-induced declines, with highest values in Neotropical montane regions and Papuan montane regions among other (smaller) hotspots.

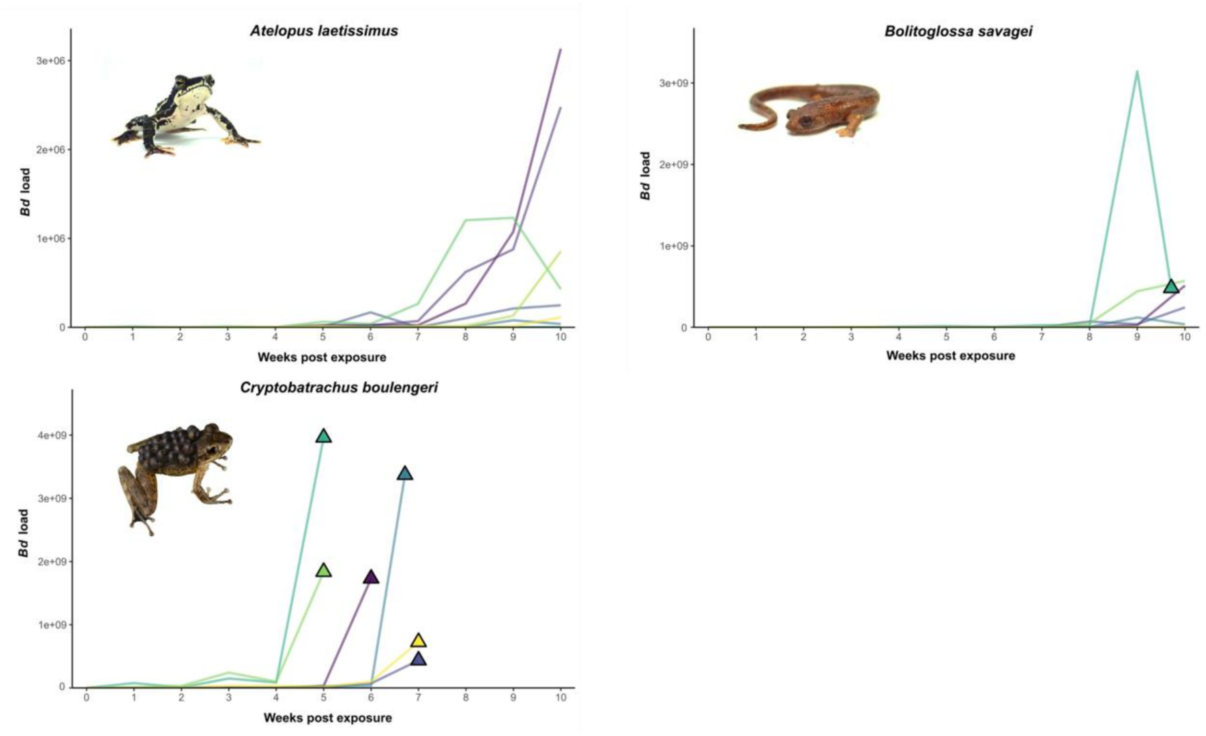

**Figure S8.** Infection loads during experimental *Bd* infection. Plots are those of Fig. 4 with linear y scale instead of log scale to highlight differences in terminal infection load between specimens.

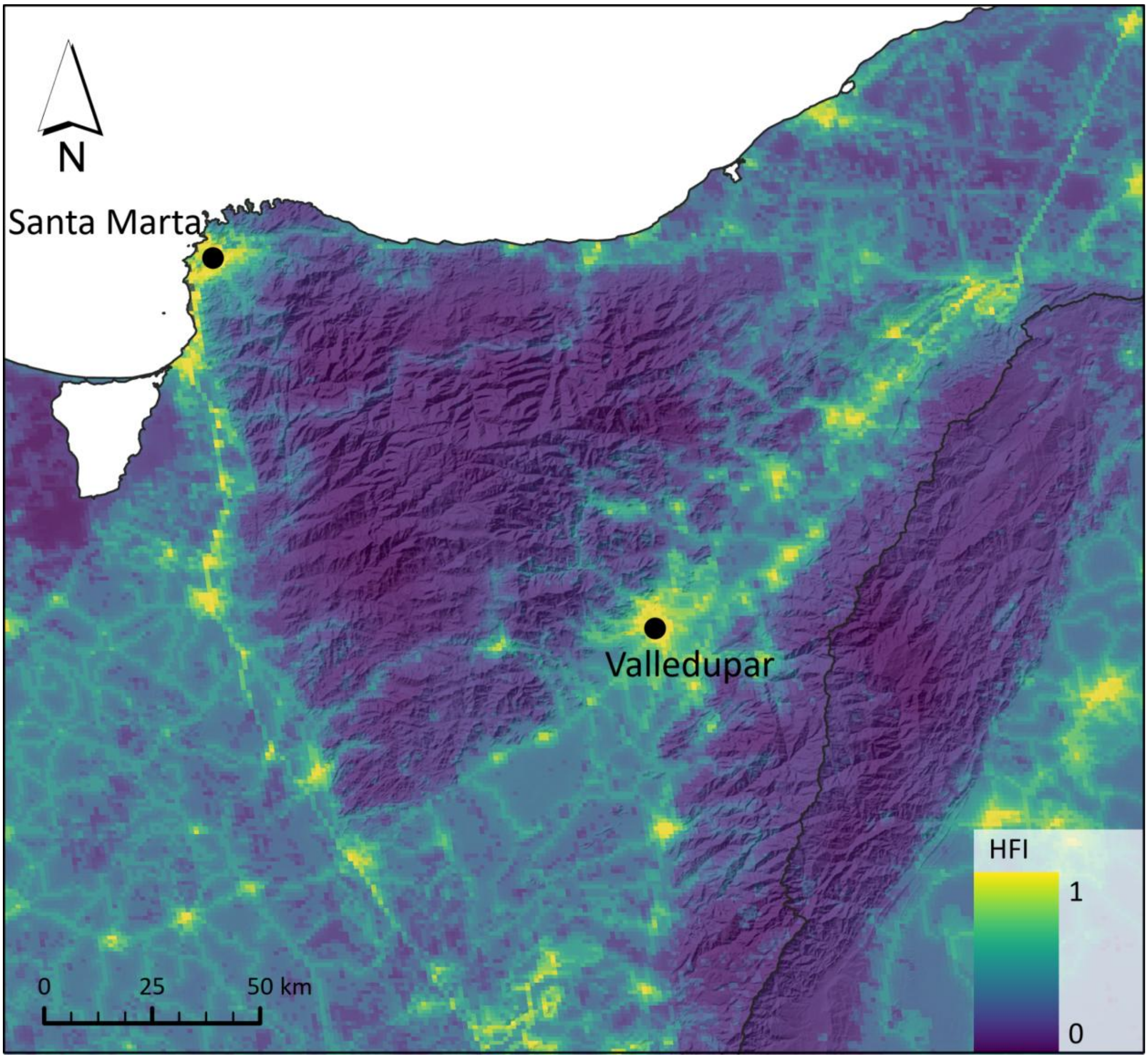

**Figure S9.** The Human Footprint Index (HFI, 2024 dataset; 100) suggests low human activity in both the SNSM and the SP. Low human activity may have limited jump-dispersal of *Bd* into the SNSM, see text.

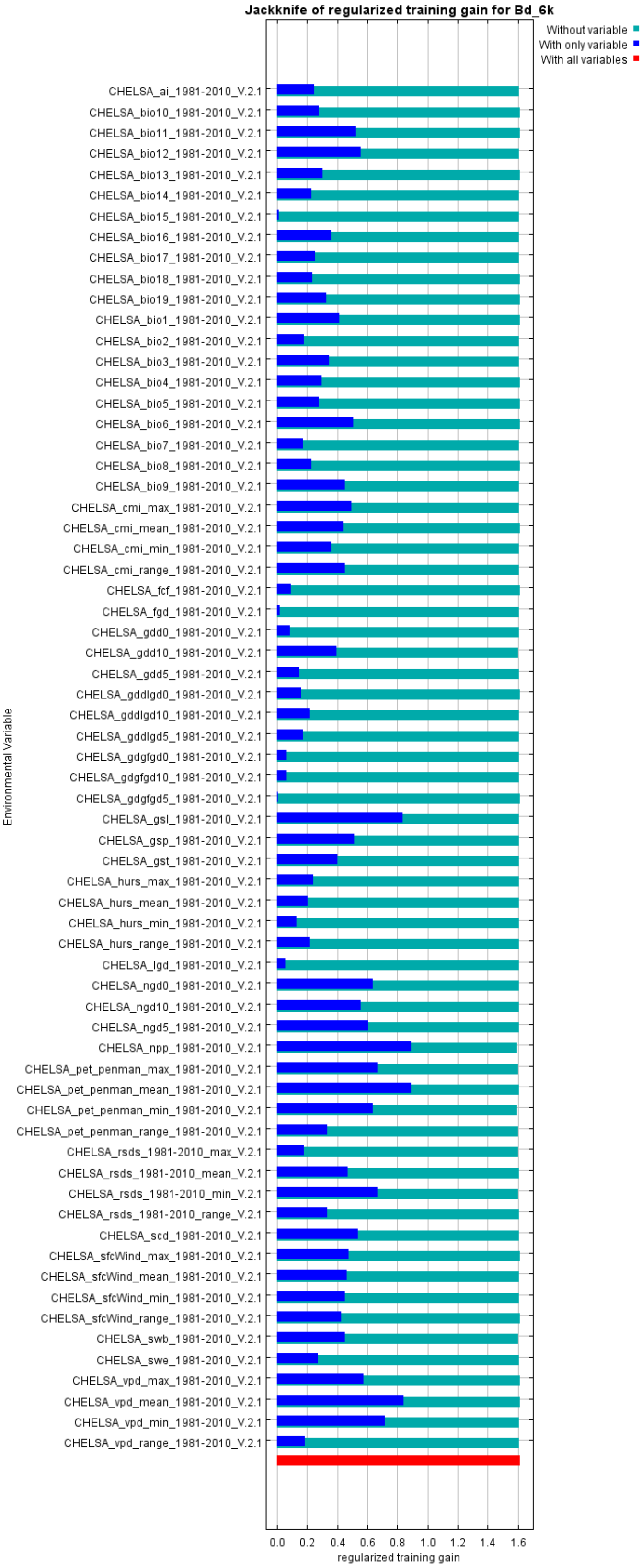

**Figure S10.** Jackknife tests from five exploratory replicate models for the environmental niche of *Bd* presence including all 66 CHELSA variables.

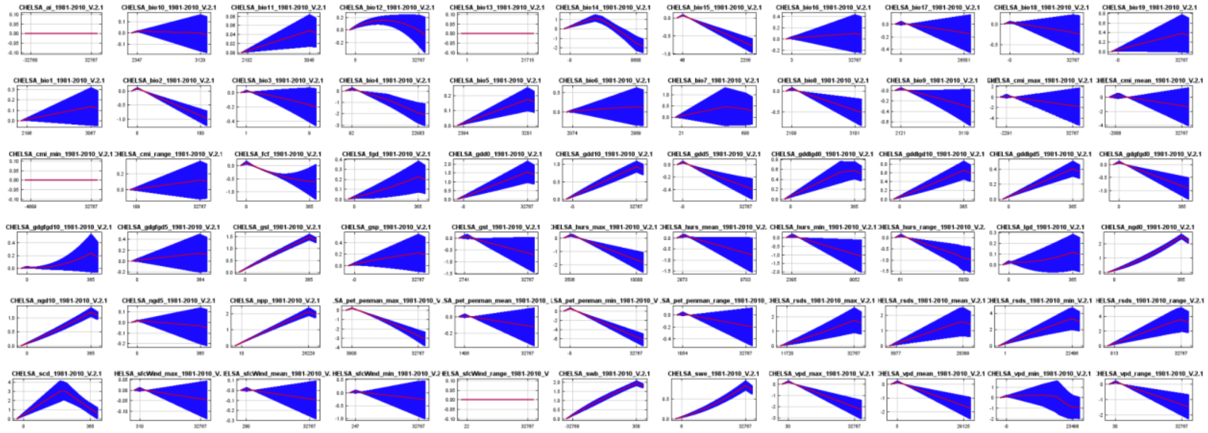

**Figure S11.** Response curves from five exploratory replicate models for the environmental niche of *Bd* presence including all 66 CHELSA variables.

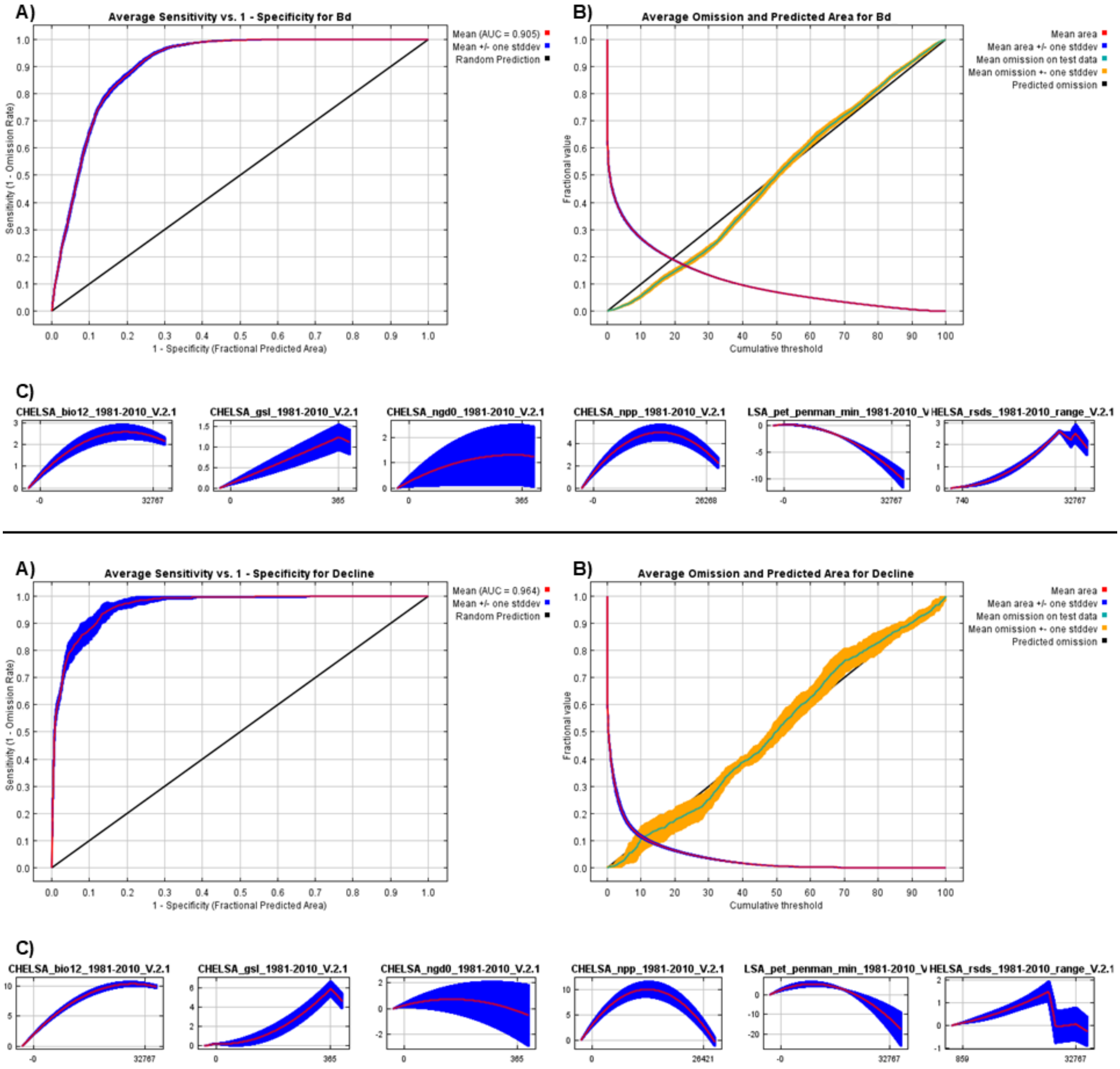

**Figure S12.** Area under the Receiver Operating Characteristic curve (A), omission rate (B) and response curves (C) from ten replicate models for the environmental niche of *Bd* presence (above) and the climatic envelope of *Bd*-induced amphibian declines (below).

**Table S1.** List of endemic amphibian species found in the SNSM. Endemic genera marked by an asterisk.

| **species** | **family** | **reference** | **elevational range in m a.s.l.** | **remarks** |
|---|---|---|---|---|
| *Atelopus arsyecue* | Bufonidae | Rueda-Almonacid (1994) | 2000-3500 | endemic clade (most basal in genus), see Lötters et al. (2025). See www.atelopus.org for cultural value of harlequin toads in indigenous SNSM communities |
| *Atelopus carrikeri* | Bufonidae | Ruthven (1916) | 2350-4800 | endemic clade (most basal in genus), see Lötters et al. (2025). See www.atelopus.org for cultural value of harlequin toads in indigenous SNSM communities |
| *Atelopus laetissimus* | Bufonidae | Ruiz-Carranza et al. (1994) | 1500-2900 | endemic clade (most basal in genus), see Lötters et al. (2025). See www.atelopus.org for cultural value of harlequin toads in indigenous SNSM communities |
| *Atelopus leoperezii* | Bufonidae | Ruiz-Carranza et al. (1994) | 2350-4800 | endemic clade (most basal in genus), see Lötters et al. (2025). For the validity of this sp., see Lötters et al. (2025). See www.atelopus.org for cultural value of harlequin toads in indigenous SNSM communities |
| *Atelopus nahumae* | Bufonidae | Ruiz-Carranza et al. (1994) | 1500-2800 | endemic clade (most basal in genus), see Lötters et al. (2025). See www.atelopus.org for cultural value of harlequin toads in indigenous SNSM communities |
| *Atelopus walkeri* | Bufonidae | Rivero (1963) | 1500-2160 | endemic clade (most basal in genus), see Lötters et al. (2025). See www.atelopus.org for cultural value of harlequin toads in indigenous SNSM communities |
| *Ikakogi ispacue** | Centrolenidae | Rada et al. (2019) | 950 | endemic genus, most basal lineage in family (Guayasamin et al. 2008, 2009) |
| *Ikakogi tayrona** | Centrolenidae | Ruiz-Carranza & Lynch (1991) | 950-2600 | endemic genus, most basal lineage in family (Guayasamin et al. 2008, 2009) |
| *Serranobatrachus carmelitae** | Craugastoridae | Ruthven (1922) | 1520-2200 | endemic genus (Arroyo et al. 2022) |
| *Serranobatrachus cristinae** | Craugastoridae | Lynch & Ruiz-Carranza (1985) | 1530-3500 | endemic genus (Arroyo et al. 2022) |
| *Serranobatrachus delicatus** | Craugastoridae | Ruthven (1917) | 1500-2600 | endemic genus (Arroyo et al. 2022) |
| *Serranobatrachus insignitus** | Craugastoridae | Ruthven (1917) | 1530-2134 | endemic genus (Arroyo et al. 2022) |
| *Serranobatrachus* | Craugastori | Ruthven (1917) | 1300-2450 | endemic genus (Arroyo et al. |

| | | | | |
|---|---|---|---|---|
| *megalops** | dae | | | 2022) |
| *Serranobatrachus ruthveni** | Craugastoridae | Lynch & Ruiz-Carranza (1985) | 1800-3500 | endemic genus (Arroyo et al. 2022) |
| *Serranobatrachus sanctaemartae** | Craugastoridae | Ruthven (1917) | 1100-2600 | endemic genus (Arroyo et al. 2022) |
| *Tachiramantis tayrona* | Craugastoridae | Lynch & Ruiz-Carranza (1985) | 1300-2700 | genus endemic to SNSM, SP and small portion of adjacent eastern Cordillera (Arroyo et al. 2022) |
| *Tachiramantis* sp. "Minca" | Craugastoridae | undescribed species; Erens et al. (2025) | 1000-1550 | genus endemic to SNSM, SP and small portion of adjacent eastern Cordillera (Arroyo et al. 2022) |
| *"Colostethus" ruthveni** | Dendrobatidae | Kaplan (1997) | 470-2100 | undescribed endemic genus that is siter to all Dendrobatini (Grant et al. 2017) |
| *"Colostethus"* sp. Valledupar* | Dendrobatidae | undescribed species; Grant et al. (2017) | unknown | undescribed endemic genus that is siter to all Dendrobatini (Grant et al. 2017) |
| *Cryptobatrachus boulengeri* | Hemiphractidae | Ruthven (1916) | 250-1790 | genus endemic to SNSM, SP and adjacent Andean Cordilleras (Lynch 2008) |
| *Cryptobatrachus ruthveni* | Hemiphractidae | Lynch (2008) | 1400-1540 | genus endemic to SNSM, SP and adjacent Andean Cordilleras (Lynch 2008) |
| *Bolitoglossa savagei* | Plethodontidae | Brame & Wake (1963) | 1000-2800 | |
| *Bolitoglossa* sp. "Sogrome" | Plethodontidae | undescribed species | 1780 | This candidate species is supported by molecular (GenBank PX105574) and bioacoustic data |
| *Geobatrachus walkeri** | unknown | Ruthven (1915) | 1550-2870 | likely endemic family based on unpubl. mitogenomic data |
| *Geobatrachus* sp. 1* | unknown | undescribed species; Erens et al. (2025) | 2200-2600 | likely endemic family based on unpubl. mitogenomic data. This candidate species is supported by molecular and bioacoustic data (L.A. Rueda Solano) |

**Table S2.** List of additional (non-endemic) amphibian species found in the lower regions of the SNSM and its periphery.

| species | family | reference | remarks |
|---|---|---|---|
| *Boana platanera* | Hylidae | This study, as *B. crepitans* in Pérez-González et al. (2016) | previously reported as *B.* aff. *xerophyla* for the region |
| *Boana pugnax* | Hylidae | Pérez-González et al. (2016) | |
| *Dendropsophus microcephalus* | Hylidae | Pérez-González et al. (2016) | |
| *Pseudis paradoxa* | Hylidae | Pérez-González et al. (2016) | |
| *Scarthyla vigilans* | Hylidae | Pérez-González et al. (2016) | |
| *Scinax rostratus* | Hylidae | Pérez-González et al. (2016) | |
| *Scinax ruber* | Hylidae | Pérez-González et al. (2016) | |
| *Trachycephalus typhonius* | Hylidae | Pérez-González et al. (2016) | |
| *Caecilia subnigricans* | Caeciliidae | iNaturalist records | |
| *Ceratophrys calcarata* | Ceratophryidae | Pérez-González et al. (2016) | |
| *Dendrobates truncatus* | Dendrobatidae | Pérez-González et al. (2016) | |
| *Elachistocleis panamensis* | Microhylidae | Pérez-González et al. (2016) | presence of *Elachistocleis pearsei* in the close periphery uncertain but likely |
| *Engystomops pustulosus* | Leptodactylidae | Pérez-González et al. (2016) | |
| *Leptodactylus fragilis* | Leptodactylidae | Pérez-González et al. (2016) | |
| *Leptodactylus fuscus* | Leptodactylidae | Pérez-González et al. (2016) | |
| *Leptodactylus insularum* | Leptodactylidae | Pérez-González et al. (2016) | |
| *Leptodactylus savagei* | Leptodactylidae | Pérez-González et al. (2016) | |
| *Pleurodema brachyops* | Leptodactylidae | Pérez-González et al. (2016) | |
| *Pseudopaludicola pusilla* | Leptodactylidae | Pérez-González et al. (2016) | |
| *Lithobates vaillanti* | Ranidae | Pérez-González et al. (2016) | |
| *Phyllomedusa venusta* | Phyllomedusidae | Collections at Universidad del Magdalena | |
| *Rhinella horribilis* | Bufonidae | This study, as *R. marina* in Pérez-González et al. (2016) | |
| *Rhinella humboldti* | Bufonidae | Pérez-González et al. (2016) | |

**Table S3.** Whole genome and sequence capture-derived sequences used for phylogenetic placement of *Bd* from the Serranía de Perijá. Newly generated sequence data from this study in bold.

| Name in tree | Reference | Host-species | Data Type | SRA | Strain | Clade | Location | PE/SS | N raw reads |
|---|---|---|---|---|---|---|---|---|---|
| South Korea: Gangwon-Do KRBOOR_317 | O'Hanlon et al. (2018) | *Bombina orientalis* | Genomic data | SRR6375581 | KRBOOR_317 | Bd-Asia1 | South Korea | PE | 19.644.468 |
| South Korea: Gangwon-Do KRBOOR_323 | O'Hanlon et al. (2018) | *Bombina orientalis* | Genomic data | SRR6375580 | KRBOOR_323 | Bd-Asia1 | South Korea | PE | 24.259.701 |
| Brazil: São Paulo CLFT001 | O'Hanlon et al. (2018) | *Hylodes japi* | Genomic data | SRR6375524 | CLFT001 | Bd-Brazil | Brazil | PE | 7.672.502 |
| South Korea: Gyeonggi-Do KB108 | O'Hanlon et al. (2018) | *Rana catesbeianus* | Genomic data | SRR6375525 | KB108 | Bd-Brazil | South Korea | PE | 18.044.501 |
| Captive specimen Cameroon LM2 | O'Hanlon et al. (2018) | *Leptopelis rufus* | Genomic data | SRR6375544 | LM2 | Bd-Cape | United Kingdom | PE | 8.796.156 |
| South Africa SAKN-5 | O'Hanlon et al. (2018) | Pyxicephalidae | Genomic data | SRR6375526 | SAKN-5 | Bd-Cape | South Africa | PE | 4.700.300 |
| Switzerland: Zürich 739 | O'Hanlon et al. (2018) | *Alytes obstetricans* | Genomic data | SRR6375451 | CH_0739 | Bd-CH | Switzerland | PE | 18.257.745 |

| | | | | | | | | | |
|---|---|---|---|---|---|---|---|---|---|
| Canada: Quebec JEL261 | O'Hanlon et al. (2018) | *Rana catesbeianus* | Genomic data | SRR6375485 | JEL261 | Bd-GPL | Canada | PE | 27.052.567 |
| Panama: Chiriqui JEL423 | O'Hanlon et al. (2018) | *Agalychnis lemur* | Genomic data | SRR6375517 | JEL423 | Bd-GPL | Panama | PE | 24.226.737 |
| USA: Arizona LMT04 | Mulder et al. (2024) | *Rana chiricahuensis* | Sequence capture | SRR27383576 | LMT04 | Bd-GPL | USA | PE | 6.619.423 |
| Venezuela: Mérida JEL429 | O'Hanlon et al. (2018) | *Rana catesbeiana* | Genomic data | SRR635201 | JEL429 | Bd-GPL | Venezuela | SE | 41.487.211 |
| Colombia: Cundinamarca EV001 | Rosenblum et al. (2013) | *Rheobates palmatus* | Genomic data | SRR634967 | EV001 | Bd-GPL | Colombia | SE | 34.211.525 |
| **Ecuador: Carchi M10** | **This study** | ***Pristimantis crucifer*** | **Sequence capture** | **SRR36443896** | **M10** | **Bd-GPL** | **Ecuador** | **PE** | **14.170.207** |
| **Colombia: Serranía de Perijá environmental sample** | **This study** | **aquatic eDNA** | **Sequence capture** | **SRR36443897** | **Perija3** | **Bd-GPL** | **Colombia** | **PE** | **20.600.397** |

**Table S4.** Amphibian community abundance/density data recorded during nocturnal skin swab collection along transects in humid montane forest of the SNSM. Note host densities are similarly high as in other pre-epizootic studies (e.g. 11, 31).

| locality | amphibian density | date | number of species | remarks |
|---|---|---|---|---|
| Estación Experimental San Lorenzo, Magdalena department | 0.049 individuals per m² (range 0.020–0.080) | yearly surveys 2019-2024 | NA, this monitoring project considered only one species, *Atelopus laetissimus*. | This estimate is only for one species, *Atelopus laetissimus*. Visual nocturnal encounter surveys are conducted in a 50x10m area along Quebrada San Lorenzo. Unpubl. data Fundación Atelopus. |
| Minca road, stream 1, Magdalena department | 31 individuals per person*hour | 11. Jan. 2024 | Three species detected, *Cryptobatrachus boulengeri, Ikakogi tayrona, Tachiramantis* sp. “Minca” | |
| Minca road, stream 2, Magdalena department | 22 individuals per person*hour | 11. Jan. 2024 | Four species detected, *Cryptobatrachus boulengeri, Ikakogi tayrona, Serranobatrachus megalops, Tachiramantis* sp. “Minca” | |
| Minca road, stream 3, Magdalena department | 17 individuals per person*hour | 11. Jan. 2024 | Three species detected, *Cryptobatrachus boulengeri, Ikakogi tayrona, Tachiramantis* sp. “Minca” | |
| Minca road, stream 4, Magdalena department | 19 individuals per person*hour | 11. Jan. 2024 | Three species detected, *Cryptobatrachus boulengeri, Ikakogi tayrona, Tachiramantis* sp. “Minca” | |
| Minca road, stream 5, Magdalena department | 27 individuals per person*hour | 24. Jan. 2024 | Three species detected, *Cryptobatrachus boulengeri, Ikakogi tayrona, Tachiramantis* sp. “Minca” | |
| Casas Viejas, stream 1, Magdalena department | 22 individuals per person*hour | 13. Jan. 2024 | Three species detected, *Colostethus ruthveni, Cryptobatrachus boulengeri, Ikakogi tayrona* | |
| Casas Viejas, stream 2, Magdalena department | 30 individuals per person*hour | 13. Jan. 2024 | One species detected, *Cryptobatrachus boulengeri* | |
| San Pedro de la Sierra, Magdalena department | 39 individuals per person*hour | 19. Jan. 2024 | Six species detected, *Bolitoglossa savagei, Cryptobatrachus ruthveni, Serranobatrachus delicatus, S. insignitus, S. megalops, S. sanctaemartae* | |
| stream above San Pedro de la Sierra, Magdalena department | 24 individuals per person*hour | 20. Jan. 2024 | Five species detected, *Atelopus laetissimus, Bolitoglossa* cf. *savagei, Serranobatrachus carmelitae, S. ruthveni, S. sanctaemartae* | |
| Palmor, Magdalena department | 34 individuals per person*hour | 26. Jan. 2024 | Five species detected, *Boana platanera, Colostethus ruthveni, Cryptobatrachus ruthveni, Lithobates vaillanti, Rhinella horribilis* | |

| | | | |
|---|---|---|---|
| Minca, transect 1 | 8 individuals recorded in 1:10 person*hours | 01. Jun. 2023 | NA, this monitoring project considered only one species, *Cryptobatrachus boulengeri*. |
| Minca, transect 2 | 3 individuals recorded in 1:50 person*hours | 01. Jun. 2023 | NA, this monitoring project considered only one species, *Cryptobatrachus boulengeri*. |
| Minca, transect 3 | 1 individual recorded in 0:30 person*hours | 02. Jun. 2023 | NA, this monitoring project considered only one species, *Cryptobatrachus boulengeri*. |
| Minca, transect 4 | 11 individuals recorded in 1:20 person*hours | 02. Jun. 2023 | NA, this monitoring project considered only one species, *Cryptobatrachus boulengeri*. |
| Minca, transect 5 | 8 individuals recorded in 1:20 person*hours | 02. Jun. 2023 | NA, this monitoring project considered only one species, *Cryptobatrachus boulengeri*. |
| Minca, transect 6 | 12 individuals recorded in 1:15 person*hours | 03. Jun. 2023 | NA, this monitoring project considered only one species, *Cryptobatrachus boulengeri*. |
| Minca, transect 7 | 8 individuals recorded in 1:00 person*hours | 03. Jun. 2023 | NA, this monitoring project considered only one species, *Cryptobatrachus boulengeri*. |
| Minca, transect 8 | 14 individuals recorded in 0:40 person*hours | 04. Jun. 2023 | NA, this monitoring project considered only one species, *Cryptobatrachus boulengeri*. |
| Minca, transect 9 | 87 individuals recorded in 2:20 person*hours | 05. Jun. 2023 | Four species detected, *Cryptobatrachus boulengeri, Engystomops pustulosus, Rhinella horribilis, R. humboldti* |
| Minca, transect 10 | 29 individuals recorded in 0:35 person*hours | 07. Jun. 2023 | Four species detected, *Boana pugnax, Cryptobatrachus boulengeri, Engystomops pustulosus, Rhinella horribilis* |
| Minca, transect 11 | 6 individuals recorded in 0:35 person*hours | 07. Jun. 2023 | Two species detected, *Boana pugnax, Cryptobatrachus boulengeri* |
| Minca, transect 12 | 9 individuals recorded in 0:15 person*hours | 07. Jun. 2023 | Two species detected, *Boana pugnax, Cryptobatrachus boulengeri* |
| Minca, transect 13 | 6 individuals recorded in 0:30 person*hours | 07. Jun. 2023 | NA, this monitoring project considered only one species, *Cryptobatrachus boulengeri*. |
| Minca, transect 14 | 5 individuals recorded in 1:10 person*hours | 07. Jun. 2023 | NA, this monitoring project considered only one species, *Cryptobatrachus boulengeri*. |
| Minca, transect 15 | 7 individuals recorded in 0:30 person*hours | 12. Jun. 2023 | NA, this monitoring project considered only one species, *Cryptobatrachus boulengeri*. |
| Minca, transect 16 | 6 individuals recorded in 0:30 person*hours | 12. Jun. 2023 | NA, this monitoring project considered only one species, *Cryptobatrachus boulengeri*. |
| Minca, transect 17 | 19 individuals recorded in 0:30 person*hours | 13. Jun. 2023 | NA, this monitoring project considered only one species, *Cryptobatrachus boulengeri*. |

| | | | |
|---|---|---|---|
| Minca, transect 18 | 8 individuals recorded in 0:20 person*hours | 13. Jun. 2023 | NA, this monitoring project considered only one species, *Cryptobatrachus boulengeri*. |
| San Pedro, transect 1 | 13 individuals recorded in 1:00 person*hours | 16. Jun. 2023 | NA, this monitoring project considered only one species, *Cryptobatrachus boulengeri*. |
| San Pedro, transect 2 | 22 individuals recorded in 0:40 person*hours | 17. Jun. 2023 | NA, this monitoring project considered only one species, *Cryptobatrachus boulengeri*. |
| San Pedro, transect 3 | 11 individuals recorded in 1:10 person*hours | 17. Jun. 2023 | NA, this monitoring project considered only one species, *Cryptobatrachus boulengeri*. |
| San Pedro, transect 4 | 9 individuals recorded in 0:40 person*hours | 18. Jun. 2023 | NA, this monitoring project considered only one species, *Cryptobatrachus boulengeri*. |
| San Pedro, transect 5 | 8 individuals recorded in 2:20 person*hours | 18. Jun. 2023 | NA, this monitoring project considered only one species, *Cryptobatrachus boulengeri*. |
| San Pedro, transect 6 | 17 individuals recorded in 1:00 person*hours | 20. Jun. 2023 | NA, this monitoring project considered only one species, *Cryptobatrachus boulengeri*. |
| San Pedro, transect 7 | 18 individuals recorded in 1:00 person*hours | 20. Jun. 2023 | NA, this monitoring project considered only one species, *Cryptobatrachus boulengeri*. |

**Table S5.** Population genomic metrics and accession numbers for whole genomic data of *Atelopus nahumae*.

| Individual | HET Exp % | HET obs % | Fhat1 | Fhat2 | Fhat3 | average coverage per genomic position | Accession number |
|---|---|---|---|---|---|---|---|
| nahumae_1 | 0.281279400929931 | 0.318722120862021 | -0.0435958 | -0.133336 | -0.088466 | 16.19149 | SAMN54183605, SAMN54183606 |
| nahumae_2 | 0.281798499970175 | 0.315542129002931 | -0.0768946 | -0.0910583 | -0.0839764 | 18.85102 | SAMN54183607, SAMN54183608 |
| nahumae_3 | 0.281604272898842 | 0.309496960235109 | -0.0663301 | -0.0690297 | -0.0676799 | 18.03145 | SAMN54183609, SAMN54183610 |
| nahumae_4 | 0.281567535826905 | 0.301865780197712 | -0.0922354 | -0.041304 | -0.0667697 | 17.79856 | SAMN54183611, SAMN54183612 |
| nahumae_5 | 0.281700103901915 | 0.321482235559133 | -0.0603125 | -0.125326 | -0.0928191 | 17.91856 | SAMN54183613, SAMN54183614 |
| nahumae_6 | 0.281729935364737 | 0.325718063009366 | -0.0427348 | -0.152522 | -0.0976285 | 18.51102 | SAMN54183615, SAMN54183616 |
| nahumae_7 | 0.281334939285426 | 0.307605099976931 | -0.0896607 | -0.0601612 | -0.074911 | 18.51102 | SAMN54183617, SAMN54183618 |
| nahumae_8 | 0.281562014752609 | 0.315464317822668 | -0.0693126 | -0.104239 | -0.0867759 | 17.49297 | SAMN54183619, SAMN54183620 |
| nahumae_9 | 0.281506678332703 | 0.308383764484839 | -0.0666165 | -0.0738759 | -0.0702462 | 17.10231 | SAMN54183621, SAMN54183622 |
| nahumae_10 | 0.281577147904147 | 0.310183180708278 | -0.0816419 | -0.0709235 | -0.0762827 | 17.58632 | SAMN54183623, SAMN54183624 |

| | | | | | | | |
|---|---|---|---|---|---|---|---|
| nahumae_11 | 0.28143267845644 | 0.309132805704844 | -0.092073 | -0.0630504 | -0.0775617 | 17.06731 | SAMN54183625, SAMN54183626 |
| nahumae_12 | 0.281253695430644 | 0.297001824333588 | -0.0860346 | -0.0225853 | -0.0543099 | 16.67084 | SAMN54183627, SAMN54183628 |
| nahumae_13 | 0.281548469489931 | 0.31541540692227 | -0.0747813 | -0.0946555 | -0.0847184 | 17.76214 | SAMN54183629, SAMN54183630 |
| nahumae_14 | 0.281259905422387 | 0.300363384958166 | -0.0933277 | -0.0376787 | -0.0655032 | 16.43575 | SAMN54183631, SAMN54183632 |
| nahumae_15 | 0.281458284026962 | 0.307827116839076 | -0.0968274 | -0.0569167 | -0.0768721 | 17.12953 | SAMN54183633, SAMN54183634 |
| nahumae_16 | 0.282145105361958 | 0.313289362953966 | -0.100613 | -0.066884 | -0.0837486 | 21.96766 | SAMN54183635, SAMN54183636 |
| nahumae_17 | 0.280972490710414 | 0.300592071560429 | -0.0775488 | -0.0448527 | -0.0612007 | 15.29783 | SAMN54183637, SAMN54183638 |
| nahumae_18 | 0.281375589546466 | 0.311780061120776 | -0.0807118 | -0.0792161 | -0.0799639 | 17.02931 | SAMN54183639, SAMN54183640 |
| nahumae_19 | 0.281974943786252 | 0.313772537518482 | -0.0827151 | -0.0849014 | -0.0838082 | 19.87845 | SAMN54183641, SAMN54183642 |

**Table S6.** Individual samples (skin swabs, environmental DNA filters) from the SP and SNSM analysed with qPCR and/or CRISPR-diagnostics (CRISPR-Dx). Positive SP sample highlighted in bold.

| Date | lat | long | species | Biogeographic region | qPCR | CRISPR-Dx | Seq. Cap. | eDNA |
|---|---|---|---|---|---|---|---|---|
| 30/8/2024 | 10,3981 | -72,936 | eDNA | SP | negative | NA | NA | Yes |
| 30/8/2024 | 10,3985 | -72,9389 | eDNA | SP | negative | NA | NA | Yes |
| **30/8/2024** | **10,39967** | **-72,9398** | **eDNA** | **SP** | **positive** | **NA** | **Yes** | **Yes** |
| 30/8/2024 | 10,3916 | -72,9666 | eDNA | SP | negative | NA | NA | Yes |
| 30/8/2024 | 10,38827 | -72,9816 | eDNA | SP | negative | NA | NA | Yes |
| 30/8/2024 | 10,39322 | -73,0112 | eDNA | SP | negative | NA | NA | Yes |
| 10/10/2021 | 10,905 | -73,964 | *Atelopus laetissimus* | SNSM | negative | NA | NA | No |
| 10/10/2021 | 10,905 | -73,964 | *Atelopus laetissimus* | SNSM | negative | NA | NA | No |
| 10/10/2021 | 10,905 | -73,964 | *Atelopus laetissimus* | SNSM | negative | NA | NA | No |
| 10/10/2021 | 10,905 | -73,964 | *Atelopus laetissimus* | SNSM | negative | NA | NA | No |
| 10/10/2021 | 10,905 | -73,964 | *Atelopus laetissimus* | SNSM | negative | NA | NA | No |
| 10/10/2021 | 10,905 | -73,964 | *Atelopus laetissimus* | SNSM | negative | NA | NA | No |
| 10/10/2021 | 10,905 | -73,964 | *Atelopus laetissimus* | SNSM | negative | NA | NA | No |
| 10/10/2021 | 10,905 | -73,964 | *Atelopus laetissimus* | SNSM | negative | NA | NA | No |
| 10/10/2021 | 10,905 | -73,964 | *Atelopus laetissimus* | SNSM | negative | NA | NA | No |
| 10/10/2021 | 10,905 | -73,964 | *Atelopus laetissimus* | SNSM | negative | NA | NA | No |
| 10/10/2021 | 10,905 | -73,964 | *Atelopus laetissimus* | SNSM | negative | NA | NA | No |
| 10/10/2021 | 10,905 | -73,964 | *Atelopus laetissimus* | SNSM | negative | NA | NA | No |
| 10/10/2021 | 10,905 | -73,964 | *Atelopus laetissimus* | SNSM | negative | NA | NA | No |
| 10/10/2021 | 10,905 | -73,964 | *Atelopus laetissimus* | SNSM | negative | NA | NA | No |
| 10/10/2021 | 10,905 | -73,964 | *Atelopus laetissimus* | SNSM | negative | NA | NA | No |
| 10/10/2021 | 10,905 | -73,964 | *Atelopus laetissimus* | SNSM | negative | NA | NA | No |
| 10/10/2021 | 10,905 | -73,964 | *Atelopus laetissimus* | SNSM | negative | NA | NA | No |
| 10/10/2021 | 10,905 | -73,964 | *Atelopus laetissimus* | SNSM | negative | NA | NA | No |
| 10/10/2021 | 10,905 | -73,964 | *Atelopus laetissimus* | SNSM | negative | NA | NA | No |
| 10/10/2021 | 10,905 | -73,964 | *Atelopus laetissimus* | SNSM | negative | NA | NA | No |
| 10/10/2021 | 10,905 | -73,964 | *Atelopus laetissimus* | SNSM | negative | NA | NA | No |
| 10/10/2021 | 10,905 | -73,964 | *Atelopus laetissimus* | SNSM | negative | NA | NA | No |
| 10/10/2021 | 10,905 | -73,964 | *Atelopus laetissimus* | SNSM | negative | NA | NA | No |
| 10/10/2021 | 10,905 | -73,964 | *Atelopus laetissimus* | SNSM | negative | NA | NA | No |
| 10/10/2021 | 10,905 | -73,964 | *Atelopus laetissimus* | SNSM | negative | NA | NA | No |
| 10/10/2021 | 10,905 | -73,964 | *Atelopus laetissimus* | SNSM | negative | NA | NA | No |
| 10/10/2021 | 10,905 | -73,964 | *Atelopus laetissimus* | SNSM | negative | NA | NA | No |

| | | | | | | | | |
|---|---|---|---|---|---|---|---|---|
| 10/10/2021 | 10,905 | -73,964 | *Atelopus laetissimus* | SNSM | negative | NA | NA | No |
| 10/10/2021 | 10,905 | -73,964 | *Atelopus laetissimus* | SNSM | negative | NA | NA | No |
| 10/10/2021 | 10,905 | -73,964 | *Atelopus laetissimus* | SNSM | negative | NA | NA | No |
| 10/10/2021 | 10,905 | -73,964 | *Atelopus laetissimus* | SNSM | negative | NA | NA | No |
| 10/10/2021 | 10,905 | -73,964 | *Atelopus laetissimus* | SNSM | negative | NA | NA | No |
| 10/10/2021 | 10,906 | -73,893 | *Atelopus carrikeri* | SNSM | negative | NA | NA | No |
| 10/10/2021 | 10,906 | -73,893 | *Atelopus carrikeri* | SNSM | negative | NA | NA | No |
| 10/10/2021 | 10,906 | -73,893 | *Atelopus carrikeri* | SNSM | negative | NA | NA | No |
| 10/10/2021 | 10,906 | -73,893 | *Atelopus carrikeri* | SNSM | negative | NA | NA | No |
| 10/10/2021 | 10,906 | -73,893 | *Atelopus carrikeri* | SNSM | negative | NA | NA | No |
| 10/10/2021 | 10,906 | -73,893 | *Atelopus carrikeri* | SNSM | negative | NA | NA | No |
| 10/10/2021 | 10,906 | -73,893 | *Atelopus carrikeri* | SNSM | negative | NA | NA | No |
| 10/10/2021 | 10,906 | -73,893 | *Atelopus carrikeri* | SNSM | negative | NA | NA | No |
| 10/10/2021 | 10,906 | -73,893 | *Atelopus carrikeri* | SNSM | negative | NA | NA | No |
| 10/10/2021 | 10,906 | -73,893 | *Atelopus carrikeri* | SNSM | negative | NA | NA | No |
| 10/10/2021 | 10,906 | -73,893 | *Atelopus carrikeri* | SNSM | negative | NA | NA | No |
| 10/10/2021 | 10,906 | -73,893 | *Atelopus carrikeri* | SNSM | negative | NA | NA | No |
| 10/10/2021 | 10,906 | -73,893 | *Atelopus carrikeri* | SNSM | negative | NA | NA | No |
| 10/10/2021 | 10,906 | -73,893 | *Atelopus carrikeri* | SNSM | negative | NA | NA | No |
| 10/10/2021 | 10,906 | -73,893 | *Atelopus carrikeri* | SNSM | negative | NA | NA | No |
| 10/10/2021 | 10,906 | -73,893 | *Atelopus carrikeri* | SNSM | negative | NA | NA | No |
| NA | 10,906 | -73,893 | *Atelopus carrikeri* | SNSM | negative | NA | NA | No |
| NA | 10,906 | -73,893 | *Atelopus carrikeri* | SNSM | negative | NA | NA | No |
| NA | 10,906 | -73,893 | *Atelopus carrikeri* | SNSM | negative | NA | NA | No |
| NA | 10,906 | -73,893 | *Atelopus carrikeri* | SNSM | negative | NA | NA | No |
| NA | 10,906 | -73,893 | *Atelopus carrikeri* | SNSM | negative | NA | NA | No |
| NA | 10,906 | -73,893 | *Atelopus carrikeri* | SNSM | negative | NA | NA | No |
| NA | 10,906 | -73,893 | *Atelopus carrikeri* | SNSM | negative | NA | NA | No |
| NA | 10,906 | -73,893 | *Atelopus carrikeri* | SNSM | negative | NA | NA | No |
| NA | 10,906 | -73,893 | *Atelopus carrikeri* | SNSM | negative | NA | NA | No |
| NA | 10,906 | -73,893 | *Atelopus carrikeri* | SNSM | negative | NA | NA | No |
| 20/9/2021 | 11,106 | -73,463 | *Atelopus aff. laetissimus* | SNSM | negative | NA | NA | No |
| 20/9/2021 | 11,106 | -73,463 | *Atelopus aff. laetissimus* | SNSM | negative | NA | NA | No |
| 20/9/2021 | 11,106 | -73,463 | *Atelopus aff. laetissimus* | SNSM | negative | NA | NA | No |
| 20/9/2021 | 11,106 | -73,463 | *Atelopus aff. laetissimus* | SNSM | negative | NA | NA | No |
| 20/9/2021 | 11,106 | -73,463 | *Atelopus aff. laetissimus* | SNSM | negative | NA | NA | No |
| 20/9/2021 | 11,106 | -73,463 | *Atelopus aff. laetissimus* | SNSM | negative | NA | NA | No |
| 20/9/2021 | 11,106 | -73,463 | *Atelopus aff. laetissimus* | SNSM | negative | NA | NA | No |
| 20/9/2021 | 11,106 | -73,463 | *Atelopus aff. laetissimus* | SNSM | negative | NA | NA | No |

| | | | | | | | | |
|---|---|---|---|---|---|---|---|---|
| 20/9/2021 | 11,106 | -73,463 | *Atelopus aff. laetissimus* | SNSM | negative | NA | NA | No |
| 20/9/2021 | 11,106 | -73,463 | *Atelopus aff. laetissimus* | SNSM | negative | NA | NA | No |
| 20/9/2021 | 11,106 | -73,463 | *Atelopus aff. laetissimus* | SNSM | negative | NA | NA | No |
| 20/9/2021 | 11,106 | -73,463 | *Atelopus aff. laetissimus* | SNSM | negative | NA | NA | No |
| 20/9/2021 | 11,106 | -73,463 | *Atelopus aff. laetissimus* | SNSM | negative | NA | NA | No |
| 20/9/2021 | 11,106 | -73,463 | *Atelopus aff. laetissimus* | SNSM | negative | NA | NA | No |
| 20/9/2021 | 11,106 | -73,463 | *Atelopus aff. laetissimus* | SNSM | negative | NA | NA | No |
| 20/9/2021 | 11,106 | -73,463 | *Atelopus aff. laetissimus* | SNSM | negative | NA | NA | No |
| 20/9/2021 | 11,106 | -73,463 | *Atelopus aff. laetissimus* | SNSM | negative | NA | NA | No |
| 20/9/2021 | 11,106 | -73,463 | *Atelopus aff. laetissimus* | SNSM | negative | NA | NA | No |
| 20/9/2021 | 11,106 | -73,463 | *Atelopus aff. laetissimus* | SNSM | negative | NA | NA | No |
| 20/9/2021 | 11,106 | -73,463 | *Atelopus aff. laetissimus* | SNSM | negative | NA | NA | No |
| 20/9/2021 | 11,106 | -73,463 | *Atelopus aff. laetissimus* | SNSM | negative | NA | NA | No |
| 20/9/2021 | 11,106 | -73,463 | *Atelopus aff. laetissimus* | SNSM | negative | NA | NA | No |
| 20/9/2021 | 11,106 | -73,463 | *Atelopus aff. laetissimus* | SNSM | negative | NA | NA | No |
| 20/9/2021 | 11,106 | -73,463 | *Atelopus aff. laetissimus* | SNSM | negative | NA | NA | No |
| 20/9/2021 | 11,106 | -73,463 | *Atelopus aff. laetissimus* | SNSM | negative | NA | NA | No |
| 20/9/2021 | 11,106 | -73,463 | *Atelopus aff. laetissimus* | SNSM | negative | NA | NA | No |
| 20/9/2021 | 11,106 | -73,463 | *Atelopus aff. laetissimus* | SNSM | negative | NA | NA | No |
| 20/9/2021 | 11,106 | -73,463 | *Atelopus aff. laetissimus* | SNSM | negative | NA | NA | No |
| 20/9/2021 | 11,106 | -73,463 | *Atelopus aff. laetissimus* | SNSM | negative | NA | NA | No |
| 04/05/21 | 11,115 | -74,05 | *Atelopus laetissimus* | SNSM | negative | NA | NA | No |
| 04/05/21 | 11,115 | -74,05 | *Atelopus laetissimus* | SNSM | negative | NA | NA | No |
| 04/05/21 | 11,115 | -74,05 | *Atelopus laetissimus* | SNSM | negative | NA | NA | No |
| 04/05/21 | 11,115 | -74,05 | *Atelopus laetissimus* | SNSM | negative | NA | NA | No |
| 04/05/21 | 11,115 | -74,05 | *Atelopus laetissimus* | SNSM | negative | NA | NA | No |
| 04/05/21 | 11,115 | -74,05 | *Atelopus laetissimus* | SNSM | negative | NA | NA | No |
| 04/05/21 | 11,115 | -74,05 | *Atelopus laetissimus* | SNSM | negative | NA | NA | No |
| 04/05/21 | 11,115 | -74,05 | *Atelopus laetissimus* | SNSM | negative | NA | NA | No |
| 04/05/21 | 11,115 | -74,05 | *Atelopus laetissimus* | SNSM | negative | NA | NA | No |
| 04/05/21 | 11,115 | -74,05 | *Atelopus laetissimus* | SNSM | negative | NA | NA | No |
| 04/05/21 | 11,115 | -74,05 | *Atelopus laetissimus* | SNSM | negative | NA | NA | No |
| 04/05/21 | 11,115 | -74,05 | *Atelopus laetissimus* | SNSM | negative | NA | NA | No |
| 04/05/21 | 11,115 | -74,05 | *Atelopus laetissimus* | SNSM | negative | NA | NA | No |
| 04/05/21 | 11,115 | -74,05 | *Atelopus laetissimus* | SNSM | negative | NA | NA | No |
| 04/05/21 | 11,115 | -74,05 | *Atelopus laetissimus* | SNSM | negative | NA | NA | No |
| 04/05/21 | 11,115 | -74,05 | *Atelopus laetissimus* | SNSM | negative | NA | NA | No |

| | | | | | | | | |
|---|---|---|---|---|---|---|---|---|
| 04/05/21 | 11,115 | -74,05 | *Atelopus laetissimus* | SNSM | negative | NA | NA | No |
| 04/05/21 | 11,115 | -74,05 | *Atelopus laetissimus* | SNSM | negative | NA | NA | No |
| 04/05/21 | 11,115 | -74,05 | *Atelopus laetissimus* | SNSM | negative | NA | NA | No |
| 04/05/21 | 11,115 | -74,05 | *Atelopus laetissimus* | SNSM | negative | NA | NA | No |
| 04/05/21 | 11,115 | -74,05 | *Atelopus laetissimus* | SNSM | negative | NA | NA | No |
| 10/2/2021 | 11,115 | -74,05 | *Atelopus laetissimus* | SNSM | negative | NA | NA | No |
| 10/2/2021 | 11,115 | -74,05 | *Atelopus laetissimus* | SNSM | negative | NA | NA | No |
| 10/2/2021 | 11,115 | -74,05 | *Atelopus laetissimus* | SNSM | negative | NA | NA | No |
| 10/2/2021 | 11,115 | -74,05 | *Atelopus laetissimus* | SNSM | negative | NA | NA | No |
| 10/2/2021 | 11,115 | -74,05 | *Atelopus laetissimus* | SNSM | negative | NA | NA | No |
| 10/2/2021 | 11,115 | -74,05 | *Atelopus laetissimus* | SNSM | negative | NA | NA | No |
| 10/2/2021 | 11,115 | -74,05 | *Atelopus laetissimus* | SNSM | negative | NA | NA | No |
| 10/2/2021 | 11,115 | -74,05 | *Atelopus laetissimus* | SNSM | negative | NA | NA | No |
| 10/2/2021 | 11,115 | -74,05 | *Atelopus laetissimus* | SNSM | negative | NA | NA | No |
| 10/2/2021 | 11,115 | -74,05 | *Atelopus laetissimus* | SNSM | negative | NA | NA | No |
| 10/2/2021 | 11,115 | -74,05 | *Atelopus laetissimus* | SNSM | negative | NA | NA | No |
| 10/2/2021 | 11,115 | -74,05 | *Atelopus laetissimus* | SNSM | negative | NA | NA | No |
| 10/2/2021 | 11,115 | -74,05 | *Atelopus laetissimus* | SNSM | negative | NA | NA | No |
| 04/05/21 | 11,115 | -74,05 | *Atelopus nahumae* | SNSM | negative | NA | NA | No |
| 04/05/21 | 11,115 | -74,05 | *Atelopus nahumae* | SNSM | negative | NA | NA | No |
| 10/2/2021 | 11,115 | -74,05 | *Atelopus nahumae* | SNSM | negative | NA | NA | No |
| 10/2/2021 | 11,115 | -74,05 | *Atelopus nahumae* | SNSM | negative | NA | NA | No |
| 10/2/2021 | 11,115 | -74,05 | *Atelopus nahumae* | SNSM | negative | NA | NA | No |
| 10/2/2021 | 11,115 | -74,05 | *Atelopus nahumae* | SNSM | negative | NA | NA | No |
| 10/2/2021 | 11,115 | -74,05 | *Atelopus nahumae* | SNSM | negative | NA | NA | No |
| 10/2/2021 | 11,115 | -74,05 | *Atelopus nahumae* | SNSM | negative | NA | NA | No |
| 10/2/2021 | 11,115 | -74,05 | *Atelopus nahumae* | SNSM | negative | NA | NA | No |
| 10/2/2021 | 11,115 | -74,05 | *Atelopus nahumae* | SNSM | negative | NA | NA | No |
| 10/2/2021 | 11,115 | -74,05 | *Atelopus nahumae* | SNSM | negative | NA | NA | No |
| 10/2/2021 | 11,115 | -74,05 | *Atelopus nahumae* | SNSM | negative | NA | NA | No |
| 10/2/2021 | 11,115 | -74,05 | *Atelopus laetissimus* | SNSM | negative | NA | NA | No |
| 10/2/2021 | 11,115 | -74,05 | *Atelopus laetissimus* | SNSM | negative | NA | NA | No |
| 5/5/2022 | 11,115 | -74,05 | *Atelopus laetissimus* | SNSM | negative | NA | NA | No |
| 5/5/2022 | 11,115 | -74,05 | *Atelopus laetissimus* | SNSM | negative | NA | NA | No |
| 5/5/2022 | 11,115 | -74,05 | *Atelopus laetissimus* | SNSM | negative | NA | NA | No |
| 5/5/2022 | 11,115 | -74,05 | *Atelopus laetissimus* | SNSM | negative | NA | NA | No |
| 5/5/2022 | 11,115 | -74,05 | *Atelopus laetissimus* | SNSM | negative | NA | NA | No |
| 5/5/2022 | 11,115 | -74,05 | *Atelopus laetissimus* | SNSM | negative | NA | NA | No |
| 5/5/2022 | 11,115 | -74,05 | *Atelopus laetissimus* | SNSM | negative | NA | NA | No |

| | | | | | | | | |
|---|---|---|---|---|---|---|---|---|
| 5/5/2022 | 11,115 | -74,05 | *Atelopus laetissimus* | SNSM | negative | NA | NA | No |
| 5/5/2022 | 11,115 | -74,05 | *Atelopus laetissimus* | SNSM | negative | NA | NA | No |
| 5/5/2022 | 11,115 | -74,05 | *Atelopus laetissimus* | SNSM | negative | NA | NA | No |
| 5/5/2022 | 11,115 | -74,05 | *Atelopus laetissimus* | SNSM | negative | NA | NA | No |
| 5/5/2022 | 11,115 | -74,05 | *Atelopus laetissimus* | SNSM | negative | NA | NA | No |
| 5/5/2022 | 11,115 | -74,05 | *Atelopus laetissimus* | SNSM | negative | NA | NA | No |
| 5/5/2022 | 11,115 | -74,05 | *Atelopus laetissimus* | SNSM | negative | NA | NA | No |
| 5/5/2022 | 11,115 | -74,05 | *Atelopus laetissimus* | SNSM | negative | NA | NA | No |
| 5/5/2022 | 11,115 | -74,05 | *Atelopus laetissimus* | SNSM | negative | NA | NA | No |
| 5/5/2022 | 11,115 | -74,05 | *Atelopus laetissimus* | SNSM | negative | NA | NA | No |
| 5/5/2022 | 11,115 | -74,05 | *Atelopus laetissimus* | SNSM | negative | NA | NA | No |
| 5/5/2022 | 11,115 | -74,05 | *Atelopus laetissimus* | SNSM | negative | NA | NA | No |
| 5/5/2022 | 11,115 | -74,05 | *Atelopus laetissimus* | SNSM | negative | NA | NA | No |
| 5/5/2022 | 11,115 | -74,05 | *Atelopus laetissimus* | SNSM | negative | NA | NA | No |
| 5/5/2022 | 11,115 | -74,05 | *Atelopus laetissimus* | SNSM | negative | NA | NA | No |
| 5/5/2022 | 11,115 | -74,05 | *Atelopus laetissimus* | SNSM | negative | NA | NA | No |
| 5/5/2022 | 11,115 | -74,05 | *Atelopus laetissimus* | SNSM | negative | NA | NA | No |
| 5/5/2022 | 11,115 | -74,05 | *Atelopus laetissimus* | SNSM | negative | NA | NA | No |
| 5/5/2022 | 11,115 | -74,05 | *Atelopus laetissimus* | SNSM | negative | NA | NA | No |
| 5/5/2022 | 11,115 | -74,05 | *Atelopus laetissimus* | SNSM | negative | NA | NA | No |
| 5/5/2022 | 11,115 | -74,05 | *Atelopus laetissimus* | SNSM | negative | NA | NA | No |
| 5/5/2022 | 11,115 | -74,05 | *Atelopus laetissimus* | SNSM | negative | NA | NA | No |
| 5/5/2022 | 11,115 | -74,05 | *Atelopus laetissimus* | SNSM | negative | NA | NA | No |
| 5/5/2022 | 11,115 | -74,05 | *Atelopus laetissimus* | SNSM | negative | NA | NA | No |
| 5/5/2022 | 11,115 | -74,05 | *Atelopus laetissimus* | SNSM | negative | NA | NA | No |
| 5/5/2022 | 11,115 | -74,05 | *Atelopus laetissimus* | SNSM | negative | NA | NA | No |
| 5/5/2022 | 11,115 | -74,05 | *Atelopus laetissimus* | SNSM | negative | NA | NA | No |
| 5/5/2022 | 11,115 | -74,05 | *Atelopus laetissimus* | SNSM | negative | NA | NA | No |
| 5/5/2022 | 11,115 | -74,05 | *Atelopus laetissimus* | SNSM | negative | NA | NA | No |
| 18/3/2022 | 11,115 | -74,05 | *Atelopus laetissimus* | SNSM | negative | NA | NA | No |
| 18/3/2022 | 11,115 | -74,05 | *Atelopus laetissimus* | SNSM | negative | NA | NA | No |
| 18/3/2022 | 11,115 | -74,05 | *Atelopus laetissimus* | SNSM | negative | NA | NA | No |
| 18/3/2022 | 11,115 | -74,05 | *Atelopus laetissimus* | SNSM | negative | NA | NA | No |
| 18/3/2022 | 11,115 | -74,05 | *Atelopus laetissimus* | SNSM | negative | NA | NA | No |
| 18/3/2022 | 11,115 | -74,05 | *Atelopus laetissimus* | SNSM | negative | NA | NA | No |
| 18/3/2022 | 11,115 | -74,05 | *Atelopus laetissimus* | SNSM | negative | NA | NA | No |
| 18/3/2022 | 11,115 | -74,05 | *Atelopus laetissimus* | SNSM | negative | NA | NA | No |

| | | | | | | | | |
|---|---|---|---|---|---|---|---|---|
| 18/3/2022 | 11,115 | -74,05 | *Atelopus laetissimus* | SNSM | negative | NA | NA | No |
| 18/3/2022 | 11,115 | -74,05 | *Atelopus laetissimus* | SNSM | negative | NA | NA | No |
| 18/3/2022 | 11,115 | -74,05 | *Atelopus laetissimus* | SNSM | negative | NA | NA | No |
| 18/3/2022 | 11,115 | -74,05 | *Atelopus laetissimus* | SNSM | negative | NA | NA | No |
| 18/3/2022 | 11,115 | -74,05 | *Atelopus laetissimus* | SNSM | negative | NA | NA | No |
| 18/3/2022 | 11,115 | -74,05 | *Atelopus laetissimus* | SNSM | negative | NA | NA | No |
| 18/3/2022 | 11,115 | -74,05 | *Atelopus laetissimus* | SNSM | negative | NA | NA | No |
| 18/3/2022 | 11,115 | -74,05 | *Atelopus laetissimus* | SNSM | negative | NA | NA | No |
| 18/3/2022 | 11,115 | -74,05 | *Atelopus laetissimus* | SNSM | negative | NA | NA | No |
| 18/3/2022 | 11,115 | -74,05 | *Atelopus laetissimus* | SNSM | negative | NA | NA | No |
| 10/4/2022 | 11,115 | -74,05 | *Atelopus laetissimus* | SNSM | negative | NA | NA | No |
| 10/4/2022 | 11,115 | -74,05 | *Atelopus laetissimus* | SNSM | negative | NA | NA | No |
| 10/4/2022 | 11,115 | -74,05 | *Atelopus laetissimus* | SNSM | negative | NA | NA | No |
| 10/4/2022 | 11,115 | -74,05 | *Atelopus laetissimus* | SNSM | negative | NA | NA | No |
| 10/4/2022 | 11,115 | -74,05 | *Atelopus laetissimus* | SNSM | negative | NA | NA | No |
| 10/4/2022 | 11,115 | -74,05 | *Atelopus laetissimus* | SNSM | negative | NA | NA | No |
| 10/4/2022 | 11,115 | -74,05 | *Atelopus laetissimus* | SNSM | negative | NA | NA | No |
| 10/4/2022 | 11,115 | -74,05 | *Atelopus laetissimus* | SNSM | negative | NA | NA | No |
| 10/4/2022 | 11,115 | -74,05 | *Atelopus laetissimus* | SNSM | negative | NA | NA | No |
| 10/4/2022 | 11,115 | -74,05 | *Atelopus laetissimus* | SNSM | negative | NA | NA | No |
| 10/4/2022 | 11,115 | -74,05 | *Atelopus laetissimus* | SNSM | negative | NA | NA | No |
| 10/4/2022 | 11,115 | -74,05 | *Atelopus laetissimus* | SNSM | negative | NA | NA | No |
| 10/4/2022 | 11,115 | -74,05 | *Atelopus laetissimus* | SNSM | negative | NA | NA | No |
| 10/4/2022 | 11,115 | -74,05 | *Atelopus laetissimus* | SNSM | negative | NA | NA | No |
| 20/6/2021 | 10,905 | -73,964 | *Atelopus laetissimus* | SNSM | negative | NA | NA | No |
| 20/6/2021 | 10,905 | -73,964 | *Atelopus laetissimus* | SNSM | negative | NA | NA | No |
| 20/6/2021 | 10,905 | -73,964 | *Atelopus laetissimus* | SNSM | negative | NA | NA | No |
| 20/6/2021 | 10,905 | -73,964 | *Atelopus laetissimus* | SNSM | negative | NA | NA | No |
| 20/6/2021 | 10,905 | -73,964 | *Atelopus laetissimus* | SNSM | negative | NA | NA | No |
| 20/6/2021 | 10,905 | -73,964 | *Atelopus laetissimus* | SNSM | negative | NA | NA | No |
| 20/6/2021 | 10,905 | -73,964 | *Atelopus laetissimus* | SNSM | negative | NA | NA | No |
| 20/6/2021 | 10,905 | -73,964 | *Atelopus laetissimus* | SNSM | negative | NA | NA | No |
| 20/6/2021 | 10,905 | -73,964 | *Atelopus laetissimus* | SNSM | negative | NA | NA | No |
| 20/6/2021 | 10,905 | -73,964 | *Atelopus laetissimus* | SNSM | negative | NA | NA | No |
| 20/6/2021 | 10,905 | -73,964 | *Atelopus laetissimus* | SNSM | negative | NA | NA | No |
| 20/6/2021 | 10,905 | -73,964 | *Atelopus laetissimus* | SNSM | negative | NA | NA | No |
| 20/6/2021 | 10,905 | -73,964 | *Atelopus laetissimus* | SNSM | negative | NA | NA | No |

| | | | | | | | | |
|---|---|---|---|---|---|---|---|---|
| 20/6/2021 | 10,905 | -73,964 | *Atelopus laetissimus* | SNSM | negative | NA | NA | No |
| 20/6/2021 | 10,905 | -73,964 | *Atelopus laetissimus* | SNSM | negative | NA | NA | No |
| 20/6/2021 | 10,905 | -73,964 | *Atelopus laetissimus* | SNSM | negative | NA | NA | No |
| 20/6/2021 | 10,905 | -73,964 | *Atelopus laetissimus* | SNSM | negative | NA | NA | No |
| 20/6/2021 | 10,905 | -73,964 | *Atelopus laetissimus* | SNSM | negative | NA | NA | No |
| 20/6/2021 | 10,905 | -73,964 | *Atelopus laetissimus* | SNSM | negative | NA | NA | No |
| 20/6/2021 | 10,905 | -73,964 | *Atelopus laetissimus* | SNSM | negative | NA | NA | No |
| 20/6/2021 | 10,905 | -73,964 | *Atelopus laetissimus* | SNSM | negative | NA | NA | No |
| 20/6/2021 | 10,905 | -73,964 | *Atelopus laetissimus* | SNSM | negative | NA | NA | No |
| 20/6/2021 | 10,905 | -73,964 | *Atelopus laetissimus* | SNSM | negative | NA | NA | No |
| 20/6/2021 | 10,905 | -73,964 | *Atelopus laetissimus* | SNSM | negative | NA | NA | No |
| 20/6/2021 | 10,905 | -73,964 | *Atelopus laetissimus* | SNSM | negative | NA | NA | No |
| 20/6/2021 | 10,905 | -73,964 | *Atelopus laetissimus* | SNSM | negative | NA | NA | No |
| 20/6/2021 | 10,905 | -73,964 | *Atelopus laetissimus* | SNSM | negative | NA | NA | No |
| 20/6/2021 | 10,905 | -73,964 | *Atelopus laetissimus* | SNSM | negative | NA | NA | No |
| 20/6/2021 | 10,905 | -73,964 | *Atelopus laetissimus* | SNSM | negative | NA | NA | No |
| 20/6/2021 | 10,905 | -73,964 | *Atelopus laetissimus* | SNSM | negative | NA | NA | No |
| 20/6/2021 | 10,905 | -73,964 | *Atelopus laetissimus* | SNSM | negative | NA | NA | No |
| 20/6/2021 | 10,905 | -73,964 | *Atelopus laetissimus* | SNSM | negative | NA | NA | No |
| 11/3/2021 | 10,905 | -73,964 | *Atelopus laetissimus* | SNSM | negative | NA | NA | No |
| 11/3/2021 | 10,905 | -73,964 | *Atelopus laetissimus* | SNSM | negative | NA | NA | No |
| 11/3/2021 | 10,905 | -73,964 | *Atelopus laetissimus* | SNSM | negative | NA | NA | No |
| 11/3/2021 | 10,905 | -73,964 | *Atelopus laetissimus* | SNSM | negative | NA | NA | No |
| 11/3/2021 | 10,905 | -73,964 | *Atelopus laetissimus* | SNSM | negative | NA | NA | No |
| 11/3/2021 | 10,905 | -73,964 | *Atelopus laetissimus* | SNSM | negative | NA | NA | No |
| 11/3/2021 | 10,905 | -73,964 | *Atelopus laetissimus* | SNSM | negative | NA | NA | No |
| 11/3/2021 | 10,905 | -73,964 | *Atelopus laetissimus* | SNSM | negative | NA | NA | No |
| 11/3/2021 | 10,905 | -73,964 | *Atelopus laetissimus* | SNSM | negative | NA | NA | No |
| 11/3/2021 | 10,905 | -73,964 | *Atelopus laetissimus* | SNSM | negative | NA | NA | No |
| 11/3/2021 | 10,905 | -73,964 | *Atelopus laetissimus* | SNSM | negative | NA | NA | No |
| 11/3/2021 | 10,905 | -73,964 | *Atelopus laetissimus* | SNSM | negative | NA | NA | No |
| 11/3/2021 | 10,905 | -73,964 | *Atelopus laetissimus* | SNSM | negative | NA | NA | No |
| 11/3/2021 | 10,905 | -73,964 | *Atelopus laetissimus* | SNSM | negative | NA | NA | No |
| 11/3/2021 | 10,905 | -73,964 | *Atelopus laetissimus* | SNSM | negative | NA | NA | No |
| 11/3/2021 | 10,905 | -73,964 | *Atelopus laetissimus* | SNSM | negative | NA | NA | No |
| 11/3/2021 | 10,905 | -73,964 | *Atelopus laetissimus* | SNSM | negative | NA | NA | No |
| 11/3/2021 | 10,905 | -73,964 | *Atelopus laetissimus* | SNSM | negative | NA | NA | No |

| | | | | | | | | |
|---|---|---|---|---|---|---|---|---|
| 11/3/2021 | 10,905 | -73,964 | *Atelopus laetissimus* | SNSM | negative | NA | NA | No |
| 11/3/2021 | 10,905 | -73,964 | *Atelopus laetissimus* | SNSM | negative | NA | NA | No |
| 11/3/2021 | 10,905 | -73,964 | *Atelopus laetissimus* | SNSM | negative | NA | NA | No |
| 11/3/2021 | 10,905 | -73,964 | *Atelopus laetissimus* | SNSM | negative | NA | NA | No |
| 11/3/2021 | 10,905 | -73,964 | *Atelopus laetissimus* | SNSM | negative | NA | NA | No |
| 11/3/2021 | 10,905 | -73,964 | *Atelopus laetissimus* | SNSM | negative | NA | NA | No |
| 11/3/2021 | 10,905 | -73,964 | *Atelopus laetissimus* | SNSM | negative | NA | NA | No |
| 11/3/2021 | 10,905 | -73,964 | *Atelopus laetissimus* | SNSM | negative | NA | NA | No |
| 11/3/2021 | 10,905 | -73,964 | *Atelopus laetissimus* | SNSM | negative | NA | NA | No |
| 11/3/2021 | 10,905 | -73,964 | *Atelopus laetissimus* | SNSM | negative | NA | NA | No |
| 11/3/2021 | 10,905 | -73,964 | *Atelopus laetissimus* | SNSM | negative | NA | NA | No |
| 11/3/2021 | 10,905 | -73,964 | *Atelopus laetissimus* | SNSM | negative | NA | NA | No |
| 30/4/2022 | 10,905 | -73,964 | *Atelopus laetissimus* | SNSM | negative | NA | NA | No |
| 30/4/2022 | 10,905 | -73,964 | *Atelopus laetissimus* | SNSM | negative | NA | NA | No |
| 30/4/2022 | 10,905 | -73,964 | *Atelopus laetissimus* | SNSM | negative | NA | NA | No |
| 30/4/2022 | 10,905 | -73,964 | *Atelopus laetissimus* | SNSM | negative | NA | NA | No |
| 30/4/2022 | 10,905 | -73,964 | *Atelopus laetissimus* | SNSM | negative | NA | NA | No |
| 30/4/2022 | 10,905 | -73,964 | *Atelopus laetissimus* | SNSM | negative | NA | NA | No |
| 30/4/2022 | 10,905 | -73,964 | *Atelopus laetissimus* | SNSM | negative | NA | NA | No |
| 30/4/2022 | 10,905 | -73,964 | *Atelopus laetissimus* | SNSM | negative | NA | NA | No |
| 30/4/2022 | 10,905 | -73,964 | *Atelopus laetissimus* | SNSM | negative | NA | NA | No |
| 30/4/2022 | 10,905 | -73,964 | *Atelopus laetissimus* | SNSM | negative | NA | NA | No |
| 30/4/2022 | 10,905 | -73,964 | *Atelopus laetissimus* | SNSM | negative | NA | NA | No |
| 30/4/2022 | 10,905 | -73,964 | *Atelopus laetissimus* | SNSM | negative | NA | NA | No |
| 30/4/2022 | 10,905 | -73,964 | *Atelopus laetissimus* | SNSM | negative | NA | NA | No |
| 30/4/2022 | 10,905 | -73,964 | *Atelopus laetissimus* | SNSM | negative | NA | NA | No |
| 30/4/2022 | 10,905 | -73,964 | *Atelopus laetissimus* | SNSM | negative | NA | NA | No |
| 30/4/2022 | 10,905 | -73,964 | *Atelopus laetissimus* | SNSM | negative | NA | NA | No |
| 30/4/2022 | 10,905 | -73,964 | *Atelopus laetissimus* | SNSM | negative | NA | NA | No |
| 30/4/2022 | 10,905 | -73,964 | *Atelopus laetissimus* | SNSM | negative | NA | NA | No |
| 30/4/2022 | 10,905 | -73,964 | *Atelopus laetissimus* | SNSM | negative | NA | NA | No |
| 30/4/2022 | 10,905 | -73,964 | *Atelopus laetissimus* | SNSM | negative | NA | NA | No |
| 30/4/2022 | 10,905 | -73,964 | *Atelopus laetissimus* | SNSM | negative | NA | NA | No |
| 30/4/2022 | 10,905 | -73,964 | *Atelopus laetissimus* | SNSM | negative | NA | NA | No |
| 30/4/2022 | 10,905 | -73,964 | *Atelopus laetissimus* | SNSM | negative | NA | NA | No |
| 30/4/2022 | 10,905 | -73,964 | *Atelopus laetissimus* | SNSM | negative | NA | NA | No |
| 30/4/2022 | 10,905 | -73,964 | *Atelopus laetissimus* | SNSM | negative | NA | NA | No |

| | | | | | | | | |
|---|---|---|---|---|---|---|---|---|
| 30/4/2022 | 10,905 | -73,964 | *Atelopus laetissimus* | SNSM | negative | NA | NA | No |
| 30/4/2022 | 10,905 | -73,964 | *Atelopus laetissimus* | SNSM | negative | NA | NA | No |
| 30/4/2022 | 10,905 | -73,964 | *Atelopus laetissimus* | SNSM | negative | NA | NA | No |
| 30/4/2022 | 10,905 | -73,964 | *Atelopus laetissimus* | SNSM | negative | NA | NA | No |
| 30/4/2022 | 10,905 | -73,964 | *Atelopus laetissimus* | SNSM | negative | NA | NA | No |
| 30/4/2022 | 10,905 | -73,964 | *Atelopus laetissimus* | SNSM | negative | NA | NA | No |
| 30/4/2022 | 10,905 | -73,964 | *Atelopus laetissimus* | SNSM | negative | NA | NA | No |
| 30/4/2022 | 10,905 | -73,964 | *Atelopus laetissimus* | SNSM | negative | NA | NA | No |
| 30/4/2022 | 10,905 | -73,964 | *Atelopus laetissimus* | SNSM | negative | NA | NA | No |
| NA | 10,905 | -73,964 | *Atelopus laetissimus* | SNSM | negative | NA | NA | No |
| NA | 10,905 | -73,964 | *Atelopus laetissimus* | SNSM | negative | NA | NA | No |
| NA | 10,905 | -73,964 | *Atelopus laetissimus* | SNSM | negative | NA | NA | No |
| NA | 10,905 | -73,964 | *Atelopus laetissimus* | SNSM | negative | NA | NA | No |
| NA | 10,905 | -73,964 | *Atelopus laetissimus* | SNSM | negative | NA | NA | No |
| NA | 10,905 | -73,964 | *Atelopus laetissimus* | SNSM | negative | NA | NA | No |
| NA | 10,905 | -73,964 | *Atelopus laetissimus* | SNSM | negative | NA | NA | No |
| NA | 10,905 | -73,964 | *Atelopus laetissimus* | SNSM | negative | NA | NA | No |
| NA | 10,905 | -73,964 | *Atelopus laetissimus* | SNSM | negative | NA | NA | No |
| NA | 10,905 | -73,964 | *Atelopus laetissimus* | SNSM | negative | NA | NA | No |
| NA | 10,905 | -73,964 | *Atelopus laetissimus* | SNSM | negative | NA | NA | No |
| NA | 10,905 | -73,964 | *Atelopus laetissimus* | SNSM | negative | NA | NA | No |
| NA | 10,905 | -73,964 | *Atelopus laetissimus* | SNSM | negative | NA | NA | No |
| 20/6/2021 | 10,905 | -73,964 | *Atelopus nahumae* | SNSM | negative | NA | NA | No |
| 20/6/2021 | 10,905 | -73,964 | *Atelopus nahumae* | SNSM | negative | NA | NA | No |
| 20/6/2021 | 10,905 | -73,964 | *Atelopus nahumae* | SNSM | negative | NA | NA | No |
| NA | 10,732 | -73,478 | *Atelopus arsyecue* | SNSM | negative | NA | NA | No |
| NA | 10,732 | -73,478 | *Atelopus arsyecue* | SNSM | negative | NA | NA | No |
| NA | 10,732 | -73,478 | *Atelopus arsyecue* | SNSM | negative | NA | NA | No |
| NA | 10,732 | -73,478 | *Atelopus arsyecue* | SNSM | negative | NA | NA | No |
| NA | 10,732 | -73,478 | *Atelopus arsyecue* | SNSM | negative | NA | NA | No |
| NA | 10,732 | -73,478 | *Atelopus arsyecue* | SNSM | negative | NA | NA | No |
| NA | 10,732 | -73,478 | *Atelopus arsyecue* | SNSM | negative | NA | NA | No |
| NA | 10,732 | -73,478 | *Atelopus arsyecue* | SNSM | negative | NA | NA | No |
| NA | 10,732 | -73,478 | *Atelopus arsyecue* | SNSM | negative | NA | NA | No |
| NA | 10,732 | -73,478 | *Atelopus arsyecue* | SNSM | negative | NA | NA | No |
| NA | 10,732 | -73,478 | *Atelopus arsyecue* | SNSM | negative | NA | NA | No |
| NA | 10,732 | -73,478 | *Atelopus arsyecue* | SNSM | negative | NA | NA | No |
| NA | 10,732 | -73,478 | *Atelopus arsyecue* | SNSM | negative | NA | NA | No |
| NA | 10,732 | -73,478 | *Atelopus arsyecue* | SNSM | negative | NA | NA | No |
| NA | 10,732 | -73,478 | *Atelopus arsyecue* | SNSM | negative | NA | NA | No |

| | | | | | | | | |
|---|---|---|---|---|---|---|---|---|
| NA | 10,732 | -73,478 | *Atelopus arsyecue* | SNSM | negative | NA | NA | No |
| NA | 10,732 | -73,478 | *Atelopus arsyecue* | SNSM | negative | NA | NA | No |
| NA | 10,732 | -73,478 | *Atelopus arsyecue* | SNSM | negative | NA | NA | No |
| NA | 10,732 | -73,478 | *Atelopus arsyecue* | SNSM | negative | NA | NA | No |
| NA | 10,732 | -73,478 | *Atelopus arsyecue* | SNSM | negative | NA | NA | No |
| NA | 10,732 | -73,478 | *Atelopus arsyecue* | SNSM | negative | NA | NA | No |
| NA | 10,732 | -73,478 | *Atelopus arsyecue* | SNSM | negative | NA | NA | No |
| NA | 10,732 | -73,478 | *Atelopus arsyecue* | SNSM | negative | NA | NA | No |
| NA | 10,732 | -73,478 | *Atelopus arsyecue* | SNSM | negative | NA | NA | No |
| 20/7/2021 | 10,732 | -73,478 | *Atelopus arsyecue* | SNSM | negative | NA | NA | No |
| 20/7/2021 | 10,732 | -73,478 | *Atelopus arsyecue* | SNSM | negative | NA | NA | No |
| 20/7/2021 | 10,732 | -73,478 | *Atelopus arsyecue* | SNSM | negative | NA | NA | No |
| 20/7/2021 | 10,732 | -73,478 | *Atelopus arsyecue* | SNSM | negative | NA | NA | No |
| 20/7/2021 | 10,732 | -73,478 | *Atelopus arsyecue* | SNSM | negative | NA | NA | No |
| 20/7/2021 | 10,732 | -73,478 | *Atelopus arsyecue* | SNSM | negative | NA | NA | No |
| 20/7/2021 | 10,732 | -73,478 | *Atelopus arsyecue* | SNSM | negative | NA | NA | No |
| 20/7/2021 | 10,732 | -73,478 | *Atelopus arsyecue* | SNSM | negative | NA | NA | No |
| 20/7/2021 | 10,732 | -73,478 | *Atelopus arsyecue* | SNSM | negative | NA | NA | No |
| 20/7/2021 | 10,732 | -73,478 | *Atelopus arsyecue* | SNSM | negative | NA | NA | No |
| 20/7/2021 | 10,732 | -73,478 | *Atelopus arsyecue* | SNSM | negative | NA | NA | No |
| 20/7/2021 | 10,732 | -73,478 | *Atelopus arsyecue* | SNSM | negative | NA | NA | No |
| 20/7/2021 | 10,732 | -73,478 | *Atelopus arsyecue* | SNSM | negative | NA | NA | No |
| 20/7/2021 | 10,732 | -73,478 | *Atelopus arsyecue* | SNSM | negative | NA | NA | No |
| 20/7/2021 | 10,732 | -73,478 | *Atelopus arsyecue* | SNSM | negative | NA | NA | No |
| 20/7/2021 | 10,732 | -73,478 | *Atelopus arsyecue* | SNSM | negative | NA | NA | No |
| 20/7/2021 | 10,732 | -73,478 | *Atelopus arsyecue* | SNSM | negative | NA | NA | No |
| 20/7/2021 | 10,732 | -73,478 | *Atelopus arsyecue* | SNSM | negative | NA | NA | No |
| 20/7/2021 | 10,732 | -73,478 | *Atelopus arsyecue* | SNSM | negative | NA | NA | No |
| 20/7/2021 | 10,732 | -73,478 | *Atelopus arsyecue* | SNSM | negative | NA | NA | No |
| 20/7/2021 | 10,732 | -73,478 | *Atelopus arsyecue* | SNSM | negative | NA | NA | No |
| 20/7/2021 | 10,732 | -73,478 | *Atelopus arsyecue* | SNSM | negative | NA | NA | No |
| 20/7/2021 | 10,732 | -73,478 | *Atelopus arsyecue* | SNSM | negative | NA | NA | No |
| 20/7/2021 | 10,732 | -73,478 | *Atelopus arsyecue* | SNSM | negative | NA | NA | No |
| 20/7/2021 | 10,732 | -73,478 | *Atelopus arsyecue* | SNSM | negative | NA | NA | No |
| 20/7/2021 | 10,732 | -73,478 | *Atelopus arsyecue* | SNSM | negative | NA | NA | No |
| 20/7/2021 | 10,732 | -73,478 | *Atelopus arsyecue* | SNSM | negative | NA | NA | No |
| 20/7/2021 | 10,732 | -73,478 | *Atelopus arsyecue* | SNSM | negative | NA | NA | No |
| 20/7/2021 | 10,732 | -73,478 | *Atelopus arsyecue* | SNSM | negative | NA | NA | No |
| 20/7/2021 | 10,732 | -73,478 | *Atelopus arsyecue* | SNSM | negative | NA | NA | No |
| 20/7/2021 | 10,732 | -73,478 | *Atelopus arsyecue* | SNSM | negative | NA | NA | No |
| 20/7/2021 | 10,732 | -73,478 | *Atelopus arsyecue* | SNSM | negative | NA | NA | No |
| 20/7/2021 | 10,732 | -73,478 | *Atelopus arsyecue* | SNSM | negative | NA | NA | No |
| 20/7/2021 | 10,732 | -73,478 | *Atelopus arsyecue* | SNSM | negative | NA | NA | No |
| 20/7/2021 | 10,732 | -73,478 | *Atelopus arsyecue* | SNSM | negative | NA | NA | No |

| | | | | | | | | |
|---|---|---|---|---|---|---|---|---|
| 26/7/2021 | 10,732 | -73,478 | *Atelopus arsyecue* | SNSM | negative | NA | NA | No |
| 26/7/2021 | 10,732 | -73,478 | *Atelopus arsyecue* | SNSM | negative | NA | NA | No |
| 26/7/2021 | 10,732 | -73,478 | *Atelopus arsyecue* | SNSM | negative | NA | NA | No |
| 26/7/2021 | 10,732 | -73,478 | *Atelopus arsyecue* | SNSM | negative | NA | NA | No |
| 26/7/2021 | 10,732 | -73,478 | *Atelopus arsyecue* | SNSM | negative | NA | NA | No |
| 26/7/2021 | 10,732 | -73,478 | *Atelopus arsyecue* | SNSM | negative | NA | NA | No |
| 26/7/2021 | 10,732 | -73,478 | *Atelopus arsyecue* | SNSM | negative | NA | NA | No |
| 26/7/2021 | 10,732 | -73,478 | *Atelopus arsyecue* | SNSM | negative | NA | NA | No |
| 26/7/2021 | 10,732 | -73,478 | *Atelopus arsyecue* | SNSM | negative | NA | NA | No |
| 26/7/2021 | 10,732 | -73,478 | *Atelopus arsyecue* | SNSM | negative | NA | NA | No |
| 26/7/2021 | 10,732 | -73,478 | *Atelopus arsyecue* | SNSM | negative | NA | NA | No |
| 26/7/2021 | 10,732 | -73,478 | *Atelopus arsyecue* | SNSM | negative | NA | NA | No |
| 26/7/2021 | 10,732 | -73,478 | *Atelopus arsyecue* | SNSM | negative | NA | NA | No |
| 26/7/2021 | 10,732 | -73,478 | *Atelopus arsyecue* | SNSM | negative | NA | NA | No |
| 26/7/2021 | 10,732 | -73,478 | *Atelopus arsyecue* | SNSM | negative | NA | NA | No |
| 26/7/2021 | 10,732 | -73,478 | *Atelopus arsyecue* | SNSM | negative | NA | NA | No |
| 26/7/2021 | 10,732 | -73,478 | *Atelopus arsyecue* | SNSM | negative | NA | NA | No |
| 26/7/2021 | 10,732 | -73,478 | *Atelopus arsyecue* | SNSM | negative | NA | NA | No |
| 26/7/2021 | 10,732 | -73,478 | *Atelopus arsyecue* | SNSM | negative | NA | NA | No |
| 26/7/2021 | 10,732 | -73,478 | *Atelopus arsyecue* | SNSM | negative | NA | NA | No |
| 26/7/2021 | 10,732 | -73,478 | *Atelopus arsyecue* | SNSM | negative | NA | NA | No |
| 26/7/2021 | 10,732 | -73,478 | *Atelopus arsyecue* | SNSM | negative | NA | NA | No |
| 26/7/2021 | 10,732 | -73,478 | *Atelopus arsyecue* | SNSM | negative | NA | NA | No |
| 26/7/2021 | 10,732 | -73,478 | *Atelopus arsyecue* | SNSM | negative | NA | NA | No |
| 26/7/2021 | 10,732 | -73,478 | *Atelopus arsyecue* | SNSM | negative | NA | NA | No |
| 26/7/2021 | 10,732 | -73,478 | *Atelopus arsyecue* | SNSM | negative | NA | NA | No |
| 26/7/2021 | 10,732 | -73,478 | *Atelopus arsyecue* | SNSM | negative | NA | NA | No |
| 26/7/2021 | 10,732 | -73,478 | *Atelopus arsyecue* | SNSM | negative | NA | NA | No |
| 26/7/2021 | 10,732 | -73,478 | *Atelopus arsyecue* | SNSM | negative | NA | NA | No |
| 26/7/2021 | 10,732 | -73,478 | *Atelopus arsyecue* | SNSM | negative | NA | NA | No |
| 26/7/2021 | 10,732 | -73,478 | *Atelopus arsyecue* | SNSM | negative | NA | NA | No |
| 26/7/2021 | 10,732 | -73,478 | *Atelopus arsyecue* | SNSM | negative | NA | NA | No |
| 26/7/2021 | 10,732 | -73,478 | *Atelopus arsyecue* | SNSM | negative | NA | NA | No |
| 26/7/2021 | 10,732 | -73,478 | *Atelopus arsyecue* | SNSM | negative | NA | NA | No |
| 26/7/2021 | 10,732 | -73,478 | *Atelopus arsyecue* | SNSM | negative | NA | NA | No |
| 26/7/2021 | 10,732 | -73,478 | *Atelopus arsyecue* | SNSM | negative | NA | NA | No |
| 26/7/2021 | 10,732 | -73,478 | *Atelopus arsyecue* | SNSM | negative | NA | NA | No |
| 26/7/2021 | 10,732 | -73,478 | *Atelopus arsyecue* | SNSM | negative | NA | NA | No |
| 26/7/2021 | 10,732 | -73,478 | *Atelopus arsyecue* | SNSM | negative | NA | NA | No |
| 26/7/2021 | 10,732 | -73,478 | *Atelopus arsyecue* | SNSM | negative | NA | NA | No |
| 26/7/2021 | 10,732 | -73,478 | *Atelopus arsyecue* | SNSM | negative | NA | NA | No |
| 26/7/2021 | 10,732 | -73,478 | *Atelopus arsyecue* | SNSM | negative | NA | NA | No |
| 26/7/2021 | 10,732 | -73,478 | *Atelopus arsyecue* | SNSM | negative | NA | NA | No |
| 26/7/2021 | 10,732 | -73,478 | *Atelopus arsyecue* | SNSM | negative | NA | NA | No |

| | | | | | | | | |
|---|---|---|---|---|---|---|---|---|
| 26/7/2021 | 10,732 | -73,478 | *Atelopus arsyecue* | SNSM | negative | NA | NA | No |
| 26/7/2021 | 10,732 | -73,478 | *Atelopus arsyecue* | SNSM | negative | NA | NA | No |
| 26/7/2021 | 10,732 | -73,478 | *Atelopus arsyecue* | SNSM | negative | NA | NA | No |
| 26/7/2021 | 10,732 | -73,478 | *Atelopus arsyecue* | SNSM | negative | NA | NA | No |
| 26/7/2021 | 10,732 | -73,478 | *Atelopus arsyecue* | SNSM | negative | NA | NA | No |
| 26/7/2021 | 10,732 | -73,478 | *Atelopus arsyecue* | SNSM | negative | NA | NA | No |
| 26/7/2021 | 10,732 | -73,478 | *Atelopus arsyecue* | SNSM | negative | NA | NA | No |
| 26/7/2021 | 10,732 | -73,478 | *Atelopus arsyecue* | SNSM | negative | NA | NA | No |
| 26/7/2021 | 10,732 | -73,478 | *Atelopus arsyecue* | SNSM | negative | NA | NA | No |
| 26/7/2021 | 10,732 | -73,478 | *Atelopus arsyecue* | SNSM | negative | NA | NA | No |
| 26/7/2021 | 10,732 | -73,478 | *Atelopus arsyecue* | SNSM | negative | NA | NA | No |
| 26/7/2021 | 10,732 | -73,478 | *Atelopus arsyecue* | SNSM | negative | NA | NA | No |
| 26/7/2021 | 10,732 | -73,478 | *Atelopus arsyecue* | SNSM | negative | NA | NA | No |
| 12/1/2024 | 11,117 | -74,086 | *Tachiramantis sp.* | SNSM | negative | NA | NA | No |
| 12/1/2024 | 11,117 | -74,086 | *Cryptobatrachus boulengeri* | SNSM | negative | NA | NA | No |
| 12/1/2024 | 11,117 | -74,086 | *Cryptobatrachus boulengeri* | SNSM | negative | NA | NA | No |
| 12/1/2024 | 11,117 | -74,086 | *Cryptobatrachus boulengeri* | SNSM | negative | NA | NA | No |
| 12/1/2024 | 11,117 | -74,086 | *Tachiramantis sp.* | SNSM | negative | NA | NA | No |
| 12/1/2024 | 11,118 | -74,084 | *Cryptobatrachus boulengeri* | SNSM | negative | NA | NA | No |
| 12/1/2024 | 11,118 | -74,084 | *Cryptobatrachus boulengeri* | SNSM | negative | NA | NA | No |
| 12/1/2024 | 11,118 | -74,084 | *Cryptobatrachus boulengeri* | SNSM | negative | NA | NA | No |
| 12/1/2024 | 11,119 | -74,081 | *Tachiramantis sp.* | SNSM | negative | NA | NA | No |
| 12/1/2024 | 11,119 | -74,081 | *Cryptobatrachus boulengeri* | SNSM | negative | NA | NA | No |
| 12/1/2024 | 11,119 | -74,081 | *Cryptobatrachus boulengeri* | SNSM | negative | NA | NA | No |
| 12/1/2024 | 11,119 | -74,081 | *Cryptobatrachus boulengeri* | SNSM | negative | NA | NA | No |
| 12/1/2024 | 11,119 | -74,081 | *Cryptobatrachus boulengeri* | SNSM | negative | NA | NA | No |
| 12/1/2024 | 11,119 | -74,081 | *Cryptobatrachus boulengeri* | SNSM | negative | NA | NA | No |
| 12/1/2024 | 11,119 | -74,081 | *Cryptobatrachus boulengeri* | SNSM | negative | NA | NA | No |
| 12/1/2024 | 11,119 | -74,081 | *Cryptobatrachus boulengeri* | SNSM | negative | NA | NA | No |
| 12/1/2024 | 11,119 | -74,081 | *Cryptobatrachus boulengeri* | SNSM | negative | NA | NA | No |
| 12/1/2024 | 11,119 | -74,081 | *Cryptobatrachus boulengeri* | SNSM | negative | NA | NA | No |
| 12/1/2024 | 11,119 | -74,081 | *Cryptobatrachus boulengeri* | SNSM | negative | NA | NA | No |
| 12/1/2024 | 11,119 | -74,081 | *Serranobatrachus sanctaemartae* | SNSM | negative | NA | NA | No |
| 12/1/2024 | 11,119 | -74,081 | *Cryptobatrachus boulengeri* | SNSM | negative | NA | NA | No |
| 12/1/2024 | 11,119 | -74,081 | *Cryptobatrachus boulengeri* | SNSM | negative | NA | NA | No |
| 12/1/2024 | 11,119 | -74,081 | *Cryptobatrachus boulengeri* | SNSM | negative | NA | NA | No |
| 12/1/2024 | 11,119 | -74,081 | *Cryptobatrachus boulengeri* | SNSM | negative | NA | NA | No |
| 12/1/2024 | 11,119 | -74,081 | *Cryptobatrachus boulengeri* | SNSM | negative | NA | NA | No |
| 12/1/2024 | 11,119 | -74,081 | *Cryptobatrachus boulengeri* | SNSM | negative | NA | NA | No |
| 12/1/2024 | 11,119 | -74,081 | *Cryptobatrachus boulengeri* | SNSM | negative | NA | NA | No |

| | | | | | | | | |
|---|---|---|---|---|---|---|---|---|
| 12/1/2024 | 11,119 | -74,081 | *Cryptobatrachus boulengeri* | SNSM | negative | NA | NA | No |
| 12/1/2024 | 11,119 | -74,081 | *Cryptobatrachus boulengeri* | SNSM | negative | NA | NA | No |
| 12/1/2024 | 11,119 | -74,081 | *Cryptobatrachus boulengeri* | SNSM | negative | NA | NA | No |
| 12/1/2024 | 11,119 | -74,081 | *Serranobatrachus sanctaemartae* | SNSM | negative | NA | NA | No |
| 12/1/2024 | 11,119 | -74,081 | *Colostethus ruthveni* | SNSM | negative | NA | NA | No |
| 12/1/2024 | 11,119 | -74,081 | *Colostethus ruthveni* | SNSM | negative | NA | NA | No |
| 12/1/2024 | 11,119 | -74,081 | *Colostethus ruthveni* | SNSM | negative | NA | NA | No |
| 12/1/2024 | 11,119 | -74,081 | *Serranobatrachus sanctaemartae* | SNSM | negative | NA | NA | No |
| 12/1/2024 | 11,119 | -74,081 | *Tachiramantis sp.* | SNSM | negative | NA | NA | No |
| 12/1/2024 | 11,119 | -74,081 | *Tachiramantis sp.* | SNSM | negative | NA | NA | No |
| 12/1/2024 | 11,119 | -74,081 | *Serranobatrachus sanctaemartae* | SNSM | negative | NA | NA | No |
| 12/1/2024 | 11,119 | -74,081 | *Tachiramantis sp.* | SNSM | negative | NA | NA | No |
| 12/1/2024 | 11,119 | -74,081 | *Tachiramantis sp.* | SNSM | negative | NA | NA | No |
| 13/1/2024 | 11,12 | -74,093 | *Cryptobatrachus boulengeri* | SNSM | negative | NA | NA | No |
| 13/1/2024 | 11,12 | -74,093 | *Cryptobatrachus boulengeri* | SNSM | negative | NA | NA | No |
| 13/1/2024 | 11,12 | -74,093 | *Cryptobatrachus boulengeri* | SNSM | negative | NA | NA | No |
| 13/1/2024 | 11,12 | -74,093 | *Cryptobatrachus boulengeri* | SNSM | negative | NA | NA | No |
| 13/1/2024 | 11,12 | -74,093 | *Cryptobatrachus boulengeri* | SNSM | negative | NA | NA | No |
| 13/1/2024 | 11,12 | -74,093 | *Cryptobatrachus boulengeri* | SNSM | negative | NA | NA | No |
| 13/1/2024 | 11,12 | -74,093 | *Cryptobatrachus boulengeri* | SNSM | negative | NA | NA | No |
| 13/1/2024 | 11,12 | -74,093 | *Cryptobatrachus boulengeri* | SNSM | negative | NA | NA | No |
| 13/1/2024 | 11,12 | -74,093 | *Cryptobatrachus boulengeri* | SNSM | negative | NA | NA | No |
| 13/1/2024 | 11,12 | -74,093 | *Cryptobatrachus boulengeri* | SNSM | negative | NA | NA | No |
| 13/1/2024 | 11,12 | -74,093 | *Cryptobatrachus boulengeri* | SNSM | negative | NA | NA | No |
| 13/1/2024 | 11,12 | -74,093 | *Cryptobatrachus boulengeri* | SNSM | negative | NA | NA | No |
| 13/1/2024 | 11,12 | -74,093 | *Cryptobatrachus boulengeri* | SNSM | negative | NA | NA | No |
| 13/1/2024 | 11,12 | -74,093 | *Cryptobatrachus boulengeri* | SNSM | negative | NA | NA | No |
| 13/1/2024 | 11,12 | -74,093 | *Cryptobatrachus boulengeri* | SNSM | negative | NA | NA | No |
| 13/1/2024 | 11,12 | -74,093 | *Cryptobatrachus boulengeri* | SNSM | negative | NA | NA | No |
| 13/1/2024 | 11,12 | -74,093 | *Cryptobatrachus boulengeri* | SNSM | negative | NA | NA | No |
| 13/1/2024 | 11,12 | -74,093 | *Cryptobatrachus boulengeri* | SNSM | negative | NA | NA | No |
| 13/1/2024 | 11,12 | -74,093 | *Cryptobatrachus boulengeri* | SNSM | negative | NA | NA | No |
| 13/1/2024 | 11,12 | -74,093 | *Cryptobatrachus boulengeri* | SNSM | negative | NA | NA | No |
| 13/1/2024 | 11,12 | -74,093 | *Cryptobatrachus boulengeri* | SNSM | negative | NA | NA | No |
| 13/1/2024 | 11,12 | -74,093 | *Cryptobatrachus boulengeri* | SNSM | negative | NA | NA | No |
| 13/1/2024 | 11,12 | -74,093 | *Cryptobatrachus boulengeri* | SNSM | negative | NA | NA | No |
| 13/1/2024 | 11,12 | -74,093 | *Cryptobatrachus boulengeri* | SNSM | negative | NA | NA | No |

| | | | | | | | | |
|---|---|---|---|---|---|---|---|---|
| 13/1/2024 | 11,12 | -74,093 | *Cryptobatrachus boulengeri* | SNSM | negative | NA | NA | No |
| 13/1/2024 | 11,12 | -74,093 | *Cryptobatrachus boulengeri* | SNSM | negative | NA | NA | No |
| 13/1/2024 | 11,12 | -74,093 | *Cryptobatrachus boulengeri* | SNSM | negative | NA | NA | No |
| 13/1/2024 | 11,12 | -74,093 | *Cryptobatrachus boulengeri* | SNSM | negative | NA | NA | No |
| 13/1/2024 | 11,12 | -74,093 | *Cryptobatrachus boulengeri* | SNSM | negative | NA | NA | No |
| 14/1/2024 | 11,113 | -74,05 | *Serranobatrachus sanctaemartae* | SNSM | negative | NA | NA | No |
| 14/1/2024 | 11,113 | -74,05 | *Serranobatrachus sanctaemartae* | SNSM | negative | NA | NA | No |
| 14/1/2024 | 11,113 | -74,05 | *Serranobatrachus sanctaemartae* | SNSM | negative | NA | NA | No |
| 14/1/2024 | 11,113 | -74,05 | *Serranobatrachus sanctaemartae* | SNSM | negative | NA | NA | No |
| 14/1/2024 | 11,113 | -74,05 | *Serranobatrachus carmelitae* | SNSM | negative | NA | NA | No |
| 14/1/2024 | 11,113 | -74,05 | *Serranobatrachus cristinae* | SNSM | negative | NA | NA | No |
| 14/1/2024 | 11,113 | -74,05 | *Serranobatrachus sanctaemartae* | SNSM | negative | NA | NA | No |
| 14/1/2024 | 11,113 | -74,05 | *Serranobatrachus megalops* | SNSM | negative | NA | NA | No |
| 14/1/2024 | 11,113 | -74,05 | *Tachiramantis tayrona* | SNSM | negative | NA | NA | No |
| 14/1/2024 | 11,113 | -74,05 | *Serranobatrachus carmelitae* | SNSM | negative | NA | NA | No |
| 14/1/2024 | 11,113 | -74,05 | *Serranobatrachus carmelitae* | SNSM | negative | NA | NA | No |
| 14/1/2024 | 11,113 | -74,05 | *Serranobatrachus sanctaemartae* | SNSM | negative | NA | NA | No |
| 14/1/2024 | 11,113 | -74,05 | *Serranobatrachus megalops* | SNSM | negative | NA | NA | No |
| 14/1/2024 | 11,113 | -74,05 | *Serranobatrachus cristinae* | SNSM | negative | NA | NA | No |
| 14/1/2024 | 11,113 | -74,05 | *Serranobatrachus sanctaemartae* | SNSM | negative | NA | NA | No |
| 14/1/2024 | 11,113 | -74,05 | *Serranobatrachus carmelitae* | SNSM | negative | NA | NA | No |
| 14/1/2024 | 11,113 | -74,05 | *Serranobatrachus delicatus* | SNSM | negative | NA | NA | No |
| 14/1/2024 | 11,113 | -74,05 | *Serranobatrachus sanctaemartae* | SNSM | negative | NA | NA | No |
| 14/1/2024 | 11,113 | -74,05 | *Serranobatrachus sanctaemartae* | SNSM | negative | NA | NA | No |
| 14/1/2024 | 11,113 | -74,05 | *Serranobatrachus carmelitae* | SNSM | negative | NA | NA | No |
| 14/1/2024 | 11,113 | -74,05 | *Serranobatrachus sanctaemartae* | SNSM | negative | NA | NA | No |
| 14/1/2024 | 11,113 | -74,05 | *Serranobatrachus carmelitae* | SNSM | negative | NA | NA | No |
| 14/1/2024 | 11,113 | -74,05 | *Serranobatrachus sanctaemartae* | SNSM | negative | NA | NA | No |
| 14/1/2024 | 11,113 | -74,05 | *Serranobatrachus sanctaemartae* | SNSM | negative | NA | NA | No |
| 14/1/2024 | 11,113 | -74,05 | *Serranobatrachus sanctaemartae* | SNSM | negative | NA | NA | No |
| 14/1/2024 | 11,113 | -74,05 | *Serranobatrachus sanctaemartae* | SNSM | negative | NA | NA | No |
| 14/1/2024 | 11,113 | -74,05 | *Serranobatrachus sanctaemartae* | SNSM | negative | NA | NA | No |
| 14/1/2024 | 11,113 | -74,05 | *Serranobatrachus sanctaemartae* | SNSM | negative | NA | NA | No |
| 14/1/2024 | 11,113 | -74,05 | *Serranobatrachus sanctaemartae* | SNSM | negative | NA | NA | No |
| 14/1/2024 | 11,113 | -74,05 | *Serranobatrachus sanctaemartae* | SNSM | negative | NA | NA | No |
| 14/1/2024 | 11,113 | -74,05 | *Serranobatrachus sanctaemartae* | SNSM | negative | NA | NA | No |
| 15/1/2024 | 11,102 | -74,063 | *Geobatrachus walkeri* | SNSM | negative | NA | NA | No |

| | | | | | | | | |
|---|---|---|---|---|---|---|---|---|
| 15/1/2024 | 11,102 | -74,063 | *Tachiramantis tayrona* | SNSM | negative | NA | NA | No |
| 15/1/2024 | 11,102 | -74,063 | *Tachiramantis tayrona* | SNSM | negative | NA | NA | No |
| 15/1/2024 | 11,102 | -74,063 | *Geobatrachus walkeri* | SNSM | negative | NA | NA | No |
| 15/1/2024 | 11,102 | -74,063 | *Geobatrachus walkeri* | SNSM | negative | NA | NA | No |
| 15/1/2024 | 11,102 | -74,063 | *Geobatrachus walkeri* | SNSM | negative | NA | NA | No |
| 15/1/2024 | 11,102 | -74,063 | *Geobatrachus walkeri* | SNSM | negative | NA | NA | No |
| 15/1/2024 | 11,102 | -74,063 | *Bolitoglossa savagei* | SNSM | negative | NA | NA | No |
| 15/1/2024 | 11,102 | -74,063 | *Bolitoglossa savagei* | SNSM | negative | NA | NA | No |
| 15/1/2024 | 11,102 | -74,063 | *Bolitoglossa savagei* | SNSM | negative | NA | NA | No |
| 15/1/2024 | 11,102 | -74,063 | *Bolitoglossa savagei* | SNSM | negative | NA | NA | No |
| 15/1/2024 | 11,11 | -74,062 | *Serranobatrachus sanctaemartae* | SNSM | negative | NA | NA | No |
| 15/1/2024 | 11,11 | -74,062 | *Serranobatrachus sanctaemartae* | SNSM | negative | NA | NA | No |
| 15/1/2024 | 11,11 | -74,062 | *Serranobatrachus sanctaemartae* | SNSM | negative | NA | NA | No |
| 15/1/2024 | 11,11 | -74,062 | *Serranobatrachus megalops* | SNSM | negative | NA | NA | No |
| 15/1/2024 | 11,11 | -74,062 | *Serranobatrachus megalops* | SNSM | negative | NA | NA | No |
| 15/1/2024 | 11,11 | -74,062 | *Tachiramantis tayrona* | SNSM | negative | NA | NA | No |
| 15/1/2024 | 11,11 | -74,062 | *Serranobatrachus sanctaemartae* | SNSM | negative | NA | NA | No |
| 15/1/2024 | 11,11 | -74,062 | *Serranobatrachus sanctaemartae* | SNSM | negative | NA | NA | No |
| 15/1/2024 | 11,11 | -74,062 | *Serranobatrachus delicatus* | SNSM | negative | NA | NA | No |
| 15/1/2024 | 11,11 | -74,062 | *Serranobatrachus delicatus* | SNSM | negative | NA | NA | No |
| 15/1/2024 | 11,11 | -74,062 | *Serranobatrachus sanctaemartae* | SNSM | negative | NA | NA | No |
| 15/1/2024 | 11,11 | -74,062 | *Serranobatrachus sanctaemartae* | SNSM | negative | NA | NA | No |
| 15/1/2024 | 11,11 | -74,062 | *Serranobatrachus sanctaemartae* | SNSM | negative | NA | NA | No |
| 15/1/2024 | 11,11 | -74,062 | *Serranobatrachus sanctaemartae* | SNSM | negative | NA | NA | No |
| 15/1/2024 | 11,11 | -74,062 | *Serranobatrachus sanctaemartae* | SNSM | negative | NA | NA | No |
| 15/1/2024 | 11,11 | -74,062 | *Serranobatrachus sanctaemartae* | SNSM | negative | NA | NA | No |
| 15/1/2024 | 11,11 | -74,062 | *Serranobatrachus megalops* | SNSM | negative | NA | NA | No |
| 15/1/2024 | 11,11 | -74,062 | *Serranobatrachus sanctaemartae* | SNSM | negative | NA | NA | No |
| 15/1/2024 | 11,11 | -74,062 | *Serranobatrachus sanctaemartae* | SNSM | negative | NA | NA | No |
| 15/1/2024 | 11,11 | -74,062 | *Serranobatrachus sanctaemartae* | SNSM | negative | NA | NA | No |
| 15/1/2024 | 11,11 | -74,062 | *Serranobatrachus sanctaemartae* | SNSM | negative | NA | NA | No |
| 15/1/2024 | 11,11 | -74,062 | *Serranobatrachus sanctaemartae* | SNSM | negative | NA | NA | No |
| 15/1/2024 | 11,11 | -74,062 | *Serranobatrachus megalops* | SNSM | negative | NA | NA | No |
| 15/1/2024 | 11,11 | -74,062 | *Serranobatrachus sanctaemartae* | SNSM | negative | NA | NA | No |
| 15/1/2024 | 11,11 | -74,062 | *Serranobatrachus sanctaemartae* | SNSM | negative | NA | NA | No |
| 15/1/2024 | 11,11 | -74,062 | *Serranobatrachus sanctaemartae* | SNSM | negative | NA | NA | No |
| 15/1/2024 | 11,11 | -74,062 | *Serranobatrachus sanctaemartae* | SNSM | negative | NA | NA | No |

| | | | | | | | | |
|---|---|---|---|---|---|---|---|---|
| 15/1/2024 | 11,11 | -74,062 | *Serranobatrachus delicatus* | SNSM | negative | NA | NA | No |
| 15/1/2024 | 11,105 | -74,063 | *Serranobatrachus insignitus* | SNSM | negative | NA | NA | No |
| 27/1/2024 | 10,768 | -74,027 | environmental DNA | SNSM | negative | NA | NA | Yes |
| 27/1/2024 | 10,768 | -74,027 | environmental DNA | SNSM | negative | NA | NA | Yes |
| 27/1/2024 | 10,768 | -74,027 | *Rhinella horribilis* | SNSM | negative | NA | NA | No |
| 27/1/2024 | 10,768 | -74,027 | *Rhinella horribilis* | SNSM | negative | NA | NA | No |
| 27/1/2024 | 10,768 | -74,027 | *Rhinella horribilis* | SNSM | negative | NA | NA | No |
| 27/1/2024 | 10,768 | -74,027 | *Rhinella horribilis* | SNSM | negative | NA | NA | No |
| 27/1/2024 | 10,768 | -74,027 | *Cryptobatrachus ruthveni* | SNSM | negative | NA | NA | No |
| 27/1/2024 | 10,768 | -74,027 | *Boana platanera* | SNSM | negative | NA | NA | No |
| 27/1/2024 | 10,768 | -74,027 | *Boana platanera* | SNSM | negative | NA | NA | No |
| 27/1/2024 | 10,768 | -74,027 | *Boana platanera* | SNSM | negative | NA | NA | No |
| 27/1/2024 | 10,768 | -74,027 | *Boana platanera* | SNSM | negative | NA | NA | No |
| 27/1/2024 | 10,768 | -74,027 | *Boana platanera* | SNSM | negative | NA | NA | No |
| 27/1/2024 | 10,768 | -74,027 | *Boana platanera* | SNSM | negative | NA | NA | No |
| 27/1/2024 | 10,768 | -74,027 | *Cryptobatrachus ruthveni* | SNSM | negative | NA | NA | No |
| 27/1/2024 | 10,768 | -74,027 | *Boana platanera* | SNSM | negative | NA | NA | No |
| 27/1/2024 | 10,768 | -74,027 | *Boana platanera* | SNSM | negative | NA | NA | No |
| 27/1/2024 | 10,768 | -74,027 | *Boana platanera* | SNSM | negative | NA | NA | No |
| 27/1/2024 | 10,768 | -74,027 | *Boana platanera* | SNSM | negative | NA | NA | No |
| 27/1/2024 | 10,768 | -74,027 | *Boana platanera* | SNSM | negative | NA | NA | No |
| 27/1/2024 | 10,768 | -74,027 | *Boana platanera* | SNSM | negative | NA | NA | No |
| 27/1/2024 | 10,768 | -74,027 | *Boana platanera* | SNSM | negative | NA | NA | No |
| 27/1/2024 | 10,768 | -74,027 | *Boana platanera* | SNSM | negative | NA | NA | No |
| 27/1/2024 | 10,768 | -74,027 | *Lithobates vaillanti* | SNSM | negative | NA | NA | No |
| 27/1/2024 | 10,768 | -74,027 | *Lithobates vaillanti* | SNSM | negative | NA | NA | No |
| 27/1/2024 | 10,768 | -74,027 | *Cryptobatrachus ruthveni* | SNSM | negative | NA | NA | No |
| 27/1/2024 | 10,768 | -74,027 | *Cryptobatrachus ruthveni* | SNSM | negative | NA | NA | No |
| 27/1/2024 | 10,768 | -74,027 | *Cryptobatrachus ruthveni* | SNSM | negative | NA | NA | No |
| 27/1/2024 | 10,768 | -74,027 | *Cryptobatrachus ruthveni* | SNSM | negative | NA | NA | No |
| 27/1/2024 | 10,768 | -74,027 | *Colostethus ruthveni* | SNSM | negative | NA | NA | No |
| 27/1/2024 | 10,768 | -74,027 | *Colostethus ruthveni* | SNSM | negative | NA | NA | No |
| 27/1/2024 | 10,768 | -74,027 | *Colostethus ruthveni* | SNSM | negative | NA | NA | No |
| 27/1/2024 | 10,768 | -74,027 | *Rhinella horribilis* | SNSM | negative | NA | NA | No |
| 27/1/2024 | 10,768 | -74,027 | *Rhinella horribilis* | SNSM | negative | NA | NA | No |
| 27/1/2024 | 10,768 | -74,027 | *Cryptobatrachus ruthveni* | SNSM | negative | NA | NA | No |
| 27/1/2024 | 10,768 | -74,027 | *Cryptobatrachus ruthveni* | SNSM | negative | NA | NA | No |
| 27/1/2024 | 10,768 | -74,027 | *Boana platanera* | SNSM | negative | NA | NA | No |
| 27/1/2024 | 10,768 | -74,027 | *Boana platanera* | SNSM | negative | NA | NA | No |
| 27/1/2024 | 10,768 | -74,027 | *Boana platanera* | SNSM | negative | NA | NA | No |
| 27/1/2024 | 10,768 | -74,027 | *Rhinella horribilis* | SNSM | negative | NA | NA | No |

| | | | | | | | | |
|---|---|---|---|---|---|---|---|---|
| 26/1/2024 | 10,79 | -73,917 | environmental DNA | SNSM | negative | NA | NA | Yes |
| 26/1/2024 | 10,79 | -73,917 | environmental DNA | SNSM | negative | NA | NA | Yes |
| 26/1/2024 | 10,79 | -73,917 | *Cryptobatrachus ruthveni* | SNSM | negative | NA | NA | No |
| 26/1/2024 | 10,79 | -73,917 | *Serranobatrachus insignitus* | SNSM | negative | NA | NA | No |
| 26/1/2024 | 10,79 | -73,917 | *Serranobatrachus insignitus* | SNSM | negative | NA | NA | No |
| 26/1/2024 | 10,79 | -73,917 | *Serranobatrachus insignitus* | SNSM | negative | NA | NA | No |
| 26/1/2024 | 10,79 | -73,917 | *Cryptobatrachus ruthveni* | SNSM | negative | NA | NA | No |
| 26/1/2024 | 10,79 | -73,917 | *Serranobatrachus sanctaemartae* | SNSM | negative | NA | NA | No |
| 26/1/2024 | 10,79 | -73,917 | *Serranobatrachus sanctaemartae* | SNSM | negative | NA | NA | No |
| 26/1/2024 | 10,79 | -73,917 | *Serranobatrachus sanctaemartae* | SNSM | negative | NA | NA | No |
| 26/1/2024 | 10,79 | -73,917 | *Serranobatrachus sanctaemartae* | SNSM | negative | NA | NA | No |
| 26/1/2024 | 10,79 | -73,917 | *Serranobatrachus sanctaemartae* | SNSM | negative | NA | NA | No |
| 26/1/2024 | 10,79 | -73,917 | *Tachiramantis sp.* | SNSM | negative | NA | NA | No |
| 26/1/2024 | 10,79 | -73,917 | *Cryptobatrachus ruthveni* | SNSM | negative | NA | NA | No |
| 26/1/2024 | 10,79 | -73,917 | *Tachiramantis sp.* | SNSM | negative | NA | NA | No |
| 26/1/2024 | 10,79 | -73,917 | *Serranobatrachus sanctaemartae* | SNSM | negative | NA | NA | No |
| 26/1/2024 | 10,79 | -73,917 | *Tachiramantis sp.* | SNSM | negative | NA | NA | No |
| 26/1/2024 | 10,79 | -73,917 | *Tachiramantis sp.* | SNSM | negative | NA | NA | No |
| 26/1/2024 | 10,79 | -73,917 | *Serranobatrachus delicatus* | SNSM | negative | NA | NA | No |
| 26/1/2024 | 10,79 | -73,917 | *Serranobatrachus megalops* | SNSM | negative | NA | NA | No |
| 26/1/2024 | 10,79 | -73,917 | *Serranobatrachus megalops* | SNSM | negative | NA | NA | No |
| 26/1/2024 | 10,79 | -73,917 | *Tachiramantis tayrona* | SNSM | negative | NA | NA | No |
| 26/1/2024 | 10,79 | -73,917 | *Cryptobatrachus ruthveni* | SNSM | negative | NA | NA | No |
| 26/1/2024 | 10,79 | -73,917 | *Cryptobatrachus ruthveni* | SNSM | negative | NA | NA | No |
| 26/1/2024 | 10,79 | -73,917 | *Cryptobatrachus ruthveni* | SNSM | negative | NA | NA | No |
| 26/1/2024 | 10,79 | -73,917 | *Cryptobatrachus ruthveni* | SNSM | negative | NA | NA | No |
| 26/1/2024 | 10,79 | -73,917 | *Cryptobatrachus ruthveni* | SNSM | negative | NA | NA | No |
| 26/1/2024 | 10,79 | -73,917 | *Colostethus ruthveni* | SNSM | negative | NA | NA | No |
| 26/1/2024 | 10,79 | -73,917 | *Serranobatrachus insignitus* | SNSM | negative | NA | NA | No |
| 1/8/2023 | 11,111 | -74,061 | *Bolitoglossa savagei* | SNSM | negative | NA | NA | No |
| 1/8/2023 | 11,111 | -74,061 | *Bolitoglossa savagei* | SNSM | negative | NA | NA | No |
| 1/8/2023 | 11,111 | -74,061 | *Bolitoglossa savagei* | SNSM | negative | NA | NA | No |
| 1/8/2023 | 11,111 | -74,061 | *Bolitoglossa savagei* | SNSM | negative | NA | NA | No |
| 1/8/2023 | 11,111 | -74,061 | *Bolitoglossa savagei* | SNSM | negative | NA | NA | No |
| 1/8/2023 | 11,111 | -74,061 | *Bolitoglossa savagei* | SNSM | negative | NA | NA | No |
| 1/8/2023 | 11,111 | -74,061 | *Bolitoglossa savagei* | SNSM | negative | NA | NA | No |
| 1/8/2023 | 11,111 | -74,061 | *Bolitoglossa savagei* | SNSM | negative | NA | NA | No |
| 1/8/2023 | 11,111 | -74,061 | *Bolitoglossa savagei* | SNSM | negative | NA | NA | No |

| | | | | | | | | |
|---|---|---|---|---|---|---|---|---|
| 1/8/2023 | 11,111 | -74,061 | *Bolitoglossa savagei* | SNSM | negative | NA | NA | No |
| 1/8/2023 | 11,111 | -74,061 | *Bolitoglossa savagei* | SNSM | negative | NA | NA | No |
| 1/8/2023 | 11,111 | -74,061 | *Bolitoglossa savagei* | SNSM | negative | NA | NA | No |
| 1/8/2023 | 11,111 | -74,061 | *Bolitoglossa savagei* | SNSM | negative | NA | NA | No |
| 1/8/2023 | 11,111 | -74,061 | *Bolitoglossa savagei* | SNSM | negative | NA | NA | No |
| 1/8/2023 | 11,111 | -74,061 | *Bolitoglossa savagei* | SNSM | negative | NA | NA | No |
| 1/7/2024 | 10,732 | -73,478 | *Serranobatrachus sp.* | SNSM | negative | NA | NA | No |
| 1/7/2024 | 10,732 | -73,478 | *Serranobatrachus sp.* | SNSM | negative | NA | NA | No |
| 1/7/2024 | 10,732 | -73,478 | *Serranobatrachus sp.* | SNSM | negative | NA | NA | No |
| 1/7/2024 | 10,732 | -73,478 | *Serranobatrachus sp.* | SNSM | negative | NA | NA | No |
| 1/7/2024 | 10,732 | -73,478 | *Serranobatrachus sp.* | SNSM | negative | NA | NA | No |
| 1/7/2024 | 10,732 | -73,478 | *Serranobatrachus sp.* | SNSM | negative | NA | NA | No |
| 1/7/2024 | 10,732 | -73,478 | *Serranobatrachus sp.* | SNSM | negative | NA | NA | No |
| 1/7/2024 | 10,732 | -73,478 | *Serranobatrachus sp.* | SNSM | negative | NA | NA | No |
| 1/7/2024 | 10,732 | -73,478 | *Serranobatrachus sp.* | SNSM | negative | NA | NA | No |
| 1/7/2024 | 10,732 | -73,478 | *Serranobatrachus sp.* | SNSM | negative | NA | NA | No |
| 11/1/2024 | 11,091 | -74,078 | environmental DNA | SNSM | negative | NA | NA | Yes |
| 11/1/2024 | 11,091 | -74,078 | *Cryptobatrachus boulengeri* | SNSM | negative | NA | NA | No |
| 11/1/2024 | 11,091 | -74,078 | *Tachiramantis sp.* | SNSM | negative | NA | NA | No |
| 11/1/2024 | 11,091 | -74,078 | *Tachiramantis sp.* | SNSM | negative | NA | NA | No |
| 11/1/2024 | 11,091 | -74,078 | *Tachiramantis sp.* | SNSM | negative | NA | NA | No |
| 11/1/2024 | 11,091 | -74,078 | *Tachiramantis sp.* | SNSM | negative | NA | NA | No |
| 11/1/2024 | 11,091 | -74,078 | *Tachiramantis sp.* | SNSM | negative | NA | NA | No |
| 11/1/2024 | 11,091 | -74,078 | *Tachiramantis sp.* | SNSM | negative | NA | NA | No |
| 11/1/2024 | 11,091 | -74,078 | *Tachiramantis sp.* | SNSM | negative | NA | NA | No |
| 11/1/2024 | 11,091 | -74,078 | *Tachiramantis sp.* | SNSM | negative | NA | NA | No |
| 11/1/2024 | 11,091 | -74,078 | *Tachiramantis sp.* | SNSM | negative | NA | NA | No |
| 11/1/2024 | 11,091 | -74,078 | *Tachiramantis sp.* | SNSM | negative | NA | NA | No |
| 11/1/2024 | 11,091 | -74,078 | *Cryptobatrachus boulengeri* | SNSM | negative | NA | NA | No |
| 11/1/2024 | 11,091 | -74,078 | *Cryptobatrachus boulengeri* | SNSM | negative | NA | NA | No |
| 11/1/2024 | 11,091 | -74,078 | *Cryptobatrachus boulengeri* | SNSM | negative | NA | NA | No |
| 11/1/2024 | 11,091 | -74,078 | *Cryptobatrachus boulengeri* | SNSM | negative | NA | NA | No |
| 11/1/2024 | 11,091 | -74,078 | *Cryptobatrachus boulengeri* | SNSM | negative | NA | NA | No |
| 11/1/2024 | 11,091 | -74,078 | *Cryptobatrachus boulengeri* | SNSM | negative | NA | NA | No |
| 11/1/2024 | 11,091 | -74,078 | *Cryptobatrachus boulengeri* | SNSM | negative | NA | NA | No |
| 11/1/2024 | 11,091 | -74,078 | *Ikakogi tayrona* | SNSM | negative | NA | NA | No |
| 24/1/2024 | 11,089 | -74,055 | environmental DNA | SNSM | negative | NA | NA | Yes |
| 24/1/2024 | 11,089 | -74,055 | *Cryptobatrachus boulengeri* | SNSM | negative | NA | NA | No |
| 24/1/2024 | 11,089 | -74,055 | *Cryptobatrachus boulengeri* | SNSM | negative | NA | NA | No |

| | | | | | | | | |
|---|---|---|---|---|---|---|---|---|
| 24/1/2024 | 11,089 | -74,055 | *Cryptobatrachus boulengeri* | SNSM | negative | NA | NA | No |
| 24/1/2024 | 11,089 | -74,055 | *Cryptobatrachus boulengeri* | SNSM | negative | NA | NA | No |
| 24/1/2024 | 11,089 | -74,055 | *Cryptobatrachus boulengeri* | SNSM | negative | NA | NA | No |
| 24/1/2024 | 11,089 | -74,055 | *Cryptobatrachus boulengeri* | SNSM | negative | NA | NA | No |
| 24/1/2024 | 11,089 | -74,055 | *Tachiramantis sp.* | SNSM | negative | NA | NA | No |
| 24/1/2024 | 11,089 | -74,055 | *Tachiramantis sp.* | SNSM | negative | NA | NA | No |
| 24/1/2024 | 11,089 | -74,055 | *Tachiramantis sp.* | SNSM | negative | NA | NA | No |
| 24/1/2024 | 11,089 | -74,055 | *Tachiramantis sp.* | SNSM | negative | NA | NA | No |
| 24/1/2024 | 11,089 | -74,055 | *Tachiramantis sp.* | SNSM | negative | NA | NA | No |
| 24/1/2024 | 11,089 | -74,055 | *Cryptobatrachus boulengeri* | SNSM | negative | NA | NA | No |
| 24/1/2024 | 11,089 | -74,055 | *Tachiramantis sp.* | SNSM | negative | NA | NA | No |
| 24/1/2024 | 11,089 | -74,055 | *Tachiramantis sp.* | SNSM | negative | NA | NA | No |
| 24/1/2024 | 11,089 | -74,055 | *Tachiramantis sp.* | SNSM | negative | NA | NA | No |
| 24/1/2024 | 11,089 | -74,055 | *Tachiramantis sp.* | SNSM | negative | NA | NA | No |
| 24/1/2024 | 11,089 | -74,055 | *Tachiramantis sp.* | SNSM | negative | NA | NA | No |
| 24/1/2024 | 11,089 | -74,055 | *Tachiramantis sp.* | SNSM | negative | NA | NA | No |
| 24/1/2024 | 11,089 | -74,055 | *Tachiramantis sp.* | SNSM | negative | NA | NA | No |
| 24/1/2024 | 11,089 | -74,055 | *Tachiramantis sp.* | SNSM | negative | NA | NA | No |
| 24/1/2024 | 11,089 | -74,055 | *Cryptobatrachus boulengeri* | SNSM | negative | NA | NA | No |
| 24/1/2024 | 11,089 | -74,055 | *Cryptobatrachus boulengeri* | SNSM | negative | NA | NA | No |
| 24/1/2024 | 11,089 | -74,055 | *Cryptobatrachus boulengeri* | SNSM | negative | NA | NA | No |
| 24/1/2024 | 11,089 | -74,055 | *Cryptobatrachus boulengeri* | SNSM | negative | NA | NA | No |
| 24/1/2024 | 11,089 | -74,055 | *Cryptobatrachus boulengeri* | SNSM | negative | NA | NA | No |
| 24/1/2024 | 11,089 | -74,055 | *Cryptobatrachus boulengeri* | SNSM | negative | NA | NA | No |
| 24/1/2024 | 11,089 | -74,055 | *Cryptobatrachus boulengeri* | SNSM | negative | NA | NA | No |
| 24/1/2024 | 11,086 | -74,05 | environmental DNA | SNSM | negative | NA | NA | Yes |
| 24/1/2024 | 11,086 | -74,05 | *Cryptobatrachus boulengeri* | SNSM | negative | NA | NA | No |
| 24/1/2024 | 11,086 | -74,05 | *Cryptobatrachus boulengeri* | SNSM | negative | NA | NA | No |
| 24/1/2024 | 11,086 | -74,05 | *Cryptobatrachus boulengeri* | SNSM | negative | NA | NA | No |
| 24/1/2024 | 11,086 | -74,05 | *Cryptobatrachus boulengeri* | SNSM | negative | NA | NA | No |
| 24/1/2024 | 11,086 | -74,05 | *Cryptobatrachus boulengeri* | SNSM | negative | NA | NA | No |
| 24/1/2024 | 11,086 | -74,05 | *Cryptobatrachus boulengeri* | SNSM | negative | NA | NA | No |
| 24/1/2024 | 11,086 | -74,05 | *Cryptobatrachus boulengeri* | SNSM | negative | NA | NA | No |
| 24/1/2024 | 11,086 | -74,05 | *Cryptobatrachus boulengeri* | SNSM | negative | NA | NA | No |
| 24/1/2024 | 11,086 | -74,05 | *Cryptobatrachus boulengeri* | SNSM | negative | NA | NA | No |
| 24/1/2024 | 11,086 | -74,05 | *Cryptobatrachus boulengeri* | SNSM | negative | NA | NA | No |
| 24/1/2024 | 11,086 | -74,05 | *Cryptobatrachus boulengeri* | SNSM | negative | NA | NA | No |
| 24/1/2024 | 11,086 | -74,05 | *Cryptobatrachus boulengeri* | SNSM | negative | NA | NA | No |
| 24/1/2024 | 11,086 | -74,05 | *Cryptobatrachus boulengeri* | SNSM | negative | NA | NA | No |

| | | | | | | | | |
|---|---|---|---|---|---|---|---|---|
| 24/1/2024 | 11,086 | -74,05 | *Cryptobatrachus boulengeri* | SNSM | negative | NA | NA | No |
| 24/1/2024 | 11,086 | -74,05 | *Cryptobatrachus boulengeri* | SNSM | negative | NA | NA | No |
| 24/1/2024 | 11,086 | -74,05 | *Cryptobatrachus boulengeri* | SNSM | negative | NA | NA | No |
| 24/1/2024 | 11,086 | -74,05 | *Tachiramantis sp.* | SNSM | negative | NA | NA | No |
| 24/1/2024 | 11,086 | -74,05 | *Tachiramantis sp.* | SNSM | negative | NA | NA | No |
| 24/1/2024 | 11,086 | -74,05 | *Tachiramantis sp.* | SNSM | negative | NA | NA | No |
| 24/1/2024 | 11,086 | -74,05 | *Tachiramantis sp.* | SNSM | negative | NA | NA | No |
| 24/1/2024 | 11,086 | -74,05 | *Tachiramantis sp.* | SNSM | negative | NA | NA | No |
| 24/1/2024 | 11,086 | -74,05 | *Tachiramantis sp.* | SNSM | negative | NA | NA | No |
| 24/1/2024 | 11,086 | -74,05 | *Cryptobatrachus boulengeri* | SNSM | negative | NA | NA | No |
| 24/1/2024 | 11,086 | -74,05 | *Tachiramantis sp.* | SNSM | negative | NA | NA | No |
| 24/1/2024 | 11,086 | -74,05 | *Cryptobatrachus boulengeri* | SNSM | negative | NA | NA | No |
| 24/1/2024 | 11,086 | -74,05 | *Cryptobatrachus boulengeri* | SNSM | negative | NA | NA | No |
| 24/1/2024 | 11,086 | -74,05 | *Cryptobatrachus boulengeri* | SNSM | negative | NA | NA | No |
| 24/1/2024 | 11,086 | -74,05 | *Cryptobatrachus boulengeri* | SNSM | negative | NA | NA | No |
| 24/1/2024 | 11,086 | -74,05 | *Cryptobatrachus boulengeri* | SNSM | negative | NA | NA | No |
| 24/1/2024 | 11,086 | -74,05 | *Cryptobatrachus boulengeri* | SNSM | negative | NA | NA | No |
| 17/6/2025 | 11,115 | -74,05 | *Atelopus laetissimus* | SNSM | negative | NA | NA | No |
| 17/6/2025 | 11,115 | -74,05 | *Atelopus laetissimus* | SNSM | negative | NA | NA | No |
| 17/6/2025 | 11,115 | -74,05 | *Atelopus laetissimus* | SNSM | negative | NA | NA | No |
| 17/6/2025 | 11,115 | -74,05 | *Atelopus laetissimus* | SNSM | negative | NA | NA | No |
| 17/6/2025 | 11,115 | -74,05 | *Atelopus laetissimus* | SNSM | negative | NA | NA | No |
| 17/6/2025 | 11,115 | -74,05 | *Atelopus laetissimus* | SNSM | negative | NA | NA | No |
| 17/6/2025 | 11,115 | -74,05 | *Atelopus laetissimus* | SNSM | negative | NA | NA | No |
| 17/6/2025 | 11,115 | -74,05 | *Atelopus laetissimus* | SNSM | negative | NA | NA | No |
| 17/6/2025 | 11,115 | -74,05 | *Atelopus laetissimus* | SNSM | negative | NA | NA | No |
| 17/6/2025 | 11,115 | -74,05 | *Atelopus laetissimus* | SNSM | negative | NA | NA | No |
| 17/6/2025 | 11,115 | -74,05 | *Atelopus laetissimus* | SNSM | negative | NA | NA | No |
| 17/6/2025 | 11,115 | -74,05 | *Atelopus laetissimus* | SNSM | negative | NA | NA | No |
| 17/6/2025 | 11,115 | -74,05 | *Atelopus laetissimus* | SNSM | negative | NA | NA | No |
| 17/6/2025 | 11,115 | -74,05 | *Atelopus laetissimus* | SNSM | negative | NA | NA | No |
| 17/6/2025 | 11,115 | -74,05 | *Atelopus laetissimus* | SNSM | negative | NA | NA | No |
| 17/6/2025 | 11,115 | -74,05 | *Atelopus laetissimus* | SNSM | negative | NA | NA | No |
| 17/6/2025 | 11,115 | -74,05 | *Atelopus laetissimus* | SNSM | negative | NA | NA | No |
| 17/6/2025 | 11,115 | -74,05 | *Atelopus laetissimus* | SNSM | negative | NA | NA | No |
| 17/6/2025 | 11,115 | -74,05 | *Atelopus laetissimus* | SNSM | negative | NA | NA | No |
| 17/6/2025 | 11,115 | -74,05 | *Atelopus laetissimus* | SNSM | negative | NA | NA | No |
| 17/6/2025 | 11,115 | -74,05 | *Atelopus laetissimus* | SNSM | negative | NA | NA | No |

| | | | | | | | | |
|---|---|---|---|---|---|---|---|---|
| 17/6/2025 | 11,115 | -74,05 | *Atelopus laetissimus* | SNSM | negative | NA | NA | No |
| 11/1/2024 | 11,089 | -74,055 | environmental DNA | SNSM | negative | NA | NA | Yes |
| 11/1/2024 | 11,106 | -74,088 | *Serranobatrachus sanctaemartae* | SNSM | negative | NA | NA | No |
| 11/1/2024 | 11,106 | -74,088 | *Cryptobatrachus boulengeri* | SNSM | negative | NA | NA | No |
| 11/1/2024 | 11,106 | -74,088 | *Cryptobatrachus boulengeri* | SNSM | negative | NA | NA | No |
| 11/1/2024 | 11,106 | -74,088 | *Cryptobatrachus boulengeri* | SNSM | negative | NA | NA | No |
| 11/1/2024 | 11,106 | -74,088 | *Cryptobatrachus boulengeri* | SNSM | negative | NA | NA | No |
| 11/1/2024 | 11,106 | -74,088 | *Cryptobatrachus boulengeri* | SNSM | negative | NA | NA | No |
| 11/1/2024 | 11,106 | -74,088 | *Cryptobatrachus boulengeri* | SNSM | negative | NA | NA | No |
| 11/1/2024 | 11,106 | -74,088 | *Cryptobatrachus boulengeri* | SNSM | negative | NA | NA | No |
| 11/1/2024 | 11,102 | -74,081 | environmental DNA | SNSM | negative | NA | NA | Yes |
| 11/1/2024 | 11,102 | -74,081 | *Cryptobatrachus boulengeri* | SNSM | negative | NA | NA | No |
| 11/1/2024 | 11,102 | -74,081 | *Cryptobatrachus boulengeri* | SNSM | negative | NA | NA | No |
| 11/1/2024 | 11,102 | -74,081 | *Cryptobatrachus boulengeri* | SNSM | negative | NA | NA | No |
| 11/1/2024 | 11,102 | -74,081 | *Cryptobatrachus boulengeri* | SNSM | negative | NA | NA | No |
| 11/1/2024 | 11,102 | -74,081 | *Cryptobatrachus boulengeri* | SNSM | negative | NA | NA | No |
| 11/1/2024 | 11,102 | -74,081 | *Cryptobatrachus boulengeri* | SNSM | negative | NA | NA | No |
| 11/1/2024 | 11,102 | -74,081 | *Serranobatrachus sanctaemartae* | SNSM | negative | NA | NA | No |
| 11/1/2024 | 11,102 | -74,081 | *Serranobatrachus sanctaemartae* | SNSM | negative | NA | NA | No |
| 11/1/2024 | 11,102 | -74,081 | *Serranobatrachus sanctaemartae* | SNSM | negative | NA | NA | No |
| 11/1/2024 | 11,102 | -74,081 | *Serranobatrachus sanctaemartae* | SNSM | negative | NA | NA | No |
| 11/1/2024 | 11,102 | -74,081 | *Serranobatrachus sanctaemartae* | SNSM | negative | NA | NA | No |
| 11/1/2024 | 11,102 | -74,081 | *Cryptobatrachus boulengeri* | SNSM | negative | NA | NA | No |
| 11/1/2024 | 11,102 | -74,081 | *Serranobatrachus sanctaemartae* | SNSM | negative | NA | NA | No |
| 11/1/2024 | 11,102 | -74,081 | *Serranobatrachus sanctaemartae* | SNSM | negative | NA | NA | No |
| 11/1/2024 | 11,102 | -74,081 | *Serranobatrachus sanctaemartae* | SNSM | negative | NA | NA | No |
| 11/1/2024 | 11,102 | -74,081 | *Serranobatrachus sanctaemartae* | SNSM | negative | NA | NA | No |
| 11/1/2024 | 11,102 | -74,081 | *Serranobatrachus sanctaemartae* | SNSM | negative | NA | NA | No |
| 11/1/2024 | 11,102 | -74,081 | *Serranobatrachus sanctaemartae* | SNSM | negative | NA | NA | No |
| 11/1/2024 | 11,102 | -74,081 | *Serranobatrachus sanctaemartae* | SNSM | negative | NA | NA | No |
| 11/1/2024 | 11,102 | -74,081 | *Serranobatrachus sanctaemartae* | SNSM | negative | NA | NA | No |
| 11/1/2024 | 11,102 | -74,081 | *Serranobatrachus sanctaemartae* | SNSM | negative | NA | NA | No |
| 11/1/2024 | 11,102 | -74,081 | *Tachiramantis sp.* | SNSM | negative | NA | NA | No |
| 11/1/2024 | 11,102 | -74,081 | *Cryptobatrachus boulengeri* | SNSM | negative | NA | NA | No |
| 11/1/2024 | 11,102 | -74,081 | *Ikakogi tayrona* | SNSM | negative | NA | NA | No |
| 11/1/2024 | 11,102 | -74,081 | *Ikakogi tayrona* | SNSM | negative | NA | NA | No |
| 11/1/2024 | 11,102 | -74,081 | *Cryptobatrachus boulengeri* | SNSM | negative | NA | NA | No |
| 11/1/2024 | 11,102 | -74,081 | *Cryptobatrachus boulengeri* | SNSM | negative | NA | NA | No |

| | | | | | | | | |
|---|---|---|---|---|---|---|---|---|
| 11/1/2024 | 11,102 | -74,081 | *Cryptobatrachus boulengeri* | SNSM | negative | NA | NA | No |
| 11/1/2024 | 11,102 | -74,081 | *Cryptobatrachus boulengeri* | SNSM | negative | NA | NA | No |
| 11/1/2024 | 11,102 | -74,081 | *Cryptobatrachus boulengeri* | SNSM | negative | NA | NA | No |
| 11/1/2024 | 11,102 | -74,081 | *Cryptobatrachus boulengeri* | SNSM | negative | NA | NA | No |
| 11/1/2024 | 11,101 | -74,08 | environmental DNA | SNSM | negative | NA | NA | Yes |
| 11/1/2024 | 11,101 | -74,08 | *Serranobatrachus sanctaemartae* | SNSM | negative | NA | NA | No |
| 11/1/2024 | 11,101 | -74,08 | *Serranobatrachus sanctaemartae* | SNSM | negative | NA | NA | No |
| 11/1/2024 | 11,101 | -74,08 | *Serranobatrachus sanctaemartae* | SNSM | negative | NA | NA | No |
| 11/1/2024 | 11,101 | -74,08 | *Serranobatrachus sanctaemartae* | SNSM | negative | NA | NA | No |
| 11/1/2024 | 11,101 | -74,08 | *Serranobatrachus megalops* | SNSM | negative | NA | NA | No |
| 11/1/2024 | 11,101 | -74,08 | *Serranobatrachus sanctaemartae* | SNSM | negative | NA | NA | No |
| 11/1/2024 | 11,101 | -74,08 | *Serranobatrachus sanctaemartae* | SNSM | negative | NA | NA | No |
| 11/1/2024 | 11,101 | -74,08 | *Serranobatrachus sanctaemartae* | SNSM | negative | NA | NA | No |
| 11/1/2024 | 11,101 | -74,08 | *Tachiramantis sp.* | SNSM | negative | NA | NA | No |
| 11/1/2024 | 11,101 | -74,08 | *Serranobatrachus megalops* | SNSM | negative | NA | NA | No |
| 11/1/2024 | 11,101 | -74,08 | *Ikakogi tayrona* | SNSM | negative | NA | NA | No |
| 11/1/2024 | 11,101 | -74,08 | *Serranobatrachus sanctaemartae* | SNSM | negative | NA | NA | No |
| 11/1/2024 | 11,101 | -74,08 | *Cryptobatrachus boulengeri* | SNSM | negative | NA | NA | No |
| 11/1/2024 | 11,101 | -74,08 | *Cryptobatrachus boulengeri* | SNSM | negative | NA | NA | No |
| 11/1/2024 | 11,101 | -74,08 | *Cryptobatrachus boulengeri* | SNSM | negative | NA | NA | No |
| 11/1/2024 | 11,101 | -74,08 | *Cryptobatrachus boulengeri* | SNSM | negative | NA | NA | No |
| 11/1/2024 | 11,101 | -74,08 | *Serranobatrachus sanctaemartae* | SNSM | negative | NA | NA | No |
| 11/1/2024 | 11,101 | -74,08 | *Serranobatrachus sanctaemartae* | SNSM | negative | NA | NA | No |
| 11/1/2024 | 11,101 | -74,08 | *Serranobatrachus sanctaemartae* | SNSM | negative | NA | NA | No |
| 11/1/2024 | 11,101 | -74,08 | *Serranobatrachus sanctaemartae* | SNSM | negative | NA | NA | No |
| 11/1/2024 | 11,101 | -74,08 | *Serranobatrachus sanctaemartae* | SNSM | negative | NA | NA | No |
| 11/1/2024 | 11,101 | -74,08 | *Serranobatrachus sanctaemartae* | SNSM | negative | NA | NA | No |
| 11/1/2024 | 11,101 | -74,08 | *Serranobatrachus sanctaemartae* | SNSM | negative | NA | NA | No |
| 11/1/2024 | 11,093 | -74,079 | *Serranobatrachus sanctaemartae* | SNSM | negative | NA | NA | No |
| 11/1/2024 | 11,093 | -74,079 | *Tachiramantis sp.* | SNSM | negative | NA | NA | No |
| 11/1/2024 | 11,093 | -74,079 | *Tachiramantis sp.* | SNSM | negative | NA | NA | No |
| 11/1/2024 | 11,093 | -74,079 | *Tachiramantis sp.* | SNSM | negative | NA | NA | No |
| 11/1/2024 | 11,093 | -74,079 | *Tachiramantis sp.* | SNSM | negative | NA | NA | No |
| 11/1/2024 | 11,093 | -74,079 | *Tachiramantis sp.* | SNSM | negative | NA | NA | No |
| 11/1/2024 | 11,093 | -74,079 | *Cryptobatrachus boulengeri* | SNSM | negative | NA | NA | No |
| 11/1/2024 | 11,093 | -74,079 | *Cryptobatrachus boulengeri* | SNSM | negative | NA | NA | No |
| 11/1/2024 | 11,093 | -74,079 | *Cryptobatrachus boulengeri* | SNSM | negative | NA | NA | No |
| 11/1/2024 | 11,093 | -74,079 | *Serranobatrachus sanctaemartae* | SNSM | negative | NA | NA | No |

| | | | | | | | | |
|---|---|---|---|---|---|---|---|---|
| 11/1/2024 | 11,093 | -74,079 | *Serranobatrachus sanctaemartae* | SNSM | negative | NA | NA | No |
| 11/1/2024 | 11,093 | -74,079 | *Tachiramantis sp.* | SNSM | negative | NA | NA | No |
| 11/1/2024 | 11,093 | -74,079 | *Tachiramantis sp.* | SNSM | negative | NA | NA | No |
| 11/1/2024 | 11,093 | -74,079 | *Tachiramantis sp.* | SNSM | negative | NA | NA | No |
| 11/1/2024 | 11,093 | -74,079 | *Tachiramantis sp.* | SNSM | negative | NA | NA | No |
| 11/1/2024 | 11,093 | -74,079 | *Tachiramantis sp.* | SNSM | negative | NA | NA | No |
| 11/1/2024 | 11,093 | -74,079 | *Tachiramantis sp.* | SNSM | negative | NA | NA | No |
| 1/7/2024 | 10,732 | -73,478 | *Atelopus arsyecue* | SNSM | negative | NA | NA | No |
| 1/7/2024 | 10,732 | -73,478 | *Atelopus arsyecue* | SNSM | negative | NA | NA | No |
| 1/7/2024 | 10,732 | -73,478 | *Atelopus arsyecue* | SNSM | negative | NA | NA | No |
| 1/7/2024 | 10,732 | -73,478 | *Atelopus arsyecue* | SNSM | negative | NA | NA | No |
| 1/7/2024 | 10,732 | -73,478 | *Atelopus arsyecue* | SNSM | negative | NA | NA | No |
| 1/7/2024 | 10,732 | -73,478 | *Atelopus arsyecue* | SNSM | negative | NA | NA | No |
| 1/7/2024 | 10,732 | -73,478 | *Atelopus arsyecue* | SNSM | negative | NA | NA | No |
| 1/7/2024 | 10,732 | -73,478 | *Atelopus arsyecue* | SNSM | negative | NA | NA | No |
| 1/7/2024 | 10,732 | -73,478 | *Atelopus arsyecue* | SNSM | negative | NA | NA | No |
| 1/7/2024 | 10,732 | -73,478 | *Atelopus arsyecue* | SNSM | negative | NA | NA | No |
| 1/7/2024 | 10,732 | -73,478 | *Atelopus arsyecue* | SNSM | negative | NA | NA | No |
| 1/7/2024 | 10,732 | -73,478 | *Atelopus arsyecue* | SNSM | negative | NA | NA | No |
| 1/7/2024 | 10,732 | -73,478 | *Atelopus arsyecue* | SNSM | negative | NA | NA | No |
| 1/7/2024 | 10,732 | -73,478 | *Atelopus arsyecue* | SNSM | negative | NA | NA | No |
| 1/7/2024 | 10,732 | -73,478 | *Atelopus arsyecue* | SNSM | negative | NA | NA | No |
| 1/7/2024 | 10,732 | -73,478 | *Atelopus arsyecue* | SNSM | negative | NA | NA | No |
| 1/7/2024 | 10,732 | -73,478 | *Atelopus arsyecue* | SNSM | negative | NA | NA | No |
| 1/7/2024 | 10,732 | -73,478 | *Atelopus arsyecue* | SNSM | negative | NA | NA | No |
| 1/7/2024 | 10,732 | -73,478 | *Atelopus arsyecue* | SNSM | negative | NA | NA | No |
| 1/7/2024 | 10,732 | -73,478 | *Atelopus arsyecue* | SNSM | negative | NA | NA | No |
| 1/7/2024 | 10,732 | -73,478 | *Atelopus arsyecue* | SNSM | negative | NA | NA | No |
| 1/7/2024 | 10,732 | -73,478 | *Atelopus arsyecue* | SNSM | negative | NA | NA | No |
| 1/7/2024 | 10,732 | -73,478 | *Atelopus arsyecue* | SNSM | negative | NA | NA | No |
| 1/7/2024 | 10,732 | -73,478 | *Atelopus arsyecue* | SNSM | negative | NA | NA | No |
| 1/7/2024 | 10,732 | -73,478 | *Atelopus arsyecue* | SNSM | negative | NA | NA | No |
| 1/7/2024 | 10,732 | -73,478 | *Atelopus arsyecue* | SNSM | negative | NA | NA | No |
| 1/7/2024 | 10,732 | -73,478 | *Atelopus arsyecue* | SNSM | negative | NA | NA | No |
| 1/7/2024 | 10,732 | -73,478 | *Atelopus arsyecue* | SNSM | negative | NA | NA | No |
| 1/7/2024 | 10,732 | -73,478 | *Atelopus arsyecue* | SNSM | negative | NA | NA | No |
| 1/7/2024 | 10,732 | -73,478 | *Atelopus arsyecue* | SNSM | negative | NA | NA | No |
| 1/7/2024 | 10,732 | -73,478 | *Atelopus arsyecue* | SNSM | negative | NA | NA | No |
| 1/7/2024 | 10,732 | -73,478 | *Atelopus arsyecue* | SNSM | negative | NA | NA | No |
| 1/7/2024 | 10,732 | -73,478 | *Atelopus arsyecue* | SNSM | negative | NA | NA | No |
| 1/7/2024 | 10,732 | -73,478 | *Atelopus arsyecue* | SNSM | negative | NA | NA | No |
| 1/7/2024 | 10,732 | -73,478 | *Atelopus arsyecue* | SNSM | negative | NA | NA | No |
| 1/7/2024 | 10,732 | -73,478 | *Atelopus arsyecue* | SNSM | negative | NA | NA | No |

| | | | | | | | | |
|---|---|---|---|---|---|---|---|---|
| 1/7/2024 | 10,732 | -73,478 | *Bolitoglossa sp. nov.* | SNSM | negative | NA | NA | No |
| 1/7/2024 | 10,732 | -73,478 | *Bolitoglossa sp. nov.* | SNSM | negative | NA | NA | No |
| 3/6/2023 | 10,732 | -73,478 | *Atelopus arsyecue* | SNSM | negative | NA | NA | No |
| 2/6/2023 | 10,732 | -73,478 | *Atelopus arsyecue* | SNSM | negative | NA | NA | No |
| 3/6/2023 | 10,732 | -73,478 | *Atelopus arsyecue* | SNSM | negative | NA | NA | No |
| 2/6/2023 | 10,732 | -73,478 | *Atelopus arsyecue* | SNSM | negative | NA | NA | No |
| 3/6/2023 | 10,732 | -73,478 | *Atelopus arsyecue* | SNSM | negative | NA | NA | No |
| 2/6/2023 | 10,732 | -73,478 | *Atelopus arsyecue* | SNSM | negative | NA | NA | No |
| 3/6/2023 | 10,732 | -73,478 | *Atelopus arsyecue* | SNSM | negative | NA | NA | No |
| 2/6/2022 | 10,732 | -73,478 | *Atelopus arsyecue* | SNSM | negative | NA | NA | No |
| 3/6/2023 | 10,732 | -73,478 | *Atelopus arsyecue* | SNSM | negative | NA | NA | No |
| 2/6/2022 | 10,732 | -73,478 | *Atelopus arsyecue* | SNSM | negative | NA | NA | No |
| 3/6/2023 | 10,732 | -73,478 | *Atelopus arsyecue* | SNSM | negative | NA | NA | No |
| 2/6/2022 | 10,732 | -73,478 | *Atelopus arsyecue* | SNSM | negative | NA | NA | No |
| 3/6/2023 | 10,732 | -73,478 | *Atelopus arsyecue* | SNSM | negative | NA | NA | No |
| 2/6/2022 | 10,732 | -73,478 | *Atelopus arsyecue* | SNSM | negative | NA | NA | No |
| 2/6/2022 | 10,732 | -73,478 | *Atelopus arsyecue* | SNSM | negative | NA | NA | No |
| 2/6/2022 | 10,732 | -73,478 | *Atelopus arsyecue* | SNSM | negative | NA | NA | No |
| 2/6/2023 | 10,732 | -73,478 | *Atelopus arsyecue* | SNSM | negative | NA | NA | No |
| 2/6/2022 | 10,732 | -73,478 | *Atelopus arsyecue* | SNSM | negative | NA | NA | No |
| 3/6/2023 | 10,732 | -73,478 | *Atelopus arsyecue* | SNSM | negative | NA | NA | No |
| 2/6/2023 | 10,732 | -73,478 | *Atelopus arsyecue* | SNSM | negative | NA | NA | No |
| 2/6/2022 | 10,732 | -73,478 | *Atelopus arsyecue* | SNSM | negative | NA | NA | No |
| 2/6/2022 | 10,732 | -73,478 | *Atelopus arsyecue* | SNSM | negative | NA | NA | No |
| 2/6/2022 | 10,732 | -73,478 | *Atelopus arsyecue* | SNSM | negative | NA | NA | No |
| 2/6/2022 | 10,732 | -73,478 | *Atelopus arsyecue* | SNSM | negative | NA | NA | No |
| 2/6/2022 | 10,732 | -73,478 | *Atelopus arsyecue* | SNSM | negative | NA | NA | No |
| 2/6/2022 | 10,732 | -73,478 | *Atelopus arsyecue* | SNSM | negative | NA | NA | No |
| 2/6/2022 | 10,732 | -73,478 | *Atelopus arsyecue* | SNSM | negative | NA | NA | No |
| 2/6/2022 | 10,732 | -73,478 | *Atelopus arsyecue* | SNSM | negative | NA | NA | No |
| 2/6/2022 | 10,732 | -73,478 | *Atelopus arsyecue* | SNSM | negative | NA | NA | No |
| 2/6/2022 | 10,732 | -73,478 | *Atelopus arsyecue* | SNSM | negative | NA | NA | No |
| 3/6/2023 | 10,732 | -73,478 | *Atelopus arsyecue* | SNSM | negative | NA | NA | No |
| 2/6/2023 | 10,732 | -73,478 | *Atelopus arsyecue* | SNSM | negative | NA | NA | No |
| 2/6/2022 | 10,732 | -73,478 | *Atelopus arsyecue* | SNSM | negative | NA | NA | No |
| 2/6/2022 | 10,732 | -73,478 | *Atelopus arsyecue* | SNSM | negative | NA | NA | No |
| 2/6/2022 | 10,732 | -73,478 | *Atelopus arsyecue* | SNSM | negative | NA | NA | No |
| 2/6/2022 | 10,732 | -73,478 | *Atelopus arsyecue* | SNSM | negative | NA | NA | No |
| 2/6/2022 | 10,732 | -73,478 | *Atelopus arsyecue* | SNSM | negative | NA | NA | No |
| 2/6/2022 | 10,732 | -73,478 | *Atelopus arsyecue* | SNSM | negative | NA | NA | No |
| 2/6/2022 | 10,732 | -73,478 | *Atelopus arsyecue* | SNSM | negative | NA | NA | No |
| 2/6/2022 | 10,732 | -73,478 | *Atelopus arsyecue* | SNSM | negative | NA | NA | No |
| 2/6/2022 | 10,732 | -73,478 | *Atelopus arsyecue* | SNSM | negative | NA | NA | No |

| | | | | | | | | |
|---|---|---|---|---|---|---|---|---|
| 2/6/2022 | 10,732 | -73,478 | *Atelopus arsyecue* | SNSM | negative | NA | NA | No |
| 3/6/2023 | 10,732 | -73,478 | *Atelopus arsyecue* | SNSM | negative | NA | NA | No |
| 2/6/2023 | 10,732 | -73,478 | *Atelopus arsyecue* | SNSM | negative | NA | NA | No |
| 2/6/2022 | 10,732 | -73,478 | *Atelopus arsyecue* | SNSM | negative | NA | NA | No |
| 2/6/2022 | 10,732 | -73,478 | *Atelopus arsyecue* | SNSM | negative | NA | NA | No |
| 2/6/2022 | 10,732 | -73,478 | *Atelopus arsyecue* | SNSM | negative | NA | NA | No |
| 2/6/2022 | 10,732 | -73,478 | *Atelopus arsyecue* | SNSM | negative | NA | NA | No |
| 2/6/2022 | 10,732 | -73,478 | *Atelopus arsyecue* | SNSM | negative | NA | NA | No |
| 2/6/2022 | 10,732 | -73,478 | *Atelopus arsyecue* | SNSM | negative | NA | NA | No |
| 2/6/2022 | 10,732 | -73,478 | *Atelopus arsyecue* | SNSM | negative | NA | NA | No |
| 2/6/2022 | 10,732 | -73,478 | *Atelopus arsyecue* | SNSM | negative | NA | NA | No |
| 2/6/2022 | 10,732 | -73,478 | *Atelopus arsyecue* | SNSM | negative | NA | NA | No |
| 2/6/2022 | 10,732 | -73,478 | *Atelopus arsyecue* | SNSM | negative | NA | NA | No |
| 3/6/2023 | 10,732 | -73,478 | *Atelopus arsyecue* | SNSM | negative | NA | NA | No |
| 2/6/2023 | 10,732 | -73,478 | *Atelopus arsyecue* | SNSM | negative | NA | NA | No |
| 2/6/2022 | 10,732 | -73,478 | *Atelopus arsyecue* | SNSM | negative | NA | NA | No |
| 2/6/2022 | 10,732 | -73,478 | *Atelopus arsyecue* | SNSM | negative | NA | NA | No |
| 2/6/2022 | 10,732 | -73,478 | *Atelopus arsyecue* | SNSM | negative | NA | NA | No |
| 2/6/2022 | 10,732 | -73,478 | *Atelopus arsyecue* | SNSM | negative | NA | NA | No |
| 2/6/2022 | 10,732 | -73,478 | *Atelopus arsyecue* | SNSM | negative | NA | NA | No |
| 2/6/2022 | 10,732 | -73,478 | *Atelopus arsyecue* | SNSM | negative | NA | NA | No |
| 2/6/2022 | 10,732 | -73,478 | *Atelopus arsyecue* | SNSM | negative | NA | NA | No |
| 2/6/2022 | 10,732 | -73,478 | *Atelopus arsyecue* | SNSM | negative | NA | NA | No |
| 2/6/2022 | 10,732 | -73,478 | *Atelopus arsyecue* | SNSM | negative | NA | NA | No |
| 2/6/2022 | 10,732 | -73,478 | *Atelopus arsyecue* | SNSM | negative | NA | NA | No |
| 3/6/2023 | 10,732 | -73,478 | *Atelopus arsyecue* | SNSM | negative | NA | NA | No |
| 2/6/2023 | 10,732 | -73,478 | *Atelopus arsyecue* | SNSM | negative | NA | NA | No |
| 3/6/2023 | 10,732 | -73,478 | *Atelopus arsyecue* | SNSM | negative | NA | NA | No |
| 2/6/2023 | 10,732 | -73,478 | *Atelopus arsyecue* | SNSM | negative | NA | NA | No |
| 3/6/2023 | 10,732 | -73,478 | *Atelopus arsyecue* | SNSM | negative | NA | NA | No |
| 2/6/2023 | 10,732 | -73,478 | *Atelopus arsyecue* | SNSM | negative | NA | NA | No |
| 3/6/2023 | 10,732 | -73,478 | *Atelopus arsyecue* | SNSM | negative | NA | NA | No |
| 2/6/2023 | 10,732 | -73,478 | *Atelopus arsyecue* | SNSM | negative | NA | NA | No |
| 1/7/2024 | 10,732 | -73,478 | *Serranobatrachus sp.* | SNSM | negative | NA | NA | No |
| 1/7/2024 | 10,732 | -73,478 | *Serranobatrachus sp.* | SNSM | negative | NA | NA | No |
| 1/7/2024 | 10,732 | -73,478 | *Serranobatrachus sp.* | SNSM | negative | NA | NA | No |
| 1/7/2024 | 10,732 | -73,478 | *Serranobatrachus sp.* | SNSM | negative | NA | NA | No |
| 1/7/2024 | 10,732 | -73,478 | *Serranobatrachus sp.* | SNSM | negative | NA | NA | No |
| 1/7/2024 | 10,732 | -73,478 | *Serranobatrachus sp.* | SNSM | negative | NA | NA | No |
| 1/7/2024 | 10,732 | -73,478 | *Serranobatrachus sp.* | SNSM | negative | NA | NA | No |
| 1/7/2024 | 10,732 | -73,478 | *Serranobatrachus sp.* | SNSM | negative | NA | NA | No |
| 1/7/2024 | 10,732 | -73,478 | *Serranobatrachus sp.* | SNSM | negative | NA | NA | No |

| | | | | | | | | |
|---|---|---|---|---|---|---|---|---|
| 19/1/2024 | 10,893 | -74,027 | environmental DNA | SNSM | negative | NA | NA | Yes |
| 19/1/2024 | 10,893 | -74,027 | environmental DNA | SNSM | negative | NA | NA | Yes |
| 20/1/2024 | 10,889 | -74,021 | environmental DNA | SNSM | negative | NA | NA | Yes |
| 21/1/2024 | 10,9 | -73,972 | environmental DNA | SNSM | negative | NA | NA | Yes |
| 29/7/2023 | 10,905 | -73,964 | *Atelopus laetissimus* | SNSM | negative | NA | NA | No |
| 20/6/2023 | 10,905 | -73,964 | *Atelopus laetissimus* | SNSM | negative | NA | NA | No |
| 19/1/2024 | 10,893 | -74,027 | *Serranobatrachus sanctaemartae* | SNSM | negative | NA | NA | No |
| 19/1/2024 | 10,893 | -74,027 | *Serranobatrachus sanctaemartae* | SNSM | negative | NA | NA | No |
| 19/1/2024 | 10,893 | -74,027 | *Serranobatrachus sanctaemartae* | SNSM | negative | NA | NA | No |
| 19/1/2024 | 10,893 | -74,027 | *Serranobatrachus delicatus* | SNSM | negative | NA | NA | No |
| 19/1/2024 | 10,893 | -74,027 | *Serranobatrachus insignitus* | SNSM | negative | NA | NA | No |
| 19/1/2024 | 10,893 | -74,027 | *Bolitoglossa savagei* | SNSM | negative | NA | NA | No |
| 19/1/2024 | 10,893 | -74,027 | *Serranobatrachus sanctaemartae* | SNSM | negative | NA | NA | No |
| 19/1/2024 | 10,893 | -74,027 | *Cryptobatrachus ruthveni* | SNSM | negative | NA | NA | No |
| 19/1/2024 | 10,893 | -74,027 | *Serranobatrachus sanctaemartae* | SNSM | negative | NA | NA | No |
| 19/1/2024 | 10,893 | -74,027 | *Serranobatrachus sanctaemartae* | SNSM | negative | NA | NA | No |
| 19/1/2024 | 10,893 | -74,027 | *Serranobatrachus sanctaemartae* | SNSM | negative | NA | NA | No |
| 19/1/2024 | 10,893 | -74,027 | *Serranobatrachus sanctaemartae* | SNSM | negative | NA | NA | No |
| 19/1/2024 | 10,893 | -74,027 | *Bolitoglossa savagei* | SNSM | negative | NA | NA | No |
| 19/1/2024 | 10,893 | -74,027 | *Serranobatrachus megalops* | SNSM | negative | NA | NA | No |
| 19/1/2024 | 10,893 | -74,027 | *Serranobatrachus sanctaemartae* | SNSM | negative | NA | NA | No |
| 19/1/2024 | 10,893 | -74,027 | *Bolitoglossa savagei* | SNSM | negative | NA | NA | No |
| 19/1/2024 | 10,893 | -74,027 | *Serranobatrachus insignitus* | SNSM | negative | NA | NA | No |
| 19/1/2024 | 10,893 | -74,027 | *Serranobatrachus sanctaemartae* | SNSM | negative | NA | NA | No |
| 19/1/2024 | 10,893 | -74,027 | *Serranobatrachus delicatus* | SNSM | negative | NA | NA | No |
| 19/1/2024 | 10,893 | -74,027 | *Cryptobatrachus ruthveni* | SNSM | negative | NA | NA | No |
| 19/1/2024 | 10,893 | -74,027 | *Serranobatrachus sanctaemartae* | SNSM | negative | NA | NA | No |
| 19/1/2024 | 10,893 | -74,027 | *Serranobatrachus insignitus* | SNSM | negative | NA | NA | No |
| 19/1/2024 | 10,893 | -74,027 | *Serranobatrachus sanctaemartae* | SNSM | negative | NA | NA | No |
| 19/1/2024 | 10,893 | -74,027 | *Serranobatrachus sanctaemartae* | SNSM | negative | NA | NA | No |
| 19/1/2024 | 10,893 | -74,027 | *Serranobatrachus sanctaemartae* | SNSM | negative | NA | NA | No |
| 19/1/2024 | 10,893 | -74,027 | *Serranobatrachus insignitus* | SNSM | negative | NA | NA | No |
| 19/1/2024 | 10,893 | -74,027 | *Cryptobatrachus ruthveni* | SNSM | negative | NA | NA | No |
| 19/1/2024 | 10,893 | -74,027 | *Cryptobatrachus ruthveni* | SNSM | negative | NA | NA | No |
| 19/1/2024 | 10,893 | -74,027 | *Cryptobatrachus ruthveni* | SNSM | negative | NA | NA | No |
| 19/1/2024 | 10,893 | -74,027 | *Serranobatrachus insignitus* | SNSM | negative | NA | NA | No |
| 19/1/2024 | 10,893 | -74,027 | *Cryptobatrachus ruthveni* | SNSM | negative | NA | NA | No |

| | | | | | | | | |
|---|---|---|---|---|---|---|---|---|
| 19/1/2024 | 10,893 | -74,027 | *Serranobatrachus delicatus* | SNSM | negative | NA | NA | No |
| 19/1/2024 | 10,893 | -74,027 | *Serranobatrachus sanctaemartae* | SNSM | negative | NA | NA | No |
| 19/1/2024 | 10,893 | -74,027 | *Serranobatrachus sanctaemartae* | SNSM | negative | NA | NA | No |
| 19/1/2024 | 10,893 | -74,027 | *Serranobatrachus sanctaemartae* | SNSM | negative | NA | NA | No |
| 19/1/2024 | 10,893 | -74,027 | *Serranobatrachus megalops* | SNSM | negative | NA | NA | No |
| 19/1/2024 | 10,893 | -74,027 | *Bolitoglossa savagei* | SNSM | negative | NA | NA | No |
| 19/1/2024 | 10,893 | -74,027 | *Bolitoglossa savagei* | SNSM | negative | NA | NA | No |
| 19/1/2024 | 10,893 | -74,027 | *Bolitoglossa savagei* | SNSM | negative | NA | NA | No |
| 29/7/2023 | 10,905 | -73,964 | *Atelopus laetissimus* | SNSM | negative | NA | NA | No |
| 20/6/2023 | 10,905 | -73,964 | *Atelopus laetissimus* | SNSM | negative | NA | NA | No |
| 20/6/2023 | 10,905 | -73,964 | *Atelopus laetissimus* | SNSM | negative | NA | NA | No |
| 20/6/2023 | 10,905 | -73,964 | *Atelopus laetissimus* | SNSM | negative | NA | NA | No |
| 20/6/2023 | 10,905 | -73,964 | *Atelopus laetissimus* | SNSM | negative | NA | NA | No |
| 20/6/2023 | 10,905 | -73,964 | *Atelopus laetissimus* | SNSM | negative | NA | NA | No |
| 20/6/2023 | 10,905 | -73,964 | *Atelopus laetissimus* | SNSM | negative | NA | NA | No |
| 29/7/2023 | 10,905 | -73,964 | *Atelopus laetissimus* | SNSM | negative | NA | NA | No |
| 20/6/2023 | 10,905 | -73,964 | *Atelopus laetissimus* | SNSM | negative | NA | NA | No |
| 20/1/2024 | 10,889 | -74,021 | *Cryptobatrachus ruthveni* | SNSM | negative | NA | NA | No |
| 20/1/2024 | 10,889 | -74,021 | *Cryptobatrachus ruthveni* | SNSM | negative | NA | NA | No |
| 20/1/2024 | 10,889 | -74,021 | *Serranobatrachus sanctaemartae* | SNSM | negative | NA | NA | No |
| 20/1/2024 | 10,889 | -74,021 | *Serranobatrachus sanctaemartae* | SNSM | negative | NA | NA | No |
| 20/1/2024 | 10,889 | -74,021 | *Serranobatrachus sanctaemartae* | SNSM | negative | NA | NA | No |
| 20/1/2024 | 10,889 | -74,021 | *Cryptobatrachus ruthveni* | SNSM | negative | NA | NA | No |
| 20/1/2024 | 10,889 | -74,021 | *Cryptobatrachus ruthveni* | SNSM | negative | NA | NA | No |
| 20/1/2024 | 10,889 | -74,021 | *Cryptobatrachus ruthveni* | SNSM | negative | NA | NA | No |
| 20/1/2024 | 10,889 | -74,021 | *Cryptobatrachus ruthveni* | SNSM | negative | NA | NA | No |
| 20/1/2024 | 10,889 | -74,021 | *Cryptobatrachus ruthveni* | SNSM | negative | NA | NA | No |
| 20/1/2024 | 10,889 | -74,021 | *Cryptobatrachus ruthveni* | SNSM | negative | NA | NA | No |
| 20/1/2024 | 10,889 | -74,021 | *Cryptobatrachus ruthveni* | SNSM | negative | NA | NA | No |
| 20/1/2024 | 10,889 | -74,021 | *Serranobatrachus sanctaemartae* | SNSM | negative | NA | NA | No |
| 20/1/2024 | 10,889 | -74,021 | *Tachiramantis tayrona* | SNSM | negative | NA | NA | No |
| 20/1/2024 | 10,889 | -74,021 | *Serranobatrachus sanctaemartae* | SNSM | negative | NA | NA | No |
| 20/1/2024 | 10,889 | -74,021 | *Cryptobatrachus ruthveni* | SNSM | negative | NA | NA | No |
| 20/1/2024 | 10,889 | -74,021 | *Cryptobatrachus ruthveni* | SNSM | negative | NA | NA | No |
| 20/1/2024 | 10,889 | -74,021 | *Cryptobatrachus ruthveni* | SNSM | negative | NA | NA | No |
| 20/1/2024 | 10,889 | -74,021 | *Cryptobatrachus ruthveni* | SNSM | negative | NA | NA | No |
| 20/1/2024 | 10,889 | -74,021 | *Cryptobatrachus ruthveni* | SNSM | negative | NA | NA | No |

| | | | | | | | | |
|---|---|---|---|---|---|---|---|---|
| 20/1/2024 | 10,889 | -74,021 | *Cryptobatrachus ruthveni* | SNSM | negative | NA | NA | No |
| 20/1/2024 | 10,889 | -74,021 | *Serranobatrachus sanctaemartae* | SNSM | negative | NA | NA | No |
| 20/1/2024 | 10,889 | -74,021 | *Cryptobatrachus ruthveni* | SNSM | negative | NA | NA | No |
| 20/1/2024 | 10,889 | -74,021 | *Cryptobatrachus ruthveni* | SNSM | negative | NA | NA | No |
| 20/1/2024 | 10,889 | -74,021 | *Cryptobatrachus ruthveni* | SNSM | negative | NA | NA | No |
| 20/1/2024 | 10,889 | -74,021 | *Cryptobatrachus ruthveni* | SNSM | negative | NA | NA | No |
| 20/1/2024 | 10,889 | -74,021 | *Serranobatrachus megalops* | SNSM | negative | NA | NA | No |
| 20/1/2024 | 10,889 | -74,021 | *Serranobatrachus sanctaemartae* | SNSM | negative | NA | NA | No |
| 20/1/2024 | 10,889 | -74,021 | *Serranobatrachus insignitus* | SNSM | negative | NA | NA | No |
| 20/1/2024 | 10,889 | -74,021 | *Cryptobatrachus ruthveni* | SNSM | negative | NA | NA | No |
| 20/1/2024 | 10,889 | -74,021 | *Serranobatrachus sanctaemartae* | SNSM | negative | NA | NA | No |
| 1/7/2024 | 10,905 | -73,964 | *Atelopus laetissimus* | SNSM | negative | NA | NA | No |
| 1/7/2024 | 10,905 | -73,964 | *Atelopus laetissimus* | SNSM | negative | NA | NA | No |
| 1/7/2024 | 10,905 | -73,964 | *Atelopus laetissimus* | SNSM | negative | NA | NA | No |
| 1/7/2024 | 10,905 | -73,964 | *Atelopus laetissimus* | SNSM | negative | NA | NA | No |
| 1/7/2024 | 10,905 | -73,964 | *Atelopus laetissimus* | SNSM | negative | NA | NA | No |
| 1/7/2024 | 10,905 | -73,964 | *Atelopus laetissimus* | SNSM | negative | NA | NA | No |
| 1/7/2024 | 10,905 | -73,964 | *Atelopus laetissimus* | SNSM | negative | NA | NA | No |
| 1/7/2024 | 10,905 | -73,964 | *Atelopus laetissimus* | SNSM | negative | NA | NA | No |
| 1/7/2024 | 10,905 | -73,964 | *Atelopus laetissimus* | SNSM | negative | NA | NA | No |
| 1/7/2024 | 10,905 | -73,964 | *Atelopus laetissimus* | SNSM | negative | NA | NA | No |
| 1/7/2024 | 10,905 | -73,964 | *Atelopus laetissimus* | SNSM | negative | NA | NA | No |
| 1/7/2024 | 10,905 | -73,964 | *Atelopus laetissimus* | SNSM | negative | NA | NA | No |
| 1/7/2024 | 10,905 | -73,964 | *Atelopus laetissimus* | SNSM | negative | NA | NA | No |
| 1/7/2024 | 10,905 | -73,964 | *Atelopus laetissimus* | SNSM | negative | NA | NA | No |
| 1/7/2024 | 10,905 | -73,964 | *Atelopus laetissimus* | SNSM | negative | NA | NA | No |
| 1/7/2024 | 10,905 | -73,964 | *Atelopus laetissimus* | SNSM | negative | NA | NA | No |
| 1/7/2024 | 10,905 | -73,964 | *Atelopus laetissimus* | SNSM | negative | NA | NA | No |
| 1/7/2024 | 10,905 | -73,964 | *Atelopus laetissimus* | SNSM | negative | NA | NA | No |
| 1/7/2024 | 10,905 | -73,964 | *Atelopus laetissimus* | SNSM | negative | NA | NA | No |
| 1/7/2024 | 10,905 | -73,964 | *Atelopus laetissimus* | SNSM | negative | NA | NA | No |
| 1/7/2024 | 10,905 | -73,964 | *Atelopus laetissimus* | SNSM | negative | NA | NA | No |
| 1/7/2024 | 10,905 | -73,964 | *Atelopus laetissimus* | SNSM | negative | NA | NA | No |
| 1/7/2024 | 10,905 | -73,964 | *Atelopus laetissimus* | SNSM | negative | NA | NA | No |
| 1/7/2024 | 10,905 | -73,964 | *Atelopus laetissimus* | SNSM | negative | NA | NA | No |
| 1/7/2024 | 10,905 | -73,964 | *Atelopus laetissimus* | SNSM | negative | NA | NA | No |
| 1/7/2024 | 10,905 | -73,964 | *Atelopus laetissimus* | SNSM | negative | NA | NA | No |
| 1/7/2024 | 10,905 | -73,964 | *Atelopus laetissimus* | SNSM | negative | NA | NA | No |

| | | | | | | | | |
|---|---|---|---|---|---|---|---|---|
| 1/7/2024 | 10,905 | -73,964 | *Atelopus laetissimus* | SNSM | negative | NA | NA | No |
| 1/7/2024 | 10,905 | -73,964 | *Atelopus laetissimus* | SNSM | negative | NA | NA | No |
| 1/7/2024 | 10,905 | -73,964 | *Atelopus laetissimus* | SNSM | negative | NA | NA | No |
| 1/7/2024 | 10,905 | -73,964 | *Atelopus laetissimus* | SNSM | negative | NA | NA | No |
| 1/7/2024 | 10,905 | -73,964 | *Atelopus laetissimus* | SNSM | negative | NA | NA | No |
| 1/7/2024 | 10,905 | -73,964 | *Atelopus laetissimus* | SNSM | negative | NA | NA | No |
| 1/7/2024 | 10,905 | -73,964 | *Atelopus laetissimus* | SNSM | negative | NA | NA | No |
| 1/7/2024 | 10,905 | -73,964 | *Atelopus laetissimus* | SNSM | negative | NA | NA | No |
| 1/7/2024 | 10,905 | -73,964 | *Atelopus laetissimus* | SNSM | negative | NA | NA | No |
| 1/7/2024 | 10,905 | -73,964 | *Atelopus laetissimus* | SNSM | negative | NA | NA | No |
| 1/7/2024 | 10,905 | -73,964 | *Atelopus laetissimus* | SNSM | negative | NA | NA | No |
| 1/7/2024 | 10,905 | -73,964 | *Atelopus laetissimus* | SNSM | negative | NA | NA | No |
| 1/7/2024 | 10,905 | -73,964 | *Atelopus laetissimus* | SNSM | negative | NA | NA | No |
| 1/7/2024 | 10,905 | -73,964 | *Atelopus laetissimus* | SNSM | negative | NA | NA | No |
| 1/7/2024 | 10,905 | -73,964 | *Atelopus laetissimus* | SNSM | negative | NA | NA | No |
| 1/7/2024 | 10,905 | -73,964 | *Atelopus laetissimus* | SNSM | negative | NA | NA | No |
| 1/7/2024 | 10,905 | -73,964 | *Atelopus laetissimus* | SNSM | negative | NA | NA | No |
| 1/7/2024 | 10,905 | -73,964 | *Atelopus laetissimus* | SNSM | negative | NA | NA | No |
| 1/7/2024 | 10,905 | -73,964 | *Atelopus laetissimus* | SNSM | negative | NA | NA | No |
| 1/7/2024 | 10,905 | -73,964 | *Atelopus laetissimus* | SNSM | negative | NA | NA | No |
| 1/7/2024 | 10,905 | -73,964 | *Atelopus laetissimus* | SNSM | negative | NA | NA | No |
| 1/7/2024 | 10,905 | -73,964 | *Atelopus laetissimus* | SNSM | negative | NA | NA | No |
| 25/6/2025 | 10,905 | -73,964 | *Atelopus laetissimus* | SNSM | negative | NA | NA | No |
| 25/6/2025 | 10,905 | -73,964 | *Atelopus laetissimus* | SNSM | negative | NA | NA | No |
| 25/6/2025 | 10,905 | -73,964 | *Atelopus laetissimus* | SNSM | negative | NA | NA | No |
| 25/6/2025 | 10,905 | -73,964 | *Atelopus laetissimus* | SNSM | negative | NA | NA | No |
| 25/6/2025 | 10,905 | -73,964 | *Atelopus laetissimus* | SNSM | negative | NA | NA | No |
| 25/6/2025 | 10,905 | -73,964 | *Atelopus laetissimus* | SNSM | negative | NA | NA | No |
| 25/6/2025 | 10,905 | -73,964 | *Atelopus laetissimus* | SNSM | negative | NA | NA | No |
| 25/6/2025 | 10,905 | -73,964 | *Atelopus laetissimus* | SNSM | negative | NA | NA | No |
| 25/6/2025 | 10,905 | -73,964 | *Atelopus laetissimus* | SNSM | negative | NA | NA | No |
| 25/6/2025 | 10,905 | -73,964 | *Atelopus laetissimus* | SNSM | negative | NA | NA | No |
| 25/6/2025 | 10,905 | -73,964 | *Atelopus laetissimus* | SNSM | negative | NA | NA | No |
| 1/7/2024 | 10,905 | -73,964 | *Atelopus laetissimus* | SNSM | negative | NA | NA | No |
| 25/6/2025 | 10,905 | -73,964 | *Atelopus laetissimus* | SNSM | negative | NA | NA | No |
| 25/6/2025 | 10,905 | -73,964 | *Atelopus laetissimus* | SNSM | negative | NA | NA | No |
| 25/6/2025 | 10,905 | -73,964 | *Atelopus laetissimus* | SNSM | negative | NA | NA | No |

| | | | | | | | | |
|---|---|---|---|---|---|---|---|---|
| 25/6/2025 | 10,905 | -73,964 | *Atelopus laetissimus* | SNSM | negative | NA | NA | No |
| 25/6/2025 | 10,905 | -73,964 | *Atelopus laetissimus* | SNSM | negative | NA | NA | No |
| 25/6/2025 | 10,905 | -73,964 | *Atelopus laetissimus* | SNSM | negative | NA | NA | No |
| 25/6/2025 | 10,905 | -73,964 | *Atelopus laetissimus* | SNSM | negative | NA | NA | No |
| 25/6/2025 | 10,905 | -73,964 | *Atelopus laetissimus* | SNSM | negative | NA | NA | No |
| 25/6/2025 | 10,905 | -73,964 | *Atelopus laetissimus* | SNSM | negative | NA | NA | No |
| 25/6/2025 | 10,905 | -73,964 | *Atelopus laetissimus* | SNSM | negative | NA | NA | No |
| 25/6/2025 | 10,905 | -73,964 | *Atelopus laetissimus* | SNSM | negative | NA | NA | No |
| 25/6/2025 | 10,905 | -73,964 | *Atelopus laetissimus* | SNSM | negative | NA | NA | No |
| 25/6/2025 | 10,905 | -73,964 | *Atelopus laetissimus* | SNSM | negative | NA | NA | No |
| 25/6/2025 | 10,905 | -73,964 | *Atelopus laetissimus* | SNSM | negative | NA | NA | No |
| 25/6/2025 | 10,905 | -73,964 | *Atelopus laetissimus* | SNSM | negative | NA | NA | No |
| 25/6/2025 | 10,905 | -73,964 | *Atelopus laetissimus* | SNSM | negative | NA | NA | No |
| 25/6/2025 | 10,905 | -73,964 | *Atelopus laetissimus* | SNSM | negative | NA | NA | No |
| 25/6/2025 | 10,905 | -73,964 | *Atelopus laetissimus* | SNSM | negative | NA | NA | No |
| 25/6/2025 | 10,905 | -73,964 | *Atelopus laetissimus* | SNSM | negative | NA | NA | No |
| 25/6/2025 | 10,905 | -73,964 | *Atelopus laetissimus* | SNSM | negative | NA | NA | No |
| 25/6/2025 | 10,905 | -73,964 | *Atelopus laetissimus* | SNSM | negative | NA | NA | No |
| 1/7/2025 | 10,905 | -73,964 | *Atelopus laetissimus* | SNSM | negative | NA | NA | No |
| 1/7/2025 | 10,905 | -73,964 | *Atelopus laetissimus* | SNSM | negative | NA | NA | No |
| 1/7/2025 | 10,905 | -73,964 | *Atelopus laetissimus* | SNSM | negative | NA | NA | No |
| 1/7/2025 | 10,905 | -73,964 | *Atelopus laetissimus* | SNSM | negative | NA | NA | No |
| 29/7/2023 | 10,905 | -73,964 | *Atelopus laetissimus* | SNSM | negative | NA | NA | No |
| 20/6/2023 | 10,905 | -73,964 | *Atelopus laetissimus* | SNSM | negative | NA | NA | No |
| 21/1/2024 | 10,9 | -73,972 | *Serranobatrachus carmelitae* | SNSM | negative | NA | NA | No |
| 21/1/2024 | 10,9 | -73,972 | *Atelopus laetissimus* | SNSM | negative | NA | NA | No |
| 21/1/2024 | 10,9 | -73,972 | *Atelopus laetissimus* | SNSM | negative | NA | NA | No |
| 21/1/2024 | 10,9 | -73,972 | *Atelopus laetissimus* | SNSM | negative | NA | NA | No |
| 21/1/2024 | 10,9 | -73,972 | *Atelopus laetissimus* | SNSM | negative | NA | NA | No |
| 21/1/2024 | 10,9 | -73,972 | *Atelopus laetissimus* | SNSM | negative | NA | NA | No |
| 21/1/2024 | 10,9 | -73,972 | *Atelopus laetissimus* | SNSM | negative | NA | NA | No |
| 21/1/2024 | 10,9 | -73,972 | *Atelopus laetissimus* | SNSM | negative | NA | NA | No |
| 21/1/2024 | 10,9 | -73,972 | *Atelopus laetissimus* | SNSM | negative | NA | NA | No |
| 21/1/2024 | 10,9 | -73,972 | *Atelopus laetissimus* | SNSM | negative | NA | NA | No |
| 21/1/2024 | 10,9 | -73,972 | *Atelopus laetissimus* | SNSM | negative | NA | NA | No |
| 21/1/2024 | 10,9 | -73,972 | *Atelopus laetissimus* | SNSM | negative | NA | NA | No |
| 21/1/2024 | 10,9 | -73,972 | *Atelopus laetissimus* | SNSM | negative | NA | NA | No |

| | | | | | | | | |
|---|---|---|---|---|---|---|---|---|
| 21/1/2024 | 10,9 | -73,972 | *Atelopus laetissimus* | SNSM | negative | NA | NA | No |
| 21/1/2024 | 10,9 | -73,972 | *Serranobatrachus carmelitae* | SNSM | negative | NA | NA | No |
| 21/1/2024 | 10,9 | -73,972 | *Serranobatrachus carmelitae* | SNSM | negative | NA | NA | No |
| 21/1/2024 | 10,9 | -73,972 | *Serranobatrachus carmelitae* | SNSM | negative | NA | NA | No |
| 21/1/2024 | 10,9 | -73,972 | *Serranobatrachus sanctaemartae* | SNSM | negative | NA | NA | No |
| 21/1/2024 | 10,9 | -73,972 | *Serranobatrachus sanctaemartae* | SNSM | negative | NA | NA | No |
| 21/1/2024 | 10,9 | -73,972 | *Serranobatrachus ruthveni* | SNSM | negative | NA | NA | No |
| 21/1/2024 | 10,9 | -73,972 | *Serranobatrachus ruthveni* | SNSM | negative | NA | NA | No |
| 21/1/2024 | 10,9 | -73,972 | *Serranobatrachus sp.* | SNSM | negative | NA | NA | No |
| 21/1/2024 | 10,9 | -73,972 | *Atelopus laetissimus* | SNSM | negative | NA | NA | No |
| 21/1/2024 | 10,9 | -73,972 | *Serranobatrachus sp.* | SNSM | negative | NA | NA | No |
| 21/1/2024 | 10,9 | -73,972 | *Serranobatrachus sp.* | SNSM | negative | NA | NA | No |
| 21/1/2024 | 10,9 | -73,972 | *Serranobatrachus sp.* | SNSM | negative | NA | NA | No |
| 21/1/2024 | 10,9 | -73,972 | *Serranobatrachus sp.* | SNSM | negative | NA | NA | No |
| 21/1/2024 | 10,9 | -73,972 | *Serranobatrachus sp.* | SNSM | negative | NA | NA | No |
| 21/1/2024 | 10,9 | -73,972 | *Serranobatrachus delicatus* | SNSM | negative | NA | NA | No |
| 21/1/2024 | 10,9 | -73,972 | *Serranobatrachus sp.* | SNSM | negative | NA | NA | No |
| 21/1/2024 | 10,9 | -73,972 | *Serranobatrachus sp.* | SNSM | negative | NA | NA | No |
| 21/1/2024 | 10,9 | -73,972 | *Serranobatrachus sp.* | SNSM | negative | NA | NA | No |
| 21/1/2024 | 10,9 | -73,972 | *Atelopus laetissimus* | SNSM | negative | NA | NA | No |
| 21/1/2024 | 10,9 | -73,972 | *Atelopus laetissimus* | SNSM | negative | NA | NA | No |
| 21/1/2024 | 10,9 | -73,972 | *Atelopus laetissimus* | SNSM | negative | NA | NA | No |
| 21/1/2024 | 10,9 | -73,972 | *Atelopus laetissimus* | SNSM | negative | NA | NA | No |
| 21/1/2024 | 10,9 | -73,972 | *Atelopus laetissimus* | SNSM | negative | NA | NA | No |
| 21/1/2024 | 10,9 | -73,972 | *Atelopus laetissimus* | SNSM | negative | NA | NA | No |
| 21/1/2024 | 10,9 | -73,972 | *Atelopus laetissimus* | SNSM | negative | NA | NA | No |
| 21/1/2024 | 10,9 | -73,972 | *Atelopus laetissimus* | SNSM | negative | NA | NA | No |
| 21/1/2024 | 10,9 | -73,972 | *Atelopus laetissimus* | SNSM | negative | NA | NA | No |
| 21/1/2024 | 10,9 | -73,972 | *Atelopus laetissimus* | SNSM | negative | NA | NA | No |
| 21/1/2024 | 10,9 | -73,972 | *Atelopus laetissimus* | SNSM | negative | NA | NA | No |
| 21/1/2024 | 10,9 | -73,972 | *Atelopus laetissimus* | SNSM | negative | NA | NA | No |
| 21/1/2024 | 10,9 | -73,972 | *Atelopus laetissimus* | SNSM | negative | NA | NA | No |
| 21/1/2024 | 10,9 | -73,972 | *Atelopus laetissimus* | SNSM | negative | NA | NA | No |
| 21/1/2024 | 10,9 | -73,972 | *Atelopus laetissimus* | SNSM | negative | NA | NA | No |
| 21/1/2024 | 10,9 | -73,972 | *Atelopus laetissimus* | SNSM | negative | NA | NA | No |
| 29/7/2023 | 10,9 | -73,972 | *Atelopus laetissimus* | SNSM | negative | NA | NA | No |
| 20/6/2023 | 10,9 | -73,972 | *Atelopus laetissimus* | SNSM | negative | NA | NA | No |

| | | | | | | | | |
|---|---|---|---|---|---|---|---|---|
| 29/7/2023 | 10,9 | -73,972 | *Atelopus laetissimus* | SNSM | negative | NA | NA | No |
| 20/6/2023 | 10,9 | -73,972 | *Atelopus laetissimus* | SNSM | negative | NA | NA | No |
| 29/7/2023 | 10,9 | -73,972 | *Atelopus laetissimus* | SNSM | negative | NA | NA | No |
| 20/6/2023 | 10,9 | -73,972 | *Atelopus laetissimus* | SNSM | negative | NA | NA | No |
| 1/6/2023 | 10,9 | -73,972 | *Atelopus laetissimus* | SNSM | negative | NA | NA | No |
| 1/6/2023 | 10,9 | -73,972 | *Atelopus laetissimus* | SNSM | negative | NA | NA | No |
| 1/6/2023 | 10,9 | -73,972 | *Atelopus laetissimus* | SNSM | negative | NA | NA | No |
| 1/6/2023 | 10,9 | -73,972 | *Atelopus laetissimus* | SNSM | negative | NA | NA | No |
| 1/6/2023 | 10,9 | -73,972 | *Atelopus laetissimus* | SNSM | negative | NA | NA | No |
| 1/6/2023 | 10,9 | -73,972 | *Atelopus laetissimus* | SNSM | negative | NA | NA | No |
| 1/6/2023 | 10,9 | -73,972 | *Atelopus laetissimus* | SNSM | negative | NA | NA | No |
| 1/6/2023 | 10,9 | -73,972 | *Atelopus laetissimus* | SNSM | negative | NA | NA | No |
| 1/6/2023 | 10,9 | -73,972 | *Atelopus laetissimus* | SNSM | negative | NA | NA | No |
| 1/6/2023 | 10,9 | -73,972 | *Atelopus laetissimus* | SNSM | negative | NA | NA | No |
| 1/6/2023 | 10,9 | -73,972 | *Atelopus laetissimus* | SNSM | negative | NA | NA | No |
| 1/6/2023 | 10,9 | -73,972 | *Atelopus laetissimus* | SNSM | negative | NA | NA | No |
| 1/6/2023 | 10,9 | -73,972 | *Atelopus laetissimus* | SNSM | negative | NA | NA | No |
| 1/6/2023 | 10,9 | -73,972 | *Atelopus laetissimus* | SNSM | negative | NA | NA | No |
| 1/6/2023 | 10,9 | -73,972 | *Atelopus laetissimus* | SNSM | negative | NA | NA | No |
| 1/6/2023 | 10,9 | -73,972 | *Atelopus laetissimus* | SNSM | negative | NA | NA | No |
| 1/6/2023 | 10,9 | -73,972 | *Atelopus laetissimus* | SNSM | negative | NA | NA | No |
| 1/6/2023 | 10,9 | -73,972 | *Atelopus laetissimus* | SNSM | negative | NA | NA | No |
| 1/6/2023 | 10,9 | -73,972 | *Atelopus laetissimus* | SNSM | negative | NA | NA | No |
| 1/6/2023 | 10,9 | -73,972 | *Atelopus laetissimus* | SNSM | negative | NA | NA | No |
| 1/6/2023 | 10,9 | -73,972 | *Atelopus laetissimus* | SNSM | negative | NA | NA | No |
| 1/6/2023 | 10,9 | -73,972 | *Atelopus laetissimus* | SNSM | negative | NA | NA | No |
| 1/6/2023 | 10,9 | -73,972 | *Atelopus laetissimus* | SNSM | negative | NA | NA | No |
| 1/6/2023 | 10,9 | -73,972 | *Atelopus laetissimus* | SNSM | negative | NA | NA | No |
| 1/6/2023 | 10,9 | -73,972 | *Atelopus laetissimus* | SNSM | negative | NA | NA | No |
| 1/6/2023 | 10,9 | -73,972 | *Atelopus laetissimus* | SNSM | negative | NA | NA | No |
| 1/6/2023 | 10,9 | -73,972 | *Atelopus laetissimus* | SNSM | negative | NA | NA | No |
| 1/6/2023 | 10,9 | -73,972 | *Atelopus laetissimus* | SNSM | negative | NA | NA | No |
| 1/6/2023 | 10,9 | -73,972 | *Atelopus laetissimus* | SNSM | negative | NA | NA | No |
| 1/6/2023 | 10,9 | -73,972 | *Atelopus laetissimus* | SNSM | negative | NA | NA | No |
| 1/6/2023 | 10,9 | -73,972 | *Atelopus laetissimus* | SNSM | negative | NA | NA | No |
| 1/6/2023 | 10,9 | -73,972 | *Atelopus laetissimus* | SNSM | negative | NA | NA | No |
| 1/6/2023 | 10,9 | -73,972 | *Atelopus laetissimus* | SNSM | negative | NA | NA | No |

| | | | | | | | | |
|---|---|---|---|---|---|---|---|---|
| 1/6/2023 | 10,9 | -73,972 | *Atelopus laetissimus* | SNSM | negative | NA | NA | No |
| 1/6/2023 | 10,9 | -73,972 | *Atelopus laetissimus* | SNSM | negative | NA | NA | No |
| 1/6/2023 | 10,9 | -73,972 | *Atelopus laetissimus* | SNSM | negative | NA | NA | No |
| 1/6/2023 | 10,9 | -73,972 | *Atelopus laetissimus* | SNSM | negative | NA | NA | No |
| 1/6/2023 | 10,9 | -73,972 | *Atelopus laetissimus* | SNSM | negative | NA | NA | No |
| 1/6/2023 | 10,9 | -73,972 | *Atelopus laetissimus* | SNSM | negative | NA | NA | No |
| 1/6/2023 | 10,9 | -73,972 | *Atelopus laetissimus* | SNSM | negative | NA | NA | No |
| 1/6/2023 | 10,9 | -73,972 | *Atelopus laetissimus* | SNSM | negative | NA | NA | No |
| 1/6/2023 | 10,9 | -73,972 | *Atelopus laetissimus* | SNSM | negative | NA | NA | No |
| 1/6/2023 | 10,9 | -73,972 | *Atelopus laetissimus* | SNSM | negative | NA | NA | No |
| 1/6/2023 | 10,9 | -73,972 | *Atelopus laetissimus* | SNSM | negative | NA | NA | No |
| 1/6/2023 | 10,9 | -73,972 | *Atelopus laetissimus* | SNSM | negative | NA | NA | No |
| 1/6/2023 | 10,9 | -73,972 | *Atelopus laetissimus* | SNSM | negative | NA | NA | No |
| 1/6/2023 | 10,9 | -73,972 | *Atelopus laetissimus* | SNSM | negative | NA | NA | No |
| 1/6/2023 | 10,9 | -73,972 | *Atelopus laetissimus* | SNSM | negative | NA | NA | No |
| 1/6/2023 | 10,9 | -73,972 | *Atelopus laetissimus* | SNSM | negative | NA | NA | No |
| 1/6/2023 | 10,9 | -73,972 | *Atelopus laetissimus* | SNSM | negative | NA | NA | No |
| 1/6/2023 | 10,9 | -73,972 | *Atelopus laetissimus* | SNSM | negative | NA | NA | No |
| 1/6/2023 | 10,9 | -73,972 | *Atelopus laetissimus* | SNSM | negative | NA | NA | No |
| 1/6/2023 | 10,9 | -73,972 | *Atelopus laetissimus* | SNSM | negative | NA | NA | No |
| 1/6/2023 | 10,9 | -73,972 | *Atelopus laetissimus* | SNSM | negative | NA | NA | No |
| 1/6/2023 | 10,9 | -73,972 | *Atelopus laetissimus* | SNSM | negative | NA | NA | No |
| 1/6/2023 | 10,9 | -73,972 | *Atelopus laetissimus* | SNSM | negative | NA | NA | No |
| 29/7/2023 | 10,9 | -73,972 | *Atelopus laetissimus* | SNSM | negative | NA | NA | No |
| 20/6/2023 | 10,9 | -73,972 | *Atelopus laetissimus* | SNSM | negative | NA | NA | No |
| 29/7/2023 | 10,9 | -73,972 | *Atelopus laetissimus* | SNSM | negative | NA | NA | No |
| 20/6/2023 | 10,9 | -73,972 | *Atelopus laetissimus* | SNSM | negative | NA | NA | No |
| 29/7/2023 | 10,9 | -73,972 | *Atelopus laetissimus* | SNSM | negative | NA | NA | No |
| 20/6/2023 | 10,9 | -73,972 | *Atelopus laetissimus* | SNSM | negative | NA | NA | No |
| 9/6/2025 | 11,111 | -74,061 | *Atelopus laetissimus* | SNSM | negative | NA | NA | No |
| 9/6/2025 | 11,111 | -74,061 | *Atelopus laetissimus* | SNSM | negative | NA | NA | No |
| 9/6/2025 | 11,111 | -74,061 | *Atelopus laetissimus* | SNSM | negative | NA | NA | No |
| 9/6/2025 | 11,111 | -74,061 | *Atelopus laetissimus* | SNSM | negative | NA | NA | No |
| 9/6/2025 | 11,111 | -74,061 | *Atelopus laetissimus* | SNSM | negative | NA | NA | No |
| 9/6/2025 | 11,111 | -74,061 | *Atelopus laetissimus* | SNSM | negative | NA | NA | No |
| 9/6/2025 | 11,111 | -74,061 | *Atelopus laetissimus* | SNSM | negative | NA | NA | No |
| 9/6/2025 | 11,111 | -74,061 | *Atelopus laetissimus* | SNSM | negative | NA | NA | No |

| | | | | | | | | |
|---|---|---|---|---|---|---|---|---|
| 9/6/2025 | 11,111 | -74,061 | *Atelopus laetissimus* | SNSM | negative | NA | NA | No |
| 9/6/2025 | 11,111 | -74,061 | *Atelopus laetissimus* | SNSM | negative | NA | NA | No |
| 9/6/2025 | 11,111 | -74,061 | *Atelopus laetissimus* | SNSM | negative | NA | NA | No |
| 9/6/2025 | 11,111 | -74,061 | *Atelopus laetissimus* | SNSM | negative | NA | NA | No |
| 9/6/2025 | 11,111 | -74,061 | *Atelopus laetissimus* | SNSM | negative | NA | NA | No |
| 9/6/2025 | 11,111 | -74,061 | *Bolitoglossa savagei* | SNSM | negative | NA | NA | No |
| 9/6/2025 | 11,111 | -74,061 | *Bolitoglossa savagei* | SNSM | negative | NA | NA | No |
| 9/6/2025 | 11,111 | -74,061 | *Bolitoglossa savagei* | SNSM | negative | NA | NA | No |
| 9/6/2025 | 11,111 | -74,061 | *Bolitoglossa savagei* | SNSM | negative | NA | NA | No |
| 9/6/2025 | 11,111 | -74,061 | *Bolitoglossa savagei* | SNSM | negative | NA | NA | No |
| 9/6/2025 | 11,111 | -74,061 | *Bolitoglossa savagei* | SNSM | negative | NA | NA | No |
| 9/6/2025 | 11,111 | -74,061 | *Bolitoglossa savagei* | SNSM | negative | NA | NA | No |
| 9/6/2025 | 11,111 | -74,061 | *Bolitoglossa savagei* | SNSM | negative | NA | NA | No |
| 9/6/2025 | 11,111 | -74,061 | *Bolitoglossa savagei* | SNSM | negative | NA | NA | No |
| 9/6/2025 | 11,111 | -74,061 | *Bolitoglossa savagei* | SNSM | negative | NA | NA | No |
| 9/6/2025 | 11,111 | -74,061 | *Bolitoglossa savagei* | SNSM | negative | NA | NA | No |
| 9/6/2025 | 11,111 | -74,061 | *Bolitoglossa savagei* | SNSM | negative | NA | NA | No |
| 9/6/2025 | 11,111 | -74,061 | *Bolitoglossa savagei* | SNSM | negative | NA | NA | No |
| 9/6/2025 | 11,111 | -74,061 | *Bolitoglossa savagei* | SNSM | negative | NA | NA | No |
| 10/6/2025 | 11,102 | -74,081 | *Cryptobatrachus boulengeri* | SNSM | negative | NA | NA | No |
| 10/6/2025 | 11,102 | -74,081 | *Cryptobatrachus boulengeri* | SNSM | negative | NA | NA | No |
| 10/6/2025 | 11,102 | -74,081 | *Cryptobatrachus boulengeri* | SNSM | negative | NA | NA | No |
| 10/6/2025 | 11,102 | -74,081 | *Cryptobatrachus boulengeri* | SNSM | negative | NA | NA | No |
| 10/6/2025 | 11,102 | -74,081 | *Cryptobatrachus boulengeri* | SNSM | negative | NA | NA | No |
| 10/6/2025 | 11,102 | -74,081 | *Cryptobatrachus boulengeri* | SNSM | negative | NA | NA | No |
| 10/6/2025 | 11,102 | -74,081 | *Cryptobatrachus boulengeri* | SNSM | negative | NA | NA | No |
| 10/6/2025 | 11,102 | -74,081 | *Cryptobatrachus boulengeri* | SNSM | negative | NA | NA | No |
| 10/6/2025 | 11,102 | -74,081 | *Cryptobatrachus boulengeri* | SNSM | negative | NA | NA | No |
| 10/6/2025 | 11,102 | -74,081 | *Cryptobatrachus boulengeri* | SNSM | negative | NA | NA | No |
| 10/6/2025 | 11,102 | -74,081 | *Cryptobatrachus boulengeri* | SNSM | negative | NA | NA | No |
| 10/6/2025 | 11,102 | -74,081 | *Cryptobatrachus boulengeri* | SNSM | negative | NA | NA | No |
| 10/6/2025 | 11,102 | -74,081 | *Cryptobatrachus boulengeri* | SNSM | negative | NA | NA | No |
| 10/6/2025 | 11,102 | -74,081 | *Cryptobatrachus boulengeri* | SNSM | negative | NA | NA | No |
| 10/6/2025 | 11,102 | -74,081 | *Cryptobatrachus boulengeri* | SNSM | negative | NA | NA | No |
| 10/6/2025 | 11,102 | -74,081 | *Cryptobatrachus boulengeri* | SNSM | negative | NA | NA | No |
| 26/6/2024 | 10,732 | -73,478 | *Atelopus arsyecue* | SNSM | NA | negative | NA | No |
| 26/6/2024 | 10,732 | -73,478 | *Atelopus arsyecue* | SNSM | NA | negative | NA | No |
| 26/6/2024 | 10,732 | -73,478 | *Atelopus arsyecue* | SNSM | NA | negative | NA | No |

| | | | | | | | | |
|---|---|---|---|---|---|---|---|---|
| 26/6/2024 | 10,732 | -73,478 | *Atelopus arsyecue* | SNSM | NA | negative | NA | No |
| 26/6/2024 | 10,732 | -73,478 | *Atelopus arsyecue* | SNSM | NA | negative | NA | No |
| 26/6/2024 | 10,732 | -73,478 | *Atelopus arsyecue* | SNSM | NA | negative | NA | No |
| 26/6/2024 | 10,732 | -73,478 | *Atelopus arsyecue* | SNSM | NA | negative | NA | No |
| 26/6/2024 | 10,732 | -73,478 | *Atelopus arsyecue* | SNSM | NA | negative | NA | No |
| 26/6/2024 | 10,732 | -73,478 | *Atelopus arsyecue* | SNSM | NA | negative | NA | No |
| 26/6/2024 | 10,732 | -73,478 | *Atelopus arsyecue* | SNSM | NA | negative | NA | No |
| 26/6/2024 | 10,732 | -73,478 | *Atelopus arsyecue* | SNSM | NA | negative | NA | No |
| 26/6/2024 | 10,732 | -73,478 | *Atelopus arsyecue* | SNSM | NA | negative | NA | No |
| 26/6/2024 | 10,732 | -73,478 | *Atelopus arsyecue* | SNSM | NA | negative | NA | No |
| 26/6/2024 | 10,732 | -73,478 | *Atelopus arsyecue* | SNSM | NA | negative | NA | No |
| 26/6/2024 | 10,732 | -73,478 | *Atelopus arsyecue* | SNSM | NA | negative | NA | No |
| 26/6/2024 | 10,732 | -73,478 | *Atelopus arsyecue* | SNSM | NA | negative | NA | No |
| 26/6/2024 | 10,732 | -73,478 | *Atelopus arsyecue* | SNSM | NA | negative | NA | No |
| 26/6/2024 | 10,732 | -73,478 | *Atelopus arsyecue* | SNSM | NA | negative | NA | No |
| 26/6/2024 | 10,732 | -73,478 | *Atelopus arsyecue* | SNSM | NA | negative | NA | No |
| 26/6/2024 | 10,732 | -73,478 | *Atelopus arsyecue* | SNSM | NA | negative | NA | No |
| 26/6/2024 | 10,732 | -73,478 | *Atelopus arsyecue* | SNSM | NA | negative | NA | No |
| 26/6/2024 | 10,732 | -73,478 | *Atelopus arsyecue* | SNSM | NA | negative | NA | No |
| 26/6/2024 | 10,732 | -73,478 | *Atelopus arsyecue* | SNSM | NA | negative | NA | No |
| 26/6/2024 | 10,732 | -73,478 | *Atelopus arsyecue* | SNSM | NA | negative | NA | No |
| 26/6/2024 | 10,732 | -73,478 | *Atelopus arsyecue* | SNSM | NA | negative | NA | No |
| 26/6/2024 | 10,732 | -73,478 | *Atelopus arsyecue* | SNSM | NA | negative | NA | No |
| 26/6/2024 | 10,732 | -73,478 | *Atelopus arsyecue* | SNSM | NA | negative | NA | No |
| 26/6/2024 | 10,732 | -73,478 | *Atelopus arsyecue* | SNSM | NA | negative | NA | No |
| 26/6/2024 | 10,732 | -73,478 | *Atelopus arsyecue* | SNSM | NA | negative | NA | No |
| 26/6/2024 | 10,732 | -73,478 | *Atelopus arsyecue* | SNSM | NA | negative | NA | No |
| 26/6/2024 | 10,732 | -73,478 | *Atelopus arsyecue* | SNSM | NA | negative | NA | No |
| 26/6/2024 | 10,732 | -73,478 | *Atelopus arsyecue* | SNSM | NA | negative | NA | No |
| 26/6/2024 | 10,732 | -73,478 | *Atelopus arsyecue* | SNSM | NA | negative | NA | No |
| 26/6/2024 | 10,732 | -73,478 | *Atelopus arsyecue* | SNSM | NA | negative | NA | No |
| 26/6/2024 | 10,732 | -73,478 | *Atelopus arsyecue* | SNSM | NA | negative | NA | No |
| 26/6/2024 | 10,732 | -73,478 | *Atelopus arsyecue* | SNSM | NA | negative | NA | No |
| 26/6/2024 | 10,732 | -73,478 | *Atelopus arsyecue* | SNSM | NA | negative | NA | No |
| 26/6/2024 | 10,732 | -73,478 | *Atelopus arsyecue* | SNSM | NA | negative | NA | No |
| 26/6/2024 | 10,732 | -73,478 | *Atelopus arsyecue* | SNSM | NA | negative | NA | No |
| 26/6/2024 | 10,732 | -73,478 | *Atelopus arsyecue* | SNSM | NA | negative | NA | No |
| 26/6/2024 | 10,732 | -73,478 | *Atelopus arsyecue* | SNSM | NA | negative | NA | No |
| 26/6/2024 | 10,732 | -73,478 | *Atelopus arsyecue* | SNSM | NA | negative | NA | No |
| 26/6/2024 | 10,732 | -73,478 | *Atelopus arsyecue* | SNSM | NA | negative | NA | No |
| 26/6/2024 | 10,732 | -73,478 | *Atelopus arsyecue* | SNSM | NA | negative | NA | No |
| 26/6/2024 | 10,732 | -73,478 | *Atelopus arsyecue* | SNSM | NA | negative | NA | No |
| 26/6/2024 | 10,732 | -73,478 | *Atelopus arsyecue* | SNSM | NA | negative | NA | No |
| 26/6/2024 | 10,732 | -73,478 | *Atelopus arsyecue* | SNSM | NA | negative | NA | No |

| | | | | | | | | |
|---|---|---|---|---|---|---|---|---|
| 26/6/2024 | 10,732 | -73,478 | *Atelopus arsyecue* | SNSM | NA | negative | NA | No |
| 26/6/2024 | 10,732 | -73,478 | *Atelopus arsyecue* | SNSM | NA | negative | NA | No |
| 26/6/2024 | 10,732 | -73,478 | *Atelopus arsyecue* | SNSM | NA | negative | NA | No |
| 26/6/2024 | 10,732 | -73,478 | *Atelopus arsyecue* | SNSM | NA | negative | NA | No |
| 26/6/2024 | 10,732 | -73,478 | *Atelopus arsyecue* | SNSM | NA | negative | NA | No |
| 26/6/2024 | 10,732 | -73,478 | *Atelopus arsyecue* | SNSM | NA | negative | NA | No |
| 26/6/2024 | 10,732 | -73,478 | *Atelopus arsyecue* | SNSM | NA | negative | NA | No |
| 26/6/2024 | 10,732 | -73,478 | *Atelopus arsyecue* | SNSM | NA | negative | NA | No |
| 26/6/2024 | 10,732 | -73,478 | *Atelopus arsyecue* | SNSM | NA | negative | NA | No |
| 26/6/2024 | 10,732 | -73,478 | *Atelopus arsyecue* | SNSM | NA | negative | NA | No |
| 26/6/2024 | 10,732 | -73,478 | *Atelopus arsyecue* | SNSM | NA | negative | NA | No |
| 26/6/2024 | 10,732 | -73,478 | *Atelopus arsyecue* | SNSM | NA | negative | NA | No |
| 26/6/2024 | 10,732 | -73,478 | *Atelopus arsyecue* | SNSM | NA | negative | NA | No |
| 26/6/2024 | 10,732 | -73,478 | *Atelopus arsyecue* | SNSM | NA | negative | NA | No |
| 26/6/2024 | 10,732 | -73,478 | *Atelopus arsyecue* | SNSM | NA | negative | NA | No |
| 22/6/2024 | 11,124 | -74,054 | environmental DNA | SNSM | negative | NA | NA | Yes |
| 19/6/2024 | 11,111 | -74,061 | environmental DNA | SNSM | negative | NA | NA | Yes |
| 23/6/2024 | 11,094 | -74,076 | environmental DNA | SNSM | negative | NA | NA | Yes |
| 25/6/2024 | 11,111 | -74,061 | environmental DNA | SNSM | negative | NA | NA | Yes |
| 3/7/2024 | 11,094 | -74,076 | environmental DNA | SNSM | negative | NA | NA | Yes |
| 4/7/2024 | 11,111 | -74,061 | environmental DNA | SNSM | negative | NA | NA | Yes |
| 6/7/2024 | 11,094 | -74,076 | environmental DNA | SNSM | negative | NA | NA | Yes |
| 21/7/2024 | 11,111 | -74,061 | environmental DNA | SNSM | negative | NA | NA | Yes |
| 22/7/2024 | 11,094 | -74,076 | environmental DNA | SNSM | negative | NA | NA | Yes |
| 24/7/2024 | 11,111 | -74,061 | environmental DNA | SNSM | negative | NA | NA | Yes |
| 26/7/2024 | 11,094 | -74,076 | environmental DNA | SNSM | negative | NA | NA | Yes |
| 13/8/2024 | 11,094 | -74,076 | environmental DNA | SNSM | negative | NA | NA | Yes |
| 20/8/2024 | 11,111 | -74,061 | environmental DNA | SNSM | negative | NA | NA | Yes |
| 19/6/2024 | 11,111 | -74,061 | *Atelopus nahumae* | SNSM | negative | negative | NA | No |
| 19/6/2024 | 11,111 | -74,061 | *Atelopus nahumae* | SNSM | NA | negative | NA | No |
| 19/6/2024 | 11,111 | -74,061 | *Atelopus laetissimus* | SNSM | NA | negative | NA | No |
| 19/6/2024 | 11,111 | -74,061 | *Atelopus laetissimus* | SNSM | NA | negative | NA | No |
| 19/6/2024 | 11,111 | -74,061 | *Atelopus laetissimus* | SNSM | NA | negative | NA | No |
| 19/6/2024 | 11,111 | -74,061 | *Atelopus laetissimus* | SNSM | NA | negative | NA | No |
| 19/6/2024 | 11,111 | -74,061 | *Atelopus laetissimus* | SNSM | NA | negative | NA | No |
| 19/6/2024 | 11,111 | -74,061 | *Atelopus laetissimus* | SNSM | NA | negative | NA | No |
| 19/6/2024 | 11,111 | -74,061 | *Atelopus laetissimus* | SNSM | NA | negative | NA | No |
| 19/6/2024 | 11,111 | -74,061 | *Atelopus laetissimus* | SNSM | NA | negative | NA | No |
| 19/6/2024 | 11,111 | -74,061 | *Atelopus laetissimus* | SNSM | NA | negative | NA | No |
| 19/6/2024 | 11,111 | -74,061 | *Atelopus laetissimus* | SNSM | NA | negative | NA | No |

| | | | | | | | | |
|---|---|---|---|---|---|---|---|---|
| 19/6/2024 | 11,111 | -74,061 | *Atelopus laetissimus* | SNSM | NA | negative | NA | No |
| 19/6/2024 | 11,111 | -74,061 | *Ikakogi tayrona* | SNSM | NA | negative | NA | No |
| 19/6/2024 | 11,111 | -74,061 | *Serranobatrachus sanctaemartae* | SNSM | NA | negative | NA | No |
| 19/6/2024 | 11,111 | -74,061 | *Serranobatrachus sanctaemartae* | SNSM | NA | negative | NA | No |
| 19/6/2024 | 11,111 | -74,061 | *Serranobatrachus sanctaemartae* | SNSM | NA | negative | NA | No |
| 19/6/2024 | 11,111 | -74,061 | *Serranobatrachus sanctaemartae* | SNSM | NA | negative | NA | No |
| 19/6/2024 | 11,111 | -74,061 | *Serranobatrachus sanctaemartae* | SNSM | NA | negative | NA | No |
| 19/6/2024 | 11,111 | -74,061 | *Serranobatrachus sanctaemartae* | SNSM | NA | negative | NA | No |
| 19/6/2024 | 11,111 | -74,061 | *Serranobatrachus sanctaemartae* | SNSM | NA | negative | NA | No |
| 19/6/2024 | 11,111 | -74,061 | *Serranobatrachus sanctaemartae* | SNSM | NA | negative | NA | No |
| 19/6/2024 | 11,111 | -74,061 | *Serranobatrachus cristinae* | SNSM | NA | negative | NA | No |
| 19/6/2024 | 11,111 | -74,061 | *Serranobatrachus sanctaemartae* | SNSM | NA | negative | NA | No |
| 19/6/2024 | 11,111 | -74,061 | *Serranobatrachus sanctaemartae* | SNSM | NA | negative | NA | No |
| 19/6/2024 | 11,111 | -74,061 | *Serranobatrachus sanctaemartae* | SNSM | NA | negative | NA | No |
| 19/6/2024 | 11,111 | -74,061 | *Serranobatrachus sanctaemartae* | SNSM | NA | negative | NA | No |
| 19/6/2024 | 11,111 | -74,061 | *Serranobatrachus megalops* | SNSM | NA | negative | NA | No |
| 19/6/2024 | 11,111 | -74,061 | *Serranobatrachus sanctaemartae* | SNSM | NA | negative | NA | No |
| 19/6/2024 | 11,111 | -74,061 | *Serranobatrachus sanctaemartae* | SNSM | NA | negative | NA | No |
| 19/6/2024 | 11,111 | -74,061 | *Serranobatrachus megalops* | SNSM | NA | negative | NA | No |
| 19/6/2024 | 11,111 | -74,061 | *Serranobatrachus megalops* | SNSM | NA | negative | NA | No |
| 19/6/2024 | 11,111 | -74,061 | *Serranobatrachus sanctaemartae* | SNSM | NA | negative | NA | No |
| 19/6/2024 | 11,111 | -74,061 | *Geobatrachus walkeri* | SNSM | NA | negative | NA | No |
| 19/6/2024 | 11,111 | -74,061 | *Serranobatrachus megalops* | SNSM | NA | negative | NA | No |
| 19/6/2024 | 11,111 | -74,061 | *Serranobatrachus sanctaemartae* | SNSM | NA | negative | NA | No |
| 19/6/2024 | 11,111 | -74,061 | *Serranobatrachus megalops* | SNSM | NA | negative | NA | No |
| 19/6/2024 | 11,111 | -74,061 | *Serranobatrachus sanctaemartae* | SNSM | NA | negative | NA | No |
| 19/6/2024 | 11,111 | -74,061 | *Serranobatrachus sanctaemartae* | SNSM | NA | negative | NA | No |
| 19/6/2024 | 11,111 | -74,061 | *Serranobatrachus sanctaemartae* | SNSM | NA | negative | NA | No |
| 19/6/2024 | 11,111 | -74,061 | *Serranobatrachus megalops* | SNSM | NA | negative | NA | No |
| 19/6/2024 | 11,111 | -74,061 | *Serranobatrachus megalops* | SNSM | NA | negative | NA | No |
| 19/6/2024 | 11,111 | -74,061 | *Serranobatrachus megalops* | SNSM | NA | negative | NA | No |
| 19/6/2024 | 11,111 | -74,061 | *Serranobatrachus sanctaemartae* | SNSM | NA | negative | NA | No |
| 19/6/2024 | 11,111 | -74,061 | *Serranobatrachus megalops* | SNSM | NA | negative | NA | No |
| 19/6/2024 | 11,111 | -74,061 | *Serranobatrachus insignitus* | SNSM | NA | negative | NA | No |
| 19/6/2024 | 11,111 | -74,061 | *Serranobatrachus megalops* | SNSM | NA | negative | NA | No |
| 19/6/2024 | 11,111 | -74,061 | *Serranobatrachus sanctaemartae* | SNSM | NA | negative | NA | No |
| 19/6/2024 | 11,111 | -74,061 | *Serranobatrachus megalops* | SNSM | NA | negative | NA | No |
| 19/6/2024 | 11,111 | -74,061 | *Geobatrachus walkeri* | SNSM | NA | negative | NA | No |

| | | | | | | | | |
|---|---|---|---|---|---|---|---|---|
| 21/6/2024 | 11,102 | -74,062 | *Serranobatrachus sanctaemartae* | SNSM | NA | negative | NA | No |
| 21/6/2024 | 11,102 | -74,062 | *Serranobatrachus megalops* | SNSM | NA | negative | NA | No |
| 21/6/2024 | 11,102 | -74,062 | *Atelopus nahumae* | SNSM | NA | negative | NA | No |
| 21/6/2024 | 11,102 | -74,062 | *Serranobatrachus megalops* | SNSM | NA | negative | NA | No |
| 21/6/2024 | 11,102 | -74,062 | *Bolitoglossa savagei* | SNSM | NA | negative | NA | No |
| 21/6/2024 | 11,102 | -74,062 | *Serranobatrachus megalops* | SNSM | NA | negative | NA | No |
| 21/6/2024 | 11,102 | -74,062 | *Serranobatrachus carmelitae* | SNSM | NA | negative | NA | No |
| 21/6/2024 | 11,102 | -74,062 | *Geobatrachus walkeri* | SNSM | NA | negative | NA | No |
| 21/6/2024 | 11,102 | -74,062 | *Serranobatrachus insignitus* | SNSM | NA | negative | NA | No |
| 21/6/2024 | 11,102 | -74,062 | *Tachiramantis tayrona* | SNSM | NA | negative | NA | No |
| 21/6/2024 | 11,102 | -74,062 | *Serranobatrachus megalops* | SNSM | NA | negative | NA | No |
| 21/6/2024 | 11,102 | -74,062 | *Serranobatrachus megalops* | SNSM | NA | negative | NA | No |
| 21/6/2024 | 11,102 | -74,062 | *Serranobatrachus carmelitae* | SNSM | NA | negative | NA | No |
| 21/6/2024 | 11,102 | -74,062 | *Serranobatrachus sanctaemartae* | SNSM | NA | negative | NA | No |
| 21/6/2024 | 11,102 | -74,062 | *Bolitoglossa savagei* | SNSM | NA | negative | NA | No |
| 21/6/2024 | 11,102 | -74,062 | *Bolitoglossa savagei* | SNSM | NA | negative | NA | No |
| 21/6/2024 | 11,102 | -74,062 | *Bolitoglossa savagei* | SNSM | NA | negative | NA | No |
| 21/6/2024 | 11,102 | -74,062 | *Tachiramantis tayrona* | SNSM | NA | negative | NA | No |
| 21/6/2024 | 11,102 | -74,062 | *Tachiramantis tayrona* | SNSM | NA | negative | NA | No |
| 21/6/2024 | 11,102 | -74,062 | *Tachiramantis tayrona* | SNSM | NA | negative | NA | No |
| 21/6/2024 | 11,102 | -74,062 | *Tachiramantis tayrona* | SNSM | NA | negative | NA | No |
| 21/6/2024 | 11,102 | -74,062 | *Tachiramantis tayrona* | SNSM | NA | negative | NA | No |
| 21/6/2024 | 11,102 | -74,062 | *Tachiramantis tayrona* | SNSM | NA | negative | NA | No |
| 21/6/2024 | 11,102 | -74,062 | *Serranobatrachus megalops* | SNSM | NA | negative | NA | No |
| 21/6/2024 | 11,102 | -74,062 | *Tachiramantis tayrona* | SNSM | NA | negative | NA | No |
| 21/6/2024 | 11,102 | -74,062 | *Serranobatrachus megalops* | SNSM | NA | negative | NA | No |
| 21/6/2024 | 11,102 | -74,062 | *Serranobatrachus megalops* | SNSM | NA | negative | NA | No |
| 21/6/2024 | 11,102 | -74,062 | *Serranobatrachus sanctaemartae* | SNSM | NA | negative | NA | No |
| 21/6/2024 | 11,102 | -74,062 | *Tachiramantis tayrona* | SNSM | NA | negative | NA | No |
| 21/6/2024 | 11,102 | -74,062 | *Serranobatrachus sanctaemartae* | SNSM | NA | negative | NA | No |
| 21/6/2024 | 11,102 | -74,062 | *Serranobatrachus megalops* | SNSM | NA | negative | NA | No |
| 21/6/2024 | 11,102 | -74,062 | *Tachiramantis tayrona* | SNSM | NA | negative | NA | No |
| 21/6/2024 | 11,102 | -74,062 | *Serranobatrachus cristinae* | SNSM | NA | negative | NA | No |
| 21/6/2024 | 11,102 | -74,062 | *Serranobatrachus delicatus* | SNSM | NA | negative | NA | No |
| 21/6/2024 | 11,102 | -74,062 | *Tachiramantis tayrona* | SNSM | NA | negative | NA | No |
| 21/6/2024 | 11,102 | -74,062 | *Serranobatrachus megalops* | SNSM | NA | negative | NA | No |
| 21/6/2024 | 11,102 | -74,062 | *Serranobatrachus sanctaemartae* | SNSM | NA | negative | NA | No |
| 21/6/2024 | 11,102 | -74,062 | *Serranobatrachus sanctaemartae* | SNSM | NA | negative | NA | No |

| | | | | | | | | |
|---|---|---|---|---|---|---|---|---|
| 21/6/2024 | 11,102 | -74,062 | *Serranobatrachus sanctaemartae* | SNSM | NA | negative | NA | No |
| 21/6/2024 | 11,102 | -74,062 | *Serranobatrachus sanctaemartae* | SNSM | NA | negative | NA | No |
| 21/6/2024 | 11,102 | -74,062 | *Serranobatrachus sanctaemartae* | SNSM | NA | negative | NA | No |
| 21/6/2024 | 11,102 | -74,062 | *Tachiramantis tayrona* | SNSM | NA | negative | NA | No |
| 21/6/2024 | 11,102 | -74,062 | *Serranobatrachus megalops* | SNSM | NA | negative | NA | No |
| 21/6/2024 | 11,102 | -74,062 | *Geobatrachus walkeri* | SNSM | NA | negative | NA | No |
| 21/6/2024 | 11,102 | -74,062 | *Tachiramantis tayrona* | SNSM | NA | negative | NA | No |
| 21/6/2024 | 11,102 | -74,062 | *Geobatrachus walkeri* | SNSM | NA | negative | NA | No |
| 21/6/2024 | 11,102 | -74,062 | *Geobatrachus walkeri* | SNSM | NA | negative | NA | No |
| 21/6/2024 | 11,102 | -74,062 | *Serranobatrachus sanctaemartae* | SNSM | NA | negative | NA | No |
| 21/6/2024 | 11,102 | -74,062 | *Tachiramantis tayrona* | SNSM | NA | negative | NA | No |
| 21/6/2024 | 11,102 | -74,062 | *Serranobatrachus ruthveni* | SNSM | NA | negative | NA | No |
| 21/6/2024 | 11,102 | -74,062 | *Serranobatrachus ruthveni* | SNSM | NA | negative | NA | No |
| 22/6/2024 | 11,126 | -74,053 | *Serranobatrachus delicatus* | SNSM | NA | negative | NA | No |
| 22/6/2024 | 11,126 | -74,053 | *Serranobatrachus megalops* | SNSM | NA | negative | NA | No |
| 22/6/2024 | 11,126 | -74,053 | *Cryptobatrachus boulengeri* | SNSM | NA | negative | NA | No |
| 22/6/2024 | 11,126 | -74,053 | *Cryptobatrachus boulengeri* | SNSM | NA | negative | NA | No |
| 22/6/2024 | 11,126 | -74,053 | *Serranobatrachus delicatus* | SNSM | NA | negative | NA | No |
| 22/6/2024 | 11,126 | -74,053 | *Serranobatrachus megalops* | SNSM | NA | negative | NA | No |
| 22/6/2024 | 11,126 | -74,053 | *Serranobatrachus delicatus* | SNSM | NA | negative | NA | No |
| 22/6/2024 | 11,126 | -74,053 | *Serranobatrachus delicatus* | SNSM | NA | negative | NA | No |
| 22/6/2024 | 11,126 | -74,053 | *Serranobatrachus delicatus* | SNSM | NA | negative | NA | No |
| 22/6/2024 | 11,126 | -74,053 | *Atelopus nahumae* | SNSM | NA | negative | NA | No |
| 22/6/2024 | 11,126 | -74,053 | *Atelopus nahumae* | SNSM | NA | negative | NA | No |
| 23/6/2024 | 11,094 | -74,076 | *Cryptobatrachus boulengeri* | SNSM | negative | negative | NA | No |
| 23/6/2024 | 11,094 | -74,076 | *Cryptobatrachus boulengeri* | SNSM | NA | negative | NA | No |
| 23/6/2024 | 11,094 | -74,076 | *Serranobatrachus sanctaemartae* | SNSM | NA | negative | NA | No |
| 23/6/2024 | 11,094 | -74,076 | *Serranobatrachus megalops* | SNSM | NA | negative | NA | No |
| 23/6/2024 | 11,094 | -74,076 | *Cryptobatrachus boulengeri* | SNSM | NA | negative | NA | No |
| 23/6/2024 | 11,094 | -74,076 | *Serranobatrachus megalops* | SNSM | NA | negative | NA | No |
| 23/6/2024 | 11,094 | -74,076 | *Serranobatrachus sanctaemartae* | SNSM | NA | negative | NA | No |
| 23/6/2024 | 11,094 | -74,076 | *Serranobatrachus sanctaemartae* | SNSM | NA | negative | NA | No |
| 23/6/2024 | 11,094 | -74,076 | *Serranobatrachus megalops* | SNSM | NA | negative | NA | No |
| 23/6/2024 | 11,094 | -74,076 | *Serranobatrachus sanctaemartae* | SNSM | NA | negative | NA | No |
| 23/6/2024 | 11,094 | -74,076 | *Serranobatrachus megalops* | SNSM | NA | negative | NA | No |
| 23/6/2024 | 11,094 | -74,076 | *Cryptobatrachus boulengeri* | SNSM | NA | negative | NA | No |
| 23/6/2024 | 11,094 | -74,076 | *Serranobatrachus sanctaemartae* | SNSM | NA | negative | NA | No |
| 23/6/2024 | 11,094 | -74,076 | *Cryptobatrachus boulengeri* | SNSM | NA | negative | NA | No |

| | | | | | | | | |
|---|---|---|---|---|---|---|---|---|
| 23/6/2024 | 11,094 | -74,076 | *Cryptobatrachus boulengeri* | SNSM | NA | negative | NA | No |
| 23/6/2024 | 11,094 | -74,076 | *Serranobatrachus megalops* | SNSM | NA | negative | NA | No |
| 23/6/2024 | 11,094 | -74,076 | *Cryptobatrachus boulengeri* | SNSM | NA | negative | NA | No |
| 23/6/2024 | 11,094 | -74,076 | *Serranobatrachus insignitus* | SNSM | NA | negative | NA | No |
| 23/6/2024 | 11,094 | -74,076 | *Cryptobatrachus boulengeri* | SNSM | NA | negative | NA | No |
| 23/6/2024 | 11,094 | -74,076 | *Cryptobatrachus boulengeri* | SNSM | negative | negative | NA | No |
| 23/6/2024 | 11,094 | -74,076 | *Cryptobatrachus boulengeri* | SNSM | NA | negative | NA | No |
| 23/6/2024 | 11,094 | -74,076 | *Serranobatrachus sanctaemartae* | SNSM | NA | negative | NA | No |
| 23/6/2024 | 11,094 | -74,076 | *Cryptobatrachus boulengeri* | SNSM | NA | negative | NA | No |
| 23/6/2024 | 11,094 | -74,076 | *Cryptobatrachus boulengeri* | SNSM | NA | negative | NA | No |
| 23/6/2024 | 11,094 | -74,076 | *Cryptobatrachus boulengeri* | SNSM | NA | negative | NA | No |
| 23/6/2024 | 11,094 | -74,076 | *Cryptobatrachus boulengeri* | SNSM | NA | negative | NA | No |
| 23/6/2024 | 11,094 | -74,076 | *Cryptobatrachus boulengeri* | SNSM | NA | negative | NA | No |
| 23/6/2024 | 11,094 | -74,076 | *Cryptobatrachus boulengeri* | SNSM | NA | negative | NA | No |
| 23/6/2024 | 11,094 | -74,076 | *Cryptobatrachus boulengeri* | SNSM | NA | negative | NA | No |
| 23/6/2024 | 11,094 | -74,076 | *Cryptobatrachus boulengeri* | SNSM | NA | negative | NA | No |
| 23/6/2024 | 11,094 | -74,076 | *Ikakogi tayrona* | SNSM | NA | negative | NA | No |
| 23/6/2024 | 11,094 | -74,076 | *Ikakogi tayrona* | SNSM | NA | negative | NA | No |
| 23/6/2024 | 11,094 | -74,076 | *Ikakogi tayrona* | SNSM | NA | negative | NA | No |
| 25/6/2024 | 11,111 | -74,061 | *Serranobatrachus carmelitae* | SNSM | NA | negative | NA | No |
| 25/6/2024 | 11,111 | -74,061 | *Ikakogi tayrona* | SNSM | NA | negative | NA | No |
| 25/6/2024 | 11,111 | -74,061 | *Serranobatrachus sanctaemartae* | SNSM | NA | negative | NA | No |
| 25/6/2024 | 11,111 | -74,061 | *Serranobatrachus carmelitae* | SNSM | NA | negative | NA | No |
| 25/6/2024 | 11,111 | -74,061 | *Serranobatrachus megalops* | SNSM | NA | negative | NA | No |
| 25/6/2024 | 11,111 | -74,061 | *Serranobatrachus megalops* | SNSM | NA | negative | NA | No |
| 25/6/2024 | 11,111 | -74,061 | *Serranobatrachus megalops* | SNSM | NA | negative | NA | No |
| 25/6/2024 | 11,111 | -74,061 | *Serranobatrachus sanctaemartae* | SNSM | NA | negative | NA | No |
| 25/6/2024 | 11,111 | -74,061 | *Serranobatrachus sanctaemartae* | SNSM | NA | negative | NA | No |
| 25/6/2024 | 11,111 | -74,061 | *Serranobatrachus sanctaemartae* | SNSM | NA | negative | NA | No |
| 25/6/2024 | 11,111 | -74,061 | *Atelopus nahumae* | SNSM | NA | negative | NA | No |
| 25/6/2024 | 11,111 | -74,061 | *Atelopus nahumae* | SNSM | NA | negative | NA | No |
| 25/6/2024 | 11,111 | -74,061 | *Serranobatrachus insignitus* | SNSM | NA | negative | NA | No |
| 25/6/2024 | 11,111 | -74,061 | *Serranobatrachus cristinae* | SNSM | NA | negative | NA | No |
| 25/6/2024 | 11,111 | -74,061 | *Serranobatrachus sanctaemartae* | SNSM | NA | negative | NA | No |
| 25/6/2024 | 11,111 | -74,061 | *Serranobatrachus sanctaemartae* | SNSM | NA | negative | NA | No |
| 25/6/2024 | 11,111 | -74,061 | *Serranobatrachus megalops* | SNSM | NA | negative | NA | No |
| 25/6/2024 | 11,111 | -74,061 | *Serranobatrachus cristinae* | SNSM | NA | negative | NA | No |
| 25/6/2024 | 11,111 | -74,061 | *Serranobatrachus sanctaemartae* | SNSM | NA | negative | NA | No |

| | | | | | | | | |
|---|---|---|---|---|---|---|---|---|
| 25/6/2024 | 11,111 | -74,061 | *Serranobatrachus sanctaemartae* | SNSM | NA | negative | NA | No |
| 25/6/2024 | 11,111 | -74,061 | *Serranobatrachus sanctaemartae* | SNSM | NA | negative | NA | No |
| 25/6/2024 | 11,111 | -74,061 | *Ikakogi tayrona* | SNSM | NA | negative | NA | No |
| 25/6/2024 | 11,111 | -74,061 | *Serranobatrachus sanctaemartae* | SNSM | NA | negative | NA | No |
| 25/6/2024 | 11,111 | -74,061 | *Atelopus nahumae* | SNSM | NA | negative | NA | No |
| 25/6/2024 | 11,111 | -74,061 | *Atelopus nahumae* | SNSM | NA | negative | NA | No |
| 25/6/2024 | 11,111 | -74,061 | *Serranobatrachus carmelitae* | SNSM | NA | negative | NA | No |
| 25/6/2024 | 11,111 | -74,061 | *Serranobatrachus sanctaemartae* | SNSM | NA | negative | NA | No |
| 25/6/2024 | 11,111 | -74,061 | *Serranobatrachus megalops* | SNSM | NA | negative | NA | No |
| 25/6/2024 | 11,111 | -74,061 | *Serranobatrachus megalops* | SNSM | NA | negative | NA | No |
| 25/6/2024 | 11,111 | -74,061 | *Serranobatrachus sanctaemartae* | SNSM | NA | negative | NA | No |
| 25/6/2024 | 11,111 | -74,061 | *Serranobatrachus carmelitae* | SNSM | NA | negative | NA | No |
| 25/6/2024 | 11,111 | -74,061 | *Ikakogi tayrona* | SNSM | NA | negative | NA | No |
| 25/6/2024 | 11,111 | -74,061 | *Serranobatrachus sanctaemartae* | SNSM | NA | negative | NA | No |
| 25/6/2024 | 11,111 | -74,061 | *Ikakogi tayrona* | SNSM | NA | negative | NA | No |
| 25/6/2024 | 11,111 | -74,061 | *Serranobatrachus sanctaemartae* | SNSM | NA | negative | NA | No |
| 25/6/2024 | 11,111 | -74,061 | *Serranobatrachus sanctaemartae* | SNSM | NA | negative | NA | No |
| 25/6/2024 | 11,111 | -74,061 | *Atelopus nahumae* | SNSM | NA | negative | NA | No |
| 25/6/2024 | 11,111 | -74,061 | *Atelopus nahumae* | SNSM | NA | negative | NA | No |
| 25/6/2024 | 11,111 | -74,061 | *Atelopus nahumae* | SNSM | NA | negative | NA | No |
| 25/6/2024 | 11,111 | -74,061 | *Serranobatrachus sanctaemartae* | SNSM | NA | negative | NA | No |
| 25/6/2024 | 11,111 | -74,061 | *Serranobatrachus sanctaemartae* | SNSM | NA | negative | NA | No |
| 25/6/2024 | 11,111 | -74,061 | *Serranobatrachus megalops* | SNSM | NA | negative | NA | No |
| 25/6/2024 | 11,111 | -74,061 | *Serranobatrachus megalops* | SNSM | NA | negative | NA | No |
| 25/6/2024 | 11,111 | -74,061 | *Ikakogi tayrona* | SNSM | NA | negative | NA | No |
| 25/6/2024 | 11,111 | -74,061 | *Serranobatrachus sanctaemartae* | SNSM | NA | negative | NA | No |
| 25/6/2024 | 11,111 | -74,061 | *Atelopus nahumae* | SNSM | NA | negative | NA | No |
| 25/6/2024 | 11,111 | -74,061 | *Serranobatrachus carmelitae* | SNSM | NA | negative | NA | No |
| 27/6/2024 | 11,108 | -74,071 | *Bolitoglossa savagei* | SNSM | NA | negative | NA | No |
| 27/6/2024 | 11,108 | -74,071 | *Bolitoglossa savagei* | SNSM | NA | negative | NA | No |
| 27/6/2024 | 11,108 | -74,071 | *Tachiramantis tayrona* | SNSM | NA | negative | NA | No |
| 27/6/2024 | 11,108 | -74,071 | *Ikakogi tayrona* | SNSM | NA | negative | NA | No |
| 27/6/2024 | 11,108 | -74,071 | *Serranobatrachus insignitus* | SNSM | NA | negative | NA | No |
| 27/6/2024 | 11,108 | -74,071 | *Serranobatrachus sanctaemartae* | SNSM | NA | negative | NA | No |
| 27/6/2024 | 11,108 | -74,071 | *Tachiramantis tayrona* | SNSM | NA | negative | NA | No |
| 27/6/2024 | 11,108 | -74,071 | *Serranobatrachus megalops* | SNSM | NA | negative | NA | No |
| 27/6/2024 | 11,108 | -74,071 | *Serranobatrachus delicatus* | SNSM | NA | negative | NA | No |
| 27/6/2024 | 11,108 | -74,071 | *Serranobatrachus carmelitae* | SNSM | NA | negative | NA | No |
| 27/6/2024 | 11,108 | -74,071 | *Serranobatrachus insignitus* | SNSM | NA | negative | NA | No |

| | | | | | | | | |
|---|---|---|---|---|---|---|---|---|
| 27/6/2024 | 11,108 | -74,071 | *Serranobatrachus insignitus* | SNSM | NA | negative | NA | No |
| 27/6/2024 | 11,108 | -74,071 | *Serranobatrachus sanctaemartae* | SNSM | NA | negative | NA | No |
| 27/6/2024 | 11,108 | -74,071 | *Serranobatrachus sanctaemartae* | SNSM | NA | negative | NA | No |
| 27/6/2024 | 11,108 | -74,071 | *Serranobatrachus sanctaemartae* | SNSM | NA | negative | NA | No |
| 27/6/2024 | 11,108 | -74,071 | *Atelopus nahumae* | SNSM | NA | negative | NA | No |
| 27/6/2024 | 11,108 | -74,071 | *Tachiramantis tayrona* | SNSM | NA | negative | NA | No |
| 27/6/2024 | 11,108 | -74,071 | *Serranobatrachus sanctaemartae* | SNSM | NA | negative | NA | No |
| 27/6/2024 | 11,108 | -74,071 | *Serranobatrachus sanctaemartae* | SNSM | NA | negative | NA | No |
| 27/6/2024 | 11,108 | -74,071 | *Tachiramantis tayrona* | SNSM | NA | negative | NA | No |
| 27/6/2024 | 11,108 | -74,071 | *Serranobatrachus sanctaemartae* | SNSM | NA | negative | NA | No |
| 27/6/2024 | 11,108 | -74,071 | *Serranobatrachus carmelitae* | SNSM | NA | negative | NA | No |
| 27/6/2024 | 11,108 | -74,071 | *Cryptobatrachus boulengeri* | SNSM | NA | negative | NA | No |
| 27/6/2024 | 11,108 | -74,071 | *Serranobatrachus sanctaemartae* | SNSM | NA | negative | NA | No |
| 27/6/2024 | 11,108 | -74,071 | *Serranobatrachus insignitus* | SNSM | NA | negative | NA | No |
| 27/6/2024 | 11,108 | -74,071 | *Serranobatrachus insignitus* | SNSM | NA | negative | NA | No |
| 27/6/2024 | 11,108 | -74,071 | *Cryptobatrachus boulengeri* | SNSM | NA | negative | NA | No |
| 27/6/2024 | 11,108 | -74,071 | *Cryptobatrachus boulengeri* | SNSM | NA | negative | NA | No |
| 27/6/2024 | 11,108 | -74,071 | *Cryptobatrachus boulengeri* | SNSM | NA | negative | NA | No |
| 27/6/2024 | 11,108 | -74,071 | *Serranobatrachus sanctaemartae* | SNSM | NA | negative | NA | No |
| 27/6/2024 | 11,108 | -74,071 | *Cryptobatrachus boulengeri* | SNSM | NA | negative | NA | No |
| 27/6/2024 | 11,108 | -74,071 | *Serranobatrachus insignitus* | SNSM | NA | negative | NA | No |
| 27/6/2024 | 11,108 | -74,071 | *Cryptobatrachus boulengeri* | SNSM | negative | negative | NA | No |
| 27/6/2024 | 11,108 | -74,071 | *Serranobatrachus sanctaemartae* | SNSM | NA | negative | NA | No |
| 27/6/2024 | 11,108 | -74,071 | *Cryptobatrachus boulengeri* | SNSM | negative | negative | NA | No |
| 27/6/2024 | 11,108 | -74,071 | *Serranobatrachus sanctaemartae* | SNSM | NA | negative | NA | No |
| 27/6/2024 | 11,108 | -74,071 | *Cryptobatrachus boulengeri* | SNSM | NA | negative | NA | No |
| 27/6/2024 | 11,108 | -74,071 | *Cryptobatrachus boulengeri* | SNSM | NA | negative | NA | No |
| 27/6/2024 | 11,108 | -74,071 | *Serranobatrachus megalops* | SNSM | NA | negative | NA | No |
| 27/6/2024 | 11,108 | -74,071 | *Cryptobatrachus boulengeri* | SNSM | NA | negative | NA | No |
| 27/6/2024 | 11,108 | -74,071 | *Bolitoglossa savagei* | SNSM | NA | negative | NA | No |
| 27/6/2024 | 11,108 | -74,071 | *Ikakogi tayrona* | SNSM | NA | negative | NA | No |
| 27/6/2024 | 11,108 | -74,071 | *Ikakogi tayrona* | SNSM | NA | negative | NA | No |
| 27/6/2024 | 11,108 | -74,071 | *Serranobatrachus megalops* | SNSM | NA | negative | NA | No |
| 27/6/2024 | 11,108 | -74,071 | *Serranobatrachus sanctaemartae* | SNSM | NA | negative | NA | No |
| 27/6/2024 | 11,108 | -74,071 | *Serranobatrachus sanctaemartae* | SNSM | NA | negative | NA | No |
| 27/6/2024 | 11,108 | -74,071 | *Cryptobatrachus boulengeri* | SNSM | NA | negative | NA | No |
| 27/6/2024 | 11,108 | -74,071 | *Serranobatrachus megalops* | SNSM | NA | negative | NA | No |
| 27/6/2024 | 11,108 | -74,071 | *Serranobatrachus sanctaemartae* | SNSM | NA | negative | NA | No |

| | | | | | | | | |
|---|---|---|---|---|---|---|---|---|
| 27/6/2024 | 11,108 | -74,071 | *Cryptobatrachus boulengeri* | SNSM | NA | negative | NA | No |
| 27/6/2024 | 11,108 | -74,071 | *Cryptobatrachus boulengeri* | SNSM | NA | negative | NA | No |
| 27/6/2024 | 11,108 | -74,071 | *Serranobatrachus megalops* | SNSM | NA | negative | NA | No |
| 27/6/2024 | 11,108 | -74,071 | *Serranobatrachus megalops* | SNSM | NA | negative | NA | No |
| 27/6/2024 | 11,108 | -74,071 | *Serranobatrachus delicatus* | SNSM | NA | negative | NA | No |
| 27/6/2024 | 11,108 | -74,071 | *Serranobatrachus megalops* | SNSM | NA | negative | NA | No |
| 27/6/2024 | 11,108 | -74,071 | *Cryptobatrachus boulengeri* | SNSM | NA | negative | NA | No |
| 27/6/2024 | 11,108 | -74,071 | *Serranobatrachus megalops* | SNSM | NA | negative | NA | No |
| 27/6/2024 | 11,108 | -74,071 | *Cryptobatrachus boulengeri* | SNSM | NA | negative | NA | No |
| 27/6/2024 | 11,108 | -74,071 | *Serranobatrachus sanctaemartae* | SNSM | NA | negative | NA | No |
| 27/6/2024 | 11,108 | -74,071 | *Cryptobatrachus boulengeri* | SNSM | NA | negative | NA | No |
| 27/6/2024 | 11,108 | -74,071 | *Serranobatrachus insignitus* | SNSM | NA | negative | NA | No |
| 27/6/2024 | 11,108 | -74,071 | *Serranobatrachus sanctaemartae* | SNSM | NA | negative | NA | No |
| 27/6/2024 | 11,108 | -74,071 | *Cryptobatrachus boulengeri* | SNSM | NA | negative | NA | No |
| 27/6/2024 | 11,108 | -74,071 | *Serranobatrachus insignitus* | SNSM | NA | negative | NA | No |
| 27/6/2024 | 11,108 | -74,071 | *Cryptobatrachus boulengeri* | SNSM | negative | negative | NA | No |
| 27/6/2024 | 11,108 | -74,071 | *Serranobatrachus delicatus* | SNSM | NA | negative | NA | No |
| 27/6/2024 | 11,108 | -74,071 | *Serranobatrachus sanctaemartae* | SNSM | NA | negative | NA | No |
| 27/6/2024 | 11,108 | -74,071 | *Serranobatrachus sanctaemartae* | SNSM | NA | negative | NA | No |
| 27/6/2024 | 11,108 | -74,071 | *Serranobatrachus megalops* | SNSM | NA | negative | NA | No |
| 27/6/2024 | 11,108 | -74,071 | *Cryptobatrachus boulengeri* | SNSM | NA | negative | NA | No |
| 28/6/2024 | 11,115 | -74,05 | *Atelopus nahumae* | SNSM | NA | negative | NA | No |
| 28/6/2024 | 11,115 | -74,05 | *Atelopus nahumae* | SNSM | NA | negative | NA | No |
| 28/6/2024 | 11,115 | -74,05 | *Atelopus laetissimus* | SNSM | NA | negative | NA | No |
| 28/6/2024 | 11,115 | -74,05 | *Atelopus laetissimus* | SNSM | NA | negative | NA | No |
| 28/6/2024 | 11,115 | -74,05 | *Atelopus laetissimus* | SNSM | NA | negative | NA | No |
| 28/6/2024 | 11,115 | -74,05 | *Atelopus laetissimus* | SNSM | NA | negative | NA | No |
| 28/6/2024 | 11,115 | -74,05 | *Atelopus laetissimus* | SNSM | NA | negative | NA | No |
| 28/6/2024 | 11,115 | -74,05 | *Atelopus laetissimus* | SNSM | NA | negative | NA | No |
| 28/6/2024 | 11,115 | -74,05 | *Atelopus laetissimus* | SNSM | NA | negative | NA | No |
| 28/6/2024 | 11,115 | -74,05 | *Atelopus laetissimus* | SNSM | NA | negative | NA | No |
| 28/6/2024 | 11,115 | -74,05 | *Atelopus laetissimus* | SNSM | NA | negative | NA | No |
| 28/6/2024 | 11,115 | -74,05 | *Atelopus laetissimus* | SNSM | NA | negative | NA | No |
| 28/6/2024 | 11,115 | -74,05 | *Atelopus laetissimus* | SNSM | NA | negative | NA | No |
| 28/6/2024 | 11,115 | -74,05 | *Atelopus laetissimus* | SNSM | NA | negative | NA | No |
| 28/6/2024 | 11,115 | -74,05 | *Atelopus nahumae* | SNSM | NA | negative | NA | No |
| 28/6/2024 | 11,115 | -74,05 | *Atelopus laetissimus* | SNSM | NA | negative | NA | No |
| 28/6/2024 | 11,115 | -74,05 | *Atelopus laetissimus* | SNSM | NA | negative | NA | No |

| | | | | | | | | |
|---|---|---|---|---|---|---|---|---|
| 28/6/2024 | 11,115 | -74,05 | *Atelopus laetissimus* | SNSM | NA | negative | NA | No |
| 28/6/2024 | 11,115 | -74,05 | *Atelopus laetissimus* | SNSM | NA | negative | NA | No |
| 28/6/2024 | 11,115 | -74,05 | *Atelopus laetissimus* | SNSM | NA | negative | NA | No |
| 28/6/2024 | 11,115 | -74,05 | *Atelopus laetissimus* | SNSM | NA | negative | NA | No |
| 28/6/2024 | 11,115 | -74,05 | *Atelopus laetissimus* | SNSM | NA | negative | NA | No |
| 28/6/2024 | 11,115 | -74,05 | *Atelopus laetissimus* | SNSM | negative | negative | NA | No |
| 28/6/2024 | 11,115 | -74,05 | *Atelopus laetissimus* | SNSM | NA | negative | NA | No |
| 28/6/2024 | 11,115 | -74,05 | *Atelopus laetissimus* | SNSM | NA | negative | NA | No |
| 28/6/2024 | 11,115 | -74,05 | *Atelopus laetissimus* | SNSM | NA | negative | NA | No |
| 28/6/2024 | 11,115 | -74,05 | *Atelopus laetissimus* | SNSM | NA | negative | NA | No |
| 28/6/2024 | 11,115 | -74,05 | *Atelopus laetissimus* | SNSM | NA | negative | NA | No |
| 28/6/2024 | 11,115 | -74,05 | *Atelopus laetissimus* | SNSM | NA | negative | NA | No |
| 28/6/2024 | 11,115 | -74,05 | *Atelopus laetissimus* | SNSM | NA | negative | NA | No |
| 28/6/2024 | 11,115 | -74,05 | *Ikakogi tayrona* | SNSM | NA | negative | NA | No |
| 28/6/2024 | 11,115 | -74,05 | *Atelopus nahumae* | SNSM | NA | negative | NA | No |
| 28/6/2024 | 11,115 | -74,05 | *Atelopus nahumae* | SNSM | NA | negative | NA | No |
| 3/7/2024 | 11,094 | -74,076 | *Cryptobatrachus boulengeri* | SNSM | NA | negative | NA | No |
| 3/7/2024 | 11,094 | -74,076 | *Cryptobatrachus boulengeri* | SNSM | NA | negative | NA | No |
| 3/7/2024 | 11,094 | -74,076 | *Cryptobatrachus boulengeri* | SNSM | NA | negative | NA | No |
| 3/7/2024 | 11,094 | -74,076 | *Cryptobatrachus boulengeri* | SNSM | NA | negative | NA | No |
| 3/7/2024 | 11,094 | -74,076 | *Cryptobatrachus boulengeri* | SNSM | NA | negative | NA | No |
| 3/7/2024 | 11,094 | -74,076 | *Cryptobatrachus boulengeri* | SNSM | NA | negative | NA | No |
| 3/7/2024 | 11,094 | -74,076 | *Cryptobatrachus boulengeri* | SNSM | NA | negative | NA | No |
| 3/7/2024 | 11,094 | -74,076 | *Cryptobatrachus boulengeri* | SNSM | NA | negative | NA | No |
| 3/7/2024 | 11,094 | -74,076 | *Cryptobatrachus boulengeri* | SNSM | NA | negative | NA | No |
| 3/7/2024 | 11,094 | -74,076 | *Cryptobatrachus boulengeri* | SNSM | NA | negative | NA | No |
| 3/7/2024 | 11,094 | -74,076 | *Cryptobatrachus boulengeri* | SNSM | NA | negative | NA | No |
| 3/7/2024 | 11,094 | -74,076 | *Cryptobatrachus boulengeri* | SNSM | negative | negative | NA | No |
| 3/7/2024 | 11,094 | -74,076 | *Cryptobatrachus boulengeri* | SNSM | NA | negative | NA | No |
| 3/7/2024 | 11,094 | -74,076 | *Serranobatrachus sanctaemartae* | SNSM | NA | negative | NA | No |
| 3/7/2024 | 11,094 | -74,076 | *Ikakogi tayrona* | SNSM | NA | negative | NA | No |
| 3/7/2024 | 11,094 | -74,076 | *Ikakogi tayrona* | SNSM | NA | negative | NA | No |
| 3/7/2024 | 11,094 | -74,076 | *Ikakogi tayrona* | SNSM | NA | negative | NA | No |
| 4/7/2024 | 11,111 | -74,061 | *Serranobatrachus megalops* | SNSM | NA | negative | NA | No |
| 4/7/2024 | 11,111 | -74,061 | *Serranobatrachus megalops* | SNSM | negative | negative | NA | No |
| 4/7/2024 | 11,111 | -74,061 | *Ikakogi tayrona* | SNSM | NA | negative | NA | No |
| 4/7/2024 | 11,111 | -74,061 | *Serranobatrachus sanctaemartae* | SNSM | NA | negative | NA | No |
| 4/7/2024 | 11,111 | -74,061 | *Serranobatrachus sanctaemartae* | SNSM | NA | negative | NA | No |

| | | | | | | | | |
|---|---|---|---|---|---|---|---|---|
| 4/7/2024 | 11,111 | -74,061 | *Serranobatrachus megalops* | SNSM | NA | negative | NA | No |
| 4/7/2024 | 11,111 | -74,061 | *Atelopus laetissimus* | SNSM | NA | negative | NA | No |
| 4/7/2024 | 11,111 | -74,061 | *Atelopus laetissimus* | SNSM | NA | negative | NA | No |
| 4/7/2024 | 11,111 | -74,061 | *Atelopus laetissimus* | SNSM | NA | negative | NA | No |
| 4/7/2024 | 11,111 | -74,061 | *Serranobatrachus sanctaemartae* | SNSM | NA | negative | NA | No |
| 4/7/2024 | 11,111 | -74,061 | *Serranobatrachus sanctaemartae* | SNSM | NA | negative | NA | No |
| 4/7/2024 | 11,111 | -74,061 | *Serranobatrachus sanctaemartae* | SNSM | NA | negative | NA | No |
| 4/7/2024 | 11,111 | -74,061 | *Serranobatrachus megalops* | SNSM | NA | negative | NA | No |
| 4/7/2024 | 11,111 | -74,061 | *Serranobatrachus sanctaemartae* | SNSM | NA | negative | NA | No |
| 4/7/2024 | 11,111 | -74,061 | *Serranobatrachus sanctaemartae* | SNSM | NA | negative | NA | No |
| 4/7/2024 | 11,111 | -74,061 | *Serranobatrachus carmelitae* | SNSM | NA | negative | NA | No |
| 4/7/2024 | 11,111 | -74,061 | *Atelopus laetissimus* | SNSM | NA | negative | NA | No |
| 4/7/2024 | 11,111 | -74,061 | *Serranobatrachus megalops* | SNSM | NA | negative | NA | No |
| 4/7/2024 | 11,111 | -74,061 | *Ikakogi tayrona* | SNSM | NA | negative | NA | No |
| 4/7/2024 | 11,111 | -74,061 | *Atelopus laetissimus* | SNSM | NA | negative | NA | No |
| 4/7/2024 | 11,111 | -74,061 | *Serranobatrachus sanctaemartae* | SNSM | NA | negative | NA | No |
| 4/7/2024 | 11,111 | -74,061 | *Serranobatrachus sanctaemartae* | SNSM | NA | negative | NA | No |
| 4/7/2024 | 11,111 | -74,061 | *Ikakogi tayrona* | SNSM | NA | negative | NA | No |
| 4/7/2024 | 11,111 | -74,061 | *Ikakogi tayrona* | SNSM | NA | negative | NA | No |
| 4/7/2024 | 11,111 | -74,061 | *Serranobatrachus sanctaemartae* | SNSM | NA | negative | NA | No |
| 4/7/2024 | 11,111 | -74,061 | *Atelopus laetissimus* | SNSM | NA | negative | NA | No |
| 4/7/2024 | 11,111 | -74,061 | *Atelopus laetissimus* | SNSM | NA | negative | NA | No |
| 4/7/2024 | 11,111 | -74,061 | *Serranobatrachus sanctaemartae* | SNSM | NA | negative | NA | No |
| 4/7/2024 | 11,111 | -74,061 | *Serranobatrachus sanctaemartae* | SNSM | NA | negative | NA | No |
| 4/7/2024 | 11,111 | -74,061 | *Serranobatrachus megalops* | SNSM | NA | negative | NA | No |
| 4/7/2024 | 11,111 | -74,061 | *Serranobatrachus megalops* | SNSM | NA | negative | NA | No |
| 4/7/2024 | 11,111 | -74,061 | *Serranobatrachus sanctaemartae* | SNSM | negative | negative | NA | No |
| 4/7/2024 | 11,111 | -74,061 | *Atelopus laetissimus* | SNSM | negative | negative | NA | No |
| 4/7/2024 | 11,111 | -74,061 | *Serranobatrachus sanctaemartae* | SNSM | negative | negative | NA | No |
| 4/7/2024 | 11,111 | -74,061 | *Serranobatrachus cristinae* | SNSM | NA | negative | NA | No |
| 6/7/2024 | 11,094 | -74,076 | *Ikakogi tayrona* | SNSM | NA | negative | NA | No |
| 6/7/2024 | 11,094 | -74,076 | *Serranobatrachus megalops* | SNSM | NA | negative | NA | No |
| 6/7/2024 | 11,094 | -74,076 | *Serranobatrachus megalops* | SNSM | NA | negative | NA | No |
| 6/7/2024 | 11,094 | -74,076 | *Serranobatrachus sanctaemartae* | SNSM | NA | negative | NA | No |
| 6/7/2024 | 11,094 | -74,076 | *Serranobatrachus sanctaemartae* | SNSM | NA | negative | NA | No |
| 6/7/2024 | 11,094 | -74,076 | *Serranobatrachus sanctaemartae* | SNSM | NA | negative | NA | No |
| 6/7/2024 | 11,094 | -74,076 | *Cryptobatrachus boulengeri* | SNSM | NA | negative | NA | No |
| 6/7/2024 | 11,094 | -74,076 | *Cryptobatrachus boulengeri* | SNSM | NA | negative | NA | No |

| | | | | | | | | |
|---|---|---|---|---|---|---|---|---|
| 6/7/2024 | 11,094 | -74,076 | *Cryptobatrachus boulengeri* | SNSM | NA | negative | NA | No |
| 6/7/2024 | 11,094 | -74,076 | *Cryptobatrachus boulengeri* | SNSM | NA | negative | NA | No |
| 6/7/2024 | 11,094 | -74,076 | *Cryptobatrachus boulengeri* | SNSM | NA | negative | NA | No |
| 6/7/2024 | 11,094 | -74,076 | *Cryptobatrachus boulengeri* | SNSM | NA | negative | NA | No |
| 6/7/2024 | 11,094 | -74,076 | *Serranobatrachus sanctaemartae* | SNSM | NA | negative | NA | No |
| 6/7/2024 | 11,094 | -74,076 | *Serranobatrachus megalops* | SNSM | NA | negative | NA | No |
| 6/7/2024 | 11,094 | -74,076 | *Cryptobatrachus boulengeri* | SNSM | NA | negative | NA | No |
| 6/7/2024 | 11,094 | -74,076 | *Cryptobatrachus boulengeri* | SNSM | NA | negative | NA | No |
| 6/7/2024 | 11,094 | -74,076 | *Serranobatrachus sanctaemartae* | SNSM | NA | negative | NA | No |
| 6/7/2024 | 11,094 | -74,076 | *Cryptobatrachus boulengeri* | SNSM | NA | negative | NA | No |
| 6/7/2024 | 11,094 | -74,076 | *Cryptobatrachus boulengeri* | SNSM | NA | negative | NA | No |
| 6/7/2024 | 11,094 | -74,076 | *Cryptobatrachus boulengeri* | SNSM | NA | negative | NA | No |
| 6/7/2024 | 11,094 | -74,076 | *Cryptobatrachus boulengeri* | SNSM | NA | negative | NA | No |
| 6/7/2024 | 11,094 | -74,076 | *Cryptobatrachus boulengeri* | SNSM | NA | negative | NA | No |
| 6/7/2024 | 11,094 | -74,076 | *Cryptobatrachus boulengeri* | SNSM | NA | negative | NA | No |
| 6/7/2024 | 11,094 | -74,076 | *Cryptobatrachus boulengeri* | SNSM | NA | negative | NA | No |
| 6/7/2024 | 11,094 | -74,076 | *Cryptobatrachus boulengeri* | SNSM | NA | negative | NA | No |
| 6/7/2024 | 11,094 | -74,076 | *Serranobatrachus insignitus* | SNSM | NA | negative | NA | No |
| 6/7/2024 | 11,094 | -74,076 | *Serranobatrachus megalops* | SNSM | NA | negative | NA | No |
| 6/7/2024 | 11,094 | -74,076 | *Serranobatrachus sanctaemartae* | SNSM | NA | negative | NA | No |
| 6/7/2024 | 11,094 | -74,076 | *Cryptobatrachus boulengeri* | SNSM | NA | negative | NA | No |
| 6/7/2024 | 11,094 | -74,076 | *Serranobatrachus megalops* | SNSM | NA | negative | NA | No |
| 6/7/2024 | 11,094 | -74,076 | *Cryptobatrachus boulengeri* | SNSM | NA | negative | NA | No |
| 6/7/2024 | 11,094 | -74,076 | *Cryptobatrachus boulengeri* | SNSM | NA | negative | NA | No |
| 6/7/2024 | 11,094 | -74,076 | *Serranobatrachus megalops* | SNSM | NA | negative | NA | No |
| 6/7/2024 | 11,094 | -74,076 | *Cryptobatrachus boulengeri* | SNSM | NA | negative | NA | No |
| 6/7/2024 | 11,094 | -74,076 | *Serranobatrachus sanctaemartae* | SNSM | NA | negative | NA | No |
| 6/7/2024 | 11,094 | -74,076 | *Serranobatrachus sanctaemartae* | SNSM | NA | negative | NA | No |
| 6/7/2024 | 11,094 | -74,076 | *Cryptobatrachus boulengeri* | SNSM | NA | negative | NA | No |
| 6/7/2024 | 11,094 | -74,076 | *Cryptobatrachus boulengeri* | SNSM | NA | negative | NA | No |
| 8/7/2024 | 11,111 | -74,061 | *Atelopus laetissimus* | SNSM | NA | negative | NA | No |
| 8/7/2024 | 11,111 | -74,061 | *Atelopus laetissimus* | SNSM | NA | negative | NA | No |
| 8/7/2024 | 11,111 | -74,061 | *Atelopus laetissimus* | SNSM | NA | negative | NA | No |
| 21/7/2024 | 11,111 | -74,061 | *Atelopus laetissimus* | SNSM | negative | NA | NA | No |
| 21/7/2024 | 11,111 | -74,061 | *Serranobatrachus cristinae* | SNSM | NA | negative | NA | No |
| 21/7/2024 | 11,111 | -74,061 | *Serranobatrachus sanctaemartae* | SNSM | negative | NA | NA | No |
| 21/7/2024 | 11,111 | -74,061 | *Atelopus laetissimus* | SNSM | NA | negative | NA | No |

| | | | | | | | | |
|---|---|---|---|---|---|---|---|---|
| 21/7/2024 | 11,111 | -74,061 | *Serranobatrachus sanctaemartae* | SNSM | NA | negative | NA | No |
| 21/7/2024 | 11,111 | -74,061 | *Serranobatrachus sanctaemartae* | SNSM | NA | negative | NA | No |
| 21/7/2024 | 11,111 | -74,061 | *Serranobatrachus megalops* | SNSM | negative | NA | NA | No |
| 21/7/2024 | 11,111 | -74,061 | *Atelopus laetissimus* | SNSM | NA | negative | NA | No |
| 21/7/2024 | 11,111 | -74,061 | *Serranobatrachus sanctaemartae* | SNSM | NA | negative | NA | No |
| 21/7/2024 | 11,111 | -74,061 | *Ikakogi tayrona* | SNSM | NA | negative | NA | No |
| 21/7/2024 | 11,111 | -74,061 | *Atelopus laetissimus* | SNSM | negative | NA | NA | No |
| 21/7/2024 | 11,111 | -74,061 | *Serranobatrachus sanctaemartae* | SNSM | NA | negative | NA | No |
| 21/7/2024 | 11,111 | -74,061 | *Atelopus laetissimus* | SNSM | NA | negative | NA | No |
| 21/7/2024 | 11,111 | -74,061 | *Serranobatrachus megalops* | SNSM | NA | negative | NA | No |
| 21/7/2024 | 11,111 | -74,061 | *Serranobatrachus sanctaemartae* | SNSM | NA | negative | NA | No |
| 21/7/2024 | 11,111 | -74,061 | *Ikakogi tayrona* | SNSM | NA | negative | NA | No |
| 21/7/2024 | 11,111 | -74,061 | *Serranobatrachus sanctaemartae* | SNSM | NA | negative | NA | No |
| 21/7/2024 | 11,111 | -74,061 | *Serranobatrachus cristinae* | SNSM | negative | NA | NA | No |
| 21/7/2024 | 11,111 | -74,061 | *Atelopus laetissimus* | SNSM | NA | negative | NA | No |
| 21/7/2024 | 11,111 | -74,061 | *Ikakogi tayrona* | SNSM | negative | NA | NA | No |
| 21/7/2024 | 11,111 | -74,061 | *Atelopus laetissimus* | SNSM | negative | NA | NA | No |
| 21/7/2024 | 11,111 | -74,061 | *Atelopus laetissimus* | SNSM | NA | negative | NA | No |
| 21/7/2024 | 11,111 | -74,061 | *Serranobatrachus sanctaemartae* | SNSM | negative | NA | NA | No |
| 22/7/2024 | 11,094 | -74,076 | *Cryptobatrachus boulengeri* | SNSM | NA | negative | NA | No |
| 22/7/2024 | 11,094 | -74,076 | *Cryptobatrachus boulengeri* | SNSM | NA | negative | NA | No |
| 22/7/2024 | 11,094 | -74,076 | *Cryptobatrachus boulengeri* | SNSM | negative | NA | NA | No |
| 22/7/2024 | 11,094 | -74,076 | *Cryptobatrachus boulengeri* | SNSM | NA | negative | NA | No |
| 22/7/2024 | 11,094 | -74,076 | *Cryptobatrachus boulengeri* | SNSM | negative | NA | NA | No |
| 22/7/2024 | 11,094 | -74,076 | *Cryptobatrachus boulengeri* | SNSM | NA | negative | NA | No |
| 22/7/2024 | 11,094 | -74,076 | *Serranobatrachus sanctaemartae* | SNSM | NA | negative | NA | No |
| 22/7/2024 | 11,094 | -74,076 | *Cryptobatrachus boulengeri* | SNSM | NA | negative | NA | No |
| 22/7/2024 | 11,094 | -74,076 | *Cryptobatrachus boulengeri* | SNSM | negative | NA | NA | No |
| 22/7/2024 | 11,094 | -74,076 | *Cryptobatrachus boulengeri* | SNSM | negative | NA | NA | No |
| 22/7/2024 | 11,094 | -74,076 | *Serranobatrachus megalops* | SNSM | NA | negative | NA | No |
| 22/7/2024 | 11,094 | -74,076 | *Cryptobatrachus boulengeri* | SNSM | negative | NA | NA | No |
| 22/7/2024 | 11,094 | -74,076 | *Ikakogi tayrona* | SNSM | NA | negative | NA | No |
| 22/7/2024 | 11,094 | -74,076 | *Serranobatrachus sanctaemartae* | SNSM | negative | NA | NA | No |
| 22/7/2024 | 11,094 | -74,076 | *Serranobatrachus megalops* | SNSM | negative | NA | NA | No |
| 22/7/2024 | 11,094 | -74,076 | *Cryptobatrachus boulengeri* | SNSM | NA | negative | NA | No |
| 22/7/2024 | 11,094 | -74,076 | *Cryptobatrachus boulengeri* | SNSM | NA | negative | NA | No |
| 22/7/2024 | 11,094 | -74,076 | *Cryptobatrachus boulengeri* | SNSM | NA | negative | NA | No |
| 22/7/2024 | 11,094 | -74,076 | *Cryptobatrachus boulengeri* | SNSM | NA | negative | NA | No |

| | | | | | | | | |
|---|---|---|---|---|---|---|---|---|
| 22/7/2024 | 11,094 | -74,076 | *Cryptobatrachus boulengeri* | SNSM | negative | NA | NA | No |
| 22/7/2024 | 11,094 | -74,076 | *Cryptobatrachus boulengeri* | SNSM | NA | negative | NA | No |
| 22/7/2024 | 11,094 | -74,076 | *Cryptobatrachus boulengeri* | SNSM | NA | negative | NA | No |
| 24/7/2024 | 11,111 | -74,061 | *Atelopus nahumae* | SNSM | NA | negative | NA | No |
| 24/7/2024 | 11,111 | -74,061 | *Serranobatrachus sanctaemartae* | SNSM | negative | NA | NA | No |
| 24/7/2024 | 11,111 | -74,061 | *Atelopus nahumae* | SNSM | negative | NA | NA | No |
| 24/7/2024 | 11,111 | -74,061 | *Atelopus laetissimus* | SNSM | NA | negative | NA | No |
| 24/7/2024 | 11,111 | -74,061 | *Atelopus nahumae* | SNSM | negative | NA | NA | No |
| 24/7/2024 | 11,111 | -74,061 | *Serranobatrachus sanctaemartae* | SNSM | negative | NA | NA | No |
| 24/7/2024 | 11,111 | -74,061 | *Atelopus laetissimus* | SNSM | negative | NA | NA | No |
| 24/7/2024 | 11,111 | -74,061 | *Serranobatrachus sanctaemartae* | SNSM | negative | NA | NA | No |
| 24/7/2024 | 11,111 | -74,061 | *Serranobatrachus megalops* | SNSM | NA | negative | NA | No |
| 24/7/2024 | 11,111 | -74,061 | *Serranobatrachus megalops* | SNSM | NA | negative | NA | No |
| 24/7/2024 | 11,111 | -74,061 | *Atelopus laetissimus* | SNSM | NA | negative | NA | No |
| 24/7/2024 | 11,111 | -74,061 | *Serranobatrachus sanctaemartae* | SNSM | NA | negative | NA | No |
| 24/7/2024 | 11,111 | -74,061 | *Atelopus laetissimus* | SNSM | NA | negative | NA | No |
| 24/7/2024 | 11,111 | -74,061 | *Serranobatrachus sanctaemartae* | SNSM | negative | NA | NA | No |
| 24/7/2024 | 11,111 | -74,061 | *Atelopus laetissimus* | SNSM | negative | NA | NA | No |
| 24/7/2024 | 11,111 | -74,061 | *Serranobatrachus sanctaemartae* | SNSM | negative | NA | NA | No |
| 24/7/2024 | 11,111 | -74,061 | *Atelopus laetissimus* | SNSM | NA | negative | NA | No |
| 24/7/2024 | 11,111 | -74,061 | *Atelopus laetissimus* | SNSM | NA | negative | NA | No |
| 24/7/2024 | 11,111 | -74,061 | *Atelopus laetissimus* | SNSM | negative | NA | NA | No |
| 24/7/2024 | 11,111 | -74,061 | *Atelopus nahumae* | SNSM | negative | NA | NA | No |
| 26/7/2024 | 11,094 | -74,076 | *Ikakogi tayrona* | SNSM | negative | NA | NA | No |
| 26/7/2024 | 11,094 | -74,076 | *Serranobatrachus sanctaemartae* | SNSM | negative | NA | NA | No |
| 26/7/2024 | 11,094 | -74,076 | *Serranobatrachus megalops* | SNSM | NA | negative | NA | No |
| 26/7/2024 | 11,094 | -74,076 | *Cryptobatrachus boulengeri* | SNSM | negative | NA | NA | No |
| 26/7/2024 | 11,094 | -74,076 | *Serranobatrachus sanctaemartae* | SNSM | negative | NA | NA | No |
| 26/7/2024 | 11,094 | -74,076 | *Cryptobatrachus boulengeri* | SNSM | negative | NA | NA | No |
| 26/7/2024 | 11,094 | -74,076 | *Cryptobatrachus boulengeri* | SNSM | NA | negative | NA | No |
| 26/7/2024 | 11,094 | -74,076 | *Serranobatrachus megalops* | SNSM | negative | NA | NA | No |
| 26/7/2024 | 11,094 | -74,076 | *Cryptobatrachus boulengeri* | SNSM | negative | NA | NA | No |
| 26/7/2024 | 11,094 | -74,076 | *Cryptobatrachus boulengeri* | SNSM | negative | NA | NA | No |
| 26/7/2024 | 11,094 | -74,076 | *Serranobatrachus megalops* | SNSM | negative | NA | NA | No |
| 26/7/2024 | 11,094 | -74,076 | *Cryptobatrachus boulengeri* | SNSM | negative | NA | NA | No |
| 26/7/2024 | 11,094 | -74,076 | *Cryptobatrachus boulengeri* | SNSM | NA | negative | NA | No |
| 26/7/2024 | 11,094 | -74,076 | *Cryptobatrachus boulengeri* | SNSM | NA | negative | NA | No |
| 26/7/2024 | 11,094 | -74,076 | *Cryptobatrachus boulengeri* | SNSM | NA | negative | NA | No |

| | | | | | | | | |
|---|---|---|---|---|---|---|---|---|
| 30/7/2024 | 11,156 | -74,135 | *Leptodactylus insularum* | Foothills SNSM | negative | NA | NA | No |
| 30/7/2024 | 11,156 | -74,135 | *Leptodactylus insularum* | Foothills SNSM | negative | NA | NA | No |
| 30/7/2024 | 11,156 | -74,135 | *Boana sp.* | Foothills SNSM | NA | negative | NA | No |
| 30/7/2024 | 11,156 | -74,135 | *Leptodactylus insularum* | Foothills SNSM | negative | NA | NA | No |
| 30/7/2024 | 11,156 | -74,135 | *Leptodactylus insularum* | Foothills SNSM | negative | NA | NA | No |
| 30/7/2024 | 11,156 | -74,135 | *Colostethus ruthveni* | Foothills SNSM | negative | NA | NA | No |
| 30/7/2024 | 11,156 | -74,135 | *Boana sp.* | Foothills SNSM | negative | NA | NA | No |
| 30/7/2024 | 11,162 | -74,141 | *Leptodactylus insularum* | Foothills SNSM | negative | NA | NA | No |
| 30/7/2024 | 11,162 | -74,141 | *Colostethus ruthveni* | Foothills SNSM | negative | NA | NA | No |
| 30/7/2024 | 11,162 | -74,141 | *Leptodactylus insularum* | Foothills SNSM | negative | NA | NA | No |
| 30/7/2024 | 11,162 | -74,141 | *Leptodactylus insularum* | Foothills SNSM | negative | NA | NA | No |
| 1/8/2024 | 11,182 | -74,149 | *Engystomops pustulosus* | Foothills SNSM | NA | negative | NA | No |
| 1/8/2024 | 11,182 | -74,149 | *Engystomops pustulosus* | Foothills SNSM | negative | NA | NA | No |
| 1/8/2024 | 11,182 | -74,149 | *Engystomops pustulosus* | Foothills SNSM | NA | negative | NA | No |
| 1/8/2024 | 11,182 | -74,149 | *Engystomops pustulosus* | Foothills SNSM | negative | NA | NA | No |
| 1/8/2024 | 11,182 | -74,149 | *Engystomops pustulosus* | Foothills SNSM | negative | NA | NA | No |
| 1/8/2024 | 11,175 | -74,146 | *Engystomops pustulosus* | Foothills SNSM | negative | NA | NA | No |
| 1/8/2024 | 11,175 | -74,146 | *Engystomops pustulosus* | Foothills SNSM | negative | NA | NA | No |
| 1/8/2024 | 11,175 | -74,146 | *Engystomops pustulosus* | Foothills SNSM | negative | NA | NA | No |
| 1/8/2024 | 11,175 | -74,146 | *Engystomops pustulosus* | Foothills SNSM | negative | NA | NA | No |
| 2/8/2024 | 11,284 | -74,005 | *Engystomops pustulosus* | Foothills SNSM | negative | NA | NA | No |
| 2/8/2024 | 11,284 | -74,005 | *Engystomops pustulosus* | Foothills SNSM | negative | NA | NA | No |
| 2/8/2024 | 11,284 | -74,005 | *Rhinella horribilis* | Foothills SNSM | negative | NA | NA | No |
| 2/8/2024 | 11,284 | -74,005 | *Engystomops pustulosus* | Foothills SNSM | negative | NA | NA | No |
| 2/8/2024 | 11,284 | -74,005 | *Engystomops pustulosus* | Foothills SNSM | negative | NA | NA | No |
| 2/8/2024 | 11,284 | -74,005 | *Engystomops pustulosus* | Foothills SNSM | negative | NA | NA | No |
| 2/8/2024 | 11,284 | -74,005 | *Engystomops pustulosus* | Foothills SNSM | negative | NA | NA | No |
| 2/8/2024 | 11,284 | -74,005 | *Engystomops pustulosus* | Foothills SNSM | negative | NA | NA | No |
| 2/8/2024 | 11,284 | -74,005 | *Engystomops pustulosus* | Foothills SNSM | negative | NA | NA | No |
| 2/8/2024 | 11,284 | -74,005 | *Engystomops pustulosus* | Foothills SNSM | negative | NA | NA | No |
| 2/8/2024 | 11,284 | -74,005 | *Engystomops pustulosus* | Foothills SNSM | negative | NA | NA | No |
| 2/8/2024 | 11,284 | -74,005 | *Engystomops pustulosus* | Foothills SNSM | negative | NA | NA | No |
| 2/8/2024 | 11,284 | -74,005 | *Engystomops pustulosus* | Foothills SNSM | negative | NA | NA | No |
| 2/8/2024 | 11,284 | -74,005 | *Engystomops pustulosus* | Foothills SNSM | negative | NA | NA | No |
| 2/8/2024 | 11,284 | -74,005 | *Engystomops pustulosus* | Foothills SNSM | negative | NA | NA | No |
| 2/8/2024 | 11,284 | -74,005 | *Engystomops pustulosus* | Foothills SNSM | negative | NA | NA | No |
| 2/8/2024 | 11,284 | -74,005 | *Engystomops pustulosus* | Foothills SNSM | negative | NA | NA | No |

| | | | | | | | | |
|---|---|---|---|---|---|---|---|---|
| 2/8/2024 | 11,284 | -74,005 | *Boana sp.* | Foothills SNSM | negative | NA | NA | No |
| 2/8/2024 | 11,284 | -74,005 | *Boana sp.* | Foothills SNSM | NA | negative | NA | No |
| 2/8/2024 | 11,284 | -74,005 | *Engystomops pustulosus* | Foothills SNSM | negative | NA | NA | No |
| 2/8/2024 | 11,284 | -74,005 | *Boana sp.* | Foothills SNSM | negative | NA | NA | No |
| 2/8/2024 | 11,284 | -74,005 | *Leptodactylus insularum* | Foothills SNSM | NA | negative | NA | No |
| 2/8/2024 | 11,284 | -74,005 | *Boana sp.* | Foothills SNSM | negative | NA | NA | No |
| 2/8/2024 | 11,284 | -74,005 | *Boana sp.* | Foothills SNSM | negative | NA | NA | No |
| 2/8/2024 | 11,284 | -74,005 | *Boana sp.* | Foothills SNSM | negative | NA | NA | No |
| 2/8/2024 | 11,284 | -74,005 | *Rhinella horribilis* | Foothills SNSM | negative | NA | NA | No |
| 2/8/2024 | 11,284 | -74,005 | *Boana sp.* | Foothills SNSM | negative | NA | NA | No |
| 2/8/2024 | 11,284 | -74,005 | *Engystomops pustulosus* | Foothills SNSM | negative | NA | NA | No |
| 2/8/2024 | 11,284 | -74,005 | *Engystomops pustulosus* | Foothills SNSM | negative | NA | NA | No |
| 2/8/2024 | 11,284 | -74,005 | *Engystomops pustulosus* | Foothills SNSM | negative | NA | NA | No |
| 2/8/2024 | 11,284 | -74,005 | *Engystomops pustulosus* | Foothills SNSM | negative | NA | NA | No |
| 2/8/2024 | 11,284 | -74,005 | *Engystomops pustulosus* | Foothills SNSM | negative | NA | NA | No |
| 2/8/2024 | 11,284 | -74,005 | *Engystomops pustulosus* | Foothills SNSM | negative | NA | NA | No |
| 2/8/2024 | 11,284 | -74,005 | *Engystomops pustulosus* | Foothills SNSM | negative | NA | NA | No |
| 2/8/2024 | 11,284 | -74,005 | *Engystomops pustulosus* | Foothills SNSM | negative | NA | NA | No |
| 2/8/2024 | 11,284 | -74,005 | *Engystomops pustulosus* | Foothills SNSM | negative | NA | NA | No |
| 2/8/2024 | 11,27 | -74,068 | *Dendrobates truncatus* | Foothills SNSM | NA | negative | NA | No |
| 2/8/2024 | 11,27 | -74,068 | *Dendrobates truncatus* | Foothills SNSM | negative | NA | NA | No |
| 2/8/2024 | 11,285 | -73,99 | *Engystomops pustulosus* | Foothills SNSM | negative | NA | NA | No |
| 2/8/2024 | 11,285 | -73,99 | *Leptodactylus sp.* | Foothills SNSM | negative | NA | NA | No |
| 2/8/2024 | 11,285 | -73,99 | *Leptodactylus sp.* | Foothills SNSM | negative | NA | NA | No |
| 2/8/2024 | 11,285 | -73,99 | *Leptodactylus sp.* | Foothills SNSM | negative | NA | NA | No |
| 2/8/2024 | 11,285 | -73,99 | *Leptodactylus sp.* | Foothills SNSM | negative | NA | NA | No |
| 2/8/2024 | 11,285 | -73,99 | *Leptodactylus sp.* | Foothills SNSM | negative | NA | NA | No |
| 2/8/2024 | 11,285 | -73,99 | *Leptodactylus sp.* | Foothills SNSM | negative | NA | NA | No |
| 2/8/2024 | 11,285 | -73,99 | *Leptodactylus sp.* | Foothills SNSM | NA | negative | NA | No |
| 2/8/2024 | 11,285 | -73,99 | *Leptodactylus sp.* | Foothills SNSM | negative | NA | NA | No |
| 2/8/2024 | 11,285 | -73,99 | *Leptodactylus sp.* | Foothills SNSM | NA | negative | NA | No |
| 3/8/2024 | 11,282 | -73,971 | *Rhinella horribilis* | Foothills SNSM | negative | NA | NA | No |
| 3/8/2024 | 11,282 | -73,971 | *Engystomops pustulosus* | Foothills SNSM | negative | NA | NA | No |
| 3/8/2024 | 11,282 | -73,971 | *Engystomops pustulosus* | Foothills SNSM | negative | NA | NA | No |
| 3/8/2024 | 11,282 | -73,971 | *Boana sp.* | Foothills SNSM | negative | NA | NA | No |
| 3/8/2024 | 11,282 | -73,971 | *Engystomops pustulosus* | Foothills SNSM | negative | NA | NA | No |
| 3/8/2024 | 11,282 | -73,971 | *Engystomops pustulosus* | Foothills SNSM | negative | NA | NA | No |

| | | | | | | | | |
|---|---|---|---|---|---|---|---|---|
| 3/8/2024 | 11,282 | -73,971 | *Engystomops pustulosus* | Foothills SNSM | negative | NA | NA | No |
| 3/8/2024 | 11,282 | -73,971 | *Engystomops pustulosus* | Foothills SNSM | negative | NA | NA | No |
| 3/8/2024 | 11,282 | -73,971 | *Rhinella horribilis* | Foothills SNSM | negative | NA | NA | No |
| 3/8/2024 | 11,282 | -73,971 | *Cryptobatrachus boulengeri* | Foothills SNSM | negative | NA | NA | No |
| 3/8/2024 | 11,282 | -73,971 | *Cryptobatrachus boulengeri* | Foothills SNSM | negative | NA | NA | No |
| 3/8/2024 | 11,282 | -73,971 | *Engystomops pustulosus* | Foothills SNSM | negative | NA | NA | No |
| 3/8/2024 | 11,282 | -73,971 | *Engystomops pustulosus* | Foothills SNSM | NA | negative | NA | No |
| 3/8/2024 | 11,282 | -73,971 | *Engystomops pustulosus* | Foothills SNSM | negative | NA | NA | No |
| 3/8/2024 | 11,282 | -73,971 | *Engystomops pustulosus* | Foothills SNSM | negative | NA | NA | No |
| 3/8/2024 | 11,282 | -73,971 | *Rhinella horribilis* | Foothills SNSM | negative | NA | NA | No |
| 3/8/2024 | 11,282 | -73,971 | *Leptodactylus insularum* | Foothills SNSM | negative | NA | NA | No |
| 3/8/2024 | 11,282 | -73,971 | *Cryptobatrachus boulengeri* | Foothills SNSM | negative | NA | NA | No |
| 3/8/2024 | 11,282 | -73,971 | *Leptodactylus insularum* | Foothills SNSM | negative | NA | NA | No |
| 3/8/2024 | 11,282 | -73,971 | *Rhinella horribilis* | Foothills SNSM | negative | NA | NA | No |
| 3/8/2024 | 11,282 | -73,971 | *Rhinella horribilis* | Foothills SNSM | negative | NA | NA | No |
| 5/8/2024 | 11,235 | -74,118 | *Boana sp.* | Foothills SNSM | negative | NA | NA | No |
| 5/8/2024 | 11,235 | -74,118 | *Engystomops pustulosus* | Foothills SNSM | negative | NA | NA | No |
| 5/8/2024 | 11,235 | -74,118 | *Engystomops pustulosus* | Foothills SNSM | negative | NA | NA | No |
| 5/8/2024 | 11,235 | -74,118 | *Engystomops pustulosus* | Foothills SNSM | negative | NA | NA | No |
| 5/8/2024 | 11,235 | -74,118 | *Engystomops pustulosus* | Foothills SNSM | NA | negative | NA | No |
| 5/8/2024 | 11,235 | -74,118 | *Engystomops pustulosus* | Foothills SNSM | negative | NA | NA | No |
| 5/8/2024 | 11,235 | -74,118 | *Engystomops pustulosus* | Foothills SNSM | negative | NA | NA | No |
| 5/8/2024 | 11,235 | -74,118 | *Engystomops pustulosus* | Foothills SNSM | negative | NA | NA | No |
| 5/8/2024 | 11,235 | -74,118 | *Engystomops pustulosus* | Foothills SNSM | NA | negative | NA | No |
| 5/8/2024 | 11,235 | -74,118 | *Engystomops pustulosus* | Foothills SNSM | negative | NA | NA | No |
| 5/8/2024 | 11,235 | -74,118 | *Engystomops pustulosus* | Foothills SNSM | negative | NA | NA | No |
| 5/8/2024 | 11,235 | -74,118 | *Engystomops pustulosus* | Foothills SNSM | negative | NA | NA | No |
| 5/8/2024 | 11,235 | -74,118 | *Engystomops pustulosus* | Foothills SNSM | NA | negative | NA | No |
| 5/8/2024 | 11,235 | -74,118 | *Engystomops pustulosus* | Foothills SNSM | negative | NA | NA | No |
| 5/8/2024 | 11,235 | -74,118 | *Engystomops pustulosus* | Foothills SNSM | negative | NA | NA | No |
| 5/8/2024 | 11,235 | -74,118 | *Engystomops pustulosus* | Foothills SNSM | negative | NA | NA | No |
| 5/8/2024 | 11,235 | -74,118 | *Engystomops pustulosus* | Foothills SNSM | negative | NA | NA | No |
| 5/8/2024 | 11,235 | -74,118 | *Engystomops pustulosus* | Foothills SNSM | negative | NA | NA | No |
| 5/8/2024 | 11,235 | -74,118 | *Engystomops pustulosus* | Foothills SNSM | NA | negative | NA | No |
| 5/8/2024 | 11,235 | -74,118 | *Engystomops pustulosus* | Foothills SNSM | negative | NA | NA | No |
| 5/8/2024 | 11,235 | -74,118 | *Engystomops pustulosus* | Foothills SNSM | negative | NA | NA | No |
| 5/8/2024 | 11,235 | -74,118 | *Engystomops pustulosus* | Foothills SNSM | negative | NA | NA | No |

| | | | | | | | | |
|---|---|---|---|---|---|---|---|---|
| 5/8/2024 | 11,235 | -74,118 | *Engystomops pustulosus* | Foothills SNSM | negative | NA | NA | No |
| 5/8/2024 | 11,235 | -74,118 | *Engystomops pustulosus* | Foothills SNSM | negative | NA | NA | No |
| 13/8/2024 | 11,094 | -74,076 | *Serranobatrachus sanctaemartae* | SNSM | negative | NA | NA | No |
| 13/8/2024 | 11,094 | -74,076 | *Ikakogi tayrona* | SNSM | negative | NA | NA | No |
| 13/8/2024 | 11,094 | -74,076 | *Ikakogi tayrona* | SNSM | negative | NA | NA | No |
| 13/8/2024 | 11,094 | -74,076 | *Serranobatrachus sanctaemartae* | SNSM | negative | NA | NA | No |
| 13/8/2024 | 11,094 | -74,076 | *Cryptobatrachus boulengeri* | SNSM | negative | NA | NA | No |
| 13/8/2024 | 11,094 | -74,076 | *Serranobatrachus sanctaemartae* | SNSM | negative | NA | NA | No |
| 13/8/2024 | 11,094 | -74,076 | *Cryptobatrachus boulengeri* | SNSM | negative | NA | NA | No |
| 13/8/2024 | 11,094 | -74,076 | *Cryptobatrachus boulengeri* | SNSM | negative | NA | NA | No |
| 13/8/2024 | 11,094 | -74,076 | *Cryptobatrachus boulengeri* | SNSM | negative | NA | NA | No |
| 13/8/2024 | 11,094 | -74,076 | *Cryptobatrachus boulengeri* | SNSM | negative | NA | NA | No |
| 13/8/2024 | 11,094 | -74,076 | *Cryptobatrachus boulengeri* | SNSM | negative | NA | NA | No |
| 13/8/2024 | 11,094 | -74,076 | *Cryptobatrachus boulengeri* | SNSM | negative | NA | NA | No |
| 13/8/2024 | 11,094 | -74,076 | *Cryptobatrachus boulengeri* | SNSM | negative | NA | NA | No |
| 13/8/2024 | 11,094 | -74,076 | *Cryptobatrachus boulengeri* | SNSM | negative | NA | NA | No |
| 13/8/2024 | 11,094 | -74,076 | *Cryptobatrachus boulengeri* | SNSM | negative | NA | NA | No |
| 13/8/2024 | 11,094 | -74,076 | *Serranobatrachus sanctaemartae* | SNSM | negative | NA | NA | No |
| 13/8/2024 | 11,094 | -74,076 | *Serranobatrachus sanctaemartae* | SNSM | negative | NA | NA | No |
| 13/8/2024 | 11,094 | -74,076 | *Cryptobatrachus boulengeri* | SNSM | negative | NA | NA | No |
| 13/8/2024 | 11,094 | -74,076 | *Cryptobatrachus boulengeri* | SNSM | negative | NA | NA | No |
| 13/8/2024 | 11,094 | -74,076 | *Cryptobatrachus boulengeri* | SNSM | negative | NA | NA | No |
| 13/8/2024 | 11,094 | -74,076 | *Serranobatrachus sanctaemartae* | SNSM | negative | NA | NA | No |
| 13/8/2024 | 11,094 | -74,076 | *Cryptobatrachus boulengeri* | SNSM | negative | NA | NA | No |
| 13/8/2024 | 11,094 | -74,076 | *Serranobatrachus sanctaemartae* | SNSM | negative | NA | NA | No |
| 13/8/2024 | 11,094 | -74,076 | *Serranobatrachus sanctaemartae* | SNSM | negative | NA | NA | No |
| 13/8/2024 | 11,094 | -74,076 | *Cryptobatrachus boulengeri* | SNSM | negative | NA | NA | No |
| 13/8/2024 | 11,094 | -74,076 | *Cryptobatrachus boulengeri* | SNSM | negative | NA | NA | No |
| 13/8/2024 | 11,094 | -74,076 | *Cryptobatrachus boulengeri* | SNSM | negative | NA | NA | No |
| 16/8/2024 | 10,902 | -74,154 | *Pleurodema brachyops* | Foothills SNSM | negative | NA | NA | No |
| 16/8/2024 | 10,902 | -74,154 | *Pleurodema brachyops* | Foothills SNSM | negative | NA | NA | No |
| 16/8/2024 | 10,902 | -74,154 | *Pleurodema brachyops* | Foothills SNSM | negative | NA | NA | No |
| 16/8/2024 | 10,902 | -74,154 | *Pleurodema brachyops* | Foothills SNSM | negative | NA | NA | No |
| 16/8/2024 | 10,902 | -74,154 | *Pleurodema brachyops* | Foothills SNSM | negative | NA | NA | No |
| 16/8/2024 | 10,902 | -74,154 | *Rhinella humboldti* | Foothills SNSM | negative | NA | NA | No |
| 16/8/2024 | 10,902 | -74,154 | *Rhinella humboldti* | Foothills SNSM | negative | NA | NA | No |
| 16/8/2024 | 10,902 | -74,154 | *Rhinella humboldti* | Foothills SNSM | negative | NA | NA | No |
| 16/8/2024 | 10,902 | -74,154 | *Pleurodema brachyops* | Foothills SNSM | negative | NA | NA | No |

| | | | | | | | | |
|---|---|---|---|---|---|---|---|---|
| 16/8/2024 | 10,902 | -74,154 | *Pleurodema brachyops* | Foothills SNSM | negative | NA | NA | No |
| 16/8/2024 | 10,902 | -74,154 | *Pleurodema brachyops* | Foothills SNSM | negative | NA | NA | No |
| 16/8/2024 | 10,902 | -74,154 | *Rhinella humboldti* | Foothills SNSM | negative | NA | NA | No |
| 16/8/2024 | 10,902 | -74,154 | *Rhinella horribilis* | Foothills SNSM | negative | NA | NA | No |
| 16/8/2024 | 10,902 | -74,154 | *Rhinella horribilis* | Foothills SNSM | negative | NA | NA | No |
| 16/8/2024 | 10,902 | -74,154 | *Pleurodema brachyops* | Foothills SNSM | negative | NA | NA | No |
| 16/8/2024 | 10,902 | -74,154 | *Pleurodema brachyops* | Foothills SNSM | negative | NA | NA | No |
| 16/8/2024 | 10,902 | -74,154 | *Pleurodema brachyops* | Foothills SNSM | negative | NA | NA | No |
| 16/8/2024 | 10,902 | -74,154 | *Pleurodema brachyops* | Foothills SNSM | negative | NA | NA | No |
| 16/8/2024 | 10,902 | -74,154 | *Rhinella humboldti* | Foothills SNSM | negative | NA | NA | No |
| 16/8/2024 | 10,902 | -74,154 | *Rhinella humboldti* | Foothills SNSM | negative | NA | NA | No |
| 16/8/2024 | 10,902 | -74,154 | *Rhinella horribilis* | Foothills SNSM | negative | NA | NA | No |
| 16/8/2024 | 10,902 | -74,154 | *Pleurodema brachyops* | Foothills SNSM | negative | NA | NA | No |
| 16/8/2024 | 10,902 | -74,154 | *Pleurodema brachyops* | Foothills SNSM | negative | NA | NA | No |
| 16/8/2024 | 10,902 | -74,154 | *Pleurodema brachyops* | Foothills SNSM | negative | NA | NA | No |
| 16/8/2024 | 10,902 | -74,154 | *Pleurodema brachyops* | Foothills SNSM | negative | NA | NA | No |
| 16/8/2024 | 10,902 | -74,154 | *Rhinella humboldti* | Foothills SNSM | negative | NA | NA | No |
| 16/8/2024 | 10,902 | -74,154 | *Pleurodema brachyops* | Foothills SNSM | NA | negative | NA | No |
| 16/8/2024 | 10,902 | -74,154 | *Pleurodema brachyops* | Foothills SNSM | NA | negative | NA | No |
| 16/8/2024 | 10,902 | -74,154 | *Pleurodema brachyops* | Foothills SNSM | negative | NA | NA | No |
| 16/8/2024 | 10,902 | -74,154 | *Pleurodema brachyops* | Foothills SNSM | negative | NA | NA | No |
| 17/8/2024 | 10,829 | -74,12 | *Pleurodema brachyops* | Foothills SNSM | negative | NA | NA | No |
| 17/8/2024 | 10,829 | -74,12 | *Engystomops pustulosus* | Foothills SNSM | negative | NA | NA | No |
| 17/8/2024 | 10,829 | -74,12 | *Rhinella horribilis* | Foothills SNSM | negative | NA | NA | No |
| 17/8/2024 | 10,829 | -74,12 | *Engystomops pustulosus* | Foothills SNSM | negative | NA | NA | No |
| 17/8/2024 | 10,829 | -74,12 | *Engystomops pustulosus* | Foothills SNSM | negative | NA | NA | No |
| 17/8/2024 | 10,829 | -74,12 | *Pleurodema brachyops* | Foothills SNSM | negative | NA | NA | No |
| 17/8/2024 | 10,829 | -74,12 | *Pleurodema brachyops* | Foothills SNSM | negative | NA | NA | No |
| 17/8/2024 | 10,829 | -74,12 | *Pleurodema brachyops* | Foothills SNSM | negative | NA | NA | No |
| 17/8/2024 | 10,829 | -74,12 | *Pleurodema brachyops* | Foothills SNSM | negative | NA | NA | No |
| 17/8/2024 | 10,829 | -74,12 | *Engystomops pustulosus* | Foothills SNSM | negative | NA | NA | No |
| 17/8/2024 | 10,829 | -74,12 | *Engystomops pustulosus* | Foothills SNSM | negative | NA | NA | No |
| 17/8/2024 | 10,829 | -74,12 | *Engystomops pustulosus* | Foothills SNSM | negative | NA | NA | No |
| 17/8/2024 | 10,829 | -74,12 | *Engystomops pustulosus* | Foothills SNSM | negative | NA | NA | No |
| 17/8/2024 | 10,829 | -74,12 | *Engystomops pustulosus* | Foothills SNSM | negative | NA | NA | No |
| 17/8/2024 | 10,829 | -74,12 | *Engystomops pustulosus* | Foothills SNSM | negative | NA | NA | No |
| 17/8/2024 | 10,829 | -74,12 | *Rhinella horribilis* | Foothills SNSM | negative | NA | NA | No |

| | | | | | | | | |
|---|---|---|---|---|---|---|---|---|
| 17/8/2024 | 10,829 | -74,12 | *Engystomops pustulosus* | Foothills SNSM | negative | NA | NA | No |
| 17/8/2024 | 10,829 | -74,12 | *Engystomops pustulosus* | Foothills SNSM | negative | NA | NA | No |
| 17/8/2024 | 10,829 | -74,12 | *Engystomops pustulosus* | Foothills SNSM | negative | NA | NA | No |
| 17/8/2024 | 10,829 | -74,12 | *Engystomops pustulosus* | Foothills SNSM | negative | NA | NA | No |
| 17/8/2024 | 10,829 | -74,12 | *Engystomops pustulosus* | Foothills SNSM | negative | NA | NA | No |
| 17/8/2024 | 10,829 | -74,12 | *Engystomops pustulosus* | Foothills SNSM | negative | NA | NA | No |
| 17/8/2024 | 10,829 | -74,12 | *Engystomops pustulosus* | Foothills SNSM | negative | NA | NA | No |
| 17/8/2024 | 10,829 | -74,12 | *Engystomops pustulosus* | Foothills SNSM | negative | NA | NA | No |
| 17/8/2024 | 10,829 | -74,12 | *Engystomops pustulosus* | Foothills SNSM | negative | NA | NA | No |
| 17/8/2024 | 10,829 | -74,12 | *Engystomops pustulosus* | Foothills SNSM | negative | NA | NA | No |
| 17/8/2024 | 10,829 | -74,12 | *Engystomops pustulosus* | Foothills SNSM | negative | NA | NA | No |
| 17/8/2024 | 10,829 | -74,12 | *Engystomops pustulosus* | Foothills SNSM | negative | NA | NA | No |
| 17/8/2024 | 10,829 | -74,12 | *Engystomops pustulosus* | Foothills SNSM | negative | NA | NA | No |
| 17/8/2024 | 10,829 | -74,12 | *Engystomops pustulosus* | Foothills SNSM | negative | NA | NA | No |
| 17/8/2024 | 10,829 | -74,12 | *Engystomops pustulosus* | Foothills SNSM | negative | NA | NA | No |
| 17/8/2024 | 10,755 | -74,097 | *Engystomops pustulosus* | Foothills SNSM | negative | NA | NA | No |
| 17/8/2024 | 10,755 | -74,097 | *Engystomops pustulosus* | Foothills SNSM | negative | NA | NA | No |
| 17/8/2024 | 10,755 | -74,097 | *Engystomops pustulosus* | Foothills SNSM | negative | NA | NA | No |
| 17/8/2024 | 10,755 | -74,097 | *Engystomops pustulosus* | Foothills SNSM | negative | NA | NA | No |
| 17/8/2024 | 10,755 | -74,097 | *Engystomops pustulosus* | Foothills SNSM | negative | NA | NA | No |
| 17/8/2024 | 10,755 | -74,097 | *Engystomops pustulosus* | Foothills SNSM | negative | NA | NA | No |
| 17/8/2024 | 10,755 | -74,097 | *Engystomops pustulosus* | Foothills SNSM | negative | NA | NA | No |
| 17/8/2024 | 10,755 | -74,097 | *Engystomops pustulosus* | Foothills SNSM | NA | negative | NA | No |
| 17/8/2024 | 10,755 | -74,097 | *Engystomops pustulosus* | Foothills SNSM | negative | NA | NA | No |
| 17/8/2024 | 10,755 | -74,097 | *Boana pugnax* | Foothills SNSM | negative | NA | NA | No |
| 17/8/2024 | 10,755 | -74,097 | *Engystomops pustulosus* | Foothills SNSM | negative | NA | NA | No |
| 17/8/2024 | 10,755 | -74,097 | *Engystomops pustulosus* | Foothills SNSM | negative | NA | NA | No |
| 17/8/2024 | 10,755 | -74,097 | *Engystomops pustulosus* | Foothills SNSM | negative | NA | NA | No |
| 17/8/2024 | 10,755 | -74,097 | *Engystomops pustulosus* | Foothills SNSM | negative | NA | NA | No |
| 17/8/2024 | 10,755 | -74,097 | *Engystomops pustulosus* | Foothills SNSM | negative | NA | NA | No |
| 17/8/2024 | 10,755 | -74,097 | *Engystomops pustulosus* | Foothills SNSM | NA | negative | NA | No |
| 17/8/2024 | 10,755 | -74,097 | *Engystomops pustulosus* | Foothills SNSM | negative | NA | NA | No |
| 17/8/2024 | 10,755 | -74,097 | *Engystomops pustulosus* | Foothills SNSM | negative | NA | NA | No |
| 17/8/2024 | 10,755 | -74,097 | *Engystomops pustulosus* | Foothills SNSM | negative | NA | NA | No |
| 17/8/2024 | 10,755 | -74,097 | *Engystomops pustulosus* | Foothills SNSM | negative | NA | NA | No |
| 19/8/2024 | 10,3 | -74,041 | *Pleurodema brachyops* | Foothills SNSM | negative | NA | NA | No |
| 19/8/2024 | 10,3 | -74,041 | *Pleurodema brachyops* | Foothills SNSM | negative | NA | NA | No |

| | | | | | | | | |
|---|---|---|---|---|---|---|---|---|
| 19/8/2024 | 10,591 | -74,169 | *Rhinella horribilis* | Foothills SNSM | negative | NA | NA | No |
| 19/8/2024 | 10,591 | -74,169 | *Boana pugnax* | Foothills SNSM | negative | NA | NA | No |
| 19/8/2024 | 10,591 | -74,169 | *Pleurodema brachyops* | Foothills SNSM | negative | NA | NA | No |
| 19/8/2024 | 10,591 | -74,169 | *Leptodactylus sp.* | Foothills SNSM | negative | NA | NA | No |
| 19/8/2024 | 10,591 | -74,169 | *Pleurodema brachyops* | Foothills SNSM | negative | NA | NA | No |
| 19/8/2024 | 10,591 | -74,169 | *Pleurodema brachyops* | Foothills SNSM | negative | NA | NA | No |
| 19/8/2024 | 10,591 | -74,169 | *Pleurodema brachyops* | Foothills SNSM | negative | NA | NA | No |
| 19/8/2024 | 10,591 | -74,169 | *Engystomops pustulosus* | Foothills SNSM | negative | NA | NA | No |
| 19/8/2024 | 10,591 | -74,169 | *Rhinella humboldti* | Foothills SNSM | negative | NA | NA | No |
| 19/8/2024 | 10,591 | -74,169 | *Engystomops pustulosus* | Foothills SNSM | negative | NA | NA | No |
| 19/8/2024 | 10,591 | -74,169 | *Rhinella humboldti* | Foothills SNSM | negative | NA | NA | No |
| 19/8/2024 | 10,591 | -74,169 | *Rhinella humboldti* | Foothills SNSM | negative | NA | NA | No |
| 19/8/2024 | 10,591 | -74,169 | *Pleurodema brachyops* | Foothills SNSM | negative | NA | NA | No |
| 19/8/2024 | 10,591 | -74,169 | *Pleurodema brachyops* | Foothills SNSM | negative | NA | NA | No |
| 19/8/2024 | 10,591 | -74,169 | *Pleurodema brachyops* | Foothills SNSM | negative | NA | NA | No |
| 19/8/2024 | 10,591 | -74,169 | *Pleurodema brachyops* | Foothills SNSM | negative | NA | NA | No |
| 19/8/2024 | 10,591 | -74,169 | *Pleurodema brachyops* | Foothills SNSM | negative | NA | NA | No |
| 19/8/2024 | 10,591 | -74,169 | *Rhinella humboldti* | Foothills SNSM | negative | NA | NA | No |
| 19/8/2024 | 10,591 | -74,169 | *Engystomops pustulosus* | Foothills SNSM | NA | negative | NA | No |
| 19/8/2024 | 10,591 | -74,169 | *Rhinella horribilis* | Foothills SNSM | negative | NA | NA | No |
| 19/8/2024 | 10,591 | -74,169 | *Pleurodema brachyops* | Foothills SNSM | negative | NA | NA | No |
| 19/8/2024 | 10,591 | -74,169 | *Engystomops pustulosus* | Foothills SNSM | negative | NA | NA | No |
| 19/8/2024 | 10,591 | -74,169 | *Rhinella horribilis* | Foothills SNSM | negative | NA | NA | No |
| 19/8/2024 | 10,591 | -74,169 | *Pleurodema brachyops* | Foothills SNSM | negative | NA | NA | No |
| 19/8/2024 | 10,591 | -74,169 | *Rhinella horribilis* | Foothills SNSM | negative | NA | NA | No |
| 19/8/2024 | 10,591 | -74,169 | *Pleurodema brachyops* | Foothills SNSM | negative | NA | NA | No |
| 19/8/2024 | 10,591 | -74,169 | *Pleurodema brachyops* | Foothills SNSM | negative | NA | NA | No |
| 19/8/2024 | 10,591 | -74,169 | *Pleurodema brachyops* | Foothills SNSM | negative | NA | NA | No |
| 19/8/2024 | 10,591 | -74,169 | *Dendropsophus microcephalus* | Foothills SNSM | negative | NA | NA | No |
| 19/8/2024 | 10,591 | -74,169 | *Leptodactylus fuscus* | Foothills SNSM | negative | NA | NA | No |
| 19/8/2024 | 10,591 | -74,169 | *Rhinella horribilis* | Foothills SNSM | negative | NA | NA | No |
| 20/8/2024 | 11,111 | -74,061 | *Serranobatrachus megalops* | SNSM | negative | NA | NA | No |
| 20/8/2024 | 11,111 | -74,061 | *Serranobatrachus sanctaemartae* | SNSM | negative | NA | NA | No |
| 20/8/2024 | 11,111 | -74,061 | *Serranobatrachus sanctaemartae* | SNSM | negative | NA | NA | No |
| 20/8/2024 | 11,111 | -74,061 | *Atelopus laetissimus* | SNSM | negative | NA | NA | No |
| 20/8/2024 | 11,111 | -74,061 | *Serranobatrachus megalops* | SNSM | negative | NA | NA | No |
| 20/8/2024 | 11,111 | -74,061 | *Serranobatrachus megalops* | SNSM | negative | NA | NA | No |

| | | | | | | | | |
|---|---|---|---|---|---|---|---|---|
| 20/8/2024 | 11,111 | -74,061 | *Atelopus laetissimus* | SNSM | negative | NA | NA | No |
| 20/8/2024 | 11,111 | -74,061 | *Serranobatrachus megalops* | SNSM | negative | NA | NA | No |
| 20/8/2024 | 11,111 | -74,061 | *Serranobatrachus megalops* | SNSM | negative | NA | NA | No |
| 20/8/2024 | 11,111 | -74,061 | *Serranobatrachus carmelitae* | SNSM | negative | NA | NA | No |
| 20/8/2024 | 11,111 | -74,061 | *Serranobatrachus megalops* | SNSM | negative | NA | NA | No |
| 20/8/2024 | 11,111 | -74,061 | *Serranobatrachus megalops* | SNSM | negative | NA | NA | No |
| 20/8/2024 | 11,111 | -74,061 | *Atelopus laetissimus* | SNSM | negative | NA | NA | No |
| 20/8/2024 | 11,111 | -74,061 | *Geobatrachus walkeri* | SNSM | negative | NA | NA | No |
| 20/8/2024 | 11,111 | -74,061 | *Serranobatrachus sanctaemartae* | SNSM | negative | NA | NA | No |
| 20/8/2024 | 11,111 | -74,061 | *Geobatrachus walkeri* | SNSM | negative | NA | NA | No |
| 20/8/2024 | 11,111 | -74,061 | *Serranobatrachus sanctaemartae* | SNSM | negative | NA | NA | No |
| 20/8/2024 | 11,111 | -74,061 | *Serranobatrachus megalops* | SNSM | negative | NA | NA | No |
| 20/8/2024 | 11,111 | -74,061 | *Ikakogi tayrona* | SNSM | negative | NA | NA | No |
| 20/8/2024 | 11,111 | -74,061 | *Atelopus laetissimus* | SNSM | negative | NA | NA | No |
| 20/8/2024 | 11,111 | -74,061 | *Serranobatrachus sanctaemartae* | SNSM | negative | NA | NA | No |
| 20/8/2024 | 11,111 | -74,061 | *Serranobatrachus sanctaemartae* | SNSM | negative | NA | NA | No |
| 20/8/2024 | 11,111 | -74,061 | *Serranobatrachus sanctaemartae* | SNSM | negative | NA | NA | No |
| 20/8/2024 | 11,111 | -74,061 | *Ikakogi tayrona* | SNSM | negative | NA | NA | No |
| 20/8/2024 | 11,111 | -74,061 | *Serranobatrachus cristinae* | SNSM | negative | NA | NA | No |
| 20/8/2024 | 11,111 | -74,061 | *Atelopus laetissimus* | SNSM | negative | NA | NA | No |
| 20/8/2024 | 11,111 | -74,061 | *Serranobatrachus megalops* | SNSM | negative | NA | NA | No |
| 20/8/2024 | 11,111 | -74,061 | *Serranobatrachus sanctaemartae* | SNSM | negative | NA | NA | No |
| 20/8/2024 | 11,111 | -74,061 | *Serranobatrachus sanctaemartae* | SNSM | negative | NA | NA | No |
| 20/8/2024 | 11,111 | -74,061 | *Atelopus laetissimus* | SNSM | negative | NA | NA | No |
| 20/8/2024 | 11,111 | -74,061 | *Serranobatrachus sanctaemartae* | SNSM | negative | NA | NA | No |
| 20/8/2024 | 11,111 | -74,061 | *Serranobatrachus cristinae* | SNSM | negative | NA | NA | No |
| 20/8/2024 | 11,111 | -74,061 | *Serranobatrachus megalops* | SNSM | negative | NA | NA | No |
| 20/8/2024 | 11,111 | -74,061 | *Atelopus laetissimus* | SNSM | negative | NA | NA | No |
| 20/8/2024 | 11,111 | -74,061 | *Atelopus laetissimus* | SNSM | negative | NA | NA | No |
| 20/8/2024 | 11,111 | -74,061 | *Serranobatrachus sanctaemartae* | SNSM | negative | NA | NA | No |
| 20/8/2024 | 11,111 | -74,061 | *Serranobatrachus sanctaemartae* | SNSM | negative | NA | NA | No |
| 20/8/2024 | 11,111 | -74,061 | *Serranobatrachus sanctaemartae* | SNSM | negative | NA | NA | No |
| 24/8/2024 | 11,285 | -73,99 | *Rhinella horribilis* | Foothills SNSM | negative | NA | NA | No |
| 24/8/2024 | 11,285 | -73,99 | *Engystomops pustulosus* | Foothills SNSM | negative | NA | NA | No |
| 24/8/2024 | 11,285 | -73,99 | *Engystomops pustulosus* | Foothills SNSM | negative | NA | NA | No |
| 24/8/2024 | 11,285 | -73,99 | *Rhinella horribilis* | Foothills SNSM | negative | NA | NA | No |
| 24/8/2024 | 11,285 | -73,99 | *Leptodactylus fuscus* | Foothills SNSM | NA | negative | NA | No |
| 24/8/2024 | 11,285 | -73,99 | *Engystomops pustulosus* | Foothills SNSM | negative | NA | NA | No |

| | | | | | | | | |
|---|---|---|---|---|---|---|---|---|
| 24/8/2024 | 11,285 | -73,99 | *Engystomops pustulosus* | Foothills SNSM | negative | NA | NA | No |
| 24/8/2024 | 11,285 | -73,99 | *Leptodactylus fuscus* | Foothills SNSM | negative | NA | NA | No |
| 24/8/2024 | 11,285 | -73,99 | *Engystomops pustulosus* | Foothills SNSM | negative | NA | NA | No |
| 24/8/2024 | 11,285 | -73,99 | *Engystomops pustulosus* | Foothills SNSM | negative | NA | NA | No |
| 24/8/2024 | 11,285 | -73,99 | *Engystomops pustulosus* | Foothills SNSM | negative | NA | NA | No |
| 24/8/2024 | 11,285 | -73,99 | *Engystomops pustulosus* | Foothills SNSM | negative | NA | NA | No |
| 24/8/2024 | 11,285 | -73,99 | *Engystomops pustulosus* | Foothills SNSM | negative | NA | NA | No |
| 24/8/2024 | 11,285 | -73,99 | *Engystomops pustulosus* | Foothills SNSM | negative | NA | NA | No |
| 24/8/2024 | 11,285 | -73,99 | *Engystomops pustulosus* | Foothills SNSM | negative | NA | NA | No |
| 24/8/2024 | 11,285 | -73,99 | *Engystomops pustulosus* | Foothills SNSM | negative | NA | NA | No |
| 24/8/2024 | 11,285 | -73,99 | *Boana sp.* | Foothills SNSM | negative | NA | NA | No |
| 24/8/2024 | 11,285 | -73,99 | *Engystomops pustulosus* | Foothills SNSM | negative | NA | NA | No |
| 24/8/2024 | 11,285 | -73,99 | *Engystomops pustulosus* | Foothills SNSM | negative | NA | NA | No |
| 24/8/2024 | 11,285 | -73,99 | *Rhinella humboldti* | Foothills SNSM | negative | NA | NA | No |
| 24/8/2024 | 11,285 | -73,99 | *Engystomops pustulosus* | Foothills SNSM | negative | NA | NA | No |
| 24/8/2024 | 11,285 | -73,99 | *Engystomops pustulosus* | Foothills SNSM | negative | NA | NA | No |
| 24/8/2024 | 11,282 | -73,971 | *Leptodactylus fuscus* | Foothills SNSM | NA | negative | NA | No |
| 24/8/2024 | 11,282 | -73,971 | *Engystomops pustulosus* | Foothills SNSM | negative | NA | NA | No |
| 24/8/2024 | 11,282 | -73,971 | *Engystomops pustulosus* | Foothills SNSM | negative | NA | NA | No |
| 24/8/2024 | 11,282 | -73,971 | *Engystomops pustulosus* | Foothills SNSM | negative | NA | NA | No |
| 24/8/2024 | 11,282 | -73,971 | *Engystomops pustulosus* | Foothills SNSM | negative | NA | NA | No |
| 24/8/2024 | 11,282 | -73,971 | *Engystomops pustulosus* | Foothills SNSM | negative | NA | NA | No |
| 24/8/2024 | 11,282 | -73,971 | *Engystomops pustulosus* | Foothills SNSM | negative | NA | NA | No |
| 24/8/2024 | 11,282 | -73,971 | *Engystomops pustulosus* | Foothills SNSM | negative | NA | NA | No |
| 24/8/2024 | 11,282 | -73,971 | *Engystomops pustulosus* | Foothills SNSM | negative | NA | NA | No |
| 24/8/2024 | 11,282 | -73,971 | *Engystomops pustulosus* | Foothills SNSM | negative | NA | NA | No |
| 24/8/2024 | 11,282 | -73,971 | *Engystomops pustulosus* | Foothills SNSM | negative | NA | NA | No |
| 29/8/2024 | 10,497 | -73,058 | *Pleurodema brachyops* | Serrania de Perrijá | negative | NA | NA | No |
| 29/8/2024 | 10,497 | -73,058 | *Engystomops pustulosus* | Serrania de Perrijá | negative | NA | NA | No |
| 29/8/2024 | 10,497 | -73,058 | *Engystomops pustulosus* | Serrania de Perrijá | negative | NA | NA | No |
| 29/8/2024 | 10,497 | -73,058 | *Engystomops pustulosus* | Serrania de Perrijá | negative | NA | NA | No |
| 29/8/2024 | 10,497 | -73,058 | *Rhinella humboldti* | Serrania de Perrijá | negative | NA | NA | No |
| 29/8/2024 | 10,497 | -73,058 | *Engystomops pustulosus* | Serrania de Perrijá | negative | NA | NA | No |
| 31/8/2024 | 10,891 | -72,836 | *Engystomops pustulosus* | Foothills SNSM | negative | NA | NA | No |
| 31/8/2024 | 10,891 | -72,836 | *Pleurodema brachyops* | Foothills SNSM | negative | NA | NA | No |
| 31/8/2024 | 10,891 | -72,836 | *Engystomops pustulosus* | Foothills SNSM | negative | NA | NA | No |
| 31/8/2024 | 10,891 | -72,836 | *Pleurodema brachyops* | Foothills SNSM | negative | NA | NA | No |

| | | | | | | | | |
|---|---|---|---|---|---|---|---|---|
| 31/8/2024 | 10,891 | -72,836 | *Pleurodema brachyops* | Foothills SNSM | negative | NA | NA | No |
| 31/8/2024 | 10,891 | -72,836 | *Engystomops pustulosus* | Foothills SNSM | negative | NA | NA | No |
| 31/8/2024 | 10,891 | -72,836 | *Engystomops pustulosus* | Foothills SNSM | negative | NA | NA | No |
| 31/8/2024 | 10,891 | -72,836 | *Engystomops pustulosus* | Foothills SNSM | negative | NA | NA | No |
| 31/8/2024 | 10,891 | -72,836 | *Engystomops pustulosus* | Foothills SNSM | negative | NA | NA | No |
| 31/8/2024 | 10,891 | -72,836 | *Engystomops pustulosus* | Foothills SNSM | negative | NA | NA | No |
| 31/8/2024 | 10,891 | -72,836 | *Pleurodema brachyops* | Foothills SNSM | negative | NA | NA | No |
| 31/8/2024 | 10,891 | -72,836 | *Engystomops pustulosus* | Foothills SNSM | negative | NA | NA | No |
| 31/8/2024 | 10,891 | -72,836 | *Engystomops pustulosus* | Foothills SNSM | negative | NA | NA | No |
| 31/8/2024 | 10,891 | -72,836 | *Engystomops pustulosus* | Foothills SNSM | negative | NA | NA | No |
| 31/8/2024 | 10,891 | -72,836 | *Engystomops pustulosus* | Foothills SNSM | negative | NA | NA | No |
| 31/8/2024 | 10,891 | -72,836 | *Pleurodema brachyops* | Foothills SNSM | negative | NA | NA | No |
| 31/8/2024 | 10,891 | -72,836 | *Engystomops pustulosus* | Foothills SNSM | negative | NA | NA | No |
| 31/8/2024 | 10,891 | -72,836 | *Engystomops pustulosus* | Foothills SNSM | negative | NA | NA | No |
| 31/8/2024 | 10,891 | -72,836 | *Engystomops pustulosus* | Foothills SNSM | negative | NA | NA | No |
| 31/8/2024 | 10,891 | -72,836 | *Engystomops pustulosus* | Foothills SNSM | negative | NA | NA | No |
| 31/8/2024 | 10,891 | -72,836 | *Engystomops pustulosus* | Foothills SNSM | negative | NA | NA | No |
| 31/8/2024 | 10,891 | -72,836 | *Engystomops pustulosus* | Foothills SNSM | negative | NA | NA | No |
| 31/8/2024 | 10,891 | -72,836 | *Engystomops pustulosus* | Foothills SNSM | negative | NA | NA | No |
| 31/8/2024 | 10,891 | -72,836 | *Engystomops pustulosus* | Foothills SNSM | negative | NA | NA | No |
| 31/8/2024 | 10,891 | -72,836 | *Engystomops pustulosus* | Foothills SNSM | negative | NA | NA | No |
| 31/8/2024 | 10,891 | -72,836 | *Pleurodema brachyops* | Foothills SNSM | negative | NA | NA | No |
| 31/8/2024 | 10,891 | -72,836 | *Engystomops pustulosus* | Foothills SNSM | negative | NA | NA | No |
| 31/8/2024 | 10,891 | -72,836 | *Leptodactylus fuscus* | Foothills SNSM | negative | NA | NA | No |
| 31/8/2024 | 10,891 | -72,836 | *Engystomops pustulosus* | Foothills SNSM | negative | NA | NA | No |
| 31/8/2024 | 10,891 | -72,836 | *Engystomops pustulosus* | Foothills SNSM | negative | NA | NA | No |
| 31/8/2024 | 10,891 | -72,836 | *Leptodactylus fuscus* | Foothills SNSM | negative | NA | NA | No |
| 31/8/2024 | 10,891 | -72,836 | *Leptodactylus fuscus* | Foothills SNSM | negative | NA | NA | No |
| 31/8/2024 | 10,781 | -72,993 | *Pleurodema brachyops* | Foothills SNSM | negative | NA | NA | No |
| 31/8/2024 | 10,781 | -72,993 | *Ceratophrys calcarata* | Foothills SNSM | negative | NA | NA | No |
| 31/8/2024 | 10,781 | -72,993 | *Pleurodema brachyops* | Foothills SNSM | negative | NA | NA | No |
| 31/8/2024 | 10,781 | -72,993 | *Rhinella humboldti* | Foothills SNSM | negative | NA | NA | No |
| 31/8/2024 | 10,781 | -72,993 | *Pleurodema brachyops* | Foothills SNSM | negative | NA | NA | No |
| 31/8/2024 | 10,781 | -72,993 | *Pleurodema brachyops* | Foothills SNSM | negative | NA | NA | No |
| 31/8/2024 | 10,781 | -72,993 | *Pleurodema brachyops* | Foothills SNSM | negative | NA | NA | No |
| 31/8/2024 | 10,781 | -72,993 | *Pleurodema brachyops* | Foothills SNSM | negative | NA | NA | No |
| 31/8/2024 | 10,781 | -72,993 | *Pleurodema brachyops* | Foothills SNSM | negative | NA | NA | No |

| | | | | | | | | |
|---|---|---|---|---|---|---|---|---|
| 1/9/2024 | 9,992 | -73,896 | *Elachistocleis panamensis* | Foothills SNSM | negative | NA | NA | No |
| 1/9/2024 | 9,992 | -73,896 | *Elachistocleis panamensis* | Foothills SNSM | negative | NA | NA | No |
| 1/9/2024 | 9,992 | -73,896 | *Elachistocleis panamensis* | Foothills SNSM | negative | NA | NA | No |
| 1/9/2024 | 9,992 | -73,896 | *Elachistocleis panamensis* | Foothills SNSM | negative | NA | NA | No |
| 1/9/2024 | 9,992 | -73,896 | *Elachistocleis panamensis* | Foothills SNSM | negative | NA | NA | No |
| 1/9/2024 | 9,992 | -73,896 | *Rhinella humboldti* | Foothills SNSM | negative | NA | NA | No |
| 1/9/2024 | 9,992 | -73,896 | *Pleurodema brachyops* | Foothills SNSM | negative | NA | NA | No |
| 1/9/2024 | 9,992 | -73,896 | *Elachistocleis panamensis* | Foothills SNSM | negative | NA | NA | No |
| 1/9/2024 | 9,992 | -73,896 | *Pleurodema brachyops* | Foothills SNSM | negative | NA | NA | No |
| 1/9/2024 | 9,992 | -73,896 | *Pleurodema brachyops* | Foothills SNSM | negative | NA | NA | No |
| 1/9/2024 | 9,992 | -73,896 | *Ceratophrys calcarata* | Foothills SNSM | negative | NA | NA | No |
| 1/9/2024 | 9,992 | -73,896 | *Engystomops pustulosus* | Foothills SNSM | negative | NA | NA | No |
| 1/9/2024 | 9,992 | -73,896 | *Engystomops pustulosus* | Foothills SNSM | negative | NA | NA | No |
| 1/9/2024 | 9,992 | -73,896 | *Elachistocleis panamensis* | Foothills SNSM | negative | NA | NA | No |
| 1/9/2024 | 9,992 | -73,896 | *Pleurodema brachyops* | Foothills SNSM | negative | NA | NA | No |
| 1/9/2024 | 9,992 | -73,896 | *Elachistocleis panamensis* | Foothills SNSM | negative | NA | NA | No |
| 1/9/2024 | 9,992 | -73,896 | *Elachistocleis panamensis* | Foothills SNSM | negative | NA | NA | No |
| 1/9/2024 | 9,992 | -73,896 | *Elachistocleis panamensis* | Foothills SNSM | negative | NA | NA | No |
| 1/9/2024 | 9,992 | -73,896 | *Leptodactylus fuscus* | Foothills SNSM | negative | NA | NA | No |
| 1/9/2024 | 9,992 | -73,896 | *Elachistocleis panamensis* | Foothills SNSM | negative | NA | NA | No |
| 1/9/2024 | 9,992 | -73,896 | *Elachistocleis panamensis* | Foothills SNSM | negative | NA | NA | No |
| 1/9/2024 | 9,992 | -73,896 | *Rhinella humboldti* | Foothills SNSM | negative | NA | NA | No |
| 1/9/2024 | 9,992 | -73,896 | *Elachistocleis panamensis* | Foothills SNSM | negative | NA | NA | No |

**Table S7.** Development of individual infection loads over the course of 10-week infection trials. As all negative controls remained negative, only experimentally infected specimens are shown. AL = *Atelopus laetissimus*, BS = *Bolitoglossa savagei*, CB = *Cryptobatrachus boulengeri*. Specimens AL9, AL11 and AL13 were inoculated at week 4 with a higher zoospore dose (see Methods: infection trials) to confirm tolerance in the species inconsistent with other harlequin toad species.

| Specimen | Sampling date | $C_T$ | $C_T$ Mean | $C_T$ SD | Quantity | Quantity Mean | Quantity SD | mortality |
|---|---|---|---|---|---|---|---|---|
| AL1 | Week 0 (before inoculation) | Undetermined | | | | | | 0 |
| AL1 | Week 0 (before inoculation) | Undetermined | | | | | | 0 |
| AL1 | Week 1 | 41,11335373 | 41,01559 | 0,138266 | 51,40075 | 54,97926712 | 5,060788631 | 0 |
| AL1 | Week 1 | 40,91781616 | 41,01559 | 0,138266 | 58,55779 | 54,97926712 | 5,060788631 | 0 |
| AL1 | Week 2 | Undetermined | | | | | | 0 |
| AL1 | Week 2 | Undetermined | | | | | | 0 |
| AL1 | Week 3 | Undetermined | 42,95041 | | | | | 0 |
| AL1 | Week 3 | 42,95040512 | 42,95041 | | 48,69255 | 48,69254684 | | 0 |
| AL1 | Week 4 | 40,89477921 | 40,29887 | 0,842735 | 95,95448 | 152,9239349 | 80,56697083 | 0 |
| AL1 | Week 4 | 39,70297241 | 40,29887 | 0,842735 | 209,8934 | 152,9239349 | 80,56697083 | 0 |
| AL1 | Week 5 | 36,08103943 | 35,68641 | 0,558092 | 1166,983 | 1597,95874 | 609,4919434 | 0 |
| AL1 | Week 5 | 35,29177856 | 35,68641 | 0,558092 | 2028,935 | 1597,95874 | 609,4919434 | 0 |
| AL1 | Week 6 | 35,71557236 | 35,75106 | 0,05019 | 1878,548 | 1835,135986 | 61,39416504 | 0 |
| AL1 | Week 6 | 35,78655243 | 35,75106 | 0,05019 | 1791,724 | 1835,135986 | 61,39416504 | 0 |
| AL1 | Week 7 | 37,2943306 | 37,16208 | 0,187035 | 1412,588 | 1546,165649 | 188,9074707 | 0 |

| | | | | | | | | |
|---|---|---|---|---|---|---|---|---|
| AL1 | Week 7 | 37,0298233 | 37,16208 | 0,187035 | 1679,743 | 1546,165649 | 188,9074707 | 0 |
| AL1 | Week 8 | 33,52910614 | 33,5538 | 0,034929 | 18045,88 | 17760,46484 | 403,6405945 | 0 |
| AL1 | Week 8 | 33,57850266 | 33,5538 | 0,034929 | 17475,05 | 17760,46484 | 403,6405945 | 0 |
| AL1 | Week 9 | 31,34446907 | 31,41468 | 0,099293 | 74777,13 | 71512,1875 | 4617,335449 | 0 |
| AL1 | Week 9 | 31,48488998 | 31,41468 | 0,099293 | 68247,23 | 71512,1875 | 4617,335449 | 0 |
| AL1 | Week 10 | 29,81492996 | 29,76653 | 0,068453 | 202313,6 | 208891 | 9301,791016 | 0 |
| AL1 | Week 10 | 29,71812248 | 29,76653 | 0,068453 | 215468,4 | 208891 | 9301,791016 | 0 |
| AL2 | Week 0 (before inoculation) | Undetermined | | | | | | 0 |
| AL2 | Week 0 (before inoculation) | Undetermined | | | | | | 0 |
| AL2 | Week 1 | 37,43198013 | 37,19976 | 0,328403 | 452,8135 | 539,904541 | 123,1653442 | 0 |
| AL2 | Week 1 | 36,96754837 | 37,19976 | 0,328403 | 626,9956 | 539,904541 | 123,1653442 | 0 |
| AL2 | Week 2 | Undetermined | | | | | | 0 |
| AL2 | Week 2 | Undetermined | | | | | | 0 |
| AL2 | Week 3 | Undetermined | | | | | | 0 |
| AL2 | Week 3 | Undetermined | | | | | | 0 |
| AL2 | Week 4 | 43,29425049 | 42,0053 | 1,822853 | 39,16265 | 119,8146515 | 114,0591431 | 0 |
| AL2 | Week 4 | 40,71634674 | 42,0053 | 1,822853 | 200,4666 | 119,8146515 | 114,0591431 | 0 |
| AL2 | Week 5 | Undetermined | 40,553 | | | | | 0 |
| AL2 | Week 5 | 40,55299759 | 40,553 | | 74,68147 | 74,68146515 | | 0 |

| | | | | | | | | |
|---|---|---|---|---|---|---|---|---|
| AL2 | Week 6 | Undetermined | | | | | | 0 |
| AL2 | Week 6 | Undetermined | | | | | | 0 |
| AL2 | Week 7 | 40,41300964 | 39,74088 | 0,950542 | 183,2502 | 312,594696 | 182,9207764 | 0 |
| AL2 | Week 7 | 39,06874084 | 39,74088 | 0,950542 | 441,9392 | 312,594696 | 182,9207764 | 0 |
| AL2 | Week 8 | 41,60984802 | 41,76759 | 0,223088 | 113,8266 | 103,5172577 | 14,57964325 | 0 |
| AL2 | Week 8 | 41,92534256 | 41,76759 | 0,223088 | 93,20789 | 103,5172577 | 14,57964325 | 0 |
| AL2 | Week 9 | 37,72635269 | 37,6494 | 0,108827 | 5019,864 | 5253,310547 | 330,1429138 | 0 |
| AL2 | Week 9 | 37,57244873 | 37,6494 | 0,108827 | 5486,757 | 5253,310547 | 330,1429138 | 0 |
| AL2 | Week 10 | 36,63441086 | 36,56449 | 0,098879 | 2392,222 | 2506,166992 | 161,1421356 | 0 |
| AL2 | Week 10 | 36,4945755 | 36,56449 | 0,098879 | 2620,112 | 2506,166992 | 161,1421356 | 0 |
| AL3 | Week 0 (before inoculation) | Undetermined | | | | | | 0 |
| AL3 | Week 0 (before inoculation) | Undetermined | | | | | | 0 |
| AL3 | Week 1 | 41,31404877 | 40,666 | 0,916476 | 44,96359 | 75,82717133 | 43,64769363 | 0 |
| AL3 | Week 1 | 40,01795578 | 40,666 | 0,916476 | 106,6908 | 75,82717133 | 43,64769363 | 0 |
| AL3 | Week 2 | 41,80137253 | 41,05312 | 1,058192 | 55,65271 | 101,2831726 | 64,53121948 | 0 |
| AL3 | Week 2 | 40,30486298 | 41,05312 | 1,058192 | 146,9136 | 101,2831726 | 64,53121948 | 0 |
| AL3 | Week 3 | Undetermined | | | | | | 0 |
| AL3 | Week 3 | Undetermined | | | | | | 0 |
| AL3 | Week 4 | Undetermined | | | | | | 0 |

| | | | | | | | | |
|---|---|---|---|---|---|---|---|---|
| AL3 | Week 4 | Undetermined | | | | | | 0 |
| AL3 | Week 5 | Undetermined | | | | | | 0 |
| AL3 | Week 5 | Undetermined | | | | | | 0 |
| AL3 | Week 6 | 39,46411133 | 39,39589 | 0,096478 | 264,0307 | 276,4737244 | 17,59704971 | 0 |
| AL3 | Week 6 | 39,32767105 | 39,39589 | 0,096478 | 288,9167 | 276,4737244 | 17,59704971 | 0 |
| AL3 | Week 7 | Undetermined | | | | | | 0 |
| AL3 | Week 7 | Undetermined | | | | | | 0 |
| AL3 | Week 8 | Undetermined | | | | | | 0 |
| AL3 | Week 8 | Undetermined | | | | | | 0 |
| AL3 | Week 9 | Undetermined | | | | | | 0 |
| AL3 | Week 9 | Undetermined | | | | | | 0 |
| AL3 | Week 10 | Undetermined | | | | | | 0 |
| AL3 | Week 10 | Undetermined | | | | | | 0 |
| AL4 | Week 0 (before inoculation) | Undetermined | | | | | | 0 |
| AL4 | Week 0 (before inoculation) | Undetermined | | | | | | 0 |
| AL4 | Week 1 | 38,97758865 | 38,47611 | 0,7092 | 153,2954 | 231,4365997 | 110,5083694 | 0 |
| AL4 | Week 1 | 37,97462845 | 38,47611 | 0,7092 | 309,5778 | 231,4365997 | 110,5083694 | 0 |
| AL4 | Week 2 | Undetermined | | | | | | 0 |
| AL4 | Week 2 | Undetermined | | | | | | 0 |

| | | | | | | | | |
|---|---|---|---|---|---|---|---|---|
| AL4 | Week 3 | Undetermined | | | | | | 0 |
| AL4 | Week 3 | Undetermined | | | | | | 0 |
| AL4 | Week 4 | Undetermined | | | | | | 0 |
| AL4 | Week 4 | Undetermined | | | | | | 0 |
| AL4 | Week 5 | 40,46007156 | 40,46007 | | 235,7992 | 235,7991791 | | 0 |
| AL4 | Week 5 | Undetermined | 40,46007 | | | | | 0 |
| AL4 | Week 6 | Undetermined | | | | | | 0 |
| AL4 | Week 6 | Undetermined | | | | | | 0 |
| AL4 | Week 7 | Undetermined | 41,32918 | | | | | 0 |
| AL4 | Week 7 | 41,32917786 | 41,32918 | | 100,5728 | 100,5728226 | | 0 |
| AL4 | Week 8 | Undetermined | | | | | | 0 |
| AL4 | Week 8 | Undetermined | | | | | | 0 |
| AL4 | Week 9 | Undetermined | | | | | | 0 |
| AL4 | Week 9 | Undetermined | | | | | | 0 |
| AL4 | Week 10 | Undetermined | | | | | | 0 |
| AL4 | Week 10 | Undetermined | | | | | | 0 |
| AL5 | Week 0 (before inoculation) | Undetermined | | | | | | 0 |
| AL5 | Week 0 (before inoculation) | Undetermined | | | | | | 0 |
| AL5 | Week 1 | Undetermined | | | | | | 0 |

| | | | | | | | | |
|---|---|---|---|---|---|---|---|---|
| AL5 | Week 1 | Undetermined | | | | | | 0 |
| AL5 | Week 2 | Undetermined | | | | | | 0 |
| AL5 | Week 2 | Undetermined | | | | | | 0 |
| AL5 | Week 3 | 38,72916794 | 38,75406 | 0,035196 | 705,84 | 694,8864746 | 15,49053288 | 0 |
| AL5 | Week 3 | 38,77894211 | 38,75406 | 0,035196 | 683,933 | 694,8864746 | 15,49053288 | 0 |
| AL5 | Week 4 | 42,70179749 | 43,4267 | 1,025162 | 31,03353 | 21,57553482 | 13,37563038 | 0 |
| AL5 | Week 4 | 44,15159607 | 43,4267 | 1,025162 | 12,11753 | 21,57553482 | 13,37563038 | 0 |
| AL5 | Week 5 | 35,82006836 | 35,93104 | 0,156935 | 4456,426 | 4164,199219 | 413,2712708 | 0 |
| AL5 | Week 5 | 36,04200745 | 35,93104 | 0,156935 | 3871,972 | 4164,199219 | 413,2712708 | 0 |
| AL5 | Week 6 | 36,18312836 | 36,06011 | 0,173974 | 2303,193 | 2506,287354 | 287,2193298 | 0 |
| AL5 | Week 6 | 35,93709183 | 36,06011 | 0,173974 | 2709,382 | 2506,287354 | 287,2193298 | 0 |
| AL5 | Week 7 | 33,05137253 | 33,09944 | 0,067983 | 18206,3 | 17646,48438 | 791,701355 | 0 |
| AL5 | Week 7 | 33,14751434 | 33,09944 | 0,067983 | 17086,67 | 17646,48438 | 791,701355 | 0 |
| AL5 | Week 8 | 31,31755638 | 31,23734 | 0,113439 | 76098,22 | 80285,04688 | 5921,063477 | 0 |
| AL5 | Week 8 | 31,15712929 | 31,23734 | 0,113439 | 84471,87 | 80285,04688 | 5921,063477 | 0 |
| AL5 | Week 9 | 31,19414139 | 31,20028 | 0,008684 | 82461,7 | 82133,5 | 464,1382751 | 0 |
| AL5 | Week 9 | 31,20642281 | 31,20028 | 0,008684 | 81805,3 | 82133,5 | 464,1382751 | 0 |
| AL5 | Week 10 | 32,81641388 | 32,82065 | 0,005994 | 28693,86 | 28614,94141 | 111,6041031 | 0 |
| AL5 | Week 10 | 32,82489014 | 32,82065 | 0,005994 | 28536,03 | 28614,94141 | 111,6041031 | 0 |

| | | | | | | | | |
|---|---|---|---|---|---|---|---|---|
| AL6 | Week 0 (before inoculation) | Undetermined | | | | | | 0 |
| AL6 | Week 0 (before inoculation) | Undetermined | | | | | | 0 |
| AL6 | Week 1 | Undetermined | 42,19857 | | | | | 0 |
| AL6 | Week 1 | 42,19856644 | 42,19857 | | 24,93223 | 24,93223 | | 0 |
| AL6 | Week 2 | Undetermined | | | | | | 0 |
| AL6 | Week 2 | Undetermined | | | | | | 0 |
| AL6 | Week 3 | 42,44096375 | 43,17588 | 1,039326 | 37,4823 | 25,80940247 | 16,50797653 | 0 |
| AL6 | Week 3 | 43,9107933 | 43,17588 | 1,039326 | 14,1365 | 25,80940247 | 16,50797653 | 0 |
| AL6 | Week 4 | 38,91572952 | 39,38847 | 0,66855 | 361,734 | 278,8191528 | 117,2593002 | 0 |
| AL6 | Week 4 | 39,86120224 | 39,38847 | 0,66855 | 195,9043 | 278,8191528 | 117,2593002 | 0 |
| AL6 | Week 5 | 37,89965439 | 37,54394 | 0,503054 | 326,2777 | 431,7182617 | 149,1154327 | 0 |
| AL6 | Week 5 | 37,18822861 | 37,54394 | 0,503054 | 537,1588 | 431,7182617 | 149,1154327 | 0 |
| AL6 | Week 6 | Undetermined | | | | | | 0 |
| AL6 | Week 6 | Undetermined | | | | | | 0 |
| AL6 | Week 7 | 38,73247528 | 38,9138 | 0,256431 | 427,9751 | 382,4159851 | 64,43035889 | 0 |
| AL6 | Week 7 | 39,09512329 | 38,9138 | 0,256431 | 336,8568 | 382,4159851 | 64,43035889 | 0 |
| AL6 | Week 8 | 38,13286972 | 38,08927 | 0,061654 | 1029,782 | 1059,019531 | 41,34831238 | 0 |
| AL6 | Week 8 | 38,04567719 | 38,08927 | 0,061654 | 1088,257 | 1059,019531 | 41,34831238 | 0 |
| AL6 | Week 9 | 34,62730789 | 34,64572 | 0,026038 | 8831,208 | 8726,660156 | 147,8536835 | 0 |

| | | | | | | | | |
|---|---|---|---|---|---|---|---|---|
| AL6 | Week 9 | 34,66413116 | 34,64572 | 0,026038 | 8622,111 | 8726,660156 | 147,8536835 | 0 |
| AL6 | Week 10 | 31,70843697 | 31,76639 | 0,081963 | 59007,8 | 56864,25 | 3031,435791 | 0 |
| AL6 | Week 10 | 31,82435036 | 31,76639 | 0,081963 | 54720,7 | 56864,25 | 3031,435791 | 0 |
| AL9 | Week 0 | Undetermined | | | | | | 0 |
| AL9 | Week 0 | Undetermined | | | | | | 0 |
| AL9 | Week 1 | Undetermined | | | | | | 0 |
| AL9 | Week 1 | Undetermined | | | | | | 0 |
| AL9 | Week 2 | Undetermined | | | | | | 0 |
| AL9 | Week 2 | Undetermined | | | | | | 0 |
| AL9 | Week 3 | Undetermined | | | | | | 0 |
| AL9 | Week 3 | Undetermined | | | | | | 0 |
| AL9 | Week 4 (before inoculation) | Undetermined | | | | | | 0 |
| AL9 | Week 4 (before inoculation) | Undetermined | | | | | | 0 |
| AL9 | Week 5 | 37,85824585 | 38,25802 | 0,565369 | 976,4153 | 777,4133911 | 281,4311829 | 0 |
| AL9 | Week 5 | 38,65779877 | 38,25802 | 0,565369 | 578,4115 | 777,4133911 | 281,4311829 | 0 |
| AL9 | Week 6 | Undetermined | | | | | | 0 |
| AL9 | Week 6 | Undetermined | | | | | | 0 |
| AL9 | Week 7 | 37,88520813 | 37,71473 | 0,241088 | 748,7463 | 843,2479248 | 133,6455536 | 0 |
| AL9 | Week 7 | 37,54425812 | 37,71473 | 0,241088 | 937,7496 | 843,2479248 | 133,6455536 | 0 |

| | | | | | | | | |
|---|---|---|---|---|---|---|---|---|
| AL9 | Week 8 | 40,9387207 | 41,36159 | 0,598024 | 145,3381 | 114,5812073 | 43,49674988 | 0 |
| AL9 | Week 8 | 41,78445435 | 41,36159 | 0,598024 | 83,82436 | 114,5812073 | 43,49674988 | 0 |
| AL9 | Week 9 | 37,92729187 | 38,28957 | 0,512344 | 1031,397 | 837,5315552 | 274,1677856 | 0 |
| AL9 | Week 9 | 38,65185547 | 38,28957 | 0,512344 | 643,6656 | 837,5315552 | 274,1677856 | 0 |
| AL9 | Week 10 | 34,90211105 | 34,87535 | 0,037847 | 7385,147 | 7516,023438 | 185,0865784 | 0 |
| AL9 | Week 10 | 34,84858704 | 34,87535 | 0,037847 | 7646,899 | 7516,023438 | 185,0865784 | 0 |
| AL11 | Week 0 | Undetermined | | | | | | 0 |
| AL11 | Week 0 | Undetermined | | | | | | 0 |
| AL11 | Week 1 | Undetermined | | | | | | 0 |
| AL11 | Week 1 | Undetermined | | | | | | 0 |
| AL11 | Week 2 | Undetermined | | | | | | 0 |
| AL11 | Week 2 | Undetermined | | | | | | 0 |
| AL11 | Week 3 | Undetermined | | | | | | 0 |
| AL11 | Week 3 | Undetermined | | | | | | 0 |
| AL11 | Week 4 (before inoculation) | Undetermined | | | | | | 0 |
| AL11 | Week 4 (before inoculation) | Undetermined | | | | | | 0 |
| AL11 | Week 5 | 39,77325058 | 39,87208 | 0,139768 | 364,3206 | 342,8831787 | 30,3170948 | 0 |
| AL11 | Week 5 | 39,97091293 | 39,87208 | 0,139768 | 321,4457 | 342,8831787 | 30,3170948 | 0 |
| AL11 | Week 6 | 37,18339539 | 37,26642 | 0,117412 | 1519,029 | 1440,772583 | 110,6715317 | 0 |

| | | | | | | | | |
|---|---|---|---|---|---|---|---|---|
| AL11 | Week 6 | 37,34944153 | 37,26642 | 0,117412 | 1362,516 | 1440,772583 | 110,6715317 | 0 |
| AL11 | Week 7 | 35,83475113 | 35,5007 | 0,472422 | 3673,911 | 4682,150391 | 1425,865967 | 0 |
| AL11 | Week 7 | 35,16664505 | 35,5007 | 0,472422 | 5690,39 | 4682,150391 | 1425,865967 | 0 |
| AL11 | Week 8 | 32,24165726 | 32,29975 | 0,082152 | 42992,42 | 41467,26953 | 2156,885498 | 0 |
| AL11 | Week 8 | 32,35783768 | 32,29975 | 0,082152 | 39942,12 | 41467,26953 | 2156,885498 | 0 |
| AL11 | Week 9 | 31,68419456 | 31,72333 | 0,055343 | 59946,04 | 58457,74219 | 2104,76416 | 0 |
| AL11 | Week 9 | 31,76246071 | 31,72333 | 0,055343 | 56969,45 | 58457,74219 | 2104,76416 | 0 |
| AL11 | Week 10 | 30,13900757 | 30,12646 | 0,017743 | 163847,2 | 165195,875 | 1907,332153 | 0 |
| AL11 | Week 10 | 30,11391449 | 30,12646 | 0,017743 | 166544,6 | 165195,875 | 1907,332153 | 0 |
| AL13 | Week 0 | Undetermined | | | | | | 0 |
| AL13 | Week 0 | Undetermined | | | | | | 0 |
| AL13 | Week 1 | Undetermined | | | | | | 0 |
| AL13 | Week 1 | Undetermined | | | | | | 0 |
| AL13 | Week 2 | Undetermined | | | | | | 0 |
| AL13 | Week 2 | Undetermined | | | | | | 0 |
| AL13 | Week 3 | Undetermined | | | | | | 0 |
| AL13 | Week 3 | Undetermined | | | | | | 0 |
| AL13 | Week 4 (before inoculation) | Undetermined | | | | | | 0 |
| AL13 | Week 4 (before inoculation) | Undetermined | | | | | | 0 |

| | | | | | | | | |
|---|---|---|---|---|---|---|---|---|
| AL13 | Week 5 | 39,21538162 | 39,26447 | 0,069412 | 518,7415 | 503,1050415 | 22,11331367 | 0 |
| AL13 | Week 5 | 39,31354523 | 39,26447 | 0,069412 | 487,4686 | 503,1050415 | 22,11331367 | 0 |
| AL13 | Week 6 | 34,32978058 | 34,36475 | 0,049457 | 11454,03 | 11205,84082 | 350,9928894 | 0 |
| AL13 | Week 6 | 34,39972305 | 34,36475 | 0,049457 | 10957,65 | 11205,84082 | 350,9928894 | 0 |
| AL13 | Week 7 | Undetermined | 42,81217 | | | | | 0 |
| AL13 | Week 7 | 42,81217194 | 42,81217 | | 38,08099 | 38,08098602 | | 0 |
| AL13 | Week 8 | 33,22111893 | 33,19158 | 0,041772 | 6708,879 | 6843,809082 | 190,819397 | 0 |
| AL13 | Week 8 | 33,16204453 | 33,19158 | 0,041772 | 6978,739 | 6843,809082 | 190,819397 | 0 |
| AL13 | Week 9 | 33,94264221 | 33,90993 | 0,046252 | 13788,29 | 14088,07031 | 423,9505615 | 0 |
| AL13 | Week 9 | 33,8772316 | 33,90993 | 0,046252 | 14387,85 | 14088,07031 | 423,9505615 | 0 |
| AL13 | Week 10 | 33,67316437 | 33,66322 | 0,014062 | 16431,09 | 16538,09375 | 151,3236084 | 0 |
| AL13 | Week 10 | 33,65327835 | 33,66322 | 0,014062 | 16645,1 | 16538,09375 | 151,3236084 | 0 |
| BS1 | Week 0 (before inoculation) | Undetermined | | | | | | 0 |
| BS1 | Week 0 (before inoculation) | Undetermined | | | | | | 0 |
| BS1 | Week 1 | 33,602005 | 33,52167 | 0,113617 | 12657,63 | 13365,85352 | 1001,578796 | 0 |
| BS1 | Week 1 | 33,44132614 | 33,52167 | 0,113617 | 14074,08 | 13365,85352 | 1001,578796 | 0 |
| BS1 | Week 2 | 34,99033356 | 35,08135 | 0,128723 | 4615,269 | 4358,25293 | 363,4749756 | 0 |
| BS1 | Week 2 | 35,17237473 | 35,08135 | 0,128723 | 4101,237 | 4358,25293 | 363,4749756 | 0 |
| BS1 | Week 3 | 34,38154602 | 34,41134 | 0,042136 | 6915,193 | 6782,491699 | 187,6684875 | 0 |

| | | | | | | | | |
|---|---|---|---|---|---|---|---|---|
| BS1 | Week 3 | 34,44113541 | 34,41134 | 0,042136 | 6649,79 | 6782,491699 | 187,6684875 | 0 |
| BS1 | Week 4 | 32,04525375 | 32,01878 | 0,037432 | 31177,67 | 31722,24609 | 770,1483765 | 0 |
| BS1 | Week 4 | 31,9923172 | 32,01878 | 0,037432 | 32266,82 | 31722,24609 | 770,1483765 | 0 |
| BS1 | Week 5 | 36,13664627 | 35,95962 | 0,250356 | 3646,679 | 4105,084473 | 648,2834473 | 0 |
| BS1 | Week 5 | 35,78258896 | 35,95962 | 0,250356 | 4563,49 | 4105,084473 | 648,2834473 | 0 |
| BS1 | Week 6 | 30,27656937 | 30,27308 | 0,004928 | 149272,2 | 149602,4063 | 466,9998474 | 0 |
| BS1 | Week 6 | 30,26959991 | 30,27308 | 0,004928 | 149932,6 | 149602,4063 | 466,9998474 | 0 |
| BS1 | Week 7 | 32,96336365 | 32,93507 | 0,040008 | 24086,21 | 24540,80664 | 642,8926392 | 0 |
| BS1 | Week 7 | 32,90678406 | 32,93507 | 0,040008 | 24995,4 | 24540,80664 | 642,8926392 | 0 |
| BS1 | Week 8 | 29,08649254 | 29,03708 | 0,069876 | 317222,3 | 327468 | 14489,56836 | 0 |
| BS1 | Week 8 | 28,98767281 | 29,03708 | 0,069876 | 337713,7 | 327468 | 14489,56836 | 0 |
| BS1 | Week 9 | 26,38513756 | 26,47687 | 0,129734 | 1884967 | 1778902,125 | 149998,5625 | 0 |
| BS1 | Week 9 | 26,56860924 | 26,47687 | 0,129734 | 1672837 | 1778902,125 | 149998,5625 | 0 |
| BS1 | Week 10 | 21,97931099 | 21,93418 | 0,063829 | 33144780 | 34147384 | 1417896,125 | 0 |
| BS1 | Week 10 | 21,88904381 | 21,93418 | 0,063829 | 35149988 | 34147384 | 1417896,125 | 0 |
| BS2 | Week 0 (before inoculation) | Undetermined | | | | | | 0 |
| BS2 | Week 0 (before inoculation) | Undetermined | | | | | | 0 |
| BS2 | Week 1 | 32,05429077 | 32,03473 | 0,027662 | 19613,94 | 19886,5 | 385,4657288 | 0 |
| BS2 | Week 1 | 32,01517105 | 32,03473 | 0,027662 | 20159,07 | 19886,5 | 385,4657288 | 0 |

| | | | | | | | | |
|---|---|---|---|---|---|---|---|---|
| BS2 | Week 2 | 33,1926384 | 33,26473 | 0,101959 | 20728,19 | 19794,32813 | 1320,680786 | 0 |
| BS2 | Week 2 | 33,33683014 | 33,26473 | 0,101959 | 18860,47 | 19794,32813 | 1320,680786 | 0 |
| BS2 | Week 3 | 29,24058151 | 29,30711 | 0,094087 | 192281,1 | 184331,375 | 11242,66602 | 0 |
| BS2 | Week 3 | 29,37364006 | 29,30711 | 0,094087 | 176381,6 | 184331,375 | 11242,66602 | 0 |
| BS2 | Week 4 | 32,71419144 | 32,6806 | 0,047515 | 23783,98 | 24326,10938 | 766,6818848 | 0 |
| BS2 | Week 4 | 32,64699554 | 32,6806 | 0,047515 | 24868,23 | 24326,10938 | 766,6818848 | 0 |
| BS2 | Week 5 | 28,71321487 | 28,73241 | 0,02714 | 203886 | 201180,625 | 3825,978027 | 0 |
| BS2 | Week 5 | 28,75159645 | 28,73241 | 0,02714 | 198475,3 | 201180,625 | 3825,978027 | 0 |
| BS2 | Week 6 | 29,26571274 | 29,22648 | 0,055477 | 138433,2 | 142345,2969 | 5532,536133 | 0 |
| BS2 | Week 6 | 29,18725586 | 29,22648 | 0,055477 | 146257,4 | 142345,2969 | 5532,536133 | 0 |
| BS2 | Week 7 | 29,32312775 | 29,37495 | 0,073284 | 213368,3 | 206313,1875 | 9977,464844 | 0 |
| BS2 | Week 7 | 29,42676735 | 29,37495 | 0,073284 | 199258 | 206313,1875 | 9977,464844 | 0 |
| BS2 | Week 8 | 24,74554062 | 24,82484 | 0,112154 | 4960672 | 4723586,5 | 335289,5313 | 0 |
| BS2 | Week 8 | 24,90415001 | 24,82484 | 0,112154 | 4486501 | 4723586,5 | 335289,5313 | 0 |
| BS2 | Week 9 | 25,89114189 | 25,89892 | 0,011001 | 2599619 | 2586526,25 | 18516,29883 | 0 |
| BS2 | Week 9 | 25,90670013 | 25,89892 | 0,011001 | 2573433 | 2586526,25 | 18516,29883 | 0 |
| BS2 | Week 10 | 23,04471588 | 23,07006 | 0,035846 | 16570123 | 16301278 | 380203,5313 | 0 |
| BS2 | Week 10 | 23,09540939 | 23,07006 | 0,035846 | 16032434 | 16301278 | 380203,5313 | 0 |
| BS3 | Week 0 (before inoculation) | Undetermined | | | | | | 0 |

| | | | | | | | | |
|---|---|---|---|---|---|---|---|---|
| BS3 | Week 0 (before inoculation) | Undetermined | | | | | | 0 |
| BS3 | Week 1 | 30,00853729 | 29,96247 | 0,065156 | 82256,29 | 84999,61719 | 3879,651855 | 0 |
| BS3 | Week 1 | 29,91639328 | 29,96247 | 0,065156 | 87742,95 | 84999,61719 | 3879,651855 | 0 |
| BS3 | Week 2 | 32,31420517 | 32,21917 | 0,134398 | 26186,56 | 27904,52344 | 2429,569092 | 0 |
| BS3 | Week 2 | 32,12413788 | 32,21917 | 0,134398 | 29622,49 | 27904,52344 | 2429,569092 | 0 |
| BS3 | Week 3 | 34,00332642 | 34,16769 | 0,232446 | 8754,731 | 7914,147461 | 1188,765259 | 0 |
| BS3 | Week 3 | 34,33205414 | 34,16769 | 0,232446 | 7073,563 | 7914,147461 | 1188,765259 | 0 |
| BS3 | Week 4 | 32,3871994 | 32,47721 | 0,127293 | 24975,58 | 23599,29688 | 1946,354248 | 0 |
| BS3 | Week 4 | 32,56721878 | 32,47721 | 0,127293 | 22223,02 | 23599,29688 | 1946,354248 | 0 |
| BS3 | Week 5 | 30,19724655 | 30,14866 | 0,068708 | 74400,73 | 76890,34375 | 3520,844971 | 0 |
| BS3 | Week 5 | 30,10007858 | 30,14866 | 0,068708 | 79379,96 | 76890,34375 | 3520,844971 | 0 |
| BS3 | Week 6 | 33,91015244 | 33,89909 | 0,015637 | 12956,75 | 13051,25098 | 133,6431885 | 0 |
| BS3 | Week 6 | 33,88803864 | 33,89909 | 0,015637 | 13145,75 | 13051,25098 | 133,6431885 | 0 |
| BS3 | Week 7 | 29,85099602 | 29,89166 | 0,057509 | 150586,9 | 146650,9531 | 5566,27832 | 0 |
| BS3 | Week 7 | 29,93232536 | 29,89166 | 0,057509 | 142715 | 146650,9531 | 5566,27832 | 0 |
| BS3 | Week 8 | 28,53771782 | 28,85857 | 0,453752 | 449085,8 | 374087,125 | 106064,0703 | 0 |
| BS3 | Week 8 | 29,17942047 | 28,85857 | 0,453752 | 299088,5 | 374087,125 | 106064,0703 | 0 |
| BS3 | Week 9 | 24,0875721 | 24,15298 | 0,092505 | 8406374 | 8063370 | 485080,9063 | 0 |
| BS3 | Week 9 | 24,21839333 | 24,15298 | 0,092505 | 7720366 | 8063370 | 485080,9063 | 0 |

| | | | | | | | | |
|---|---|---|---|---|---|---|---|---|
| BS3 | Week 10 | 25,97806931 | 25,96886 | 0,01303 | 2456652 | 2471468,75 | 20954,75586 | 0 |
| BS3 | Week 10 | 25,95964241 | 25,96886 | 0,01303 | 2486286 | 2471468,75 | 20954,75586 | 0 |
| BS4 | Week 0 (before inoculation) | Undetermined | | | | | | 0 |
| BS4 | Week 0 (before inoculation) | Undetermined | | | | | | 0 |
| BS4 | Week 1 | 30,81869125 | 30,82807 | 0,013259 | 46623,72 | 46319,40625 | 430,3711548 | 0 |
| BS4 | Week 1 | 30,8374424 | 30,82807 | 0,013259 | 46015,09 | 46319,40625 | 430,3711548 | 0 |
| BS4 | Week 2 | 30,7380619 | 30,7303 | 0,010973 | 75686,48 | 76074,13281 | 548,2286987 | 0 |
| BS4 | Week 2 | 30,72254372 | 30,7303 | 0,010973 | 76461,79 | 76074,13281 | 548,2286987 | 0 |
| BS4 | Week 3 | 30,77762794 | 30,85212 | 0,105343 | 73745,08 | 70308,28125 | 4860,364746 | 0 |
| BS4 | Week 3 | 30,92660522 | 30,85212 | 0,105343 | 66871,48 | 70308,28125 | 4860,364746 | 0 |
| BS4 | Week 4 | 27,43879509 | 27,42044 | 0,025964 | 618743,7 | 626200,5625 | 10545,61328 | 0 |
| BS4 | Week 4 | 27,40207672 | 27,42044 | 0,025964 | 633657,4 | 626200,5625 | 10545,61328 | 0 |
| BS4 | Week 5 | 27,12504959 | 27,19448 | 0,098197 | 1102098 | 1054195,125 | 67744,89844 | 0 |
| BS4 | Week 5 | 27,26392174 | 27,19448 | 0,098197 | 1006292 | 1054195,125 | 67744,89844 | 0 |
| BS4 | Week 6 | 27,7618084 | 27,63514 | 0,179132 | 377332,9 | 412045,75 | 49091,39844 | 0 |
| BS4 | Week 6 | 27,50847816 | 27,63514 | 0,179132 | 446758,6 | 412045,75 | 49091,39844 | 0 |
| BS4 | Week 7 | 26,41394997 | 26,35983 | 0,076543 | 1755744 | 1820233,5 | 91201,83594 | 0 |
| BS4 | Week 7 | 26,30570221 | 26,35983 | 0,076543 | 1884723 | 1820233,5 | 91201,83594 | 0 |
| BS4 | Week 8 | 25,62593842 | 25,61875 | 0,010168 | 1068224 | 1073375,5 | 7285,763184 | 0 |

| | | | | | | | | |
|---|---|---|---|---|---|---|---|---|
| BS4 | Week 8 | 25,61155891 | 25,61875 | 0,010168 | 1078527 | 1073375,5 | 7285,763184 | 0 |
| BS4 | Week 9 | 19,10505104 | 19,14663 | 0,058794 | 2,15E+08 | 209462912 | 8011791,5 | 0 |
| BS4 | Week 9 | 19,18819809 | 19,14663 | 0,058794 | 2,04E+08 | 209462912 | 8011791,5 | 0 |
| BS4 | 18/08/2025 | 22,04802132 | 22,0266 | 0,030293 | 31695472 | 32143484 | 633586,0625 | 1 |
| BS4 | 18/08/2025 | 22,00518036 | 22,0266 | 0,030293 | 32591498 | 32143484 | 633586,0625 | 1 |
| BS5 | Week 0 (before inoculation) | Undetermined | | | | | | 0 |
| BS5 | Week 0 (before inoculation) | Undetermined | | | | | | 0 |
| BS5 | Week 1 | 33,63772202 | 33,59853 | 0,055423 | 6466,413 | 6648,969727 | 258,174408 | 0 |
| BS5 | Week 1 | 33,55934143 | 33,59853 | 0,055423 | 6831,527 | 6648,969727 | 258,174408 | 0 |
| BS5 | Week 2 | 37,17825317 | 36,73495 | 0,626929 | 1101,825 | 1537,116577 | 615,5953369 | 0 |
| BS5 | Week 2 | 36,29164124 | 36,73495 | 0,626929 | 1972,408 | 1537,116577 | 615,5953369 | 0 |
| BS5 | Week 3 | 31,1672287 | 31,20067 | 0,047299 | 55104,7 | 53934,79688 | 1654,489014 | 0 |
| BS5 | Week 3 | 31,23411942 | 31,20067 | 0,047299 | 52764,89 | 53934,79688 | 1654,489014 | 0 |
| BS5 | Week 4 | 29,30998039 | 29,29003 | 0,028213 | 227561,7 | 230613,6875 | 4316,20166 | 0 |
| BS5 | Week 4 | 29,27008057 | 29,29003 | 0,028213 | 233665,7 | 230613,6875 | 4316,20166 | 0 |
| BS5 | Week 5 | 31,96535492 | 31,92632 | 0,055202 | 22890,01 | 23501,45117 | 864,7142334 | 0 |
| BS5 | Week 5 | 31,88728714 | 31,92632 | 0,055202 | 24112,9 | 23501,45117 | 864,7142334 | 0 |
| BS5 | Week 6 | 29,48901749 | 29,50663 | 0,024905 | 119298,5 | 117914,2188 | 1957,730103 | 0 |
| BS5 | Week 6 | 29,52423859 | 29,50663 | 0,024905 | 116529,9 | 117914,2188 | 1957,730103 | 0 |

| | | | | | | | | |
|---|---|---|---|---|---|---|---|---|
| BS5 | Week 7 | 30,30894279 | 30,32014 | 0,015828 | 136994,1 | 135997,3125 | 1409,650513 | 0 |
| BS5 | Week 7 | 30,33132744 | 30,32014 | 0,015828 | 135000,5 | 135997,3125 | 1409,650513 | 0 |
| BS5 | Week 8 | 24,02208328 | 23,98343 | 0,054671 | 3116417 | 3198954 | 116725,125 | 0 |
| BS5 | Week 8 | 23,944767 | 23,98343 | 0,054671 | 3281491 | 3198954 | 116725,125 | 0 |
| BS5 | Week 9 | 22,19887924 | 22,15863 | 0,056927 | 28731872 | 29504536 | 1092713,375 | 0 |
| BS5 | Week 9 | 22,11837196 | 22,15863 | 0,056927 | 30277202 | 29504536 | 1092713,375 | 0 |
| BS5 | Week 10 | 21,81987 | 21,76993 | 0,070629 | 36768344 | 38002944 | 1745990,875 | 0 |
| BS5 | Week 10 | 21,71998596 | 21,76993 | 0,070629 | 39237548 | 38002944 | 1745990,875 | 0 |
| BS6 | Week 0 (before inoculation) | Undetermined | | | | | | 0 |
| BS6 | Week 0 (before inoculation) | Undetermined | | | | | | 0 |
| BS6 | Week 1 | 35,26822281 | 35,11723 | 0,213534 | 4213,458 | 4678,242188 | 657,3044434 | 0 |
| BS6 | Week 1 | 34,96623993 | 35,11723 | 0,213534 | 5143,027 | 4678,242188 | 657,3044434 | 0 |
| BS6 | Week 2 | 37,15199661 | 37,23351 | 0,115276 | 1135,647 | 1078,667847 | 80,5812149 | 0 |
| BS6 | Week 2 | 37,31502151 | 37,23351 | 0,115276 | 1021,688 | 1078,667847 | 80,5812149 | 0 |
| BS6 | Week 3 | 28,83594322 | 28,75997 | 0,107448 | 371782,4 | 390564,3125 | 26561,69336 | 0 |
| BS6 | Week 3 | 28,68398857 | 28,75997 | 0,107448 | 409346,3 | 390564,3125 | 26561,69336 | 0 |
| BS6 | Week 4 | 30,54592133 | 30,45074 | 0,13461 | 100231 | 106977,2969 | 9540,632813 | 0 |
| BS6 | Week 4 | 30,35555458 | 30,45074 | 0,13461 | 113723,5 | 106977,2969 | 9540,632813 | 0 |
| BS6 | Week 5 | 33,4858017 | 33,41037 | 0,106682 | 17107,4 | 17995,67188 | 1256,205566 | 0 |

| | | | | | | | | |
|---|---|---|---|---|---|---|---|---|
| BS6 | Week 5 | 33,33493042 | 33,41037 | 0,106682 | 18883,94 | 17995,67188 | 1256,205566 | 0 |
| BS6 | Week 6 | 33,00737762 | 33,00314 | 0,005988 | 11427,27 | 11459,62402 | 45,74925613 | 0 |
| BS6 | Week 6 | 32,998909 | 33,00314 | 0,005988 | 11491,97 | 11459,62402 | 45,74925613 | 0 |
| BS6 | Week 7 | 32,21985245 | 32,04176 | 0,251857 | 39194,72 | 44342,92969 | 7280,661621 | 0 |
| BS6 | Week 7 | 31,86367226 | 32,04176 | 0,251857 | 49491,13 | 44342,92969 | 7280,661621 | 0 |
| BS6 | Week 8 | 32,92210007 | 32,82557 | 0,136513 | 27938,63 | 29755,76172 | 2569,819336 | 0 |
| BS6 | Week 8 | 32,72904205 | 32,82557 | 0,136513 | 31572,9 | 29755,76172 | 2569,819336 | 0 |
| BS6 | Week 9 | 29,86458588 | 29,87396 | 0,01326 | 195880,9 | 194693 | 1679,964233 | 0 |
| BS6 | Week 9 | 29,88333893 | 29,87396 | 0,01326 | 193505,1 | 194693 | 1679,964233 | 0 |
| BS6 | Week 10 | 28,00899315 | 27,99989 | 0,012879 | 655228,9 | 659134,875 | 5523,785645 | 0 |
| BS6 | Week 10 | 27,99077988 | 27,99989 | 0,012879 | 663040,8 | 659134,875 | 5523,785645 | 0 |
| CB1 | Week 0 (before inoculation) | Undetermined | | | | | | 0 |
| CB1 | Week 0 (before inoculation) | Undetermined | | | | | | 0 |
| CB1 | Week 1 | 32,45859146 | 32,3428 | 0,163762 | 14774,74 | 16076,43945 | 1840,87793 | 0 |
| CB1 | Week 1 | 32,22699738 | 32,3428 | 0,163762 | 17378,14 | 16076,43945 | 1840,87793 | 0 |
| CB1 | Week 2 | 29,5784111 | 29,46932 | 0,154275 | 190440,5 | 205270,3438 | 20972,52148 | 0 |
| CB1 | Week 2 | 29,36023331 | 29,46932 | 0,154275 | 220100,2 | 205270,3438 | 20972,52148 | 0 |
| CB1 | Week 3 | 28,96891022 | 28,98292 | 0,019807 | 285343,2 | 282716,4375 | 3714,851807 | 0 |
| CB1 | Week 3 | 28,99692154 | 28,98292 | 0,019807 | 280089,6 | 282716,4375 | 3714,851807 | 0 |

| | | | | | | | | |
|---|---|---|---|---|---|---|---|---|
| CB1 | Week 4 | 32,05082703 | 32,06387 | 0,018439 | 48516,38 | 48118,97266 | 562,0228271 | 0 |
| CB1 | Week 4 | 32,0769043 | 32,06387 | 0,018439 | 47721,56 | 48118,97266 | 562,0228271 | 0 |
| CB1 | Week 5 | 26,06640244 | 26,08336 | 0,023979 | 2204478 | 2180270,5 | 34234,92969 | 0 |
| CB1 | Week 5 | 26,10031319 | 26,08336 | 0,023979 | 2156063 | 2180270,5 | 34234,92969 | 0 |
| CB1 | Week 6 | 20,06487846 | 20,0226 | 0,059793 | 1,12E+08 | 115442720 | 4519204,5 | 1 |
| CB1 | Week 6 | 19,98031807 | 20,0226 | 0,059793 | 1,19E+08 | 115442720 | 4519204,5 | 1 |
| CB2 | Week 0 (before inoculation) | Undetermined | | | | | | 0 |
| CB2 | Week 0 (before inoculation) | Undetermined | | | | | | 0 |
| CB2 | Week 1 | 33,33405685 | 33,18739 | 0,207416 | 7999,832 | 8912,650391 | 1290,919434 | 0 |
| CB2 | Week 1 | 33,04072571 | 33,18739 | 0,207416 | 9825,468 | 8912,650391 | 1290,919434 | 0 |
| CB2 | Week 2 | 30,48740959 | 30,438 | 0,069871 | 104198,3 | 107727,9922 | 4991,765137 | 0 |
| CB2 | Week 2 | 30,38859749 | 30,438 | 0,069871 | 111257,7 | 107727,9922 | 4991,765137 | 0 |
| CB2 | Week 3 | 30,01311493 | 29,96984 | 0,061199 | 142729,4 | 146946,875 | 5964,412598 | 0 |
| CB2 | Week 3 | 29,92656708 | 29,96984 | 0,061199 | 151164,3 | 146946,875 | 5964,412598 | 0 |
| CB2 | Week 4 | 31,76364899 | 31,7564 | 0,010251 | 58195,56 | 58464 | 379,6362305 | 0 |
| CB2 | Week 4 | 31,74915123 | 31,7564 | 0,010251 | 58732,45 | 58464 | 379,6362305 | 0 |
| CB2 | Week 5 | 27,70198822 | 27,7268 | 0,035092 | 762491,2 | 750693 | 16685,20117 | 0 |
| CB2 | Week 5 | 27,75161552 | 27,7268 | 0,035092 | 738894,8 | 750693 | 16685,20117 | 0 |
| CB2 | Week 6 | 24,60109329 | 24,61499 | 0,062221 | 4819068 | 4778102,5 | 197824,1719 | 0 |

| | | | | | | | | |
|---|---|---|---|---|---|---|---|---|
| CB2 | Week 6 | 24,53364372 | 24,61499 | 0,062221 | 5038499 | 4778102,5 | 197824,1719 | 0 |
| CB2 | Week 7 | 22,20224571 | 22,12926 | 0,103209 | 27688232 | 29076824 | 1963765,625 | 1 |
| CB2 | Week 7 | 22,05628586 | 22,12926 | 0,103209 | 30465416 | 29076824 | 1963765,625 | 1 |
| CB3 | Week 0 (before inoculation) | Undetermined | | | | | | 0 |
| CB3 | Week 0 (before inoculation) | Undetermined | | | | | | 0 |
| CB3 | Week 1 | 34,17605591 | 33,93199 | 0,345162 | 4434,319 | 5338,600586 | 1278,847168 | 0 |
| CB3 | Week 1 | 33,68792343 | 33,93199 | 0,345162 | 6242,882 | 5338,600586 | 1278,847168 | 0 |
| CB3 | Week 2 | 36,70080948 | 36,51255 | 0,266244 | 1507,619 | 1719,096436 | 299,0739746 | 0 |
| CB3 | Week 2 | 36,3242836 | 36,51255 | 0,266244 | 1930,574 | 1719,096436 | 299,0739746 | 0 |
| CB3 | Week 3 | 29,99464035 | 29,9734 | 0,030033 | 117900 | 119546,6563 | 2328,729004 | 0 |
| CB3 | Week 3 | 29,95216751 | 29,9734 | 0,030033 | 121193,3 | 119546,6563 | 2328,729004 | 0 |
| CB3 | Week 4 | 29,43483543 | 29,38852 | 0,0655 | 254411,2 | 262098,2813 | 10871,20313 | 0 |
| CB3 | Week 4 | 29,34220505 | 29,38852 | 0,0655 | 269785,4 | 262098,2813 | 10871,20313 | 0 |
| CB3 | Week 5 | 26,28526306 | 26,32906 | 0,061943 | 1870552 | 1820067,5 | 71395,07031 | 0 |
| CB3 | Week 5 | 26,37286377 | 26,32906 | 0,061943 | 1769584 | 1820067,5 | 71395,07031 | 0 |
| CB3 | Week 6 | 25,81943703 | 25,6935 | 0,178104 | 2156042 | 2351060 | 275796,9375 | 0 |
| CB3 | Week 6 | 25,5675602 | 25,6935 | 0,178104 | 2546078 | 2351060 | 275796,9375 | 0 |
| CB3 | 28/07/2025 | 19,05291367 | 19,00511 | 0,067602 | 2,18E+08 | 224797856 | 9948676 | 1 |
| CB3 | 28/07/2025 | 18,95730972 | 19,00511 | 0,067602 | 2,32E+08 | 224797856 | 9948676 | 1 |

| | | | | | | | | |
|---|---|---|---|---|---|---|---|---|
| CB5 | Week 0 (before inoculation) | Undetermined | | | | | | 0 |
| CB5 | Week 0 (before inoculation) | Undetermined | | | | | | 0 |
| CB5 | Week 1 | 23,96832085 | 24,13413 | 0,234486 | 5668236 | 5080557 | 831103,25 | 0 |
| CB5 | Week 1 | 24,29993439 | 24,13413 | 0,234486 | 4492879 | 5080557 | 831103,25 | 0 |
| CB5 | Week 2 | 27,84185028 | 27,8875 | 0,064564 | 602689,8 | 584977,875 | 25048,41797 | 0 |
| CB5 | Week 2 | 27,93315697 | 27,8875 | 0,064564 | 567266 | 584977,875 | 25048,41797 | 0 |
| CB5 | Week 3 | 23,29094505 | 23,16729 | 0,174868 | 9119795 | 9913194 | 1122034,875 | 0 |
| CB5 | Week 3 | 23,04364395 | 23,16729 | 0,174868 | 10706592 | 9913194 | 1122034,875 | 0 |
| CB5 | Week 4 | 24,55656815 | 24,51622 | 0,057062 | 5591471 | 5738094,5 | 207357,6563 | 0 |
| CB5 | Week 4 | 24,47587013 | 24,51622 | 0,057062 | 5884719 | 5738094,5 | 207357,6563 | 0 |
| CB5 | Week 5 | 17,91008759 | 17,9353 | 0,035654 | 2,69E+08 | 264189664 | 6279187,5 | 1 |
| CB5 | Week 5 | 17,96051025 | 17,9353 | 0,035654 | 2,6E+08 | 264189664 | 6279187,5 | 1 |
| CB6 | Week 0 (before inoculation) | Undetermined | | | | | | 0 |
| CB6 | Week 0 (before inoculation) | Undetermined | | | | | | 0 |
| CB6 | Week 1 | 27,00330162 | 26,96324 | 0,056653 | 675744 | 695256,6875 | 27595,10742 | 0 |
| CB6 | Week 1 | 26,92318153 | 26,96324 | 0,056653 | 714769,4 | 695256,6875 | 27595,10742 | 0 |
| CB6 | Week 2 | 26,05195427 | 26,05728 | 0,007537 | 2168462 | 2161167 | 10317,04102 | 0 |
| CB6 | Week 2 | 26,06261253 | 26,05728 | 0,007537 | 2153872 | 2161167 | 10317,04102 | 0 |
| CB6 | Week 3 | 22,4307003 | 22,41149 | 0,027164 | 15933866 | 16134885 | 284283,7813 | 0 |

| | | | | | | | | |
|---|---|---|---|---|---|---|---|---|
| CB6 | Week 3 | 22,39228439 | 22,41149 | 0,027164 | 16335904 | 16134885 | 284283,7813 | 0 |
| CB6 | Week 4 | 24,23659706 | 24,29351 | 0,080493 | 6847762 | 6609570 | 336854,7188 | 0 |
| CB6 | Week 4 | 24,35043144 | 24,29351 | 0,080493 | 6371378 | 6609570 | 336854,7188 | 0 |
| CB6 | Week 5 | 19,64338303 | 19,68419 | 0,057709 | 1,26E+08 | 122473160 | 4476008,5 | 1 |
| CB6 | Week 5 | 19,72499657 | 19,68419 | 0,057709 | 1,19E+08 | 122473160 | 4476008,5 | 1 |
| CB7 | Week 0 (before inoculation) | Undetermined | | | | | | 0 |
| CB7 | Week 0 (before inoculation) | Undetermined | | | | | | 0 |
| CB7 | Week 1 | 28,76430511 | 28,70966 | 0,077286 | 196715,5 | 204545,0625 | 11072,64063 | 0 |
| CB7 | Week 1 | 28,65500641 | 28,70966 | 0,077286 | 212374,6 | 204545,0625 | 11072,64063 | 0 |
| CB7 | Week 2 | 28,48572922 | 28,51944 | 0,047677 | 313749,6 | 307036,5 | 9493,726563 | 0 |
| CB7 | Week 2 | 28,55315399 | 28,51944 | 0,047677 | 300323,4 | 307036,5 | 9493,726563 | 0 |
| CB7 | Week 3 | 25,40340996 | 25,33787 | 0,092692 | 2316813 | 2419620,5 | 145391,9375 | 0 |
| CB7 | Week 3 | 25,27232361 | 25,33787 | 0,092692 | 2522428 | 2419620,5 | 145391,9375 | 0 |
| CB7 | Week 4 | 25,93728447 | 26,05656 | 0,168684 | 2398986 | 2225501,75 | 245344,125 | 0 |
| CB7 | Week 4 | 26,17584038 | 26,05656 | 0,168684 | 2052017 | 2225501,75 | 245344,125 | 0 |
| CB7 | Week 5 | 26,90460587 | 26,94304 | 0,054347 | 1263546 | 1233525,5 | 42455,92969 | 0 |
| CB7 | Week 5 | 26,98146439 | 26,94304 | 0,054347 | 1203505 | 1233525,5 | 42455,92969 | 0 |
| CB7 | Week 6 | 24,08247375 | 24,12265 | 0,056821 | 6786629 | 6611309,5 | 247938,5 | 0 |
| CB7 | Week 6 | 24,16283035 | 24,12265 | 0,056821 | 6435991 | 6611309,5 | 247938,5 | 0 |

| | | | | | | | | |
|---|---|---|---|---|---|---|---|---|
| CB7 | Week 7 | 21,12580299 | 21,1258 | | 47791180 | 47791180 | | 1 |
| CB7 | Week 7 | 21,3266983 | 21,3267 | | 49125412 | 49125412 | | 1 |

**Table S8.** Contribution metrics from five exploratory replicate models for the environmental niche of *Bd* presence and including all 66 CHELSA variables excluding cloud cover variables (cltmax, cltmean, cltmin, cltrange, cmimax, cmimean, cmimin and cmirange) and Köppen-Geiger Climate Classifications (kg0, kg1, kg2, kg3, kg4 and kg5).

| **Variable** | **Percent contribution** | **Permutation importance** |
|---|---|---|
| CHELSA_ai_1981-2010_V.2.1 | 0 | 0 |
| CHELSA_bio10_1981-2010_V.2.1 | 0.9 | 0 |
| CHELSA_bio11_1981-2010_V.2.1 | 0.2 | 0 |
| CHELSA_bio12_1981-2010_V.2.1 | 0.4 | 0.3 |
| CHELSA_bio13_1981-2010_V.2.1 | 0 | 0 |
| CHELSA_bio14_1981-2010_V.2.1 | 0.1 | 0.3 |
| CHELSA_bio15_1981-2010_V.2.1 | 0.3 | 0.7 |
| CHELSA_bio16_1981-2010_V.2.1 | 0 | 0 |
| CHELSA_bio17_1981-2010_V.2.1 | 0 | 0 |
| CHELSA_bio18_1981-2010_V.2.1 | 0 | 0.1 |
| CHELSA_bio19_1981-2010_V.2.1 | 0.2 | 0.1 |
| CHELSA_bio1_1981-2010_V.2.1 | 1.1 | 0 |
| CHELSA_bio2_1981-2010_V.2.1 | 0.3 | 0.7 |
| CHELSA_bio3_1981-2010_V.2.1 | 0.8 | 0.4 |
| CHELSA_bio4_1981-2010_V.2.1 | 1 | 1.4 |
| CHELSA_bio5_1981-2010_V.2.1 | 1.1 | 0 |
| CHELSA_bio6_1981-2010_V.2.1 | 0.5 | 0.1 |
| CHELSA_bio7_1981-2010_V.2.1 | 0.2 | 0.5 |
| CHELSA_bio8_1981-2010_V.2.1 | 0.4 | 0.2 |
| CHELSA_bio9_1981-2010_V.2.1 | 0.7 | 0.1 |
| CHELSA_cmi_max_1981-2010_V.2.1 | 0 | 0 |
| CHELSA_cmi_mean_1981-2010_V.2.1 | 0.7 | 0 |
| CHELSA_cmi_min_1981-2010_V.2.1 | 0 | 0 |
| CHELSA_cmi_range_1981-2010_V.2.1 | 0.1 | 0.1 |
| CHELSA_fcf_1981-2010_V.2.1 | 0.6 | 0.2 |
| CHELSA_fgd_1981-2010_V.2.1 | 0.2 | 0.1 |
| CHELSA_gdd0_1981-2010_V.2.1 | 13.9 | 1.4 |
| CHELSA_gdd10_1981-2010_V.2.1 | 3.6 | 0.9 |
| CHELSA_gdd5_1981-2010_V.2.1 | 0.5 | 0.3 |
| CHELSA_gddlgd0_1981-2010_V.2.1 | 1.3 | 0.7 |

| | | |
|---|---|---|
| CHELSA_gddlgd10_1981-2010_V.2.1 | 4.3 | 1.7 |
| CHELSA_gddlgd5_1981-2010_V.2.1 | 1 | 0.8 |
| CHELSA_gdgfgd0_1981-2010_V.2.1 | 0.4 | 0.3 |
| CHELSA_gdgfgd10_1981-2010_V.2.1 | 0.7 | 0.5 |
| CHELSA_gdgfgd5_1981-2010_V.2.1 | 0 | 0.1 |
| CHELSA_gsl_1981-2010_V.2.1 | 17.7 | 4.4 |
| CHELSA_gsp_1981-2010_V.2.1 | 0.4 | 0.4 |
| CHELSA_gst_1981-2010_V.2.1 | 7.7 | 0 |
| CHELSA_hurs_max_1981-2010_V.2.1 | 0.5 | 1.2 |
| CHELSA_hurs_mean_1981-2010_V.2.1 | 0.7 | 0.4 |
| CHELSA_hurs_min_1981-2010_V.2.1 | 0.3 | 0.6 |
| CHELSA_hurs_range_1981-2010_V.2.1 | 2.4 | 0.6 |
| CHELSA_lgd_1981-2010_V.2.1 | 0.8 | 0.8 |
| CHELSA_ngd0_1981-2010_V.2.1 | 3.4 | 7 |
| CHELSA_ngd10_1981-2010_V.2.1 | 0.3 | 2.5 |
| CHELSA_ngd5_1981-2010_V.2.1 | 0.6 | 0.1 |
| CHELSA_npp_1981-2010_V.2.1 | 1.7 | 6.1 |
| CHELSA_pet_penman_max_1981-2010_V.2.1 | 0.2 | 5.9 |
| CHELSA_pet_penman_mean_1981-2010_V.2.1 | 0 | 0.1 |
| CHELSA_pet_penman_min_1981-2010_V.2.1 | 0.8 | 19.1 |
| CHELSA_pet_penman_range_1981-2010_V.2.1 | 0.3 | 0.1 |
| CHELSA_rsds_1981-2010_max_V.2.1 | 2.7 | 2.3 |
| CHELSA_rsds_1981-2010_mean_V.2.1 | 0.7 | 1.1 |
| CHELSA_rsds_1981-2010_min_V.2.1 | 1.2 | 6.2 |
| CHELSA_rsds_1981-2010_range_V.2.1 | 2.9 | 5.3 |
| CHELSA_scd_1981-2010_V.2.1 | 3.2 | 6.6 |
| CHELSA_sfcWind_max_1981-2010_V.2.1 | 0 | 0 |
| CHELSA_sfcWind_mean_1981-2010_V.2.1 | 0.6 | 0.1 |
| CHELSA_sfcWind_min_1981-2010_V.2.1 | 0.3 | 0.9 |

| | | |
|---|---|---|
| CHELSA_sfcWind_range_1981-2010_V.2.1 | 4.1 | 0 |
| CHELSA_swb_1981-2010_V.2.1 | 4.4 | 3.8 |
| CHELSA_swe_1981-2010_V.2.1 | 1.8 | 2 |
| CHELSA_vpd_max_1981-2010_V.2.1 | 2.2 | 4.3 |
| CHELSA_vpd_mean_1981-2010_V.2.1 | 0.4 | 1.6 |
| CHELSA_vpd_min_1981-2010_V.2.1 | 2.1 | 3.6 |
| CHELSA_vpd_range_1981-2010_V.2.1 | 0.2 | 1 |

**Table S9.** Threshold values from ten replicate models and the average model for environmental niche of *Bd* presence (*Bd* presence model; above) and for the climatic envelope of *Bd*-induced amphibian declines (*Bd*-induced decline model; below).

| ***Bd* presence model** | **Model 1** | **Model 2** | **Model 3** | **Model 4** | **Model 5** | **Model 6** | **Model 7** | **Model 8** | **Model 9** | **Model 10** | **Average** |
|---|---|---|---|---|---|---|---|---|---|---|---|
| $10^{th}$ percentile training presence | 0.3065 | 0.2951 | 0.2981 | 0.3038 | 0.2984 | 0.3097 | 0.2966 | 0.2987 | 0.3022 | 0.2989 | 0.3008 |
| maximum training sensitivity plus specificity | 0.2765 | 0.2526 | 0.2605 | 0.2609 | 0.2666 | 0.2695 | 0.2457 | 0.2529 | 0.2692 | 0.2745 | 0.2629 |
| maximum test sensitivity plus specificity. | 0.2239 | 0.2785 | 0.2206 | 0.2702 | 0.2866 | 0.2316 | 0.2025 | 0.3118 | 0.2605 | 0.2506 | 0.2537 |
| ***Bd-induced decline model*** | **Model 1** | **Model 2** | **Model 3** | **Model 4** | **Model 5** | **Model 6** | **Model 7** | **Model 8** | **Model 9** | **Model 10** | **Average** |
| $10^{th}$ percentile training presence | 0.1176 | 0.1277 | 0.1439 | 0.1087 | 0.1163 | 0.1003 | 0.0909 | 0.1148 | 0.0925 | 0.0957 | 0.1108 |
| maximum training sensitivity plus specificity | 0.1585 | 0.0761 | 0.1350 | 0.0718 | 0.1265 | 0.0553 | 0.0495 | 0.1249 | 0.0608 | 0.0498 | 0.0908 |
| maximum test sensitivity plus specificity. | 0.0328 | 0.0657 | 0.0585 | 0.1380 | 0.0229 | 0.1410 | 0.0592 | 0.0276 | 0.0615 | 0.0669 | 0.0674 |
